# Single cell transcriptomic analyses reveal the impact of bHLH factors ATOH7 and Neurog2 on human retinal organoid development

**DOI:** 10.1101/2020.10.27.358135

**Authors:** Xiangmei Zhang, Igor Mandric, Kevin H. Nguyen, Thao T. T. Nguyen, Matteo Pellegrini, James C. R. Grove, Steven Barnes, Xian-Jie Yang

## Abstract

The developing retina expresses multiple bHLH transcription factors. Their precise functions and interactions in uncommitted retinal progenitors remain to be fully elucidated. Here, we investigate the roles of bHLH factors ATOH7 and Neurog2 in developing human ES cell-derived 3D retinal organoids. Single cell transcriptome analyses identify three states of proliferating retinal progenitors: pre-neurogenic, neurogenic, and cell cycle-exiting progenitors. Each shows different expression profile of bHLH factors. The distinct cell cycle-exiting progenitors feed into a postmitotic heterozygous neuroblast pool that gives rise to early born neuronal lineages. Elevating ATOH7 or Neurog2 expression accelerates the transition from the pre-neurogenic to the neurogenic state, and expands the exiting progenitor and neuroblast populations. In addition, ATOH7 and Neurog2 significantly, yet differentially, enhance retinal ganglion cell and cone photoreceptor production. Moreover, single cell transcriptome analyses reveal that ATOH7 and Neurog2 assert positive autoregulation, suppress key bHLH factors associated with the neurogenic progenitors, and elevate bHLH factors expressed by exiting progenitors and differentiating neuroblasts. This study thus provides novel insight regarding how ATOH7 and Neurog2 impact human retinal progenitor behaviors and neuroblast fate choices.

## Introduction

As an integral component of the central nervous system, the vertebrate neural retina retains a highly conserved laminar structure that senses, processes, and delivers visual information to the brain. Classic cell birth dating and lineage tracing studies have established that the seven major neuronal cell types constituting the retinal network are generated in a temporal order from a common ocular progenitor pool during development ^1–3^. The ensuing research has ruled out a rigid deterministic cell fate specification mechanism, but instead supports a view that multipotent progenitors progressively evolve through different competence states to enable the sequential production of distinct cell types ^1, 4^. Cumulative molecular genetic studies have uncovered important roles of cell intrinsic factors involved in retinal development ^5^. Among these, transcription factors containing the basic helix-loop-helix (bHLH) motif have emerged as important players regulating the production and differentiation of various retinal cell types ^6^. Multiple bHLH factors are expressed during retinal development either in proliferating progenitors or in postmitotic neurons; however, their dynamic regulation and function in specific cellular contexts remain to be fully elucidated.

The bHLH factors Atoh7 and Neurog2 are both expressed early in the developing retinal epithelium. Atoh7 plays a critical role in the development of an early born retinal neuronal type, the retinal ganglion cells (RGCs), which project axons through the optic nerve to multiple higher visual centers ^2, 7^. Both Atoh7 and Neurog2 mRNAs are expressed by subsets of early retinal progenitors, with some co-expression at the protein level during certain time windows ^8–10^. Loss of Atoh7 function in mouse, zebrafish, and humans results in a severe reduction of RGCs leading to a diminished optic nerve and blindness ^11–15^. Atoh7 deficiency also causes a minor abnormality in the production of cone photoreceptors, another early-born cell type in the retina ^11^. In contrast, genetic ablation of Neurog2 yields a transient stall of neurogenesis but without severe lasting deficits ^16^. In the mouse retina, Atoh7 protein is not detected in fully differentiated RGCs ^9, 10^, suggesting that its main biological activity is transiently required in uncommitted early progenitors. Ectopic expression of Atoh7 in different late stage retinal progenitors either redirects progenitors towards an RGC fate ^17^ or fails to specify the RGC fate ^18^. Therefore, Atoh7 is thought to confer a competent state of progenitors to adopt early cell fates^19^. In the absence of Atoh7, co-expression of two downstream transcription factors Islet1 and Pou4f2 is sufficient to rescue the RGC production deficit and ensure full execution of the RGC differentiation program in the mouse retina ^20–24^. We have shown previously that viral mediated expression of human ATOH7 in the developing chicken retina results in precocious neurogenesis and a significant increase in RGC production ^25^, supporting a hypothesis that a critically high threshold of ATOH7 expression triggers uncommitted early progenitors to exit the cell cycle and predominantly adopt the RGC fate ^26^.

In the human retina, RGC development occurs during the first trimester and remains a minor cell population in the mature retina ^27–29^. This scarcity of human RGCs has hindered research on human RGC development as well as blinding diseases caused by RGC loss. The advancements of pluripotent stem cell technologies in the preceding decade have led to robust stem cell based retinal organoid culture systems ^30–33^, thus providing an excellent opportunity to produce and study human RGC development in vitro. Recent single cell transcriptomic analyses have revealed that primate retinas, including humans, show distinct molecular features and RGC subtype proportions compared to rodents ^34–42^. In this study, we have examined the function of bHLH factors ATOH7 and Neurog2 on human RGC development using embryonic stem cell (ESC)-derived 3D retinal organoids. Our results demonstrate that elevating these two factors in uncommitted human retinal progenitors assert powerful neurogenic effects. By performing single cell transcriptomic analysis, we identify distinct statuses of human retinal progenitors, including a population poised to exit the cell cycle. Our results show that ATOH7 and Neurog2 participate and regulate an interactive gene network to accelerate progenitors through two transitional stages to adopt postmitotic neuronal identities.

## Results

### Viral mediated bHLH factor expression affects progenitor proliferation in 3D human retinal organoids

To investigate the roles of bHLH neurogenic factors during development of the human retina, we established H9 embryonic stem cell (ESC)-derived 3D retinal organoid cultures ^30, 43^. These human retinal organoids showed typical morphology of the retinal neural epithelium and co-expressed PAX6 and VSX2 transcription factors (Supplementary Fig. 1), a characteristic feature specific to retinal progenitors. To regulate gene expression during retinogenesis, we constructed lentiviral vectors that encode the doxycycline (Dox) inducible TetO promoter upstream of the human ATOH7 cDNA fused with the Flag epitope tag (LV-ATOH7f) or co-expressing EGFP and puromycin resistant genes (LV-AEP) (Fig. 1a). We also produced a previously described lentiviral TetO vector co-expressing the mouse Neurog2, EGFP, and puromycin resistant genes (LV-NEP) ^44^(Fig. 1a). Immunohistochemistry confirmed Dox-induced bHLH protein expression following in vitro co-infections of H9 ESCs with LV-rtTA and either LV-ATOH7f, LV-AEP, or LV-NEP (Fig. 1b). We next carried out co-infection of retinal organoids with LV-rtTA and either LV-AEP or LV-NEP at the onset of retinogenesis followed by Dox inductions (Fig. 1c). Live imaging of the EGFP reporter showed that at 24 hours post Dox induction, significant numbers of LV-AEP or LV-NEP infected cells had already migrated towards the inner retina compared to cells from the control LV-GFP infected organoids. This trend continued and became more obvious as the Dox induction times lengthened (Supplementary Fig. 2). By 6-days after Dox induction, the majority of LV-AEP and LV-NEP infected cells were located in the inner layer of the organoids (Fig. 1d, 1e), indicating that viral mediated ATOH7 and Neurog2 expression impacted retinal organoid development.

**Fig. 1.**
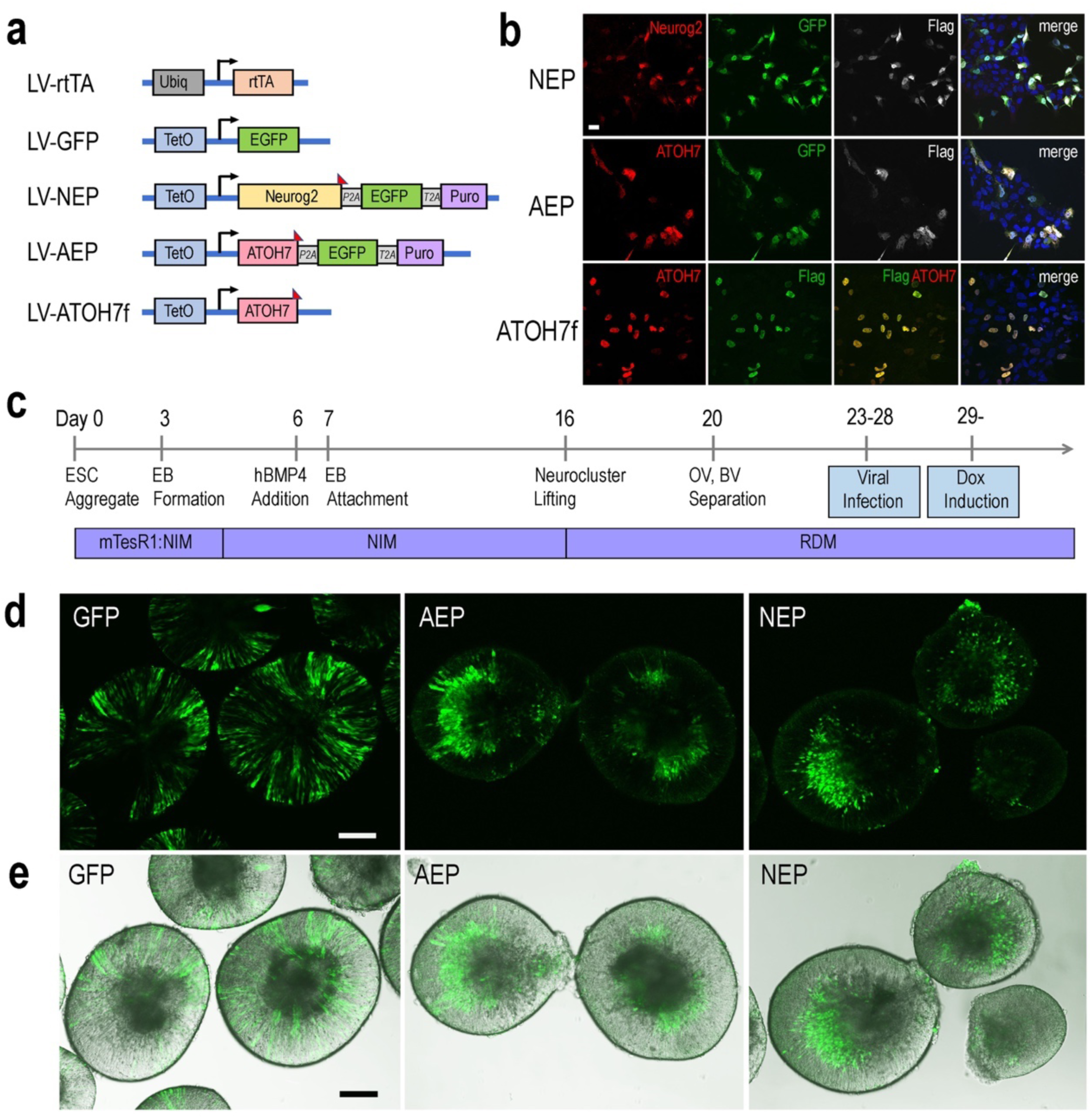
Inducible lentivirus-mediated gene expression in human ES cell-derived retinal organoids. **a** Schematics of lentiviral vectors used in the study. A vector expressing rtTA under the control of the constitutive Ubiq promoter was used to co-infect human ES cells with lentiviral vectors encoding the inducible TetO promoter. Puro, puromycin-resistant gene; *P2A* and *T2A*, intervening 2T sequences in frame with the coding sequences of the other genes in LV-NEP and LV-AEP. The Flag epitope tag was fused to the c-terminus of Neurog2 and ATOH7 coding regions. **b** Expression of Neurog2 and ATOH7 proteins as detected by immunofluorescent microscopy in H9 ES cells co-infected with LV-rtTA and either LV-NEP, LV-AEP, or LV-ATOH7f after 36 hours of Dox induction. Scale Bar for all, 20μm. **c** Experimental time course of human retinal organoid derivation and lentiviral infection. **d, e** Confocal images (**d**) and merged confocal and bright field images (**e**) show distribution of viral infected GFP^+^ cells in live retinal organoids at Day 36 after 6 days of Dox induction. Names of lentiviruses are abbreviated as: GFP for LV-GFP, NEP for LV-NEP, AEP for LV-AEP. Scale Bars, 100 μm.

To investigate whether virally expressed bHLH factors affected retinal progenitor proliferation, we performed BrdU pulse-labeling and examined progenitor marker expression by immunohistochemistry. In the control LV-GFP infected retinal organoids, BrdU-labeled and the proliferating cell nuclear antigen (PCNA)-labeled progenitors occupied the ventricular zone, whereas phospho-histone 3 (PH3) labeled M phase cells were detected at the ventricular surface of the organoids (Fig. 2a, 2b, 2c). Furthermore, PCNA showed co-labeling with PAX6-expressing progenitors in the ventricular zone, but was absent from the PAX6^+^ cells located in the inner layer, where postmitotic POU4F^+^ RGCs resided (Fig. 2b, 2c). In contrast to LV-GFP infected retinal organoids, in which GFP^+^ cells were distributed throughout the neural epithelium, the majority of GFP^+^ cells in LV-AEP and LV-NEP infected organoids were located in the inner retinal layer and devoid of co-labeling with BrdU or PCNA (Fig. 2d, 2e). Quantification of dissociated retinal organoids confirmed that percentages of GFP^+^PCNA^+^ double labeled cells were reduced significantly from 52.7 ± 8.0 % for LV-GFP infection to 9.0 ± 3.8 % and 2.0 ± 1.9 % for LV-AEP and LV-NEP infections, respectively (Fig. 2f). These results demonstrate that viral driven ATOH7 or Neurog2 expression promoted cell cycle exit.

**Fig. 2.**
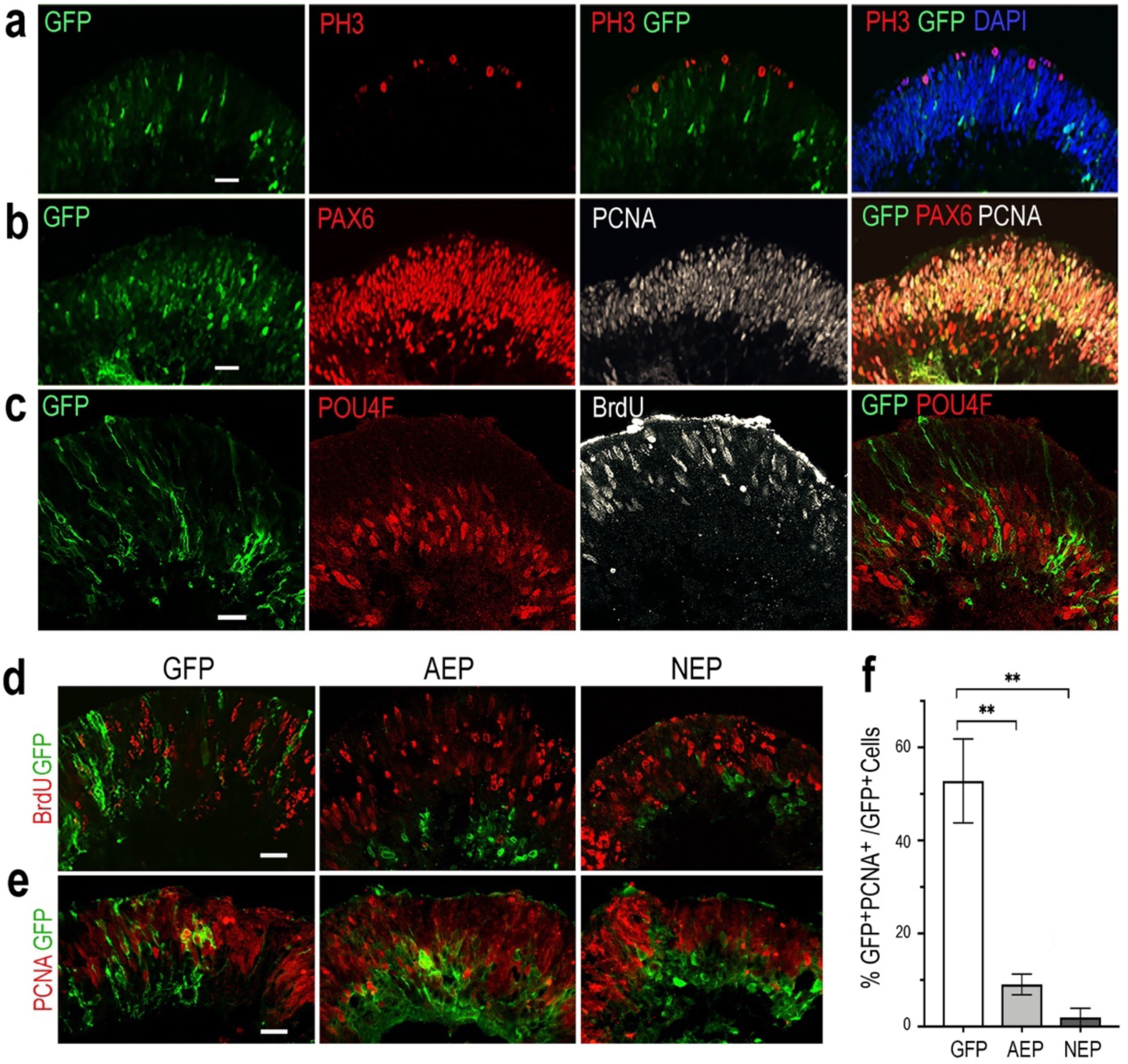
Influences of viral mediated neurogenic factor expression on cell proliferation. **a-c** Immunofluorescence labeling of cross sections from retinal organoids infected with LV-GFP. At Day 34, PH3^+^ dividing cells are located at the ventricular surface (**a**), while PCNA^+^ progenitors occupy the ventricular zone and are co-labeled with PAX6-expressing progenitors (**b**). Note that PAX6^+^PCNA**^-^** postmitotic neurons are located to the inner layer of the retinal organoid. At Day 40, proliferating progenitors labeled with BrdU are distributed in the ventricular zone, whereas POU4F^+^ postmitotic retinal ganglion cells (RGCs) reside in the inner layer of the retinal organoid (**c**). Scale bars, 20 μm. **d, e** Confocal images of cross sections from retinal organoids infected by LV-GFP, LV-AEP, or LV-NEP after 6- day Dox induction. Co-staining of GFP with BrdU at Day 36 (**d**, 3-hour labeling) or PCNA at Day 40 (**e**) shows that most LV-AEP and LV-NEP infected cells are postmitotic. Scale bars, 20 μm. **f** Quantification of double labeled PCNA^+^GFP^+^ progenitor cells among total viral infected GFP^+^ cells at Day 47. Mean ± s.e.m. of n= 3 independent samples. One-way ANOVA, ** p< 0.01.

### Elevated neurogenic factor expression promotes RGC production in 3D retinal organoids

The retinal projection neurons (RGCs) are among the earliest neuronal cells produced during retinogenesis ^2, 28^. In our human retinal organoid cultures, RGC genesis was detected as early as Day 25 and continued through Day 60 (Fig. 2c). By Day 40, retinal organoid-derived neurons exhibited voltage-gated Na^+^, K^+^ and Ca^2+^ channels as well as spontaneous and provoked electrophysiological excitability in vitro during whole cell patch clamp recording, characteristic of native RGCs (Supplementary Fig. 3). TTX-sensitive Na^+^ channels, TEA-sensitive K^+^ channels, and Cd^2+^-sensitive Ca^2+^ channel currents were recorded. Average Na^+^ current amplitude was 219 ± 76 pA (n=12) but in cells with large Na^+^ currents (∼1 nA), multiple action potentials were observed, while cells expressing smaller Na^+^ currents (200-400 pA) typically produced a single spike. Average K^+^ current amplitude at +40 mV was 438 ± 49 pA (n=40).

To determine whether viral mediated ATOH7 or Neurog2 expression promoted RGC production, we performed immunohistochemical analyses of retinal organoid sections using known RGC markers (Fig. 3). In LV-ATOH7f infected retinal organoids, signals of the viral reporter Flag closely correlated with POU4F-expressing RGCs in the inner retina as detected by a pan-POU4F/BRN3 antibody (Fig. 3a). Compared to control LV-GFP infected organoids, both LV-AEP and LV-NEP infected cells showed increased co-labeling with the RGC markers NF145, NeuN, and DCX in the inner retina (Fig. 3b). Similarly, in attached organoid cultures that displayed extensive neurite outgrowth, the ATOH7f- expressing cells showed extensive co-labeling for the RGC markers POU4F, NF145, and RBPMS (Fig. 4a).

**Fig. 3.**
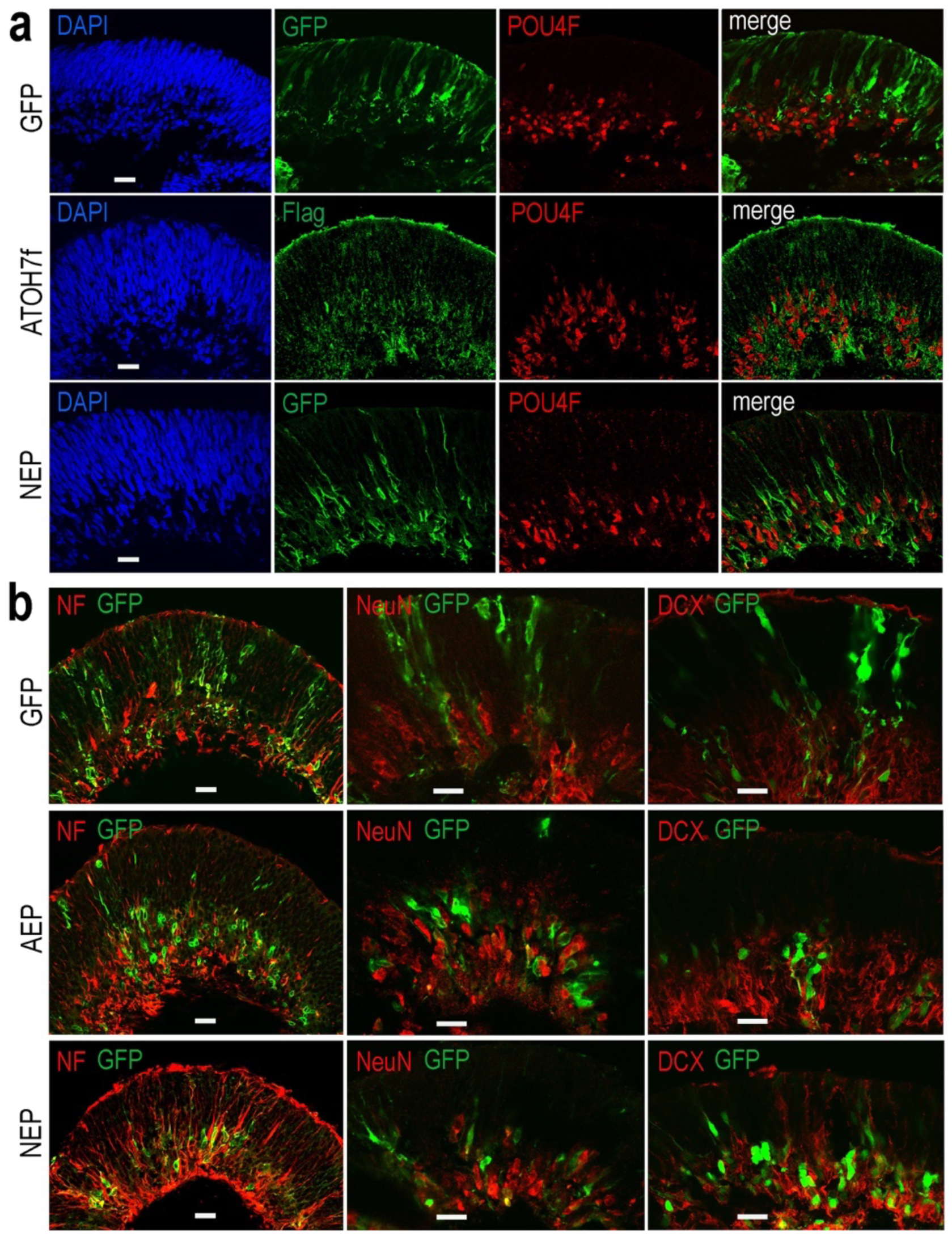
Effects of viral mediated neurogenic factor expression on retinal ganglion cell genesis. Confocal images of retinal organoid sections immunolabeled with retinal ganglion cell (RGC) markers. **a** Co-labeling of viral markers with a pan-POU4F antibody in LV-NEP and LV-ATOH7f infected retinal organoids at Day 43. **b** Merged images of viral marker GFP co-labeling with NF145 and NeuN at Day 36, and DCX at Day 46. Scale Bars, 20 μm.

**Fig. 4.**
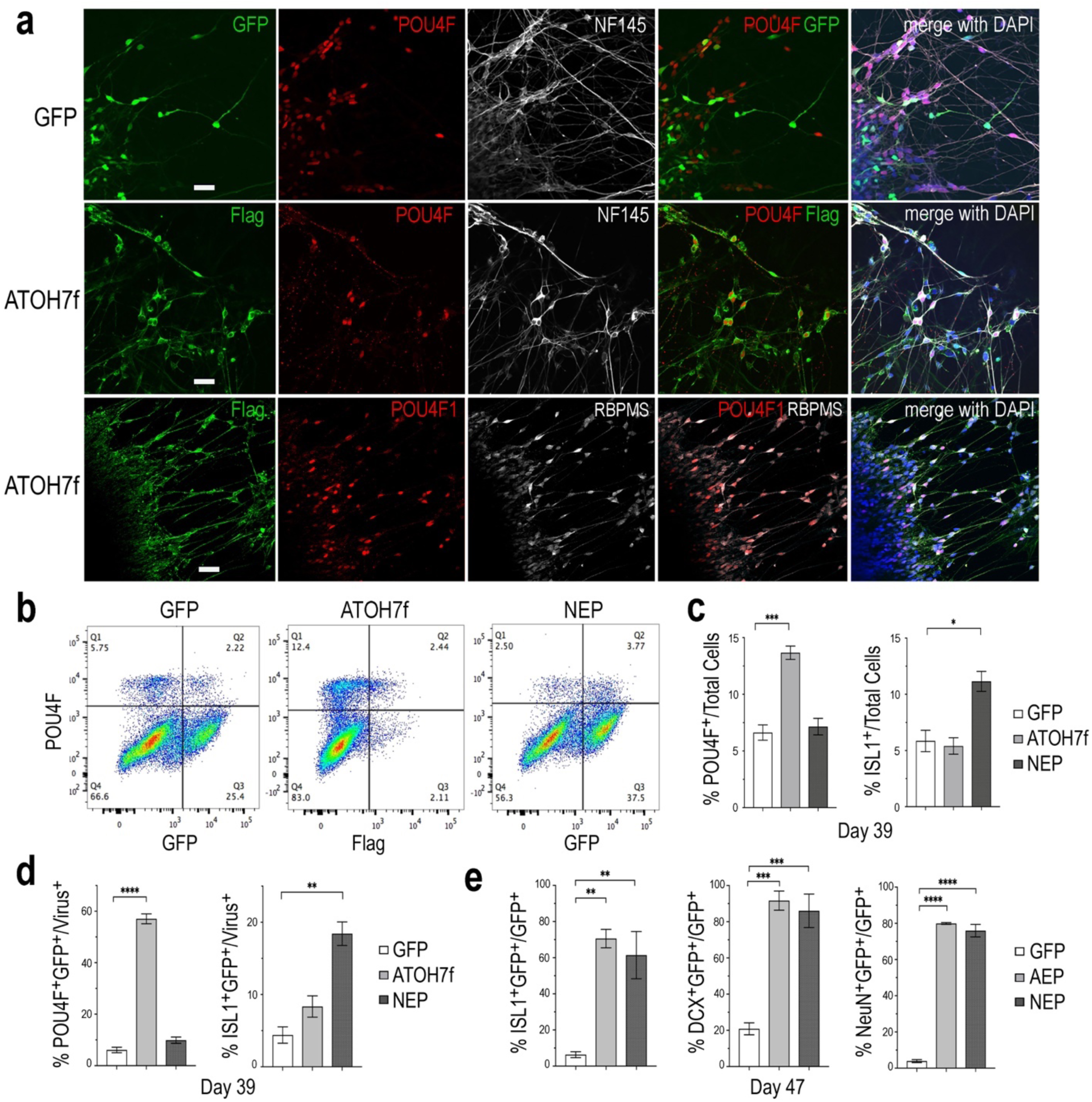
Impacts of elevating neurogenic factor expression on retinal ganglion cell genesis. **a** Immunofluorescent images of attached retinal organoid cultures infected with LV-GFP or LV-ATOH7f after 4-day Dox induction at Day 39. Co-labeling for viral vector markers with POU4F and NF145 at Day 38, or POU4F1 and RBPMS at Day 41 shows high correspondence of LV-ATOH7f infection and RGC marker expression. Scale bars, 20 μm. **b** Representative flow cytometry profiles of dissociated cells at Day 39 from LV-GFP, LV-ATOH7f, and LV-NEP infected retinal organoids after 4-day Dox induction. Cells were co-labeled for RGC marker POU4F and viral marker Flag for LV-ATOH7f, or GFP for LV-GFP and LV-NEP, respectively. **c, d** Bar graphs show flow cytometry quantification of RGC markers POU4F and ISL1 among total cells (**c**) and viral infected cells (**d**) at Day 39. **e** Quantification of cultured monolayer cells from LV-GFP, LV-AEP, and LV-NEP infected retinal organoids at Day 47 after 7-day Dox induction. Percentages of RGC marker ISL1, DCX, and NeuN positive cells among viral infected GFP^+^ cells are shown. For **c, d, e** Mean ± s.e.m. of n= 3 independent samples. One-way ANOVA, **** p< 0.0001, *** p< 0.001, **p< 0.01.

We also performed flow cytometry analyses of dissociated retinal organoid cells to quantify the effects of viral mediated ATOH7 or Neurog2 expression on RGC genesis (Fig. 4b, 4c, 4d). After 4 days of Dox induction at Day 39, ATOH7 expression led to a 2-fold increase of POU4F^+^ cells, from 6.6 ± 0.68 % of total cells in LV-GFP infected organoids to 13.7 ± 0.59 % of total cells in LV-ATOH7f infected organoids (Fig. 4c). Among LV-ATOH7f infected cells, 57.1 ± 1.9 % were POU4F^+^ compared to 6.1 ± 1.1 % among LV-GFP infected cells (Fig. 4d). However, LV-ATOH7f did not increase expression of the RGC marker ISLET1 at this stage (Fig. 4c). In the parallel analysis, LV-NEP infection did not significantly promote POU4F^+^ cells, but instead increased ISLET^+^ cells from 5.9 ± 0.95 % to 11.1 ± 0.89 % of total cells (Fig. 4c), and from 4.4 ± 1.1 % to 18.4 ± 1.6 % among viral infected cells (Fig. 4d). Similar analyses at Day 47 following a 7-day Dox induction showed significantly increased co-labeling with the RGC markers ISLET1 (> 10-fold of 6.3%), DCX (>4-fold of 21%), and NeuN (>20-fold of 4%) of LV-AEP and LV-NEP infected cells compared with the control LV-GFP infected cells (Fig. 4e). These results demonstrate that elevated ATOH7 or Neurog2 expression promoted human RGC production in retinal organoids.

### Single cell RNA-sequencing analysis reveals accelerated neurogenesis

To elucidate the influences asserted by viral driven ATOH7 and Neurog2 expression on transcriptome, we performed single cell RNA-sequencing (sc RNA-seq) analysis. We first used fluorescent activated cell sorting to enrich for LV-GFP, LV-AEP, and LV-NEP infected retinal organoid cells between Day 45 to Day 48 (Supplementary Fig.4), followed by 10X Genomics automated single-cell capture, mRNA barcoding, and cDNA library preparation. The high throughput DNA sequencing resulted in 192,932, 153,154, and 132,651 mean readings per cell for LV-GFP, LV-AEP, and LV-NEP samples, respectively. After aligning to the reference human genome and eliminating poor quality cells, the final single cell datasets had a mean gene range between 2935-3079 per cell, and consisted of 3004 cells for LV-GFP, 2063 cells for LV-AEP, and 3909 cells for LV-NEP infected samples.

The sc RNA-seq datasets were subjected to Seurat cell clustering analysis ^45^, which resolved into 12-15 clusters, and visualized as 2-dimensional UMAPs (Fig. 5a). We applied dot plot analysis using known genes to assign various cell clusters into seven categories or states; each was represented with one or two highly or uniquely expressed genes (Fig. 5b; Supplementary Fig. 5). The pre-neurogenic progenitor (PNP), neurogenic progenitor (NP), and cell cycle-exiting progenitor (EP) categories all expressed CCDN1, encoding cyclin D1, indicating that they all belonged to proliferative cells. However, the pre-neurogenic progenitor cluster cells showed distinctively high levels of PLOD2 transcripts (procollagen-lysine,2-oxoglutarate 5-dioxygenase 2), whereas the exiting progenitor cells exhibited upregulation of GADD45A (growth arrest and DNA damage inducible alpha) and downregulation of the retinal progenitor marker VSX2. The exiting progenitor cells and postmitotic neuroblasts (NB) were the main populations expressing ATOH7 mRNA (see Fig. 9). Markers identifying the early born retinal neuronal types included ISL1 and POU4F2 for RGCs, CRX and RCVRN for photoreceptors, and PRDM13 and TFAP2A for horizontal and amacrine cells (HC/AC) (Fig. 5b; Supplementary Fig. 5). These sc RNA-seq analyses thus not only allowed identification of the early postmitotic neuronal types in the human retinal organoids at the time of analysis, but also revealed sub-classes of neural progenitors with distinct molecular signatures, and transitional states including cell cycle exiting progenitors and postmitotic neuroblasts.

**Fig. 5.**
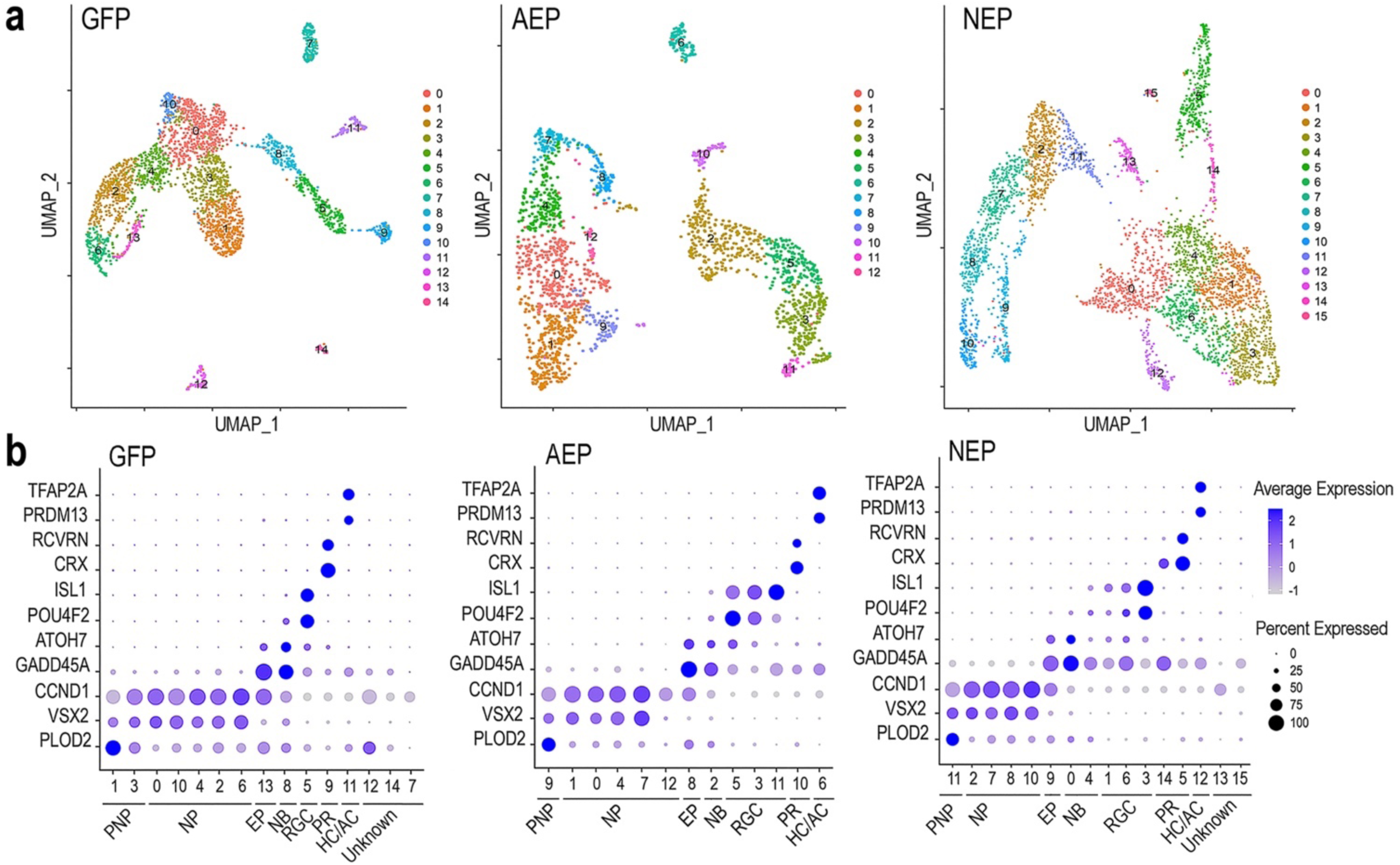
Characterization of lentivirus-infected human retinal organoids by single cell RNA-seq. **a** UMAP visualization of cell clustering based on sequencing of single cell cDNA libraries from lentivirus infected retinal organoids between culture Day 45-48 after 8-day Dox induction. The numbers of single cell that passed quality control and used for downstream analysis are 3004 for LV-GFP, 2063 for LV-AEP, and 3909 for LV-NEP samples. **b** Dot plots show assignments of UMAP clusters into seven major cell categories based on expression of known genes for different cell states or types during early retinal development. PNP, pre-neurogenic progenitor; NP, neurogenic progenitor; EP, cell cycle-exiting progenitor; NB, neuroblast; RGC, retinal ganglion cell; PR, photoreceptor; HC/AC, horizontal cell and amacrine cell. Minor cell clusters in LV-GFP and LV-NEP infected retinal organoids that do not show characteristic retinal gene expression are designated as “unknown”. The average expression levels are represented by the intensity bar, and the percentages of cells expressing a given gene in a cluster are indicated as the dot size.

Next, we combined different cell clusters assigned to each of the seven cell categories (Fig. 6a) and performed a pseudotime trajectory analysis (Fig. 6b). Without defining the starting and ending points, the trajectory of LV-GFP infected cells revealed that the pre-neurogenic progenitors were closely related to the neurogenic progenitors, which in turn produced exiting progenitors that developed into the postmitotic neuroblasts and a single trajectory including three types of retinal neurons. In both LV-AEP and LV-NEP infected samples, the pseudotime trajectory displayed a more clearly defined progression from the neurogenic progenitors toward the existing progenitors, which in turn gave rise to neuroblasts. The neuroblast cells showed a single node for bifurcated trajectories separating the RGC and HC/AC branch from the photoreceptor branch (Fig. 6b). Quantification of the different cell categories/states further demonstrated that LV-AEP and LV-NEP infection significantly reduced the percentage cells in the pre-neurogenic progenitor category from 25.2 % to less than 4% (Fig. 6c). Neurog2 expression also caused a significant reduction of neurogenic progenitors from 45 % to 26.9 %. Concomitantly, ATOH7f and Neurog2 induction increased the percentage of exiting progenitors from 2.2 % to 4.5 % as well as the percentage of neuroblasts from 5.0 % to 14.9 % and 20.2 %, respectively. Moreover, consistent with marker analysis, elevated ATOH7f and Neurog2 expression resulted in increased proportion of RGCs from 7.0 % to 22.1 % and 27.7 % of total cells, respectively (Fig. 6c). Interestingly, Neurog2, but not ATOH7f, enhanced the photoreceptor population from 3.4 % to 9.8 %; whereas ATOH7f, but not Neurog2, promoted HC/AC proportions from 3.1 % to 4.6 %; (Fig. 6c). These sc RNA-seq data demonstrated that elevating ATOH7f and Neurog2 expression promoted transitions from the pre-neurogenic to the neurogenic state and enhanced neurogenesis of early retinal cell types in retinal organoids.

**Fig. 6.**
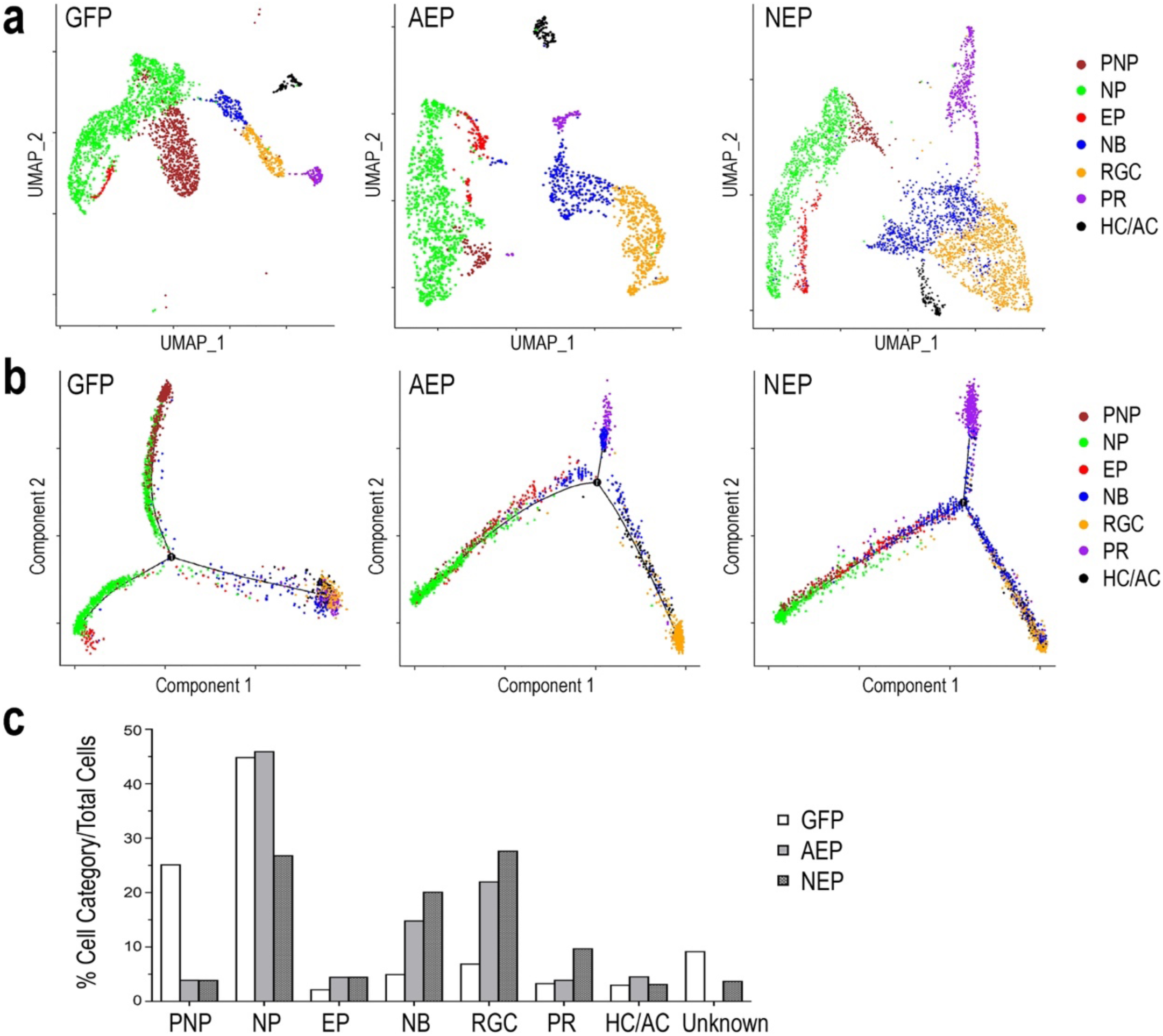
Distinct cell categories based on single cell RNA-seq profiles and their developmental trajectory. **a** UMAP presentations based on cell category assignments for single cell clusters from LV-GFP, LV-AEP, and LV-NEP infected retinal organoids. Each cell category is represented by a single color. The side legend shows the color codes used to represent distinct cell categories. **b** Pseudotime trajectory analysis using Monocle of different cell categories from LV-GFP, LV-AEP, and LV-NEP infected retinal organoid. **c** Bar graph shows the percentages of different cell categories among total cells in LV-GFP, LV-AEP, and LV-NEP infected retinal organoid cells. PNP, pre-neurogenic progenitor; NP, neurogenic progenitor; EP, cell cycle-exiting progenitor; NB, neuroblast; RGC, retinal ganglion cell; PR, photoreceptor; HC/AC, horizontal cell and amacrine cell.

### Neurogenic factors promote transitions of distinct developmental states

Since viral mediated expression of ATOH7 and Neurog2 affected transitions between developmental states, we explored the key characteristics of each cell state. We compiled differentially expressed genes (DEGs, adjusted p values <0.05) of each cell category for LV-GFP, LV-AEP and LV-NEP samples (Supplement Tables 1-3), and constructed heatmaps for the top 10 DEGs displaying more than Log 1.5-fold change of expression levels (Fig. 7a) (Supplement Tables 4-6). In the control LV-GFP infected sample, the top 10 DEGs in the pre-neurogenic progenitor category included SLC2A1, encoding glucose transporter protein type 1 (GLUT1), and GPI, glucose-6 phosphate isomerase, and were quite distinct from those of the neurogenic progenitors. In LV-AEP and LV-NEP infected retinal organoids, the pre-neurogenic progenitors were not only reduced, but also shared genes, such as SFRP2, IFITM3, and VIM with the neurogenic progenitors. In all three virus infected samples, the exiting progenitors shared top DEGs, including HES6, a suppressor of HES1 ^46^, and HMGB2, a member of the chromosomal high mobility group of proteins ^47^, indicating common processes involved in cell cycle withdrawal. The heatmaps also revealed that neuroblasts in each virus infected sample expressed some genes associated with the RGC, photoreceptor, and HC/AC cell lineages, suggesting that the newly postmitotic neuroblasts were poised at an intermediate developmental state and in the process of committing to specific cell fates.

**Fig. 7.**
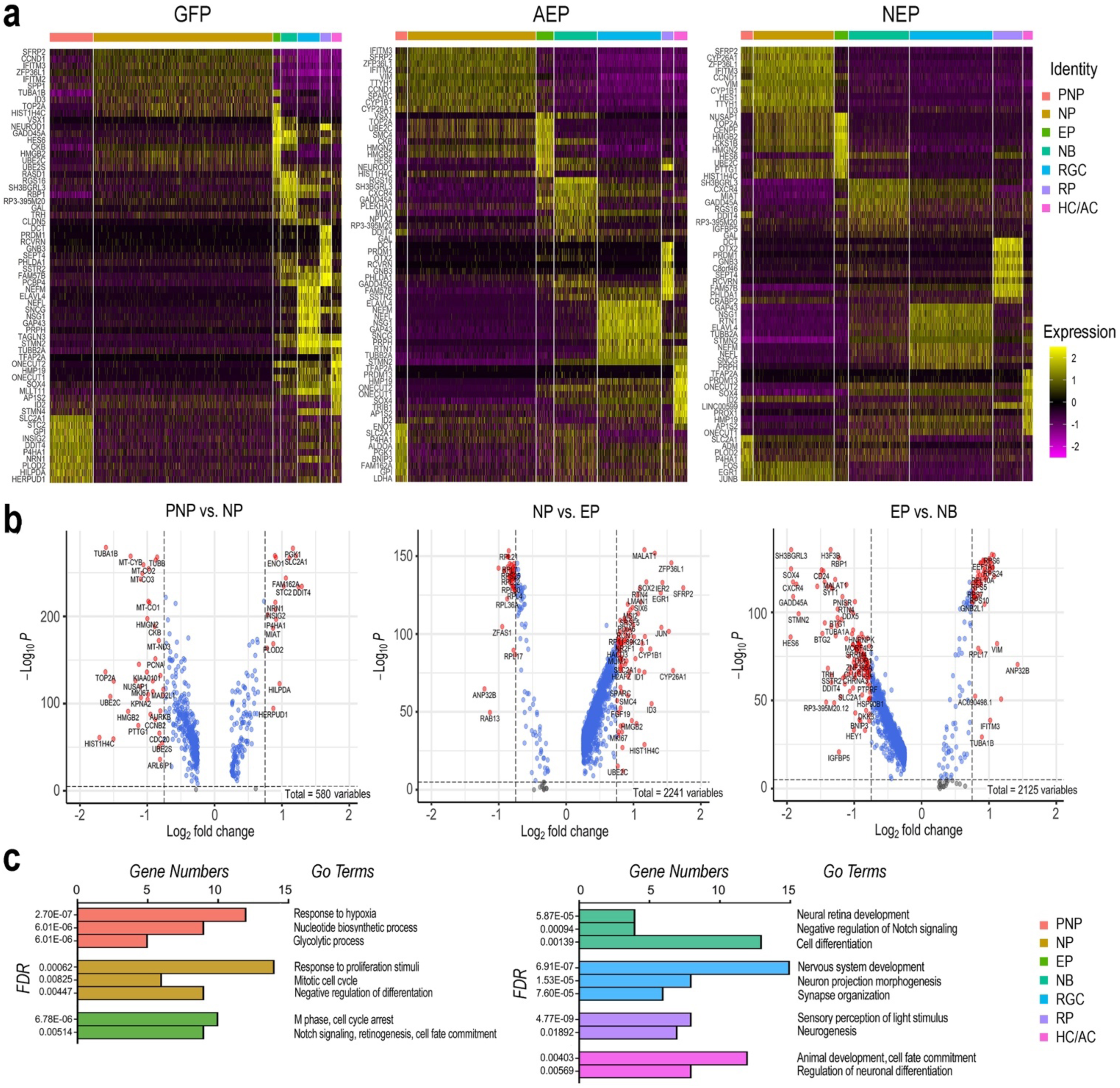
Effects of elevating neurogenic factor expression on developmental transitions. **a** Heatmaps of the top ten DEGs in each cell type category from cells of LV-GFP, LV-AEP, and LV-NEP infected retinal organoids. The side legend indicates the color codes for cell category identities atop the heatmaps. **b** Volcano plots show significant DEGs in LV-GFP infected retinal organoids during the three transitions: from the neural stem cell to the neurogenic progenitor state, from the cycling progenitor to the exiting progenitor state, and from the exiting progenitor to the neuroblast states. Genes with *p* value < 0.01 and fold change > Log_2_ 0.75 are shown as red dots. **c** Bar graphs show the predominant biological processes using GO pathway analysis for each cell category of LV-GFP infected retinal organoids. The false positive rates (FDR) and the gene numbers among the top 25 DEGs associated with a given GO term are shown. PNP, pre-neurogenic progenitor; NP, neurogenic progenitor; EP, cell cycle-exiting progenitor; NB, neuroblast; RGC, retinal ganglion cell; PR, photoreceptor; HC/AC, horizontal cell and amacrine cell.

To further decipher the changes of gene expression and the key biological processes that occur during normal developmental transitions, we constructed volcano plots (Fig. 7b) and performed gene ontology (GO) analyses (Fig. 7c) using significant DEGs from the LV-GFP infected cell categories. The pre-neurogenic progenitor state was associated with higher levels of transcripts for several glycolytic pathway enzymes such as SLC2A1, GPI, ENO1, and PGK1, as well as low levels of mitochondrial respiratory chain components and proteins required for rapid cell proliferation such as PCNA and TOP2A (Fig. 7b). This was consistent with the dominant GO pathways for the pre-neurogenic state (Fig. 7c). In contrast, cells in the neurogenic progenitor state expressed high levels of SFRP2, FGF19, and SOX2 (Fig. 7b), correlating with the biological responses to growth stimulation and the mitotic cell cycle (Fig. 7c). The dominant GO terms associated with the exiting progenitors included cell cycle arrest and Notch signaling (Fig. 7c). The exiting progenitor to neuroblast transition was marked by elevated expression of the Notch signaling inhibitor HES6 and the antiproliferation factors BTG1 ^48^, BTG2, and GADD45A ^49^ (Fig. 7b), consistent with the GO pathway analysis (Fig. 7c). The predominant GO pathways for the three neuronal types reflected their phenotypic differentiation with high axonal growth and light transduction processes associated with RGCs and photoreceptors, respectively (Fig. 7c).

### ATOH7 and Neurog2 assert differential effects on cell cycle and neuronal differentiation

To examine the potential impact of neurogenic factor expression on the cell cycle, we combined the LV-GFP, LV-AEP, and LV-NEP sc-RNA-seq datasets and carried out cell clustering analysis (Fig. 8a). Based on feature plots of known genes (Supplementary Fig. 6), cell clusters in the combined UMAP were identified as distinct cell categories. Furthermore, the progenitor cell clusters were assigned to different phases of the cell cycle (Fig. 8a) based on known functions of cell cycle genes, including CDK2 associated with G1/S phase ^50^, MCM4 as a DNA replication licensing factor ^51^, CCNB2 encoding M phase cyclin B2 ^52^, and PLK1 involved in spindle assembly and cytokinesis ^53^ (Supplementary Fig. 6). Slingshot analysis performed for the combined dataset illustrated the developmental trajectory of cell clusters from the pre-neurogenic state through the different phases of the cell cycle toward cell cycle exit (Fig.8a). Although cell clusters 5 and 6 both expressed M phase genes, they clearly had distinct transcript profiles since cluster 6 was poised to exit the cell cycle (Fig. 8a). As expected, individual UMAPs of ATOH7 and Neurog2 infected samples displayed significantly reduced pre-neurogenic progenitor populations (Fig. 8b; Fig. 6c). In addition, quantitative analyses showed that ATOH7 and Neurog2 infection altered cell cycle distributions among progenitors compared to controls (Fig. 8c). For example, LV-AEP increased G1 distribution by 5 % while decreasing S phase cells by 3.7 %, whereas LV-NEP caused a 7 % S phase cell reduction and a 4.5% increase of M phase cells, which included the exiting progenitors (Fig. 8c).

**Fig. 8.**
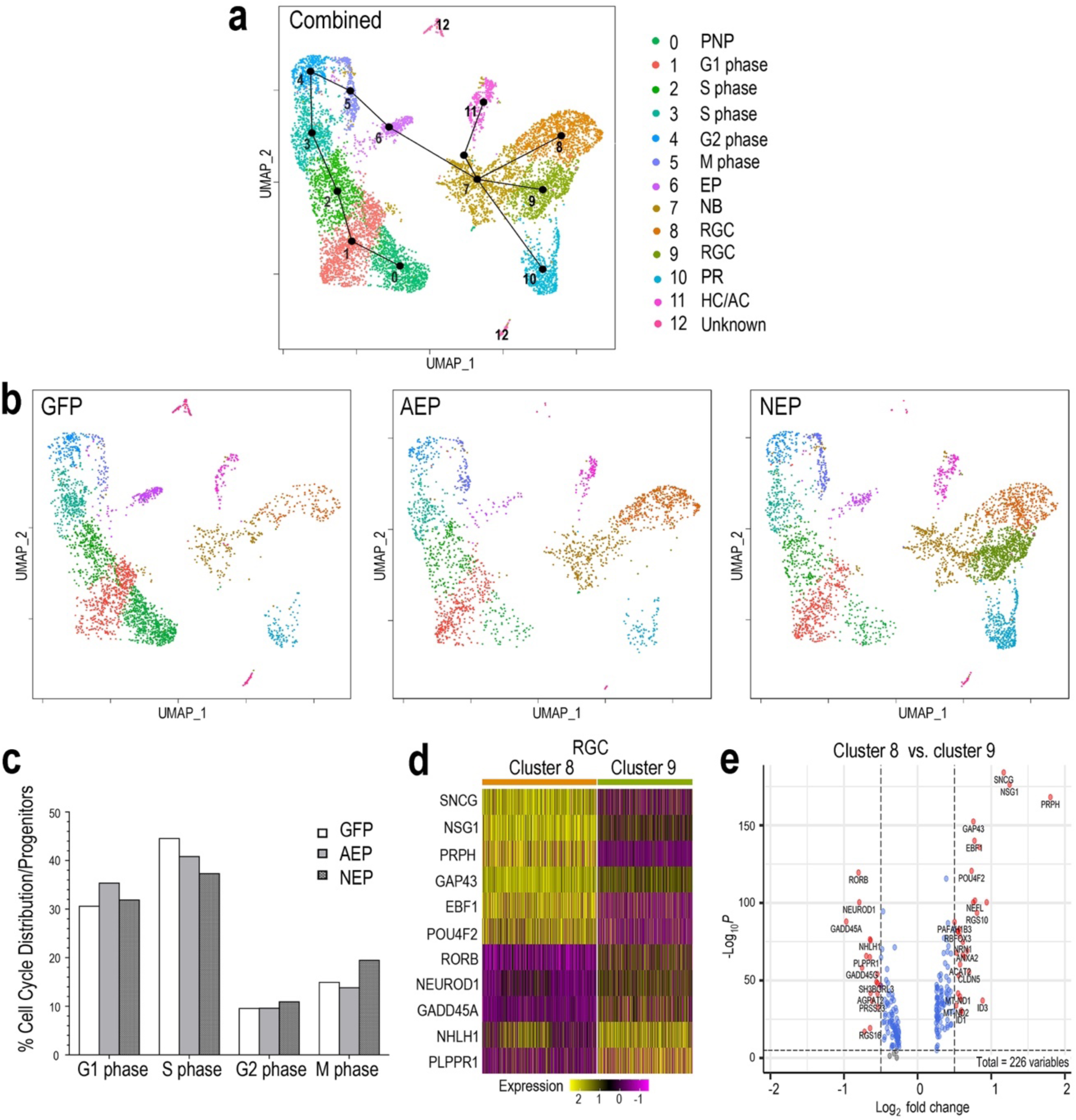
Influence of neurogenic factors on cell cycle progression and retinal ganglion cell differentiation. **a** UMAP of the combined sc RNA-seq dataset from LV-GFP, LV-AEP, and LV-NEP infected retinal organoids. Each cell cluster is assigned to an identity as shown on the side color legend based of feature plots of known genes. Line edges connecting individual clusters are generated by SLINGSHOT analysis to show the predicted developmental trajectory of the entire cell cohort. PNP, pre-neurogenic progenitor; NP, neurogenic progenitor; EP, cell cycle-exiting progenitor; NB, neuroblast; RGC, retinal ganglion cell; PR, photoreceptor; HC/AC, horizontal cell and amacrine cell. **b** UMAPs show cell clusters for individual samples from LV-GFP, LV-AEP, and LV-NEP infected retinal organoids with the same cell identity assignments as in the combined UMAP. **c** Distribution of cells in different phases of the cell cycle as percentages of total proliferative progenitor cells. The M phase cells include the exiting progenitors (cluster #6). **d** Heatmap shows differential expression levels of genes in the two RGC clusters 8 and 9. **e** Volcano plot shows DEGs between the two RGC clusters 8 and 9. Genes with *p* value < 0.01 and fold change > Log_2_ 0.5 are shown as red dots.

The Slingshot analysis demonstrated that the postmitotic neuroblast pool, cluster 7, served as the root source giving rise to three neuronal cell lineages (Fig. 8a). Interestingly, UMAPs of individual virus infected retinal organoids revealed differential effects of ATOH7 and Neurog2 on neuronal fate specification. ATOH7 elevation resulted in clear enhancement of the RGC cluster 8, without affecting the photoreceptor cluster 10 (Fig. 8b, also see Fig. 6c). In contrast, Neurog2 overexpression not only significantly enhanced the production of photoreceptor cluster 10 and RGC cluster 8, but also generated a distinct RGC cluster 9, which was largely absent in LV-GFP and LV-AEP infected retinal organoids (Fig. 8b, also see Fig. 6c). Comparison analyses revealed that the two RGC clusters consisted of cells with differential gene expression levels. For example, GAP43 and NSG1 were expressed by cells in both clusters, however, they were frequently detected at higher levels in cluster 8 (Fig. 8d, 8e). Although both clusters expressed ISL1 and POU4F2 (Supplementary Fig. 6), we often observed lower levels of POU4F2 and SNCG in many cluster 9 cells (Fig. 8d). These results suggest that ATOH7 and Neurog2 might differentially influence neuroblast fate specification and/or neuronal differentiation.

### Neurogenic factors modulate early retinal gene network

Next, we examined the effects of viral mediated ATOH7f and Neurog2 elevation on endogenous bHLH gene expression. Since the lentiviral vectors did not contain poly-A sequences associated with ATOH7 and Neurog2 cDNAs, and the single cell cDNA libraries were constructed using oligo dT priming, we were able to analyze expression of the endogenous genes using the sc RNA-seq datasets, while excluding transgenes encoded by the viruses. We first examined expression of ASCL1, an early onset bHLH neurogenic factor. Consistent with previous studies ^34, 36^, feature plots showed that ASCL1 was predominantly expressed in pre-neurogenic and neurogenic progenitors (Fig. 9a). Viral mediated ATOH7f and Neurog2 expression resulted in significant reduction of ASCL1^+^ cell from 29.7 % to 17.1 % and 13.2 %, respectively (Fig. 9c); but neither affected the median levels of ASCL1 expression (Fig. 9b). In retinal organoids, ATOH7 and NEUROG2 were detected in exiting progenitors, neuroblasts, and some postmitotic neurons (Fig. 9a). Both virally expressed ATOH7f and Neurog2 increased the number of endogenous ATOH7-expressing cells from 9.1 % to 22.0 % among total cells (Fig. 9c), and LV-NEP infection also increased the median ATOH7 expression level (Fig. 9b). Interestingly, LV-NEP but not LV-AEP infection caused a 2-fold increase of endogenous NEUROG2-expressing cells as well as elevated the median expression level (Fig. 9b, 9c), suggesting that Neurog2 positively regulates endogenous NEUROG 2 expression. In addition, elevated levels of ATOH7f and Neurog2 both correlated with the increased percentages of cells expressing OLIG2, another bHLH gene in retinal organoids ^54^ (Fig. 9c).

**Fig. 9.**
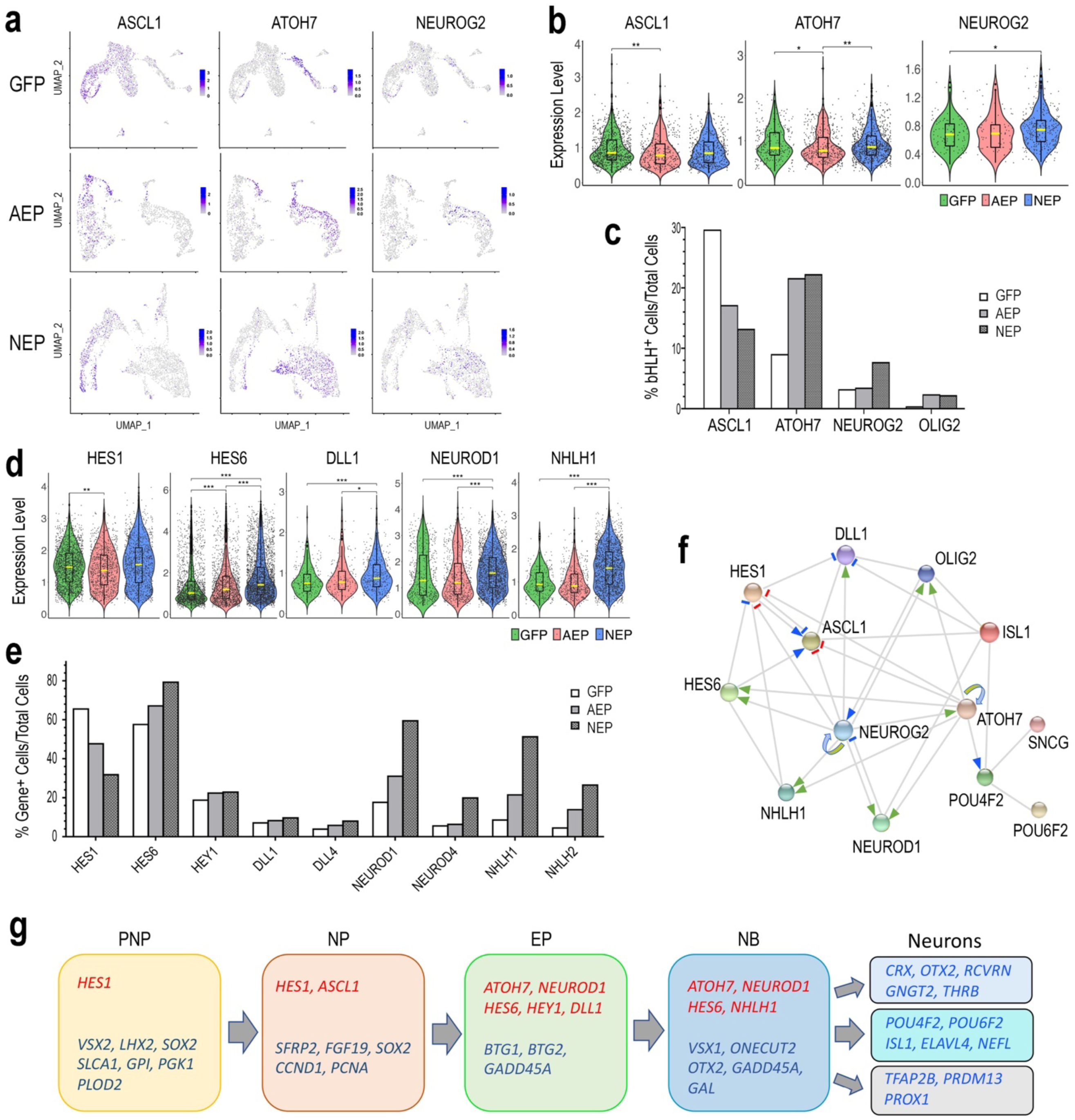
Regulatory relationship among genes playing roles in early human retinogenesis. **a** Feature plots show distribution of ASCL1, ATOH7, and NEUROG2 expressing cells among cell clusters in LV-GFP, LV-AEP, and LV-NEP infected retinal organoids. **b, d** Violin plots show comparisons of LV-GFP (green) versus LV-AEP (red) or LV-NEP (blue) induction on expression levels of endogenous genes in the retinal organoids. The box within the violin represents the middle 50% of the data, and the yellow line within the box indicates the median expression level. Statistical analysis generated P-values were based on both the cell counts and expression levels. ***p<0.001, ** p< 0.01, *p< 0.05. **c, e** Bar graphs show percentages of cells expressing endogenous genes among total viral infected cells in LV-GFP, LV-AEP, and LV-NEP samples. **f** Schematic model based on STRING analysis summarizes a gene network involved in cell cycle exit and early retinogenesis. Grey edges represent protein-protein associations. Positive and negative regulatory relationships are indicated as arrows and short bars, respectively. Previously known molecular interactions from curated databases are shown as blue arrows and short bars. Effects of lentiviral induced ATOH7f and Neurog2 on endogenous human gene expression in retinal organoids are indicated as green arrows and red bars. Looping arrows indicate positive auto-regulation. **g** Summary of sequential expression of bHLH genes (red) and other key genes (blue) during early human retinal organoid development. PNP, pre-neurogenic progenitor; NP, neurogenic progenitor; EP, cell cycle-exiting progenitor; NB, neuroblast; RGC, retinal ganglion cell; PR, photoreceptor; HC/AC, horizontal cell and amacrine cell.

In addition to ASCL1, ATOH7, NEUROG2, and OLIG2, sc RNA-seq analysis identified additional Notch signaling pathway genes and bHLH genes affected by ATOH7f and Neurog2 expression (Supplementary Table 1-3; Supplementary Fig. 7). Violin plots and quantification showed that the Notch ligands DLL1 and DLL4 were upregulated, especially by Neurog2 (Fig. 9d, 9e). In addition, the percentage of cells expressing the Notch signaling effector HES1 was significantly reduced compared to the control LV-GFP infected cells (Fig. 9e), while changes in median expression levels were mild (Fig. 9d). In contrast, not only the percentages of HES6 and HEY1 expressing cells were increased (Fig. 9e), the median HES6 expression levels were significantly elevated by bot LV-AEP and LV-NEP (Fig. 9d). Since HES1 was predominantly expressed by progenitors, whereas HES6 and HEY1 were upregulated in exiting progenitors and neuroblasts (Supplementary Fig. 7), these changes reflected the trend of enhanced neurogenesis. Data from sc RNA-seq also showed that viral expression of ATOH7f or Neurog2 caused substantial increases of several bHLH genes involved in neuronal fate specification or differentiation, including NEUROD1, NEUROD4, NHLH1 and NHLH2 (Fig. 9d, 9e; Supplementary Fig. 7). In all cases, elevated Neurog2 strongly impacted both the number of cells expressing these genes as well as their expression levels.

Together, these results suggested that viral mediated ATOH7f and Neurog2 expression in the developing retinal organoids affected an interactive gene network that plays important roles in cell cycle exit and cell fate specification. We therefore performed STRING network analysis by including bHLH genes and a few selected genes known to be involved in RGC development. The resulting network model included novel regulatory relationships revealed in this study as well as previously reported molecular interactions (Fig. 9f). We also summarize the observed temporal progression of gene expression as retinal organoid cells advanced through the developmental states as defined by our sc RNA-seq analysis (Fig. 9g).

## Discussion

In this study we have used ESC-derived organoid cultures as a model system to investigate human embryonic retinal development. The 3D retinal organoids retain the unique molecular signature and correct tissue polarity of the retinal epithelium in vivo, and generate the expected early retinal cell lineages and functional neurons. By transducing human retinal organoids with inducible viral vectors, we were able to determine whether elevating ATOH7 or Neurog2 affected retinal progenitors and neuronal production, and examine the biological processes and the gene network involving these neurogenic factors.

We have identified three classes of retinal progenitors by performing single cell transcriptome analysis. The pre-neurogenic progenitors express PAX6, VSX2, LHX2, and SOX2, and thus clearly possess the identity of the retinal primordium. Their main GO pathways show cellular metabolic characteristics common to stem cells ^55^, with prominently featured DEGs of the glycolytic pathway, including SLC2A1, GPI, PGK1, and PLOD2, and low mitochondrial respiratory chain genes. This state likely represents a naïve pre-neurogenic state found in the retinal primordium before the onset of neurogenesis and later at the ciliary margin of the mature retina. These pre-neurogenic progenitors express cyclin D1, but have relatively low levels of PCNA and TOP2A, suggesting that they might be slow cycling cells. The pre-neurogenic progenitor category was significantly reduced in LV-AEP and LV-NEP transduced retinal organoids, indicating that expression of these neurogenic factors promotes the transition from the pre-neurogenic to the neurogenic state. This finding is consistent with our previous observation in the embryonic chicken retina, where ectopic ATOH7 expression in the pre-neurogenic peripheral retina induces precocious neurogenesis ahead of the neurogenic wave front ^25^. These results indicate that the onset of neurogenic factor expression among naïve retinal progenitors can trigger or accelerate the transition into a neurogenic state in the context of the retinal primordium. Noticeably, the transition from the pre-neurogenic state to the neurogenic state is also accompanied by upregulation of the chromosomal high mobility group genes, such as HMGB2 and HMGN2, reflecting the underlying epigenetic changes accompanying this transition.

Our sc RNA-seq analysis also identified a novel and distinct progenitor state among the proliferating neurogenic progenitors, the cell cycle exiting progenitors. This group of cells still express the characteristic M phase genes such as CDC20 that interacts with the mitosis complex, and MZT1 and PLK1 that organize the mitotic spindles. However, the exiting progenitors show significant upregulation of the Notch ligands DLL1 and DLL4, as well as the bHLH genes ATOH7 and NEUROD1. Both viral mediated ATOH7 and Neurog2 elevation caused expansion of the exiting progenitor state. The developmental trajectory analysis points to a progression from the exiting progenitors to the postmitotic neuroblasts, which as a group exhibits transcriptome profile partially overlapping with all three early neuronal lineages, including VSX1, NHLH1, ATOH7, ONECUT2, NEUROD1, and the growth arrest gene GADD45A. However, transcription profiles of individual cells among neuroblasts are heterozygous, likely reflecting the dynamic gene expression of neuroblasts that serve as an important transitional cell pool poised for terminal cell fate choices and differentiation. Interestingly, we have observed that relative to ATOH7, elevating Neurog2 asserted more potent effects in promoting cell cycle withdrawal, neuroblast expansion, and neuronal differentiation in retinal organoids.

Among the early bHLH factors detected by sc RNA-seq in retinal organoids, a higher percentage of cells expressed ATOH7 (9%) than NEUROG2 (3.2%) endogenously. Although elevating either ATOH7 or Neurog2 caused enhanced neurogenesis, we also detected differential effects of these two factors. For example, at Day 39, ATOH7 promoted POU4F^+^ cells, whereas Neurog2 increased ISL1^+^ cells; but both factors enhanced ISL1^+^ cells by Day 47. Furthermore, the RGC cluster enhanced by ATOH7 elevation shared transcriptomic signatures with RGCs from the control organoids between Day 45-48. In contrast, Neurog2 induction promoted two groups of RGCs that both expressed ISL1 and POU4F2, but exhibited different levels of known RGC markers including SNCG, POU4F2, and GAP43. Recent transcriptome analysis of mature human RGCs has indeed detected differential expression levels of RGC genes such as POU4F2 among RGC subtypes ^42^. Therefore, the promotion of the two RGC subtypes may reflect the more potent neurogenic effect of Neurog2, which might have accelerated the RGC differentiation program to acquire RGC subtype features. In addition, the effect of elevated Neurog2 to induce ISL1^+^ neurons preferentially earlier on may also influence outcomes of neuronal differentiation. However, we cannot rule out that virally expressed exogenous Neurog2 has promoted a hybrid neuronal cell type, which does not naturally occur during human retinogenesis. Future research using long term cultures may provide data to address these possibilities. In addition to the RGC phenotypes, ATOH7 increased the HA/AC lineage, consistent with the developmental trajectory. The lack of ATOH7 enhancement on cone photoreceptor production is somewhat unexpected, as ATOH7 is associated with cone cell lineage in the mouse retina ^19^, and we have previously detected a mild enhancement of ATOH7 on cone cell genesis in the chicken retina ^25^. In contrast, Neurog2 significantly enhanced photoreceptor production without affecting the HC/AC lineage, which likely reflects the effect of Neurog2-induced NEUROD1 upregulation.

Expression of cell intrinsic factors can be profoundly influenced by extrinsic signaling events during neural development ^56^. Notch mediated cell-to-cell signaling plays important roles during retinogenesis, and disruption of Notch signaling can influence RGC and cone photoreceptor development ^6, 57–60^. In addition, differentiated RGCs have been shown to produce secreted signals that modulate progenitor behaviors. Among the signaling molecules released by differentiated RGCs are Shh, GDF11, and VEGF, all of which promote retinal progenitor proliferation while simultaneously suppressing RGC production ^61–64^. Accumulating evidence indicate that the Notch signaling downstream effector Hes1 may serve as an important integration node for distinct signaling pathways that converge upon retinal progenitors to influence cell proliferation and control neurogenesis. Hes1 is known to directly control cell proliferation by repressing the CDK inhibitor p27(Kip1) ^46^. The negative feedback of extrinsic signals on RGC genesis is in part mediated by Hes1 suppression of ATOH7 ^65–67^. At the present time, the temporal expression sequence and the complex regulatory relationships among the various bHLH factors in the retina are not fully understood, but their elucidation is likely crucial for a better understanding of cell fate selection and the progressive changes in progenitor competent states.

As revealed by our sc RNA-seq data, HES1 is predominantly expressed by both pre-neurogenic and neurogenic progenitors, and is down regulated among exiting progenitors. Consistent with observations in embryonic human retinas ^36^, our data indicate that ASCL1 level is low in pre-neurogenic progenitors, but is upregulated and expressed by neurogenic progenitors, and then down-regulated in exiting progenitors. In the developing mouse retina, Ascl1 can block cell cycle exit but not specify RGC fate ^68^, while in the mature retina ASCL1 promotes cell cycle reentry and regeneration of new neurons from Muller glia ^69–71^. Furthermore, forced expression of ASCL1 in pluripotent stem cells can promote neurogenesis^72^. These data together support that ASCL1 plays a role in establishing competence for the neurogenic state. Intriguingly, our results show that viral mediated ATOH7 and Neurog2 expression decrease both HES1 and ASCL1 expression. Concomitantly, ATOH7 and Neurog2 significantly upregulate the expression of the HES1 inhibitor HES6 ^46^ among exiting progenitors. These data suggest that ATOH7 and NEUROG2 comprise a selective group of neurogenic factors that can dampen the effect of HES1 and ASCL1 on maintaining cell proliferation and relieve the inhibitory effect of HES1 on neurogenesis. Moreover, viral mediated ATOH7 and Neurog2 expression significantly increase transcripts of other bHLH factors including NEUROD1, which initiates its expression among exiting progenitors, as well as NEUROD4, NHLH1 and NHLH4, which are expressed by postmitotic neurons. Future investigations are necessary to determine whether the regulatory effects asserted by ATOH7 and Neurog2 are direct or indirect.

Results from this study support the hypothesis that high levels of AOTH7 or NEUROG2 trigger a withdrawal from the cell cycle which leads to the birth of a neuroblast. Critical questions remain regarding how the level of ATOH7 is regulated in neurogenic progenitors to result in higher levels among cell cycle exiting progenitors. It is known that the ATOH7 gene contains enhancers that mediate direct positive regulation by Pax6 ^73^. In Hes1 mutants, Atoh7 is precociously expressed along with the formation of RGC and HC/AC ^74^, indicating that Hes1 negatively regulates Atoh7 expression. Our sc RNA-seq analysis show that viral mediated ATOH7 expression elevates endogenous ATOH7 without affecting endogenous NEUROG2 expression, whereas Neurog2 induction leads to increases in both ATOH7 and NEUROG2 expression. These results reveal the novel finding that ATOH7 and NEUROG2 are both under positive autoregulation, as well as confirm cross-regulation of ATOH7 by NEUROG2 ^16, 75^. Since only a fraction of ATOH7 protein expressing cells appear to co-express Neurog2 protein during a given window of time ^9^, progenitors that have coincidental expression of both factors are more likely to reach higher level of Atoh7. It is known that ATOH7 dosage and expression levels can affect RGC production ^25, 26, 76^. We therefore propose that integrated cell extrinsic signals and interacting cell-intrinsic factors could converge, resulting in stochastic expression of the ATOH7 gene, and thus enabling a limited subset of progenitors that have reached a threshold level of ATOH7 to exit the cell cycle and initiate downstream RGC and/or cone photoreceptor differentiation programs. Further investigations will be necessary to determine the ATOH7 protein threshold, the time, and cellular events involved. The sc RNA-seq analysis enables us to survey multiple genes and to construct a gene regulatory network model (Fig. 9f) that integrates our new findings and previously regulation. This model focuses on bHLH factors expressed during RGC development, but does not exclude other genes involved in the developmental process. In fact, RGC and other neuronal fate determinations are known to be regulated by multiple transcription factors ^77–83^, which together coordinate epigenetic changes and orchestrate transcriptomic responses required for neuronal differentiation.

In summary, our study show that elevating ATOH7 and Neurog2 expression in human retinal organoids significantly enhances retinogenesis, which can serve as a useful approach to produce authentic human RGCs for studying development and degenerative diseases. Our single cell transcriptome analysis provides novel insights into the interactive network of bHLH factors and their functions during human retinogenesis.

## Materials and Methods

### Lentiviral construction and production

The inducible lentiviral vector plasmids LV-GFP and LV-NEP (YS-TetO-FUW-Ng2-P2A-EGFP-T2A-Puro) were generous gifts from Dr. Thomas Sudhof ^44^. The human ATOH7 cDNA was obtained as described previously ^25^. The LV-ATOH7f vector was constructed by replacing the EGFP gene in the LV-GFP vector with ATOH7 cDNA fused to the Flag epitope tag at the c-terminus. The LV-AEP vector was constructed by custom synthesizing the continuous open reading frame of ATOH7f-P2A-EGFP-T2A-Puro by GenScript, and replacing the EGFP gene in the LV-GFP vector. All lentiviral vector plasmids were verified by DNA sequencing. The LV-rtTA lentiviral vector (FUW-M2rtTA) with the Ubi promoter was obtained from AddGene (Plasmid #20342).

Lentiviral stocks were produced by co-transfection of HEK 293T cells with a given viral vector DNA and the third-generation lentiviral helper plasmids with VSVG pseudotyping as described ^84–86^. The 293T cell medium DMEM containing 10% fetal bovine serum (SigmaAldrich, 12103C) was changed to serum free CD293 (ThermoFisher, 11913) one day post transfection, and viral supernatants were harvested every 24 hours. Combined viral stocks were concentrated by ultracentrifugation as previously described ^86^.

### Human retinal organoid derivation

Human H9 ES cells were cultured and passaged on Matrigel (Corning, 356231) coated dishes in mTeSR1 medium (Stemcell Technologies, 05850). Retinal organoids were generated based on a previously described protocol ^30^ with modifications. At the start of the culture (Day 0), H9 ES cells (at 80-90% confluency) were enzymatic detached using dispase (1 mg/ml, Stemcell Technologies, 07923). Detached cells were transferred into medium at 3:1 ratio of mTeSR1 to Neural Induction Medium (NIM) that consists of DMEM/F12 with 1x N2 supplement (ThermoFisher, 17502048), 1x non-essential amino acid (NEAA; ThermoFisher, 11140050), 2 μg/ml Heparin (ThermoFisher, H7482), and 1x Antibiotic-Antimycotic (Anti-Anti; ThermoFisher, 15240112) in low-attachment plates (Corning, 3471) to allow the formation of embryonic bodies (EBs). During the next three days, the medium was replaced daily with the ratio of NIM to mTeSR1 increased to 100%. At Day 6, human BMP4 (R&D Systems, 314-BP-010) was added to the NIM medium to a final concentration of 55 ng/ml ^43^. At Day 7, EBs were collected and seeded in NIM as adherent cultures in 6-well dishes (Corning, 3516) till Day 16. At Day 16, the visible neural rosettes formed from attached EBs were manually lifted, collected, and further cultured as suspensions in Retinal Differentiation Medium (RDM) consisting of DMEM to F12 at 3:1, 1x B27 supplement (ThermoFisher,1754044), 1x NEAA, and 1x Anti-Anti. During the 7 days after lifting neural rosettes, 5 μM SU-5402 (SigmaMillipore, SML0443) and 3 μM GSK inhibitor CHIR99021 (Stemgent, 04-0004) were added to RDM. At Day 20, the translucent optic vesicle-like structures were manually separated from the rest of the suspension culture, collected, and cultured as retinal organoids in RDM. From Day 24, 10 % FBS (SigmaAldrich, 12103C), 100 μM Taurine (SigmaAldrich, 0625), and 500 μM retinoic acid (SigmaAldrich, R2625) were added, and the medium was changed twice a week.

### Lentiviral Infection and transgene induction

Retinal organoids were infected by different lentiviruses in conjunction with LV-rtTA three times between Day 23 and Day 40. TetO promoter induction was carried out by adding doxycycline (Dox) to a final concentration of 2 μg/ml (SigmaAldrich, D3072) according to experimental designs as indicated in the results.

### Attached and dissociated retinal organoid cultures

Retinal organoids between Day 30-35 were cut into small pieces (0.1-0.5 mm) and plated on Matrigel coated culture dishes, or dissociated with Trypsin (SigmaAldrich, T9935) to single cells and plated on poly-D-lysine and laminin coated glass coverslips (Corning, 354087). After attachment, retinal organoid cells were cultured in RDM or BrainPhys neuronal medium (Stemcell Technologies, 05790) with SM1 (Stemcell Technologies, 05711) and N2 supplements (ThermoFisher, 17502048), 20 ng/ml BDNF (PeproTech, 450-02), 20 ng/ml GDNF (Stemcell Technologies, 78058), 1 mM dibutyryl cyclic-AMP (Stemcell Technologies, 73882), and 200 nM ascorbic acid (Stemcell Technologies,72132) till desired time, followed by immunofluorescent labeling or electrophysiological recordings.

### Electrophysiological recording

Electrophysiological recordings were performed using dissociated retinal organoid cells cultured as a monolayer on glass coverslips between Day 40-45 using previously described methodologies ^87–90^. Whole cell patch clamp was performed at room temperature using an Axopatch 200B amplifier controlled by pClamp 11 data acquisition software (Molecular Devices). The pipette solution contained 20 mM KCl, 120 mM K-gluconate 0.1 mM CaCl_2_, 1 mM EGTA, 10 mM HEPES, 3 mM Mg-ATP, 0.2 mM Li-GTP, and 8 mM phosphocreatine, at pH7.2. The bathing solution (for recording K^+^ currents, Na^+^ currents and for current clamp) contained125 mM NaCl, 3mM KCl, 2 mM CaCl_2_, 1.25 mM NaH_2_PO_4_, 1 mM MgCl_2_, 25 mM NaHCO_3_ and 10 mM glucose bubbled continuously with 95% O_2_ – 5% CO_2_. A high barium external solution for Ca^2+^ channel current recordings containing 110 mM choline chloride, 5 mM KCl, 1 mM MgCl_2_ 7 mM BaCl_2_, 15 mM TEACl, 0.1 mM 4-aminopyridine, 20 mM glucose, 10 mM HEPES and 1 µM tetrodotoxin, adjusted to pH 7.4, was used with a CsCl intracellular solution containing 140 mM CsCl, 1 mM CaCl_2_, 11 mM EGTA, 2 mM MgCl_2_, and 10 mM HEPES, at pH7.2.

### Immunohistochemistry and imaging

For live imaging of whole mounts retinal organoids, EGFP signals were first captured using a Leica MZ10F fluorescent dissecting microscope, followed by image acquisition using an Olympus Flowview FV1000-IX81 (inverted) scanning laser confocal microscope. For whole mount immunolabeling, retinal organoids were fixed with 4% paraformaldehyde (PFA) in PBS for 30 minutes, followed by primary and secondary antibody incubations overnight at 4°C, with extensive washes in between ^25^. Cryosections (14 μm) or attached cultures were processed for immunolabeling as previously described ^25^.

The following primary antibodies were used: mouse anti-GFP(1:200; Millipore, MAB#3580); goat anti-GFP (1:200; Rockland Inc, 600-101-215); rabbit anti-GFP (1:200; Rockland Inc, 400-401-215); mouse anti-FLAG (Clone M2)(1:500; SigmaAldrich, F3165); rabbit anti-ATOH7 (1:100; NovusBio, NBP1-88639); goat anti-Ngn2 (1:200; Santa Cruz Biotechnology, sc-19233); mouse anti-BrdU (1:1; GE Healthcare, RPN202); mouse anti-PCNA (1:500; SigmaAldrich, P3825); rabbit anti-phospho-histone 3 (ser10) (1:3000; Upstate Biotechnology, 06-570); rabbit anti-Pax6 (1:200; Chemicon, ab5409); goat anti-CHX10/VSX2 (N-18) (1:50; Santa Cruz Biotechnology, sc-21690); goat anti-BRN3a (1:100; Santa Cruz Biotechnology, sc-31984); goat anti-pan BRN3 (1:50; Santa Cruz Biotechnology, sc-6026); mouse anti-Iselt1 (1:10; Developmental Study Hybridoma Bank, 39.4D5); mouse anti-doublecortin (E-6) (1:50; Santa Cruz Biotechnology, sc-271390); rabbit anti-NeuN (1:200; Abcam, ab177487); rabbit anti-NF145 (1:750; Millipore, AB1987); rabbit anti-RBPMS (1:200; ^91^. Secondary antibodies used were: Alexa Fluor 488 donkey anti-mouse (1:500; ThermoFisher, A32766); Alexa Fluor 488 donkey anti-rabbit (1:500; ThermoFisher, A32790); Alexa Fluor 488 donkey anti-goat (1:500; ThermoFisher, A32814); Alexa Fluor 594 donkey anti-mouse (1:500; ThermoFisher, A32744); Alexa Fluor 594 donkey anti-rabbit (1:500; ThermoFisher, A32754); Alexa Fluor 594 donkey anti-goat (1:500; ThermoFisher, A32758); Alexa Fluor 647 donkey anti-mouse (1:500; ThermoFisher, A32787); Alexa Fluor 647 donkey anti-rabbit (1:500; ThermoFisher, A32795); Alexa Fluor 647 donkey anti-goat (1:500; ThermoFisher, A32849).

Slides were mounted with Fluoro-Gel (Electron Microscopy Sciences, 17985-10) after staining with DAPI (SigmaAldrich, D9542). Confocal images were acquired using an Olympus Flowview FV1000-BX61 (upright) scanning laser microscope with Plan-APO objectives. Images were arranged using Adobe Photoshop.

### Cell marker quantification and statistics

Lentivirus infected retinal organoids at Day 39 were pooled (10-15 organoids in each sample n), dissociated into single cell suspensions using trypsin (SigmaAldrich, T-9935), and immunolabeled as previously described ^63^. Flow cytometry was performed using LSRII Analytic Flow Cytometer for cell marker analyses. Quantification of FACS data was performed using FlowJo software (Tree Star, Inc.). In addition, pooled retinal organoids at Day 47 were dissociated with trypsin and plated as a monolayer for 3 hours followed by immunolabeling for the various cell markers listed above. Monolayer cell quantification was performed by counting marker-positive cells in multiple fields of independent samples (n=3) using captured confocal images. Bar graphs were constructed using Prism (Graphic Pad). Ordinary one-way ANOVA with Tukey’s multiple comparison test was used for statistical analysis of cell marker quantifications, with *p* value < 0.05 considered significant.

### Single cell cDNA library preparation and sequencing

Distinct pools of H9 ES cell-derived retinal organoids (12-20 retinal organoids/pool) co-infected by LV-rtTA and LV-EGFP, LV-AEP, or LV-NEP were induced by Dox and dissociated between Day 45 and Day 48 using trypsin and manual trituration. Dissociated cell suspensions were subjected to fluorescence activated cell sorting using FACSAriaII (BD Biosciences). Non-infected retinal organoid cells were used to set thresholds for selecting EGFP-positive cells. Sorted EGFP-positive cells were collected in HBSS without Ca^2+^ and Mg^2+^ (ThermoFisher, 14170-112) containing 1% FBS and 0.4% BSA. The cells were washed with PBS containing 0.04% BSA, then counted with Countess II Cell Counter (ThermoFisher).

Automated single-cell capture, barcoding, and cDNA library preparation were carried out using 10X Genomics Chromium Controller with Chromium Single Cell 3’ Library & Gel Bead Kit v2 reagents, with 12 cycles of cDNA amplification and 12 cycles of library amplification, following the manufacturer’s instructions. Qubit dsDNA Assay kit (Life Technologies) and TapeStation 4200 (Agilent) were used to assess the quality and concentration of the libraries. Illumina NovaSeq6000 S2 paired-end 2×50bp mode was used to sequence the libraries.

### Single cell RNA-sequencing data processing and quality control

10X Genomics Cell Ranger version 2.1.1 was used to demultiplex the raw base calls into FASTQ files (cellranger mkfastq). Spliced Transcripts Alignment to a Reference (STAR) version 2.5.1b (cellranger count) was used to perform sequence alignments to the reference human genome (GRCh38), barcode counts, and UMI counts to yield summary reports and t-Stochastic Neighboring Embedding (t-SNE) dimensionality reduction. For downstream analyses, cells with a number of unique molecular identifiers (UMI) > 2500 per cell and < 0.1 % mitochondrial gene expression were used. For LV-GFP, LV-AEP, LV-NEP samples, the mean reads per cell ranged from 139,000-195,000, with mean gene per cell ranging from 2935-3079. The resulting total single cell counts used for analysis were 3004 for LV-EGFP, 2063 for LV-AEP, and 3909 for LV-NEP infected samples.

### Single cell RNA-sequencing data analysis and visualization

The analysis of sc RNA-seq data was performed using Seurat R package (https://satijalab.org/seurat/v2.2) ^45, 92^. Clustering of cells was performed by using Seurat FindCluster function (top 20 principal components, resolution 0.8) that implements the shared nearest neighbor modularity optimization algorithm. Nonlinear dimensionality reduction using UMAP (Uniform Manifold Approximation and Projection) was applied for the visualization of cells in two-dimensional space. Feature plots of known genes were used to designate clusters observed in the UMAP space into six major cell categories/states. Cell counts of each category were obtained using custom R code.

Pseudotime developmental progression of cell states was obtained by using the Monocle R package (version 2) to process the datasets with cell labels corresponding to the six cell categories and visualized as UMAPs. Pseudotime cell cycle progression and cell fate adoption analysis was performed using Slingshot R (version1.6.1) by combining the LV-GFP, LV-AEP, and LV-NEP sc RNA-seq datasets and assigning the start point as neural stem cells and end points as differentiated neuronal cell types.

Differentially expressed genes (DEGs) were identified using edgeR with significantly enriched genes in each cell category defined as those with adjusted *p* values < 0.05 for LV-GFP, LV-AEP, and LV-NEP data sets (Supplementary Tables 1-3). The top 10 enriched DEGs in each cell category were defined as those with adjusted *p*-value < 0.05, and log fold change >1.5 (Supplementary Tables 4-6). Heatmaps of the top 10 DEGs were generated using the Seurat package. Volcano plots were generated using EnhancedVolcano R package to show *p* values and fold changes of DEGs between two datasets. Gene Ontology (GO) Enrichment analysis was performed using ShinyGO v0.61 (http://bioinformatics.sdstate.edu/go) ^93^ and the Homo Sapiens background using p-value (FDR) cutoff at 0.05. The top 25 DEGs from each cell category within the LV-GFP dataset were used as inputs, and the redundancy of the output biological processes was manually reduced to the most predominant GO terms.

Feature plots of individual gene expression patterns in different cell clusters were presented as UMAPs. Violin plots for individual genes in all cell clusters were constructed to show expression levels and cell distributions. Kruskal-Wallis one-way ANOVA rank sum test and Tukey-Kramer-Nemenyi all-pairs test were used for statistical analysis, taken into consideration of both gene expression levels and cell numbers between different samples, with *p* value < 0.05 considered significant. The statistical tests were performed on R Studio using ‘PMCMRPlus’ ^94^ and ‘FSA’ ^95^ packages. Data were plotted using R Studio ‘ggplot2’ ^96^ and ‘ggsignif’ ^97^ packages. Cells with gene expression level < 0.2 were exclude from the violin plots and statistical analyses.

STRING analysis exploring protein-protein association network was performed using human protein database (version 11.0; https://string-db.org/) by inputting relevant genes involved in retinal development. The schematic network model shows known molecular interactions reported previously and new regulatory relationships described in this study.

## Acknowledgments

This work was supported by NIH grant R01EY026319 to XJY, NIH core grant P30EY000331, and an unrestricted grant from the Research to Prevent Blindness to the Department of Ophthalmology at University of California Los Angeles.

## Author Contributions

X-J Y. conceived the project and designed lentiviral vectors; X.Z. carried out human retinal organoid and neuronal cultures, performed immunohistochemistry and imaging, and FACS analysis; X.Z. and K.H.N. prepared viral stocks and sc RNA-seq libraries; T.T.T.N, I.M., and M.P. performed bioinformatic analysis; T.T.T.N. performed statistical analysis; J.C.R.G and S.B. performed electrophysiological recordings; X.Z, T.T.T.N., and X-J Y. prepared the figures, and analyzed the data; X-J Y. wrote the manuscript. All authors reviewed and commented on the manuscript.

## Competing Interests

The authors declare no competing interests.

## Supplementary information

Description of additional supplementary files, Supplementary Figures and Tables.

## Data availability

The authors have deposited the sc RNA-seq datasets into the public archive Dryad (https://doi.org/10.5068/D1ZD5S).

## Supplementary Information

**Supplementary Fig 1.**
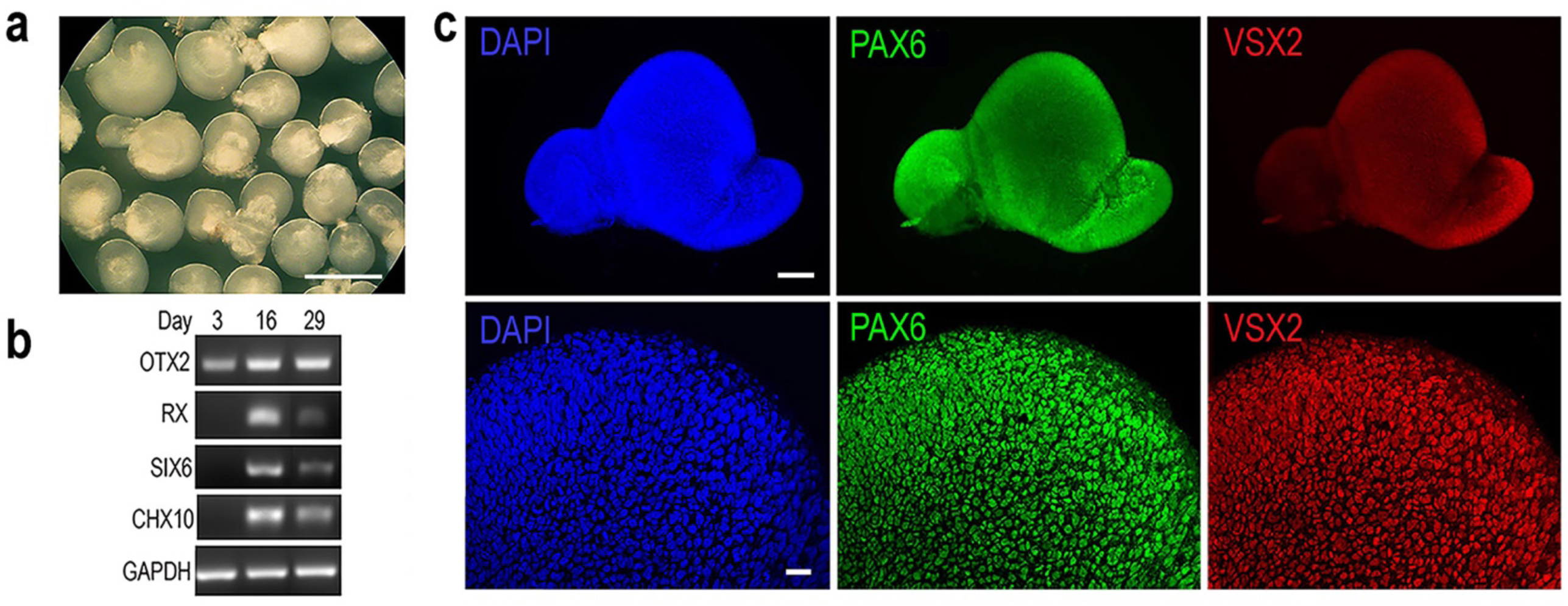
Characterization of human ES cell-derived 3D retinal organoids. **a** Bright field image shows morphology of a group of H9 ES cell-derived 3D retinal organoid at Day 33. Scale bar, 1 mm. **b** RT-PCR assay detects expression of eye field and neural retina genes at Day 16 and Day 29 in H9 ES cell-derived cultures. **c** Whole mount images show a 3D retinal organoid co-immunolabeled for PAX6 and VSX2 at Day 24. The top panels show low magnification images of the entire retinal organoid (scale bar, 100 μm), and the bottom panels show confocal images with nuclear labeling of PAX6 and VSX2 in retinal progenitor cells (scale bar, 20 μm).

**Supplementary Fig 2.**
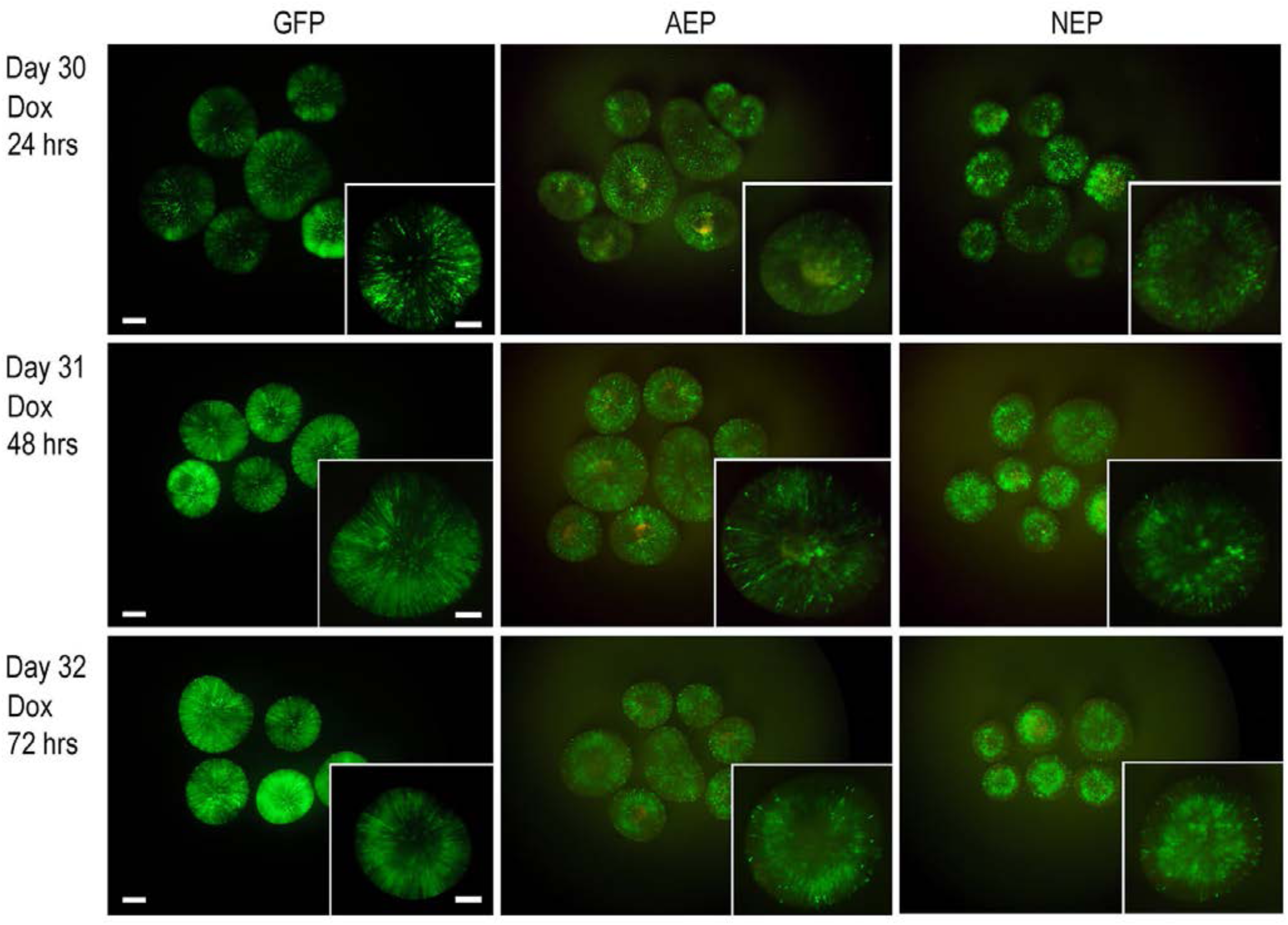
Effect of Dox induction time course on retinal organoid development. Whole mount images show effects of different Dox induction durations on retinal organoid development. After 24-hour of Dox induction, both LV-AEP and LV-NEP infected cells showed a tendency toward localizing to the inner layer, compared to the control LV-GFP virus infected retinal organoids. This trend became more pronounced after 48-hour Dox induction. By 72-hour after the onset of Dox treatments, the majority of GFP^+^ cells were concentrated in the inner layer of the retinal organoids. In contrast, most LV-GFP infected cells remained a ventricular zone distribution pattern after 48- and 72-hour induction. Scale bars, 200 μm for the lower magnification; 100 μm for the insets.

**Supplementary Fig 3.**
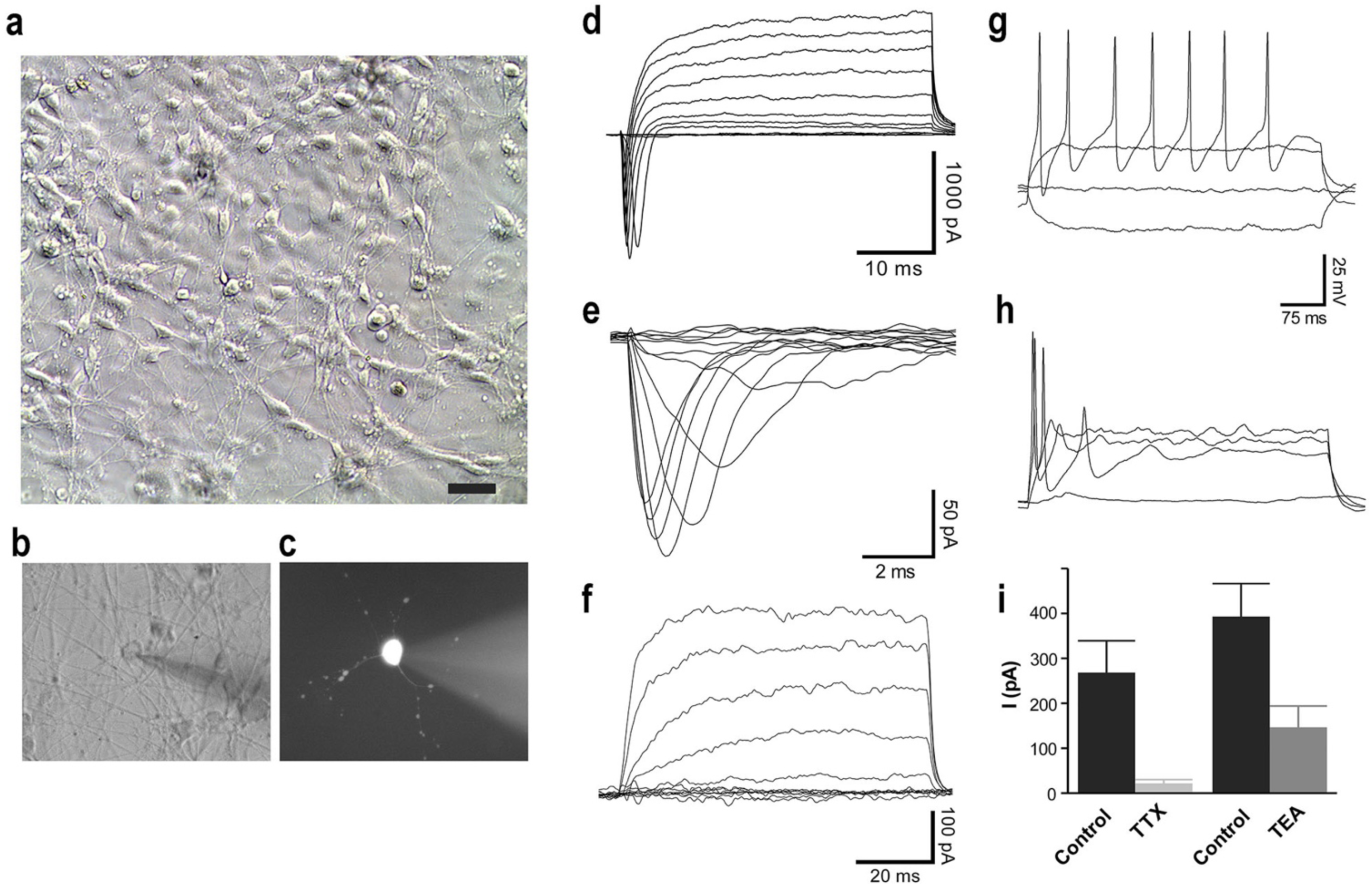
Electrophysiological properties and functionality of Human H9 ES cell-derived retinal neurons. **a** Bright field view of the dissociated cell culture derived from retinal organoids at Day 40. Scale bar, 20 μm. **b, c** A neuron being patch recorded is shown in bright field (**b**) and after filling with lucifer yellow dye (**c**). **d -i** Whole cell patch clamp recording of dissociated neurons (Day 37-40) derived from H9 ES retinal organoids cultured as a monolayer. **d** Whole cell voltage clamp of a cell with multipolar neurites stepped from -60 mV to +30 mV in 10 mV steps of 40 ms duration. **e** Example of well-clamped I_Na_ isolated by digital substraction following block with 100 nM TTX. Steps from -90 to +20 mV are shown. **f** Outward K^+^ currents isolated by digital substraction following 10 mM TEA application. Steps from -70 to +30 mV are shown. **g** Train of action potentials elicited with depolarizing current in current clamped cells having large I_Na._ **h** Phasic action potential generation in cells having smaller I_Na._ **i** Summary of block of peak I_Na_ and I_K_ at +40 mV by TTX and TEA, respectively. TTX blocked 92% of the transient inward current (n=7) and TEA blocked 62% of the sustained outward current at +40 mV (n=6).

**Supplementary Fig 4.**
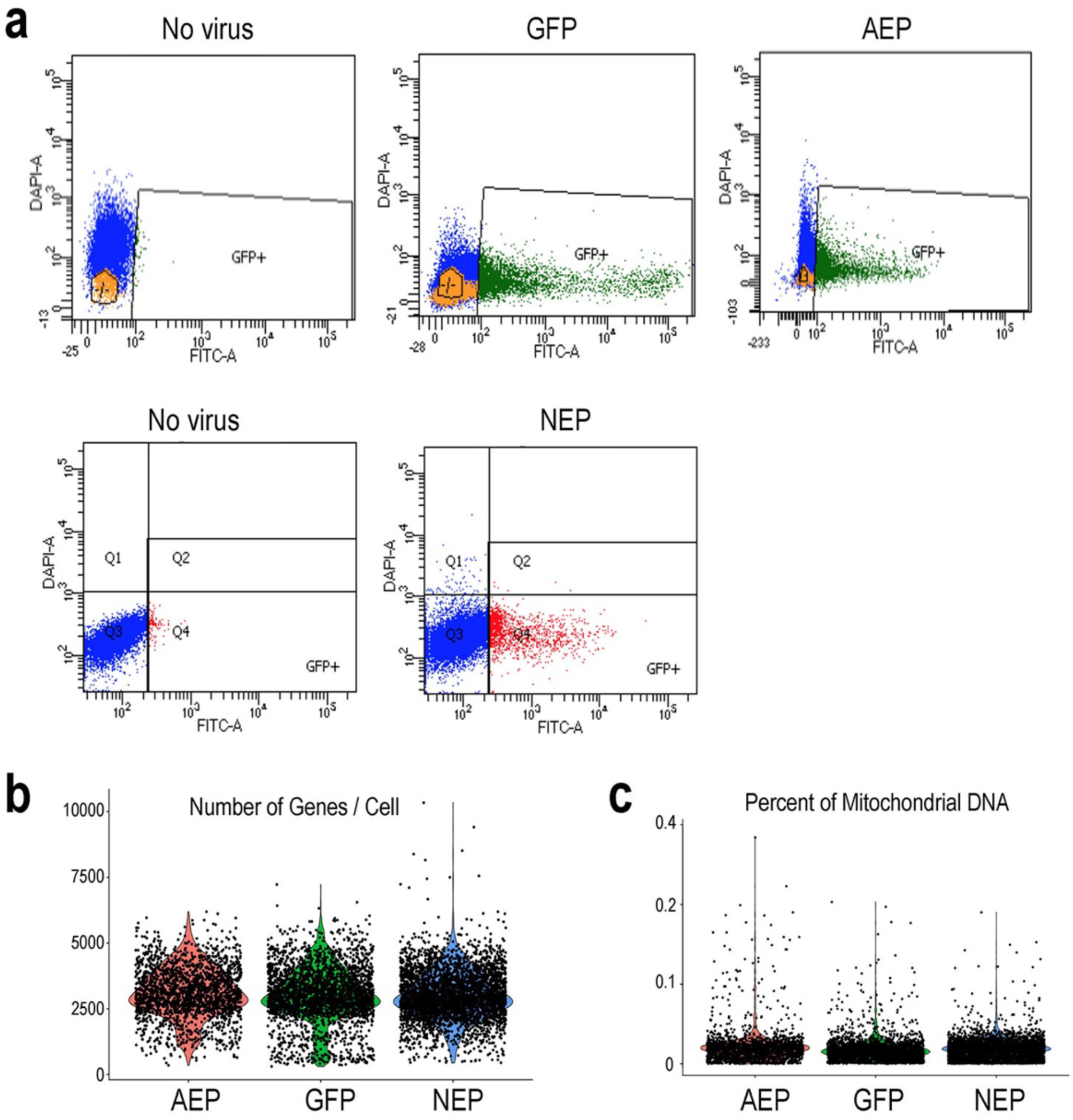
Fluorescent activated cell sorting of lentivirus infected retinal organoids for single cell RNA-sequencing. **a** FACS profiles of dissociated retinal organoids infected with LV-GFP, LV-AEP, and LV-NEP between Day 45-48 after 8 day ox treatment. Cells from non-infected retinal organoids were used as negative controls to set the thresholds for GFP^+^ cells. FACS enriched GFP^+^ cells were used for single cell RNA-seq analyses. **b** Violin plots show the numbers of genes detected in single cell RNA-seq analyses using 10X Genomics Chromium and NovaSeq work flow. The cutoff used in this study was 2500 UMI per cell, resulting in mean gene per cell ranging from 2935-3079. For downstream analysis 3004 cells for LV-GFP, 2063 cells for LV-AEP, and 3909 cells for LV-NEP were used. **c** Violin plots show percentage of mitochondrial encoded genes detected in single cell RNA-seq analyses using 10X Genomics Chromium and NovaSeq work flow. The low rates (<0.03%) of transcripts from the mitochondrial genome indicate that the transcripts analyzed in this study are from the nuclear genome.

**Supplementary Fig 5.**
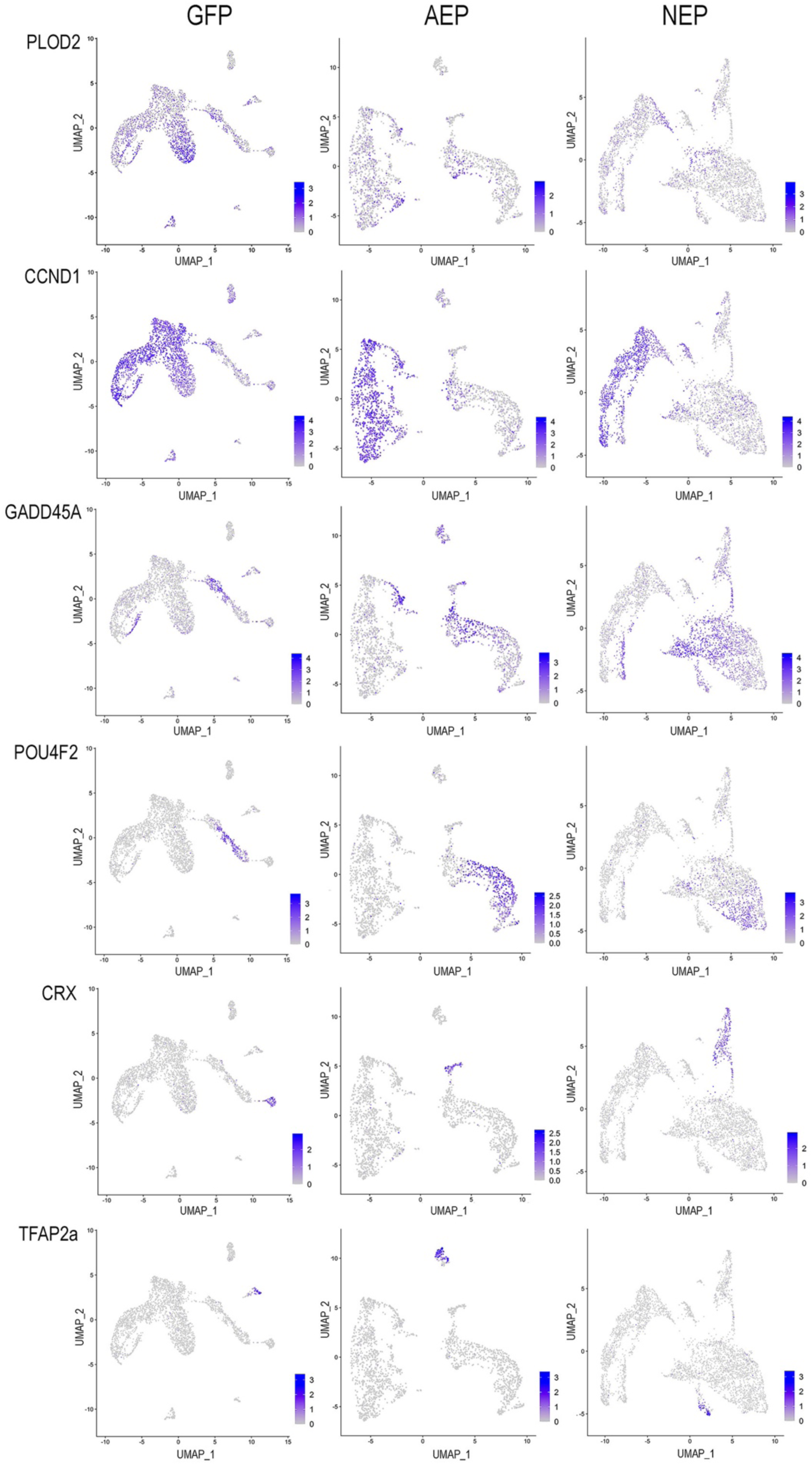
Expression of known genes used to assign cell categories of cell clusters. Feature plots of known genes in LV-GFP, LV-AEP, or LV-NEP infected retinal organoids shown as UMAPs. Genes representing different cell category or states are used to assign cell cluster identities in Figure 5.

**Supplementary Fig 6.**
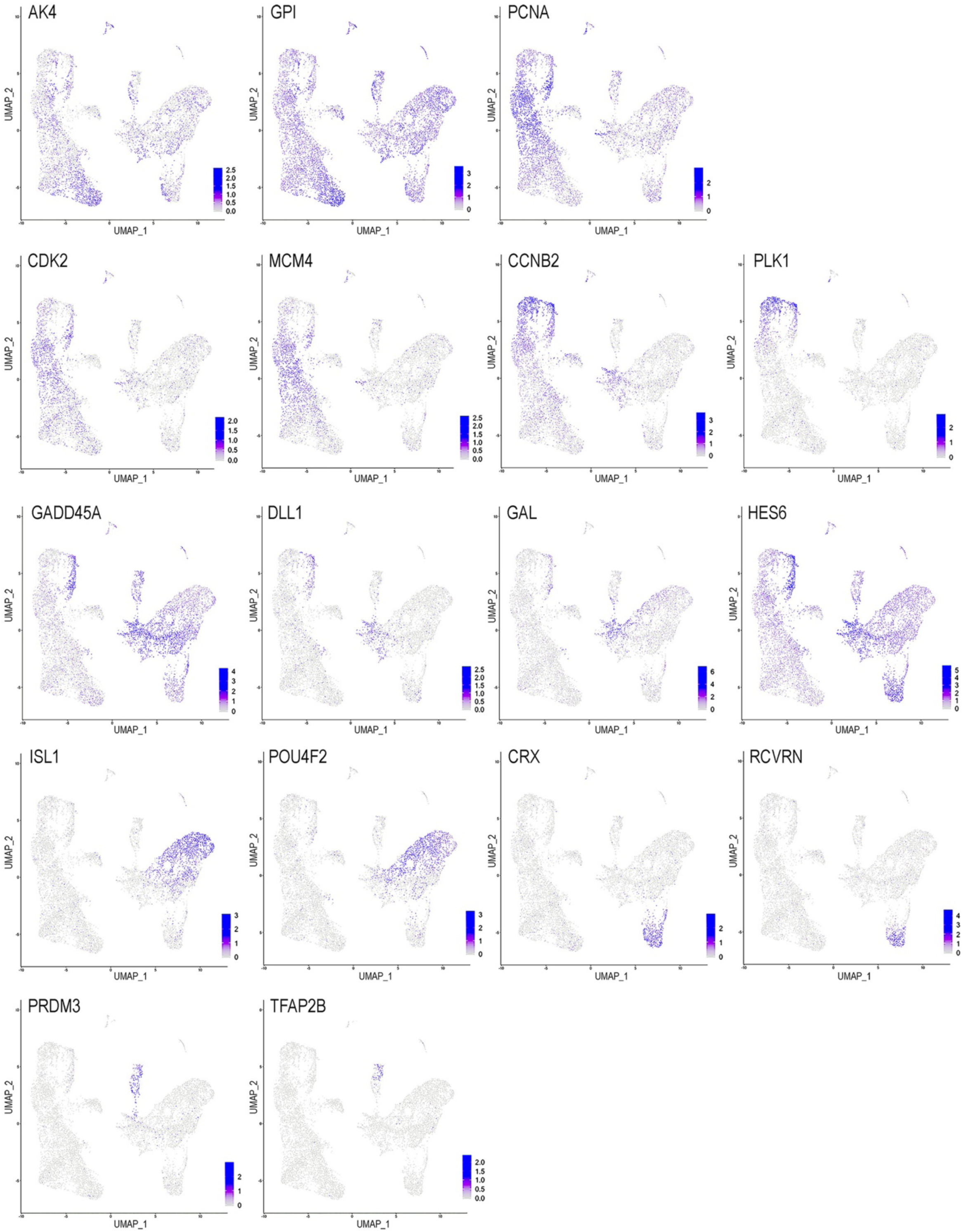
Expression of featured known genes in combined sample clusters. Feature plots of known genes in the combined LV-GFP, LV-AEP, LV-NEP sample clusters shown as UMAPs. Genes representing different cell cycle phases and cell type categories are used to assign cluster identities in Figure 8.

**Supplementary Fig 7.**
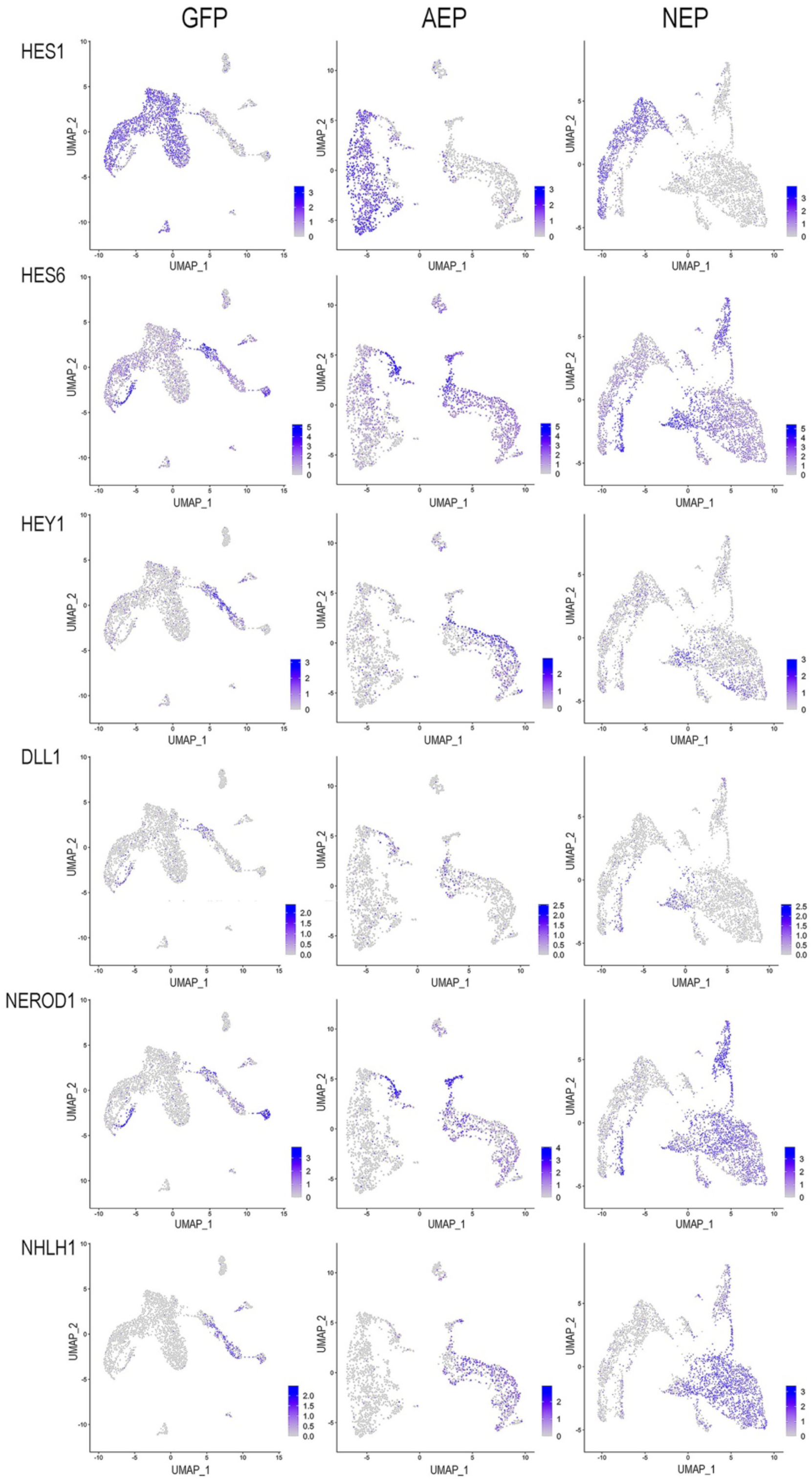
Expression patterns of Notch signaling components and selected bHLH genes in lentiviral infected retinal organoid cells. Feature plots of selected genes quantified in Figure 9 are shown in UMAPs in LV-GFP, LV-AEP, and LV-NEP infected retinal organoid cells. HES1 Is predominantly expressed among neural stem cells and progenitors. HES6, DLL1, and NEUROD1 are upregulated in exiting progenitors and neuroblasts. NHLH1 is expressed among postmitotic cells.

**Supplementary Fig 8.**
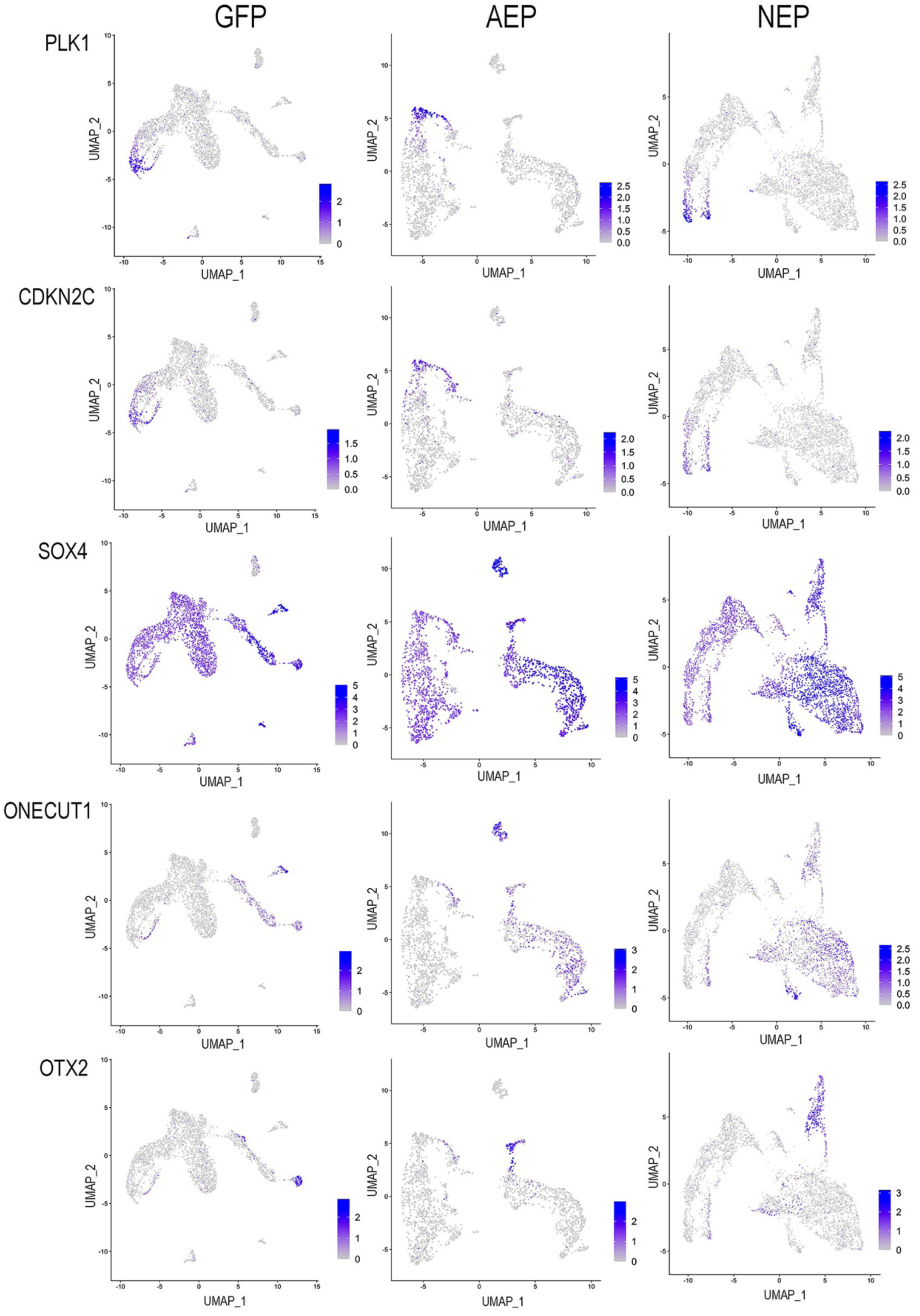
Expression patterns of genes involved in cell cycle exit and early retinogenesis. Feature plots of selected genes involved in cell cycle exit (PLK1, CDKN2C) and differentiation of early retinal neurons (SOX4, ONECUT1, OTX2) are shown in UMAPs in LV-GFP, LV-AEP, and LV-NEP infected retinal organoid cells.

**SupTable 1.**
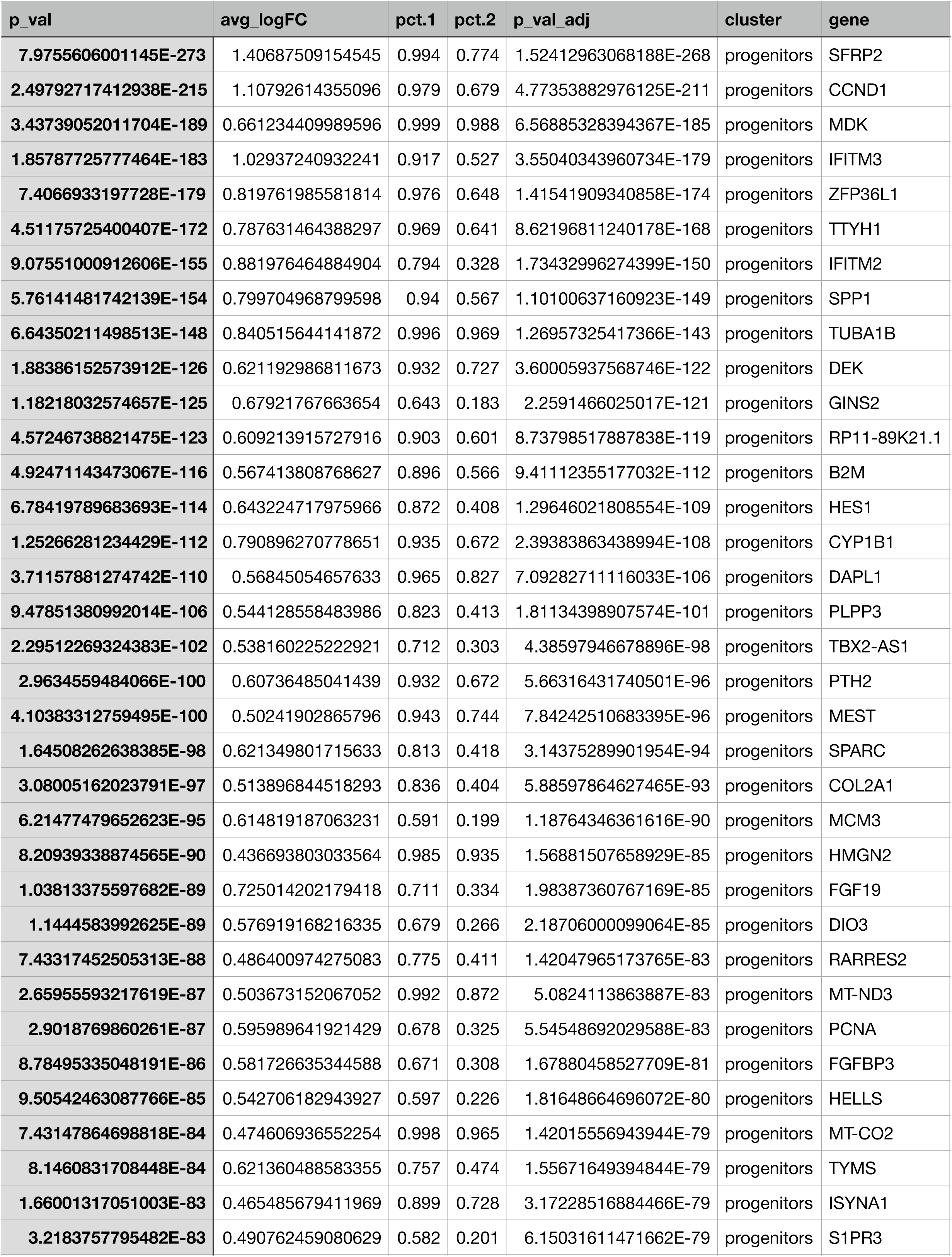
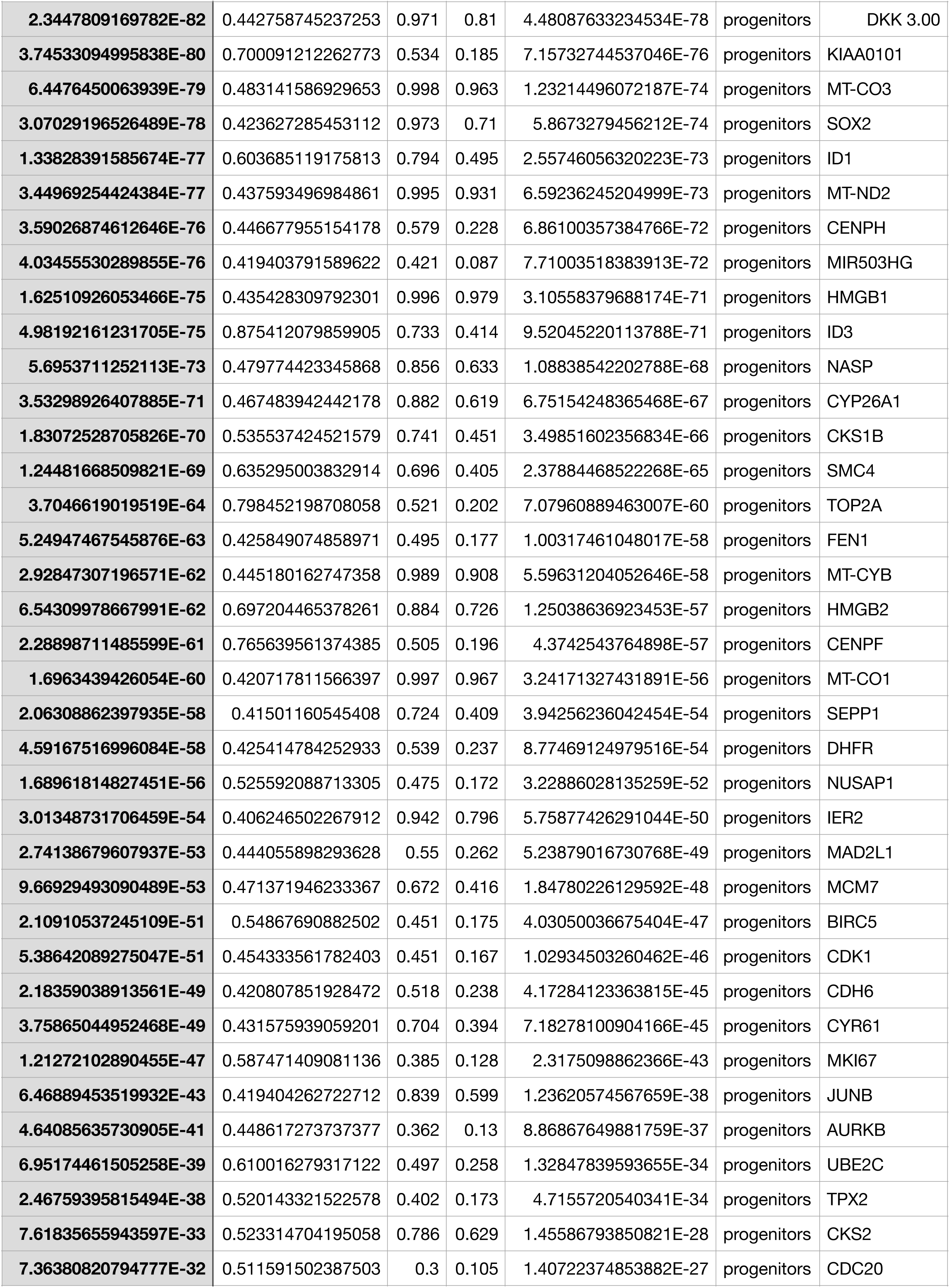
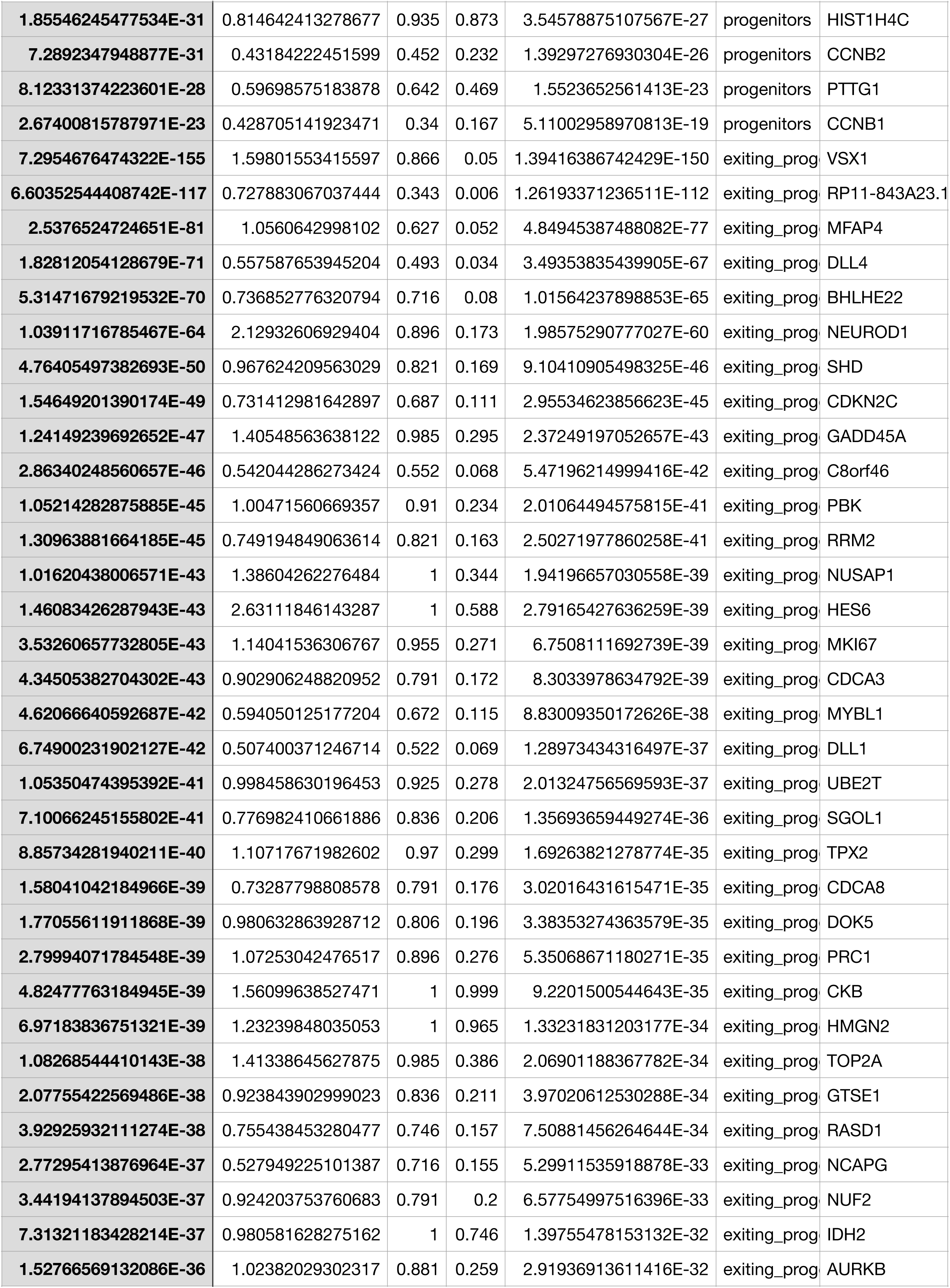
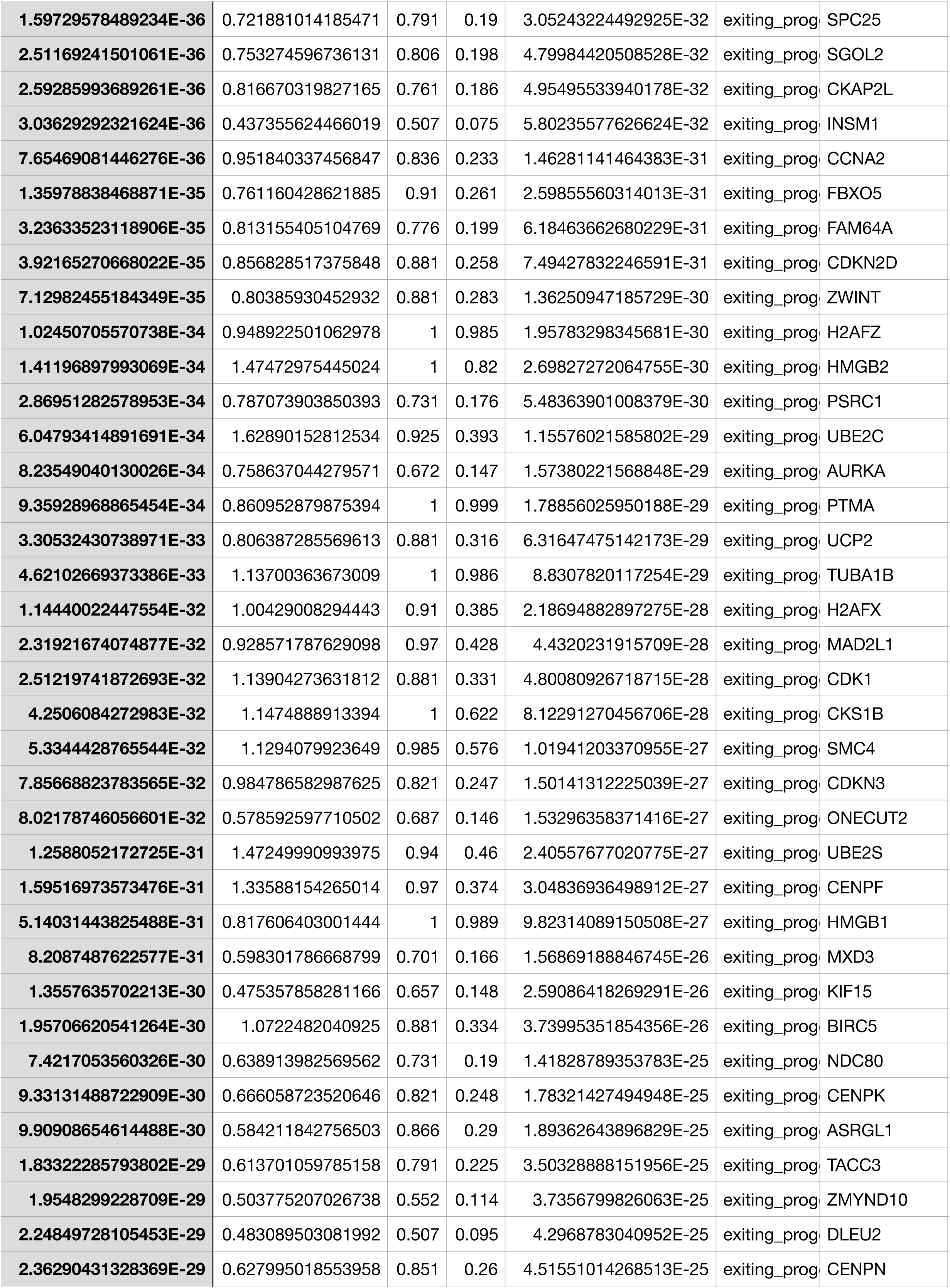
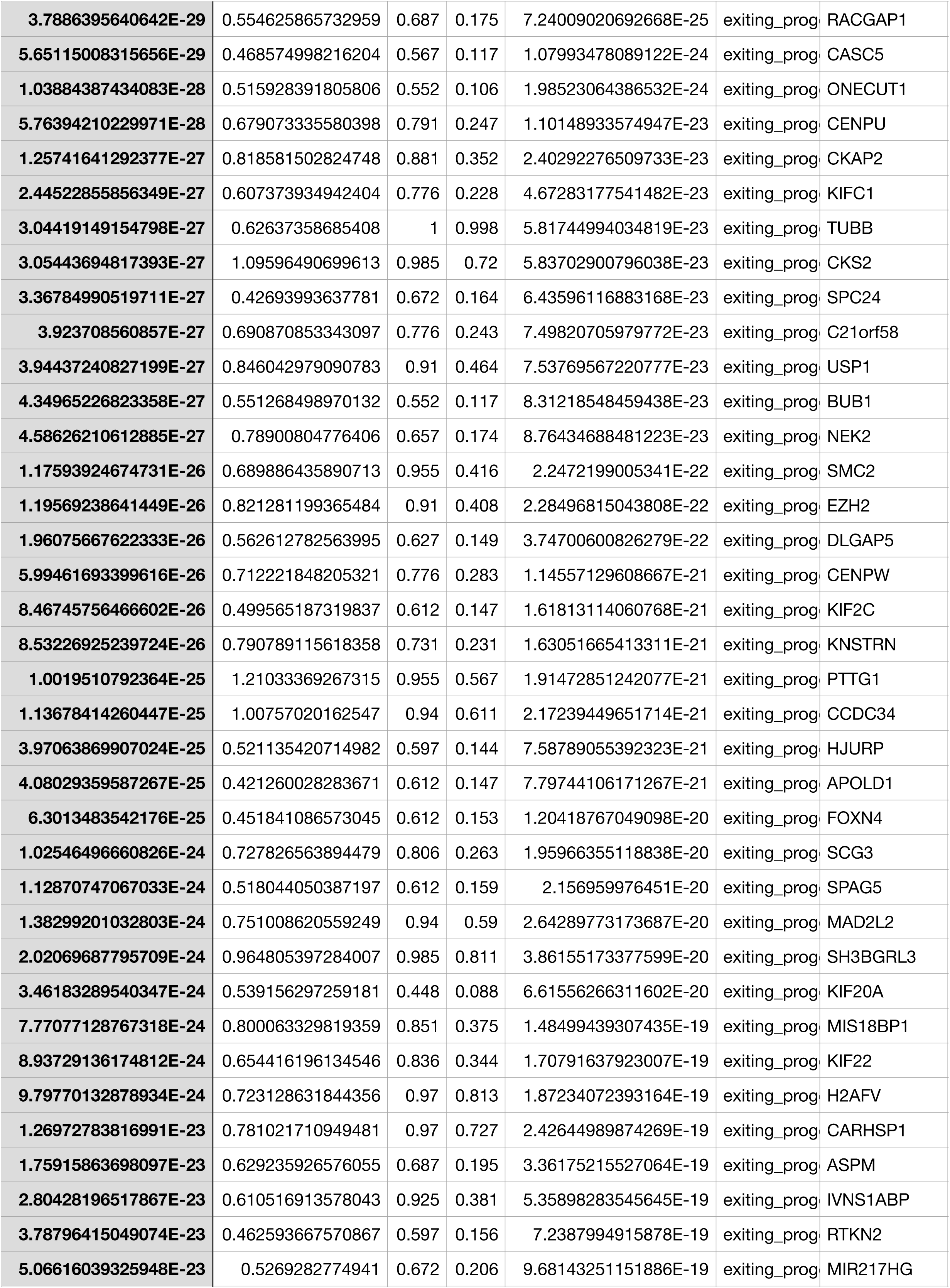
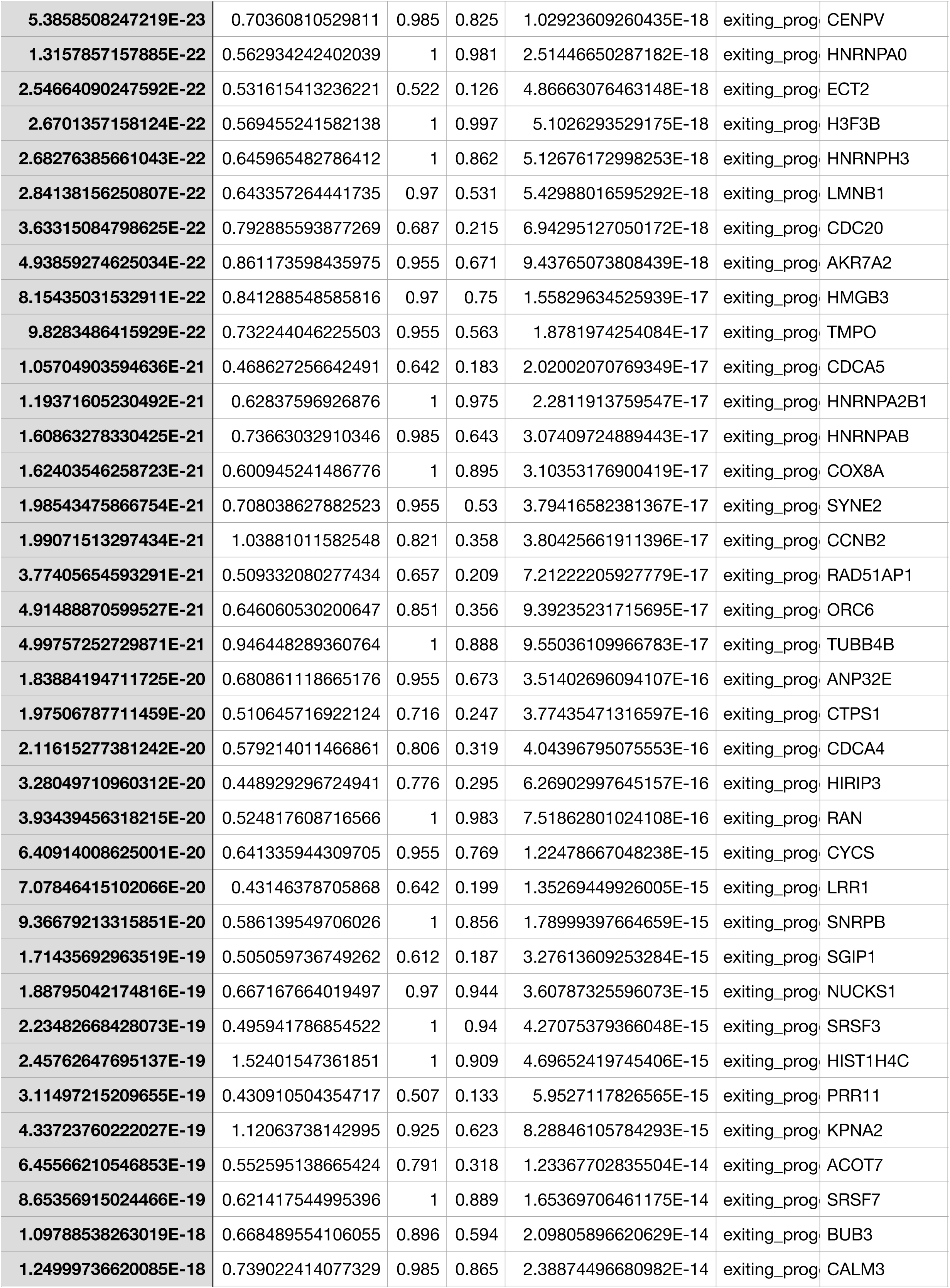
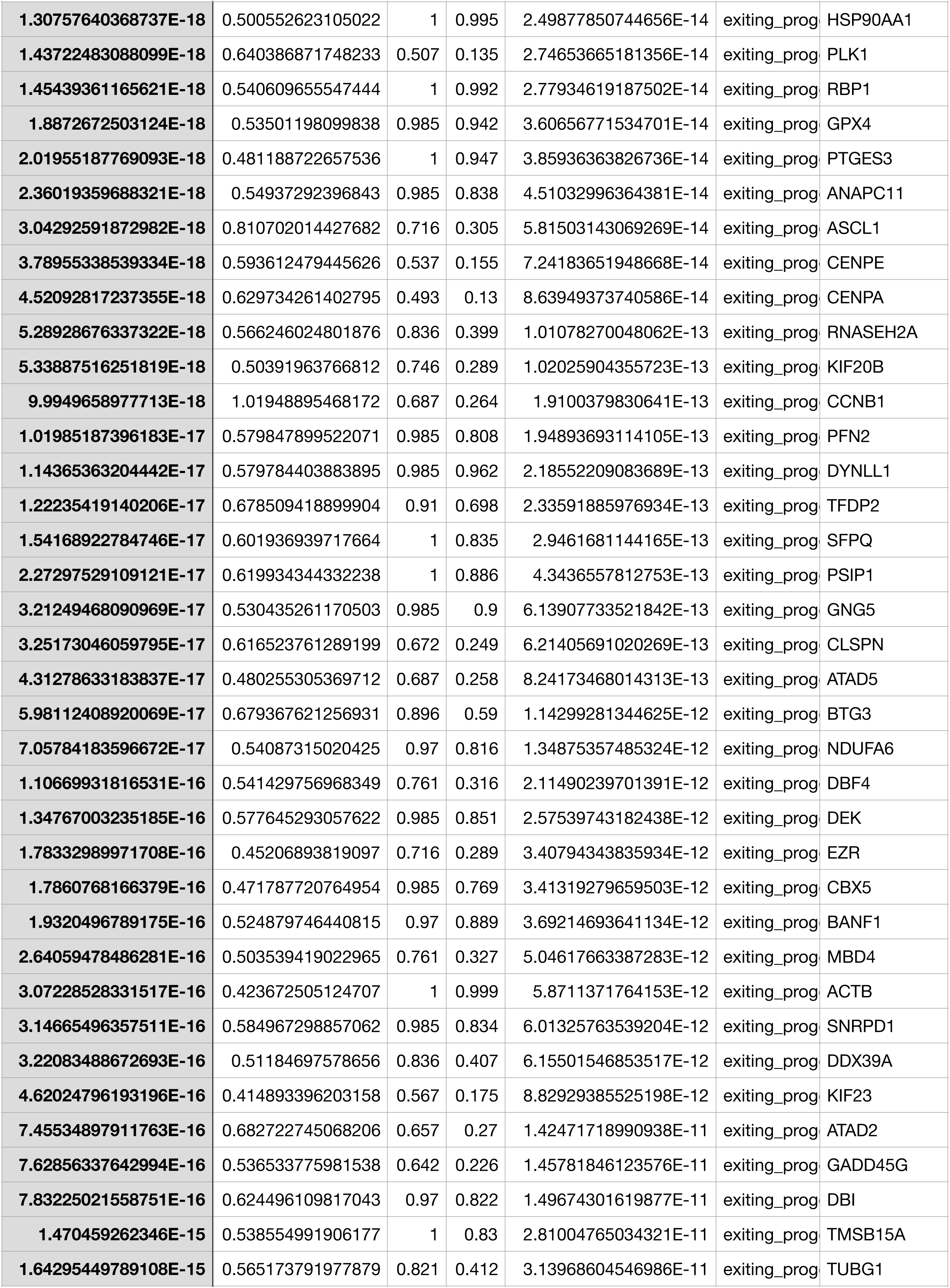
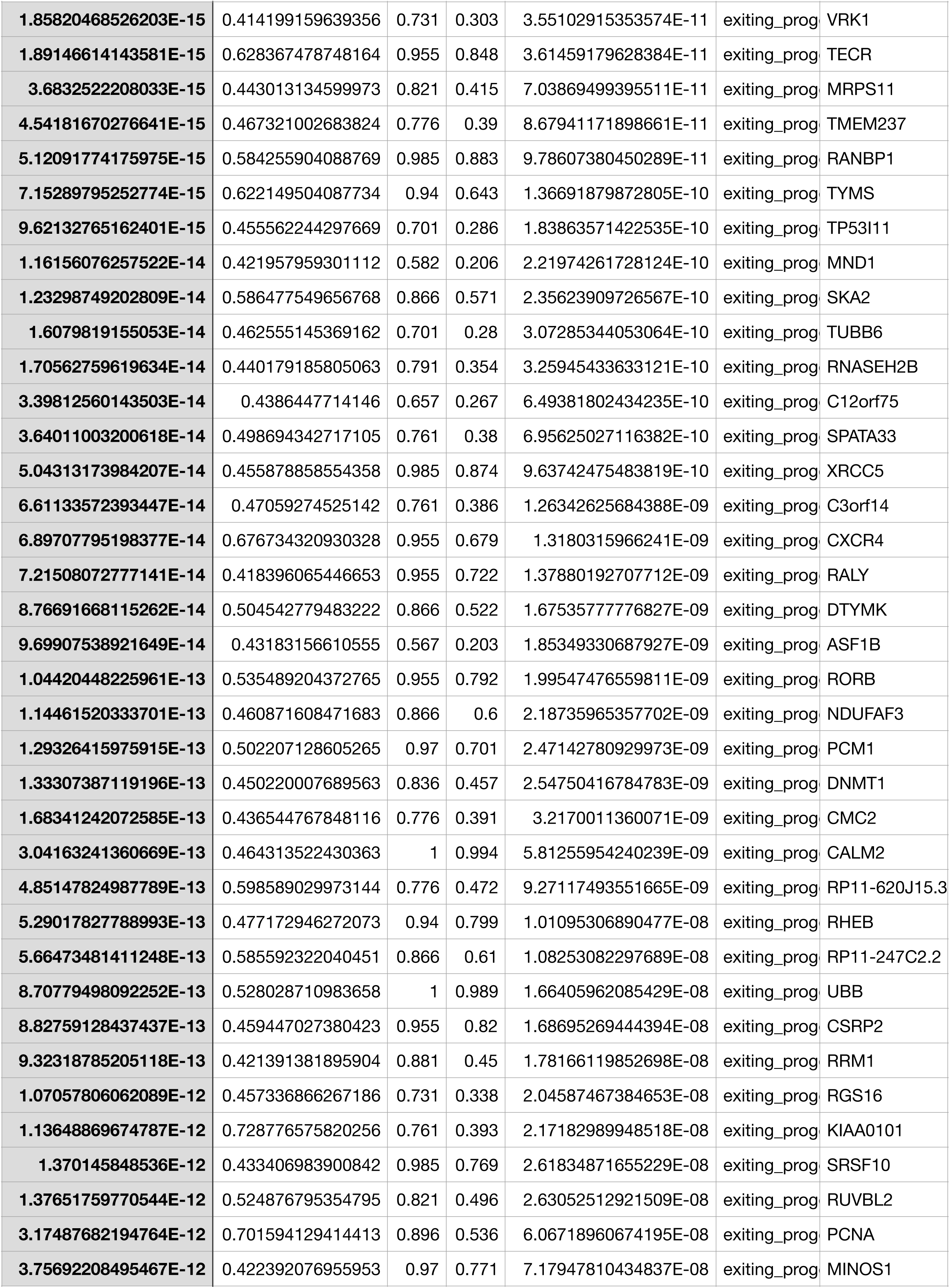
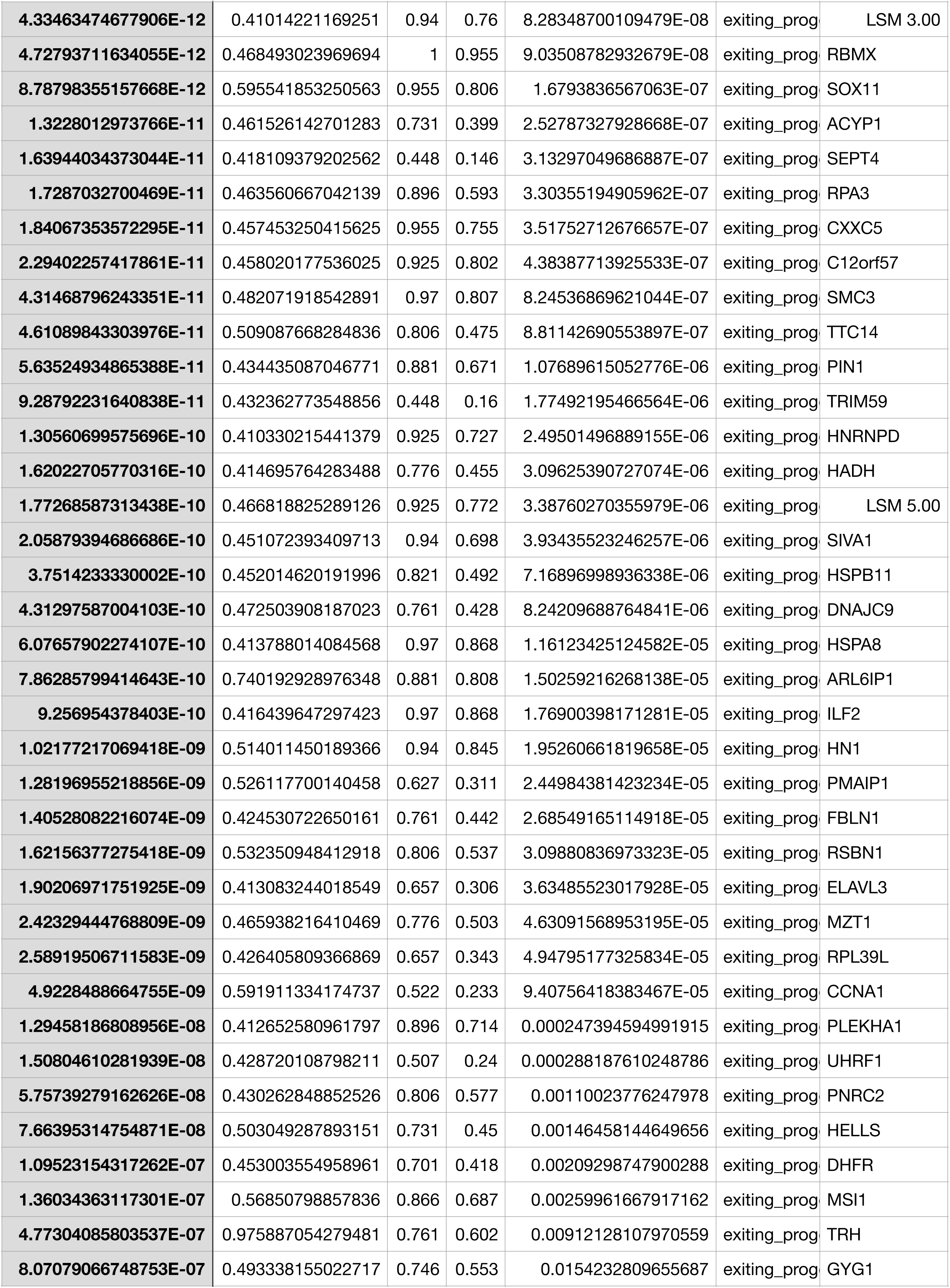
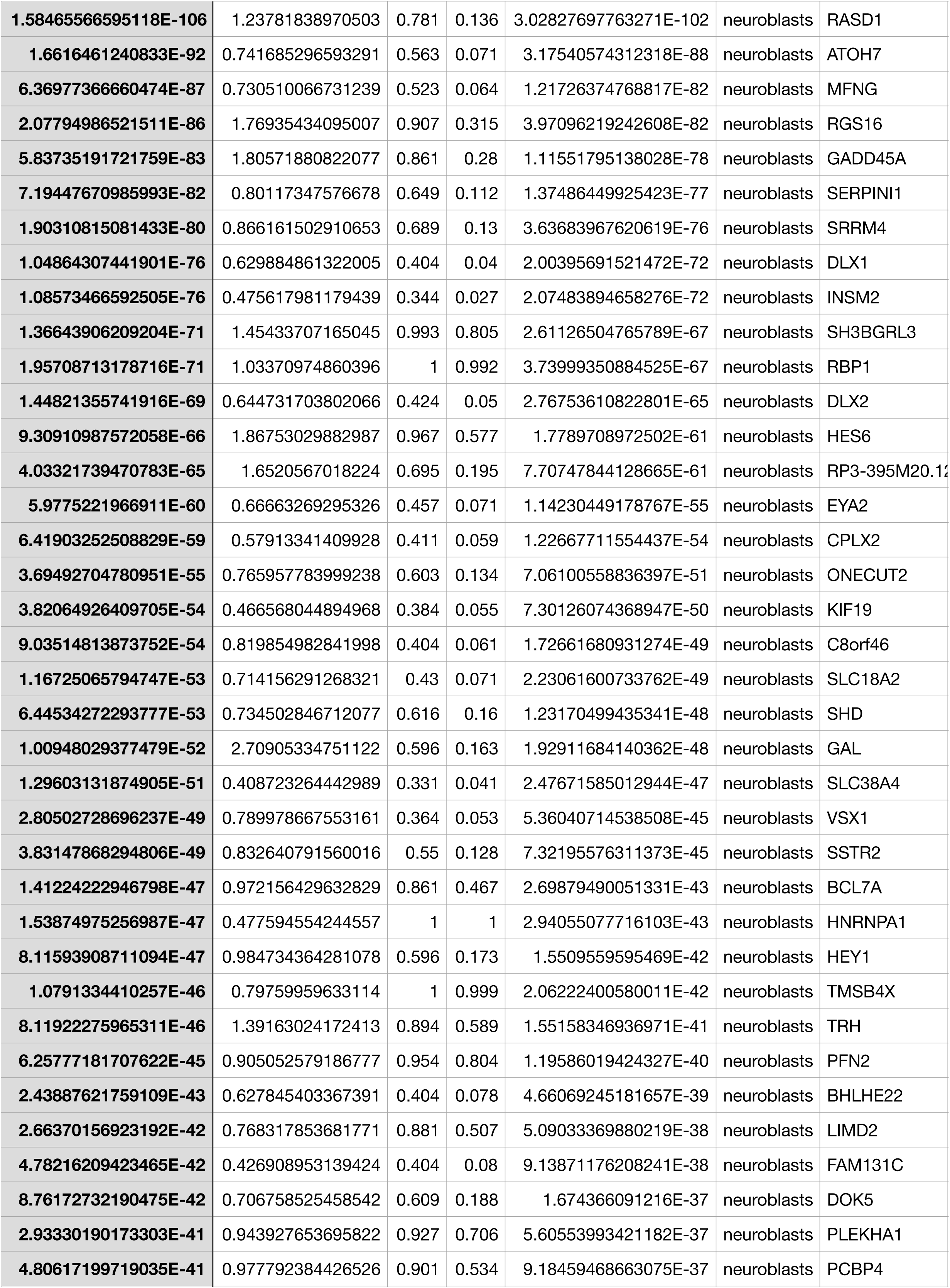
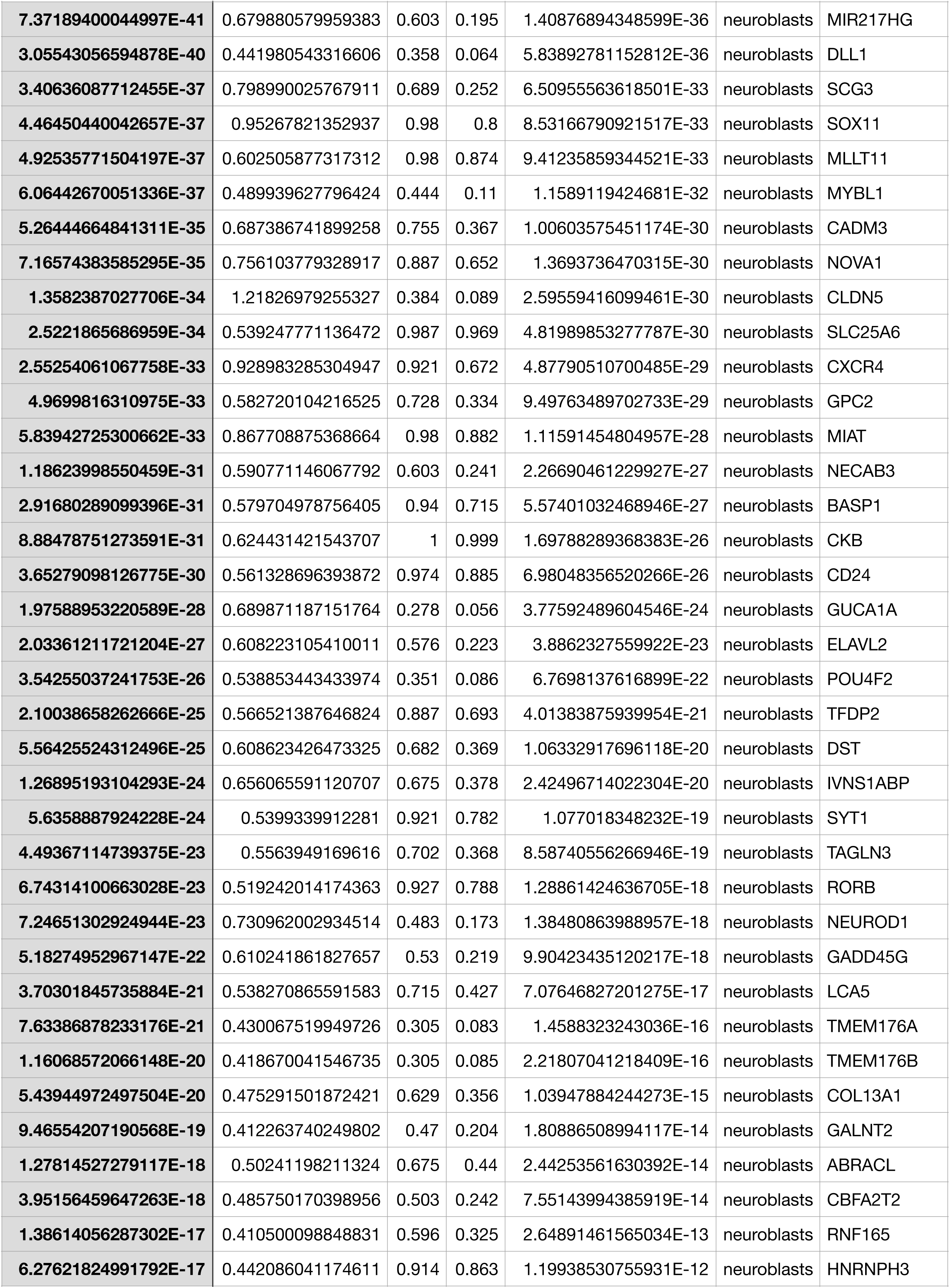
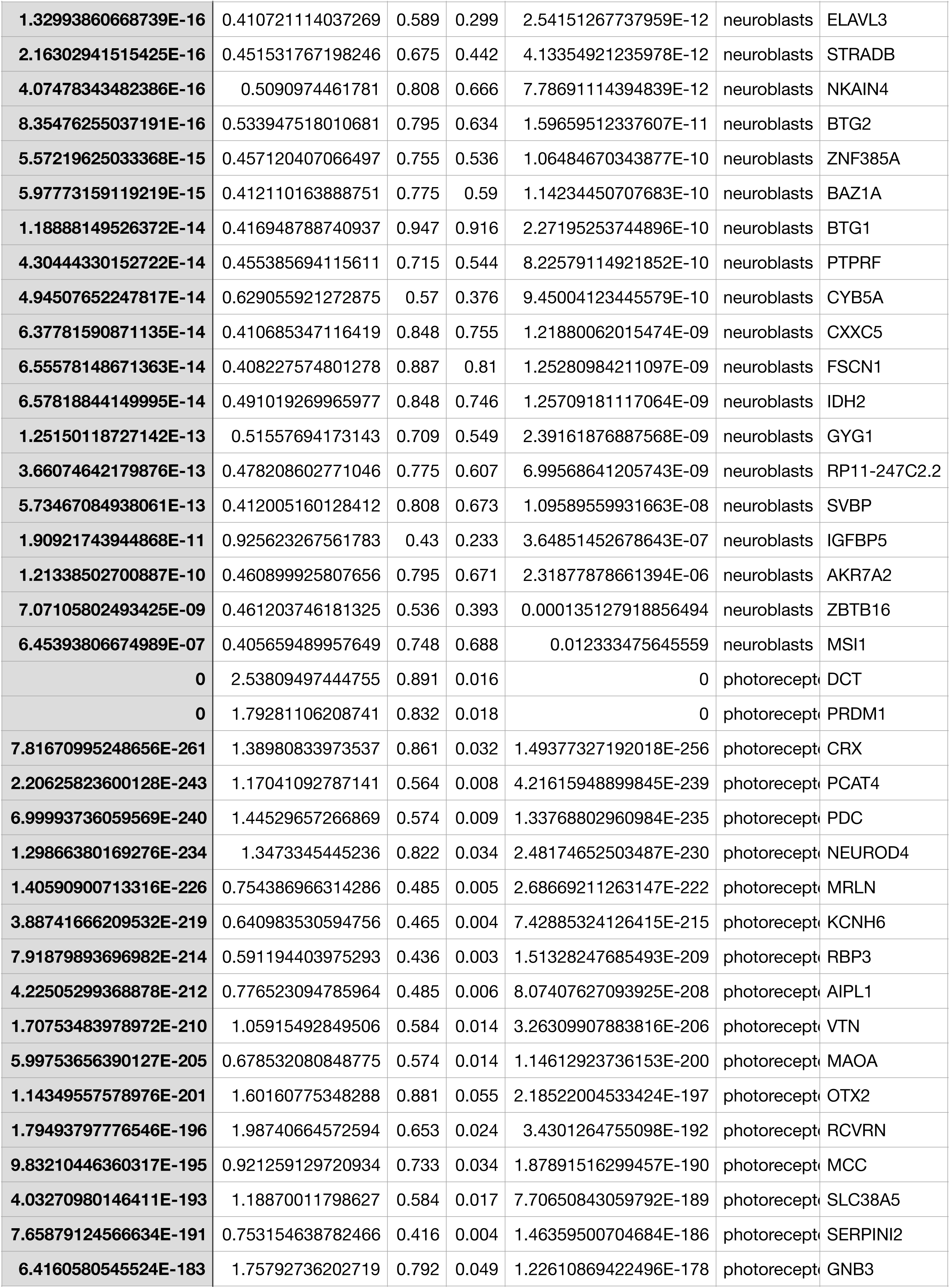
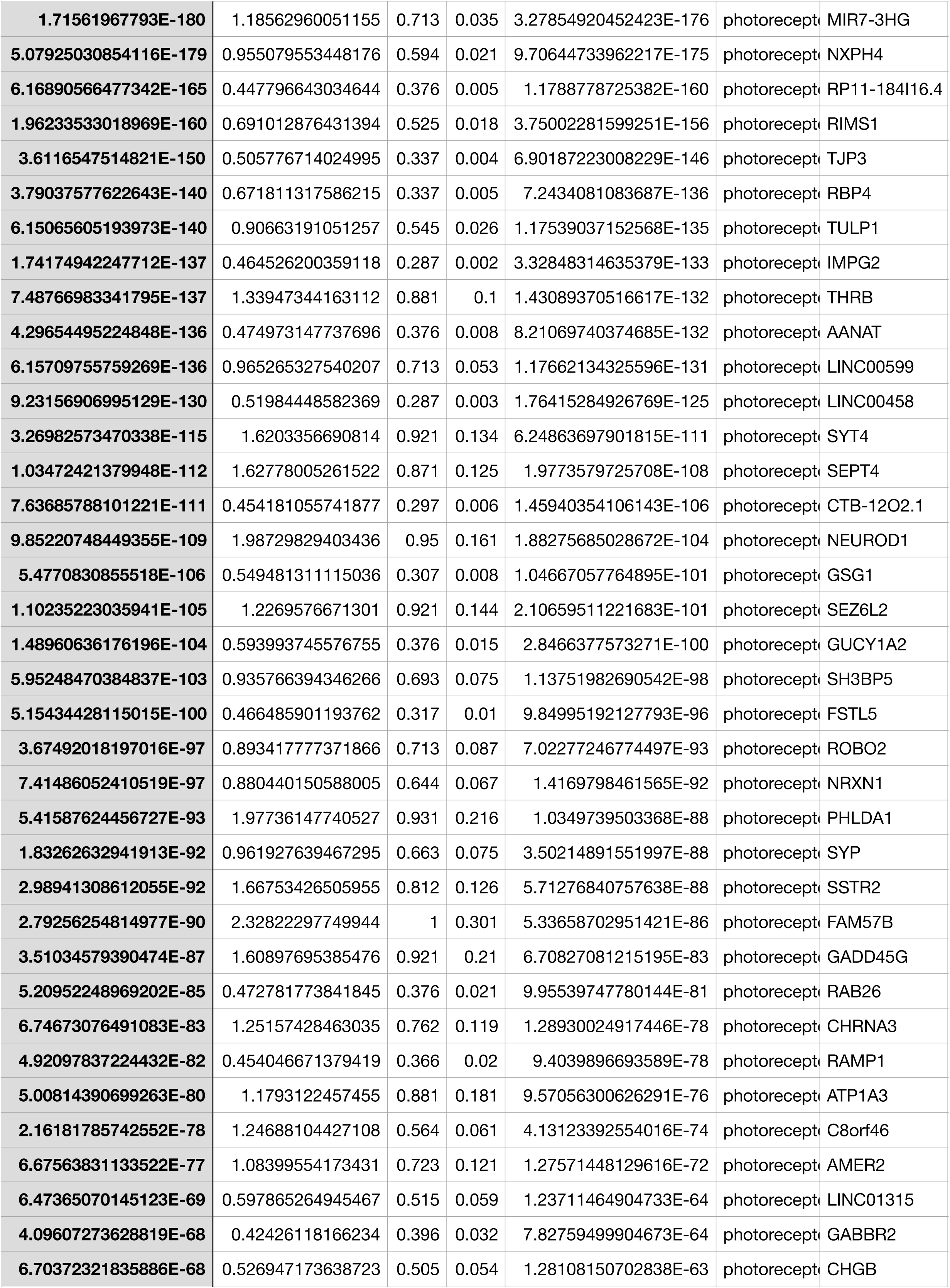
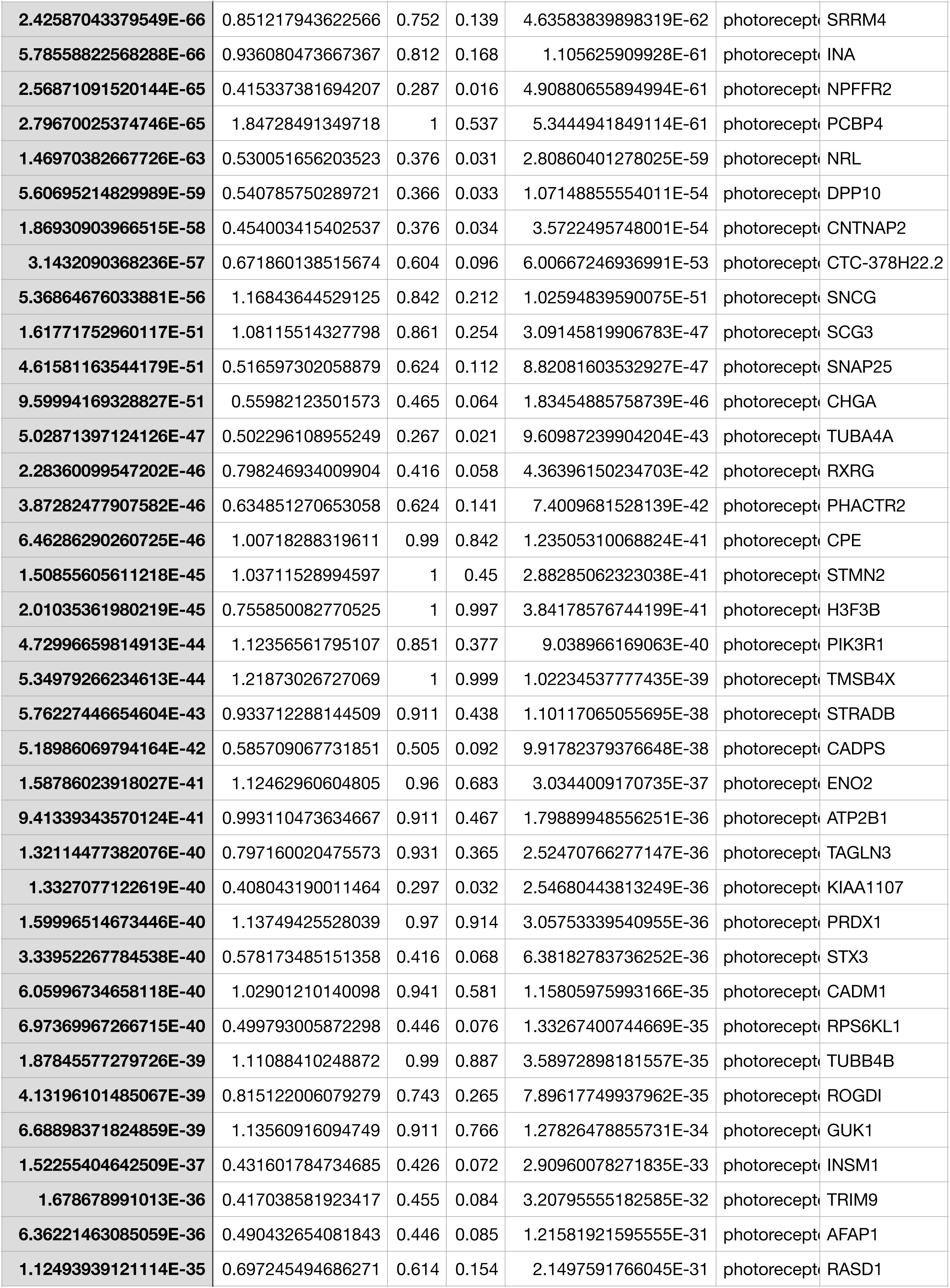
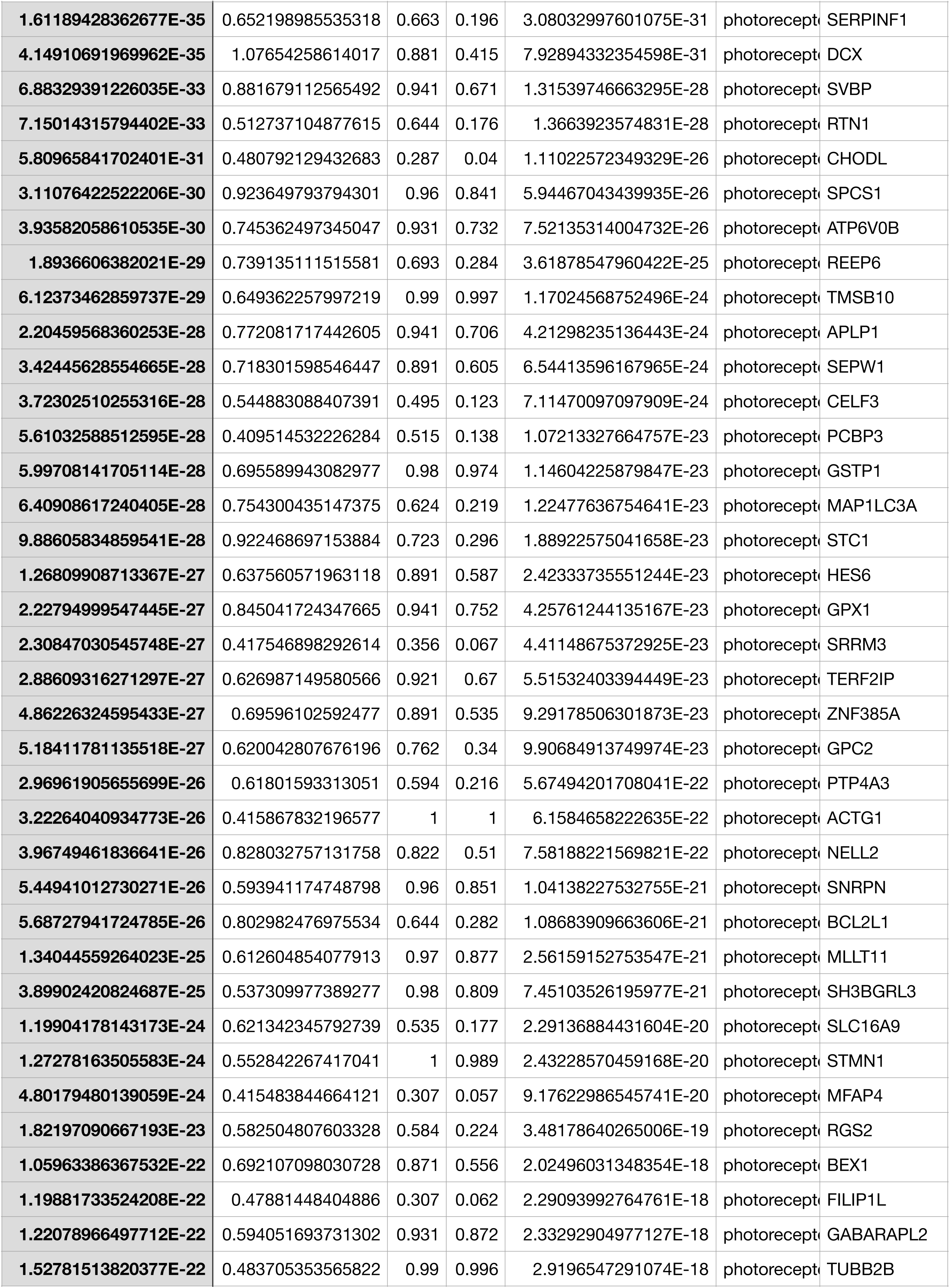
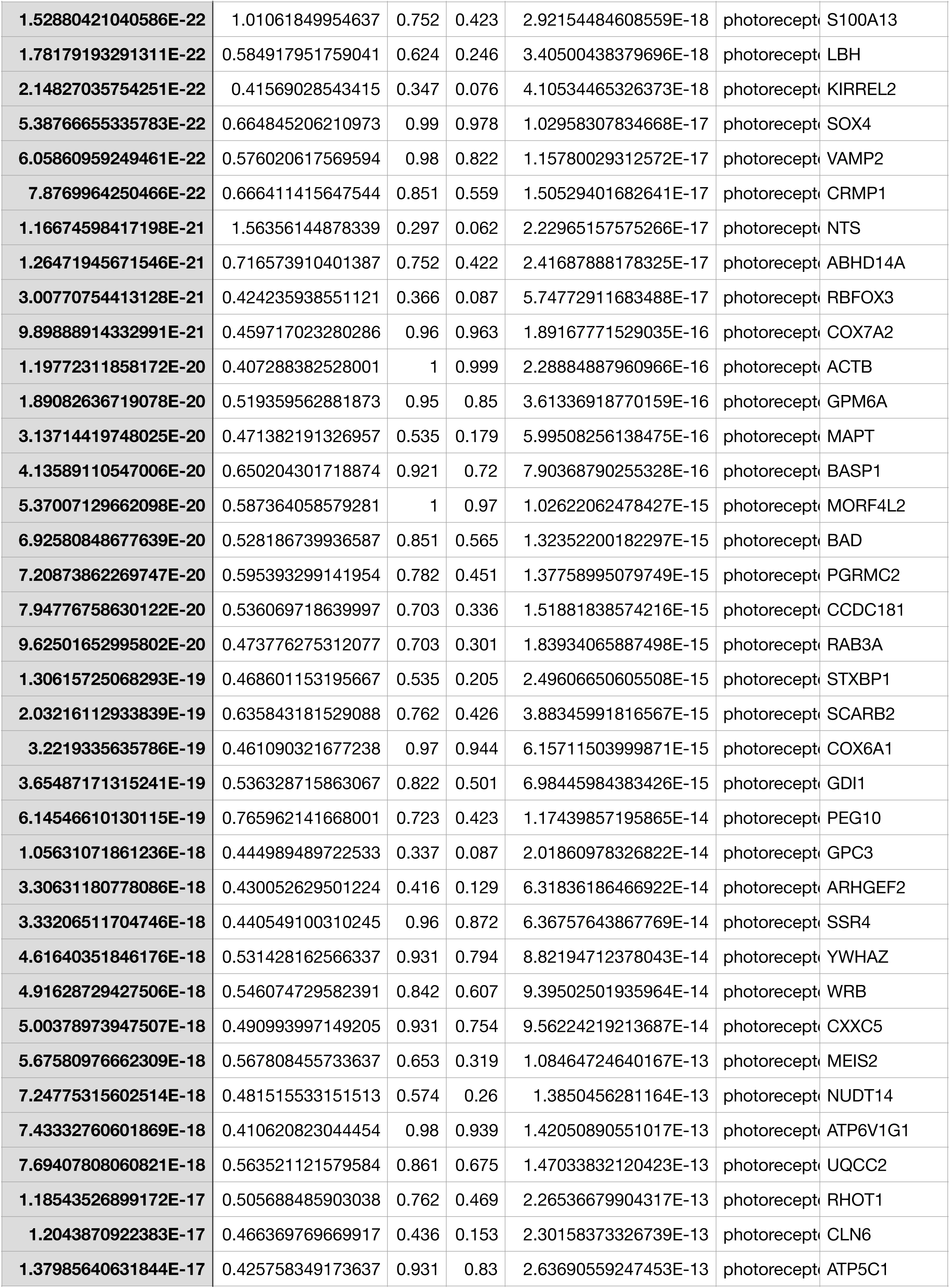
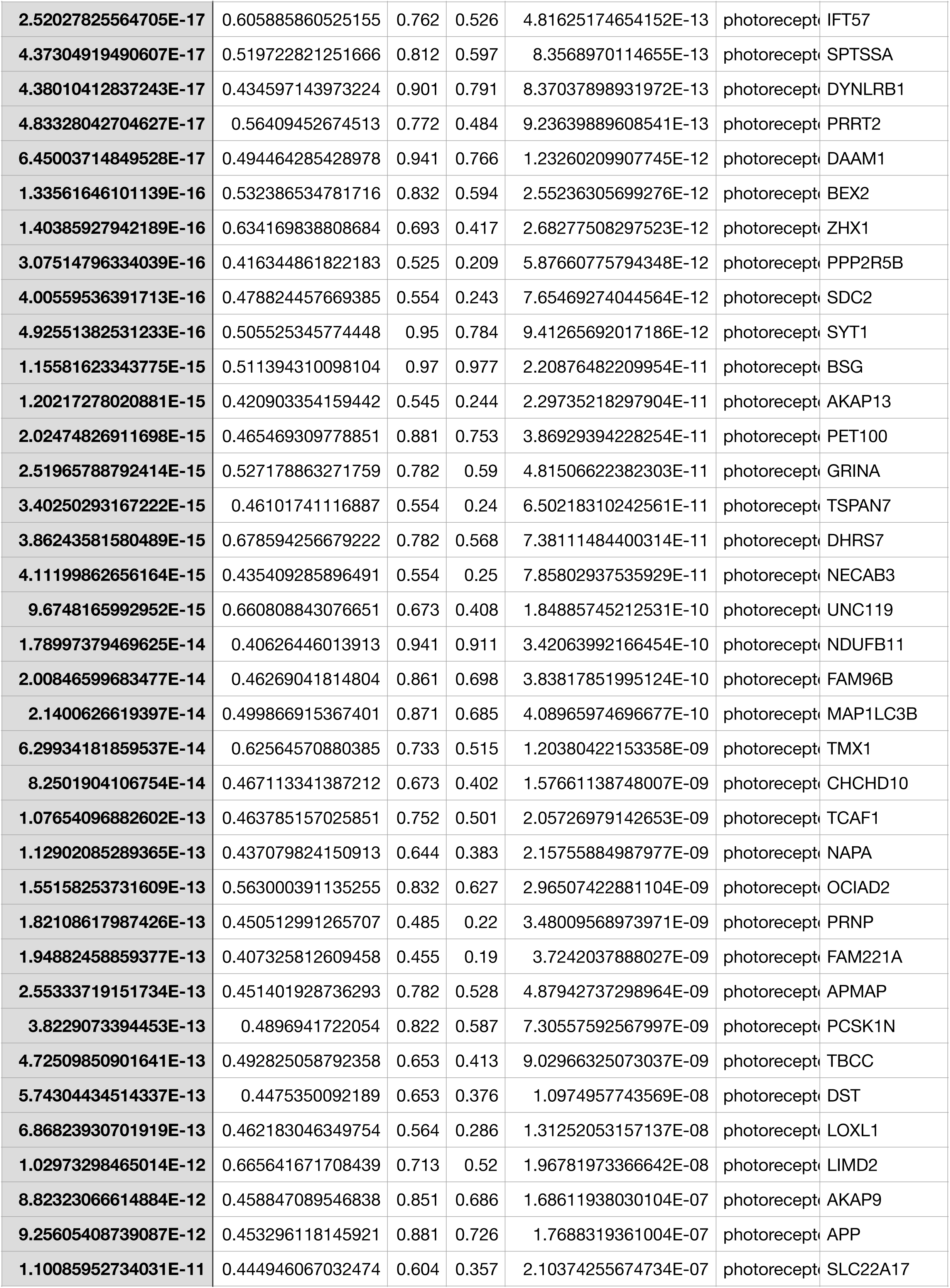
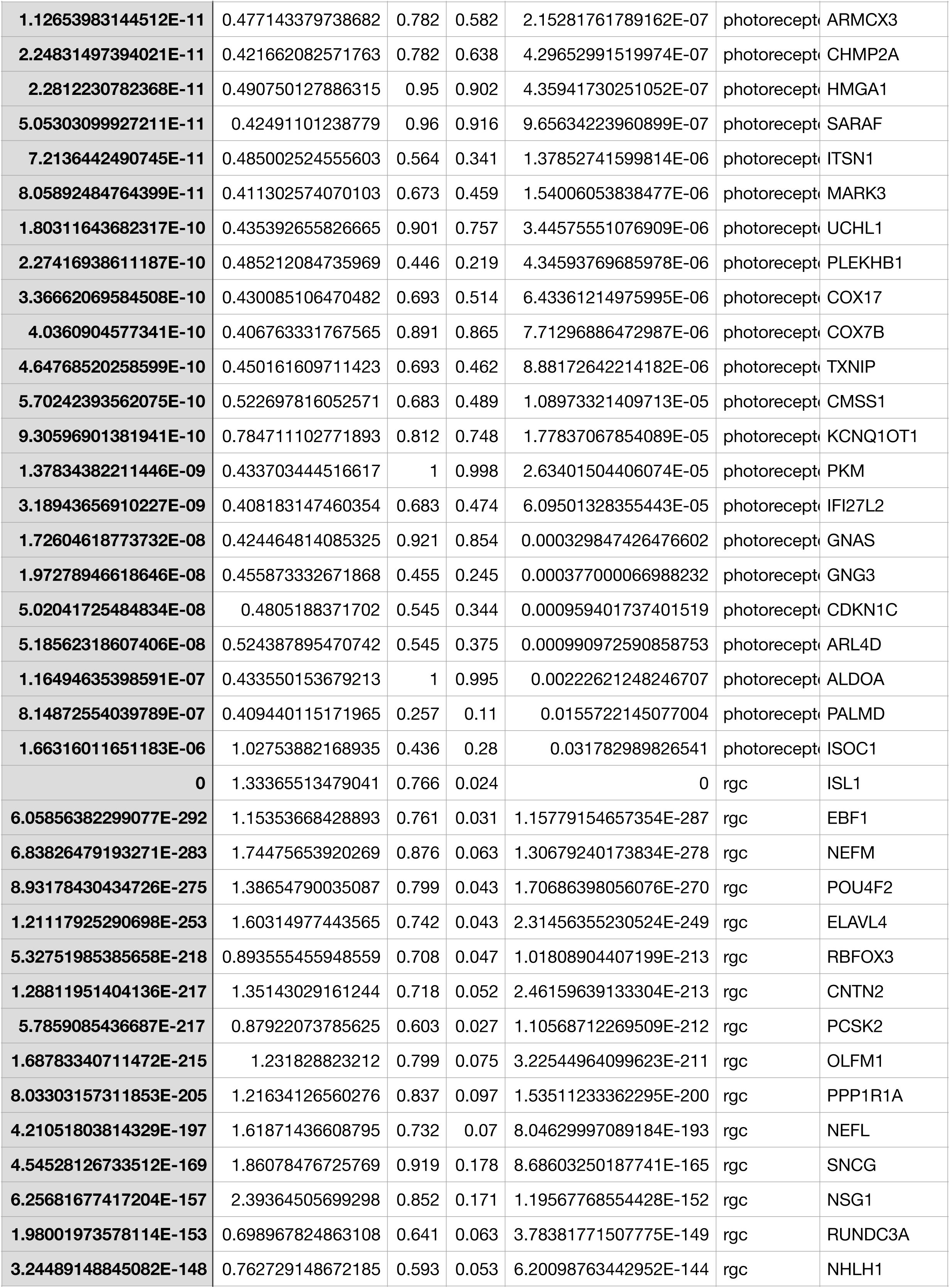
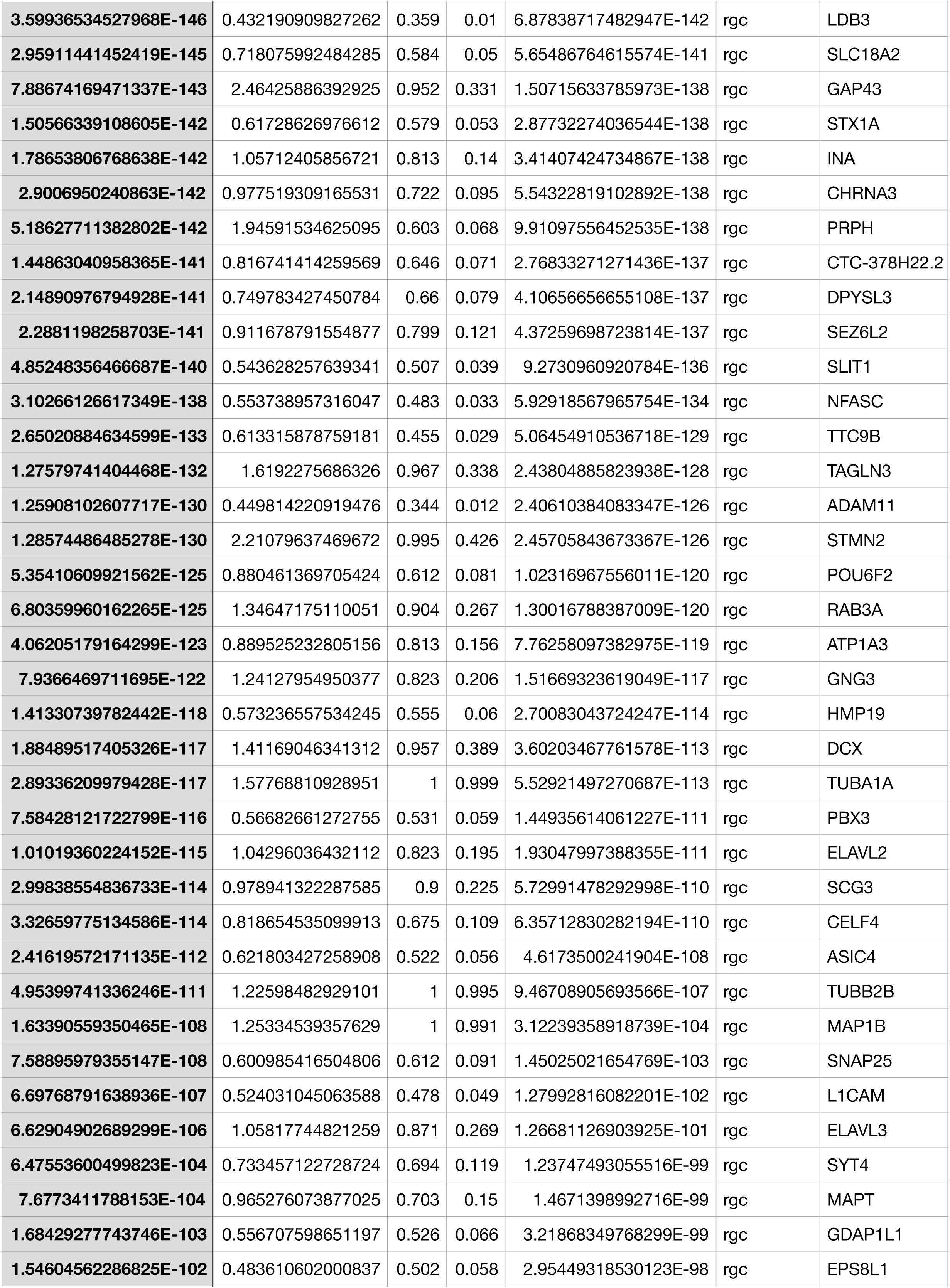
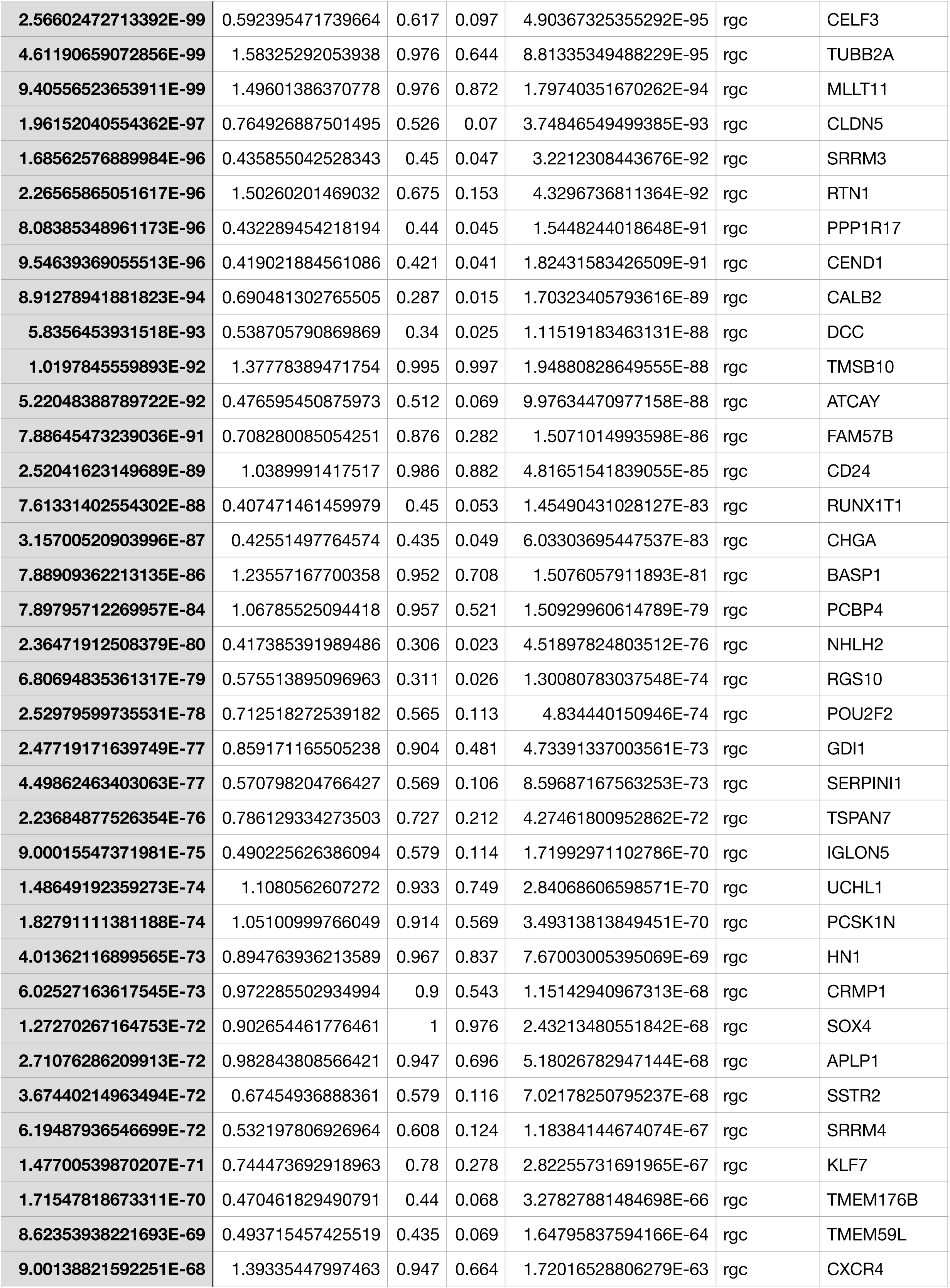
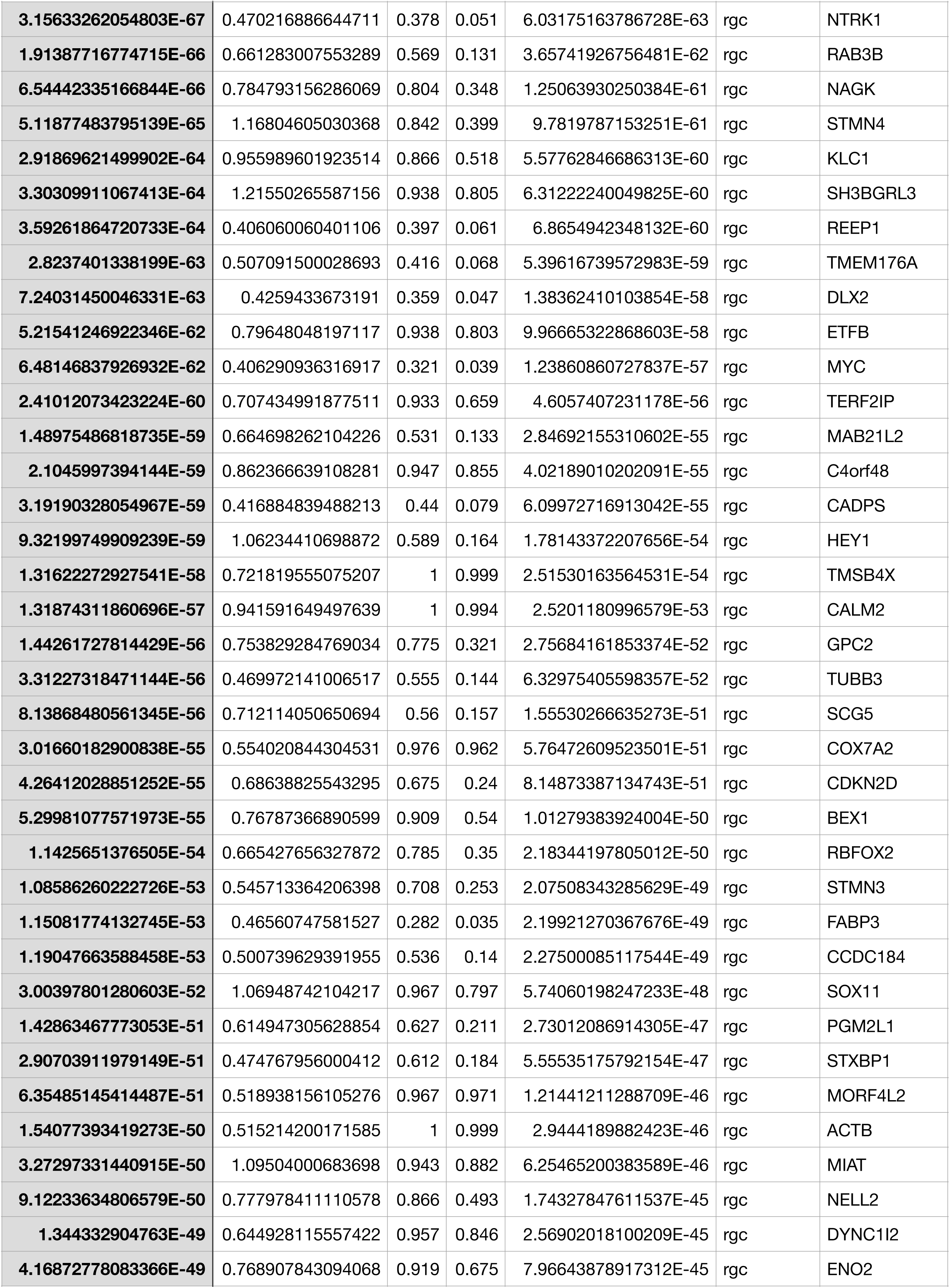
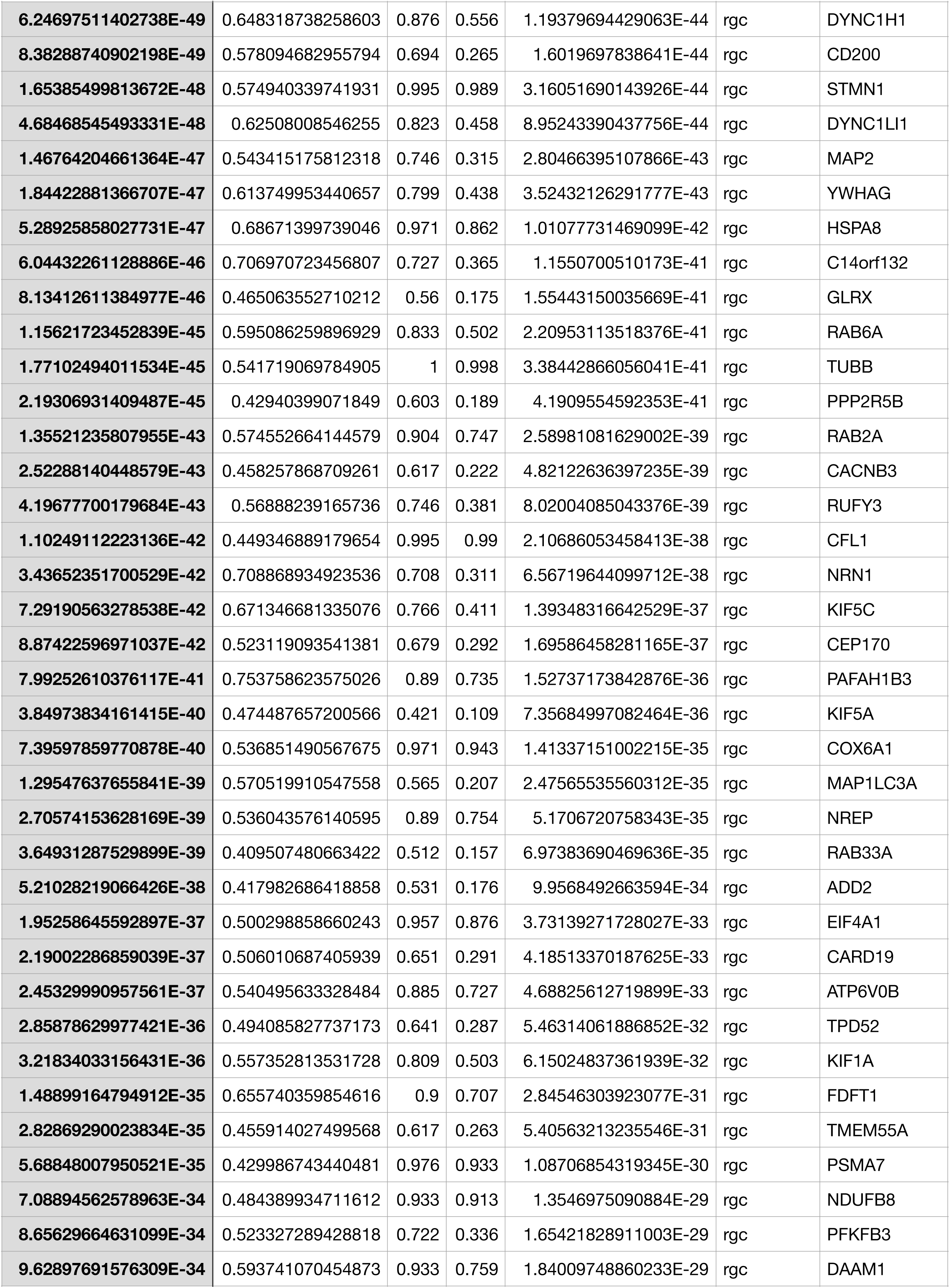
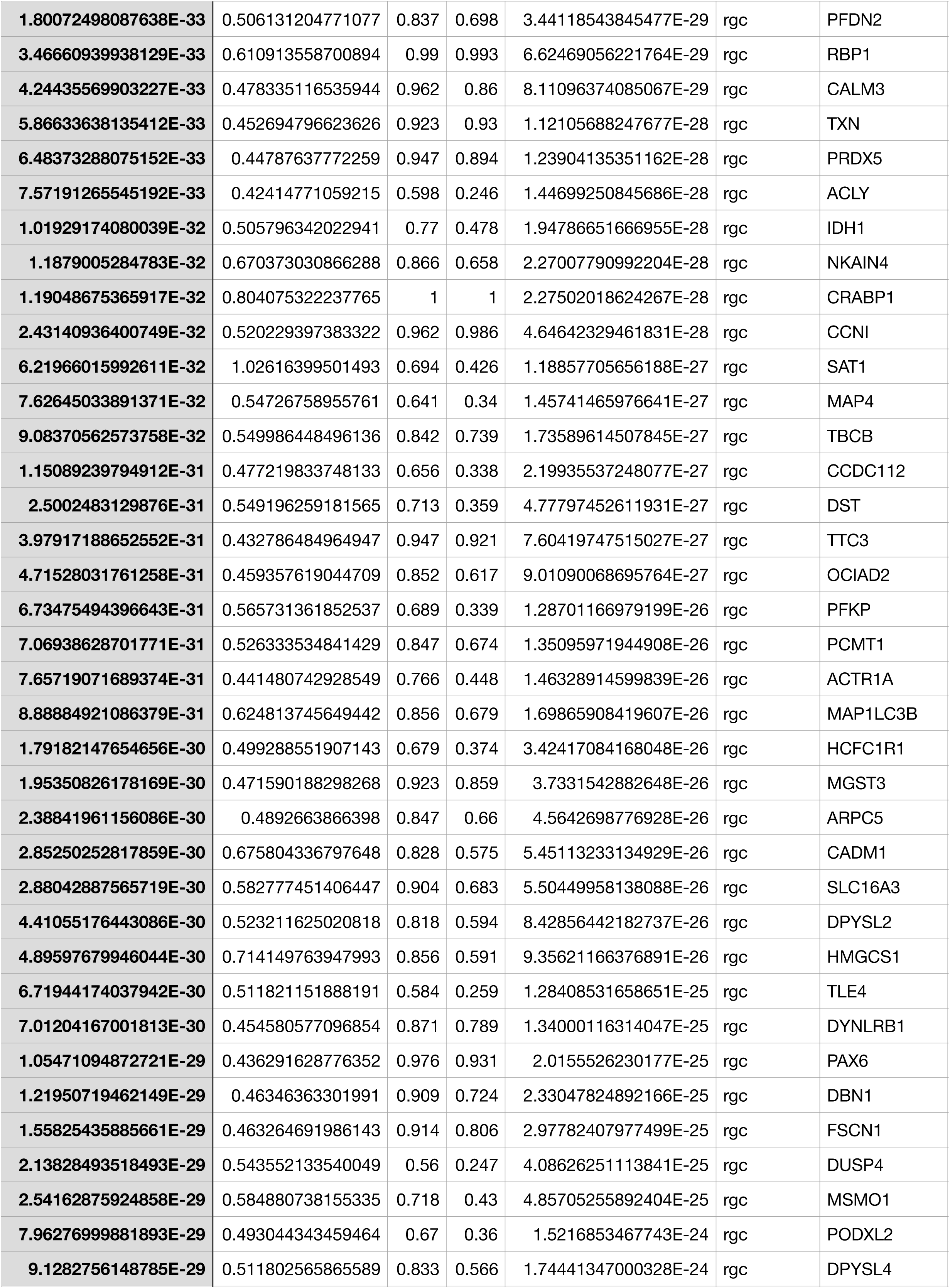
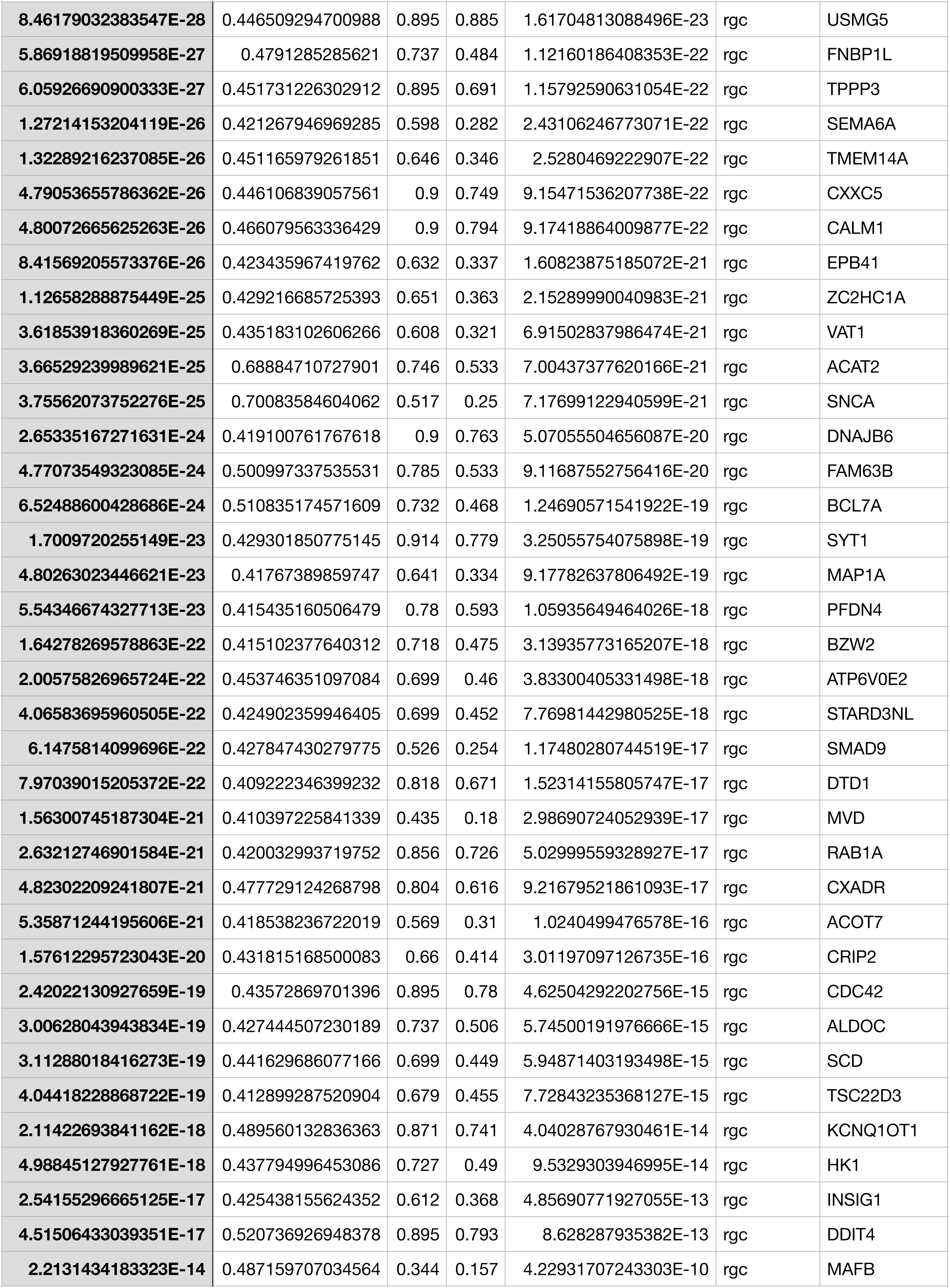
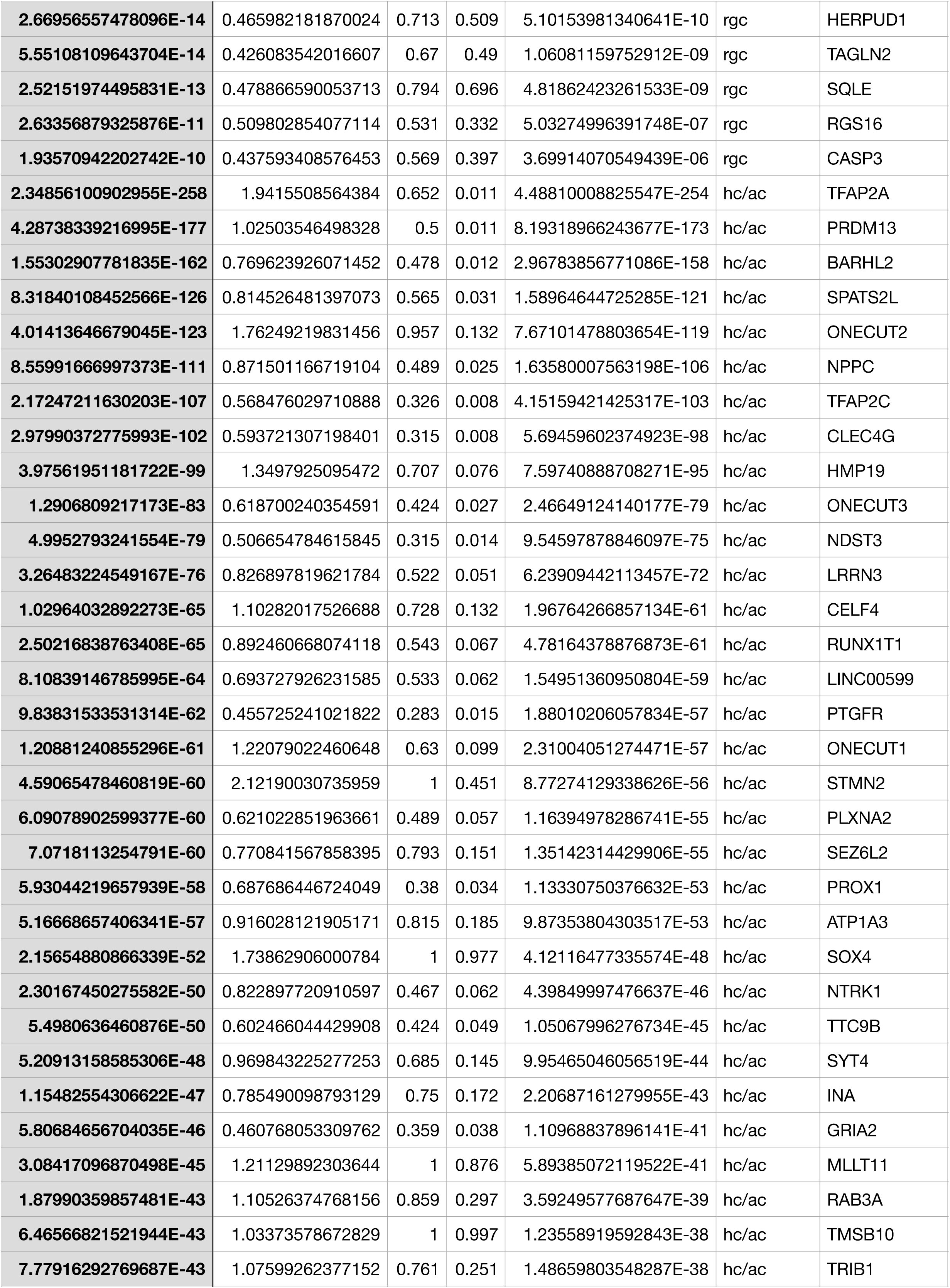
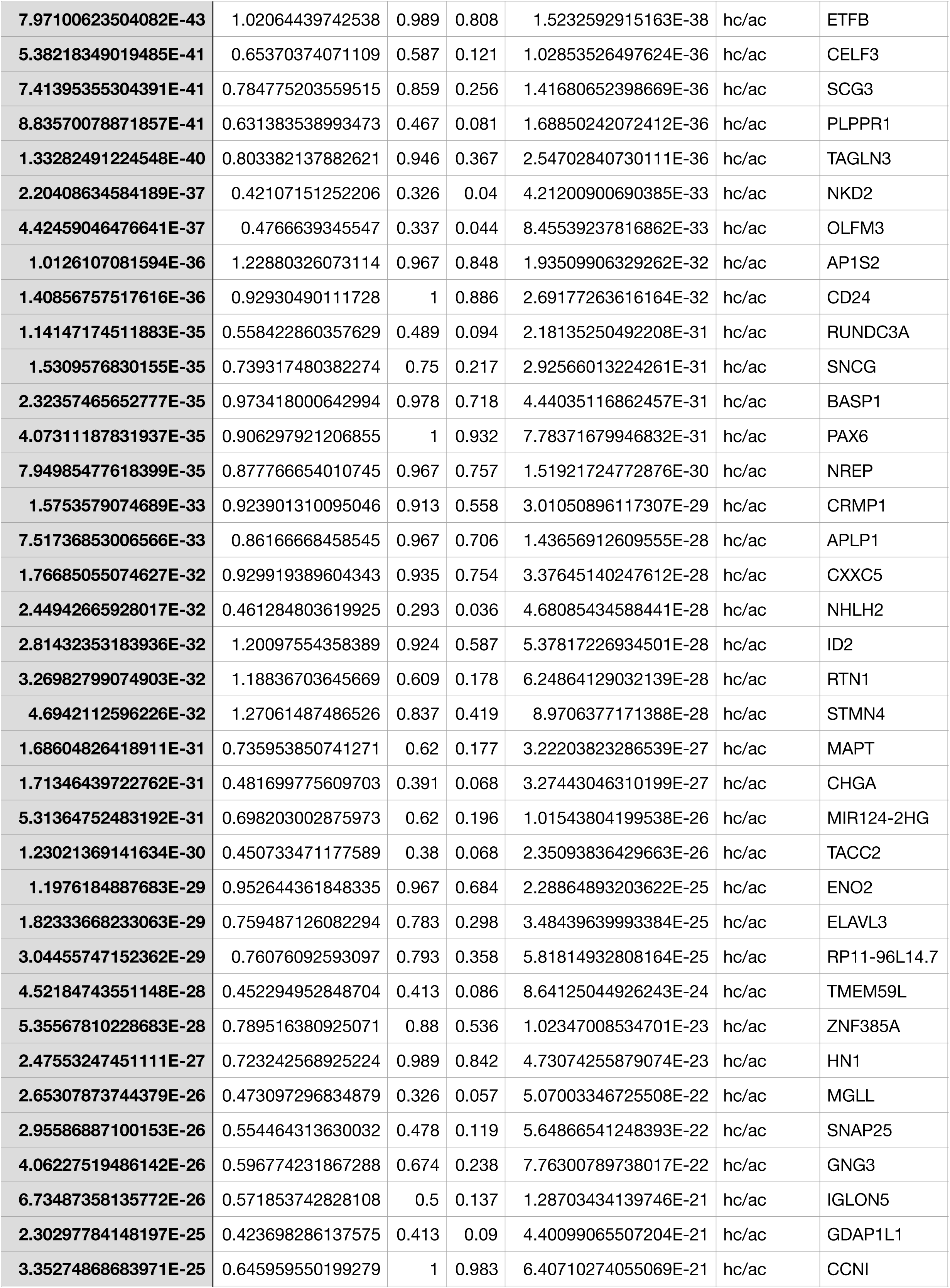
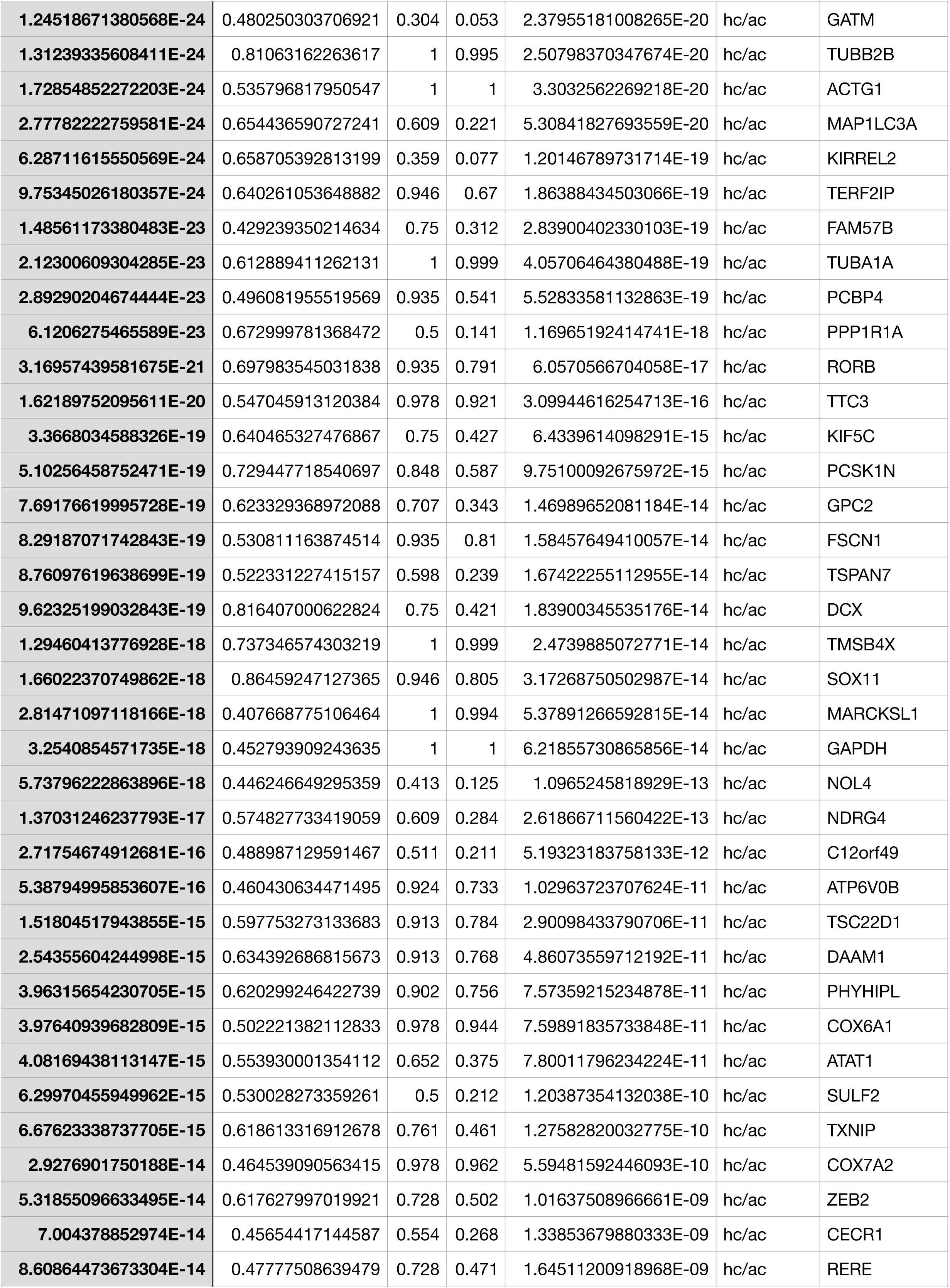
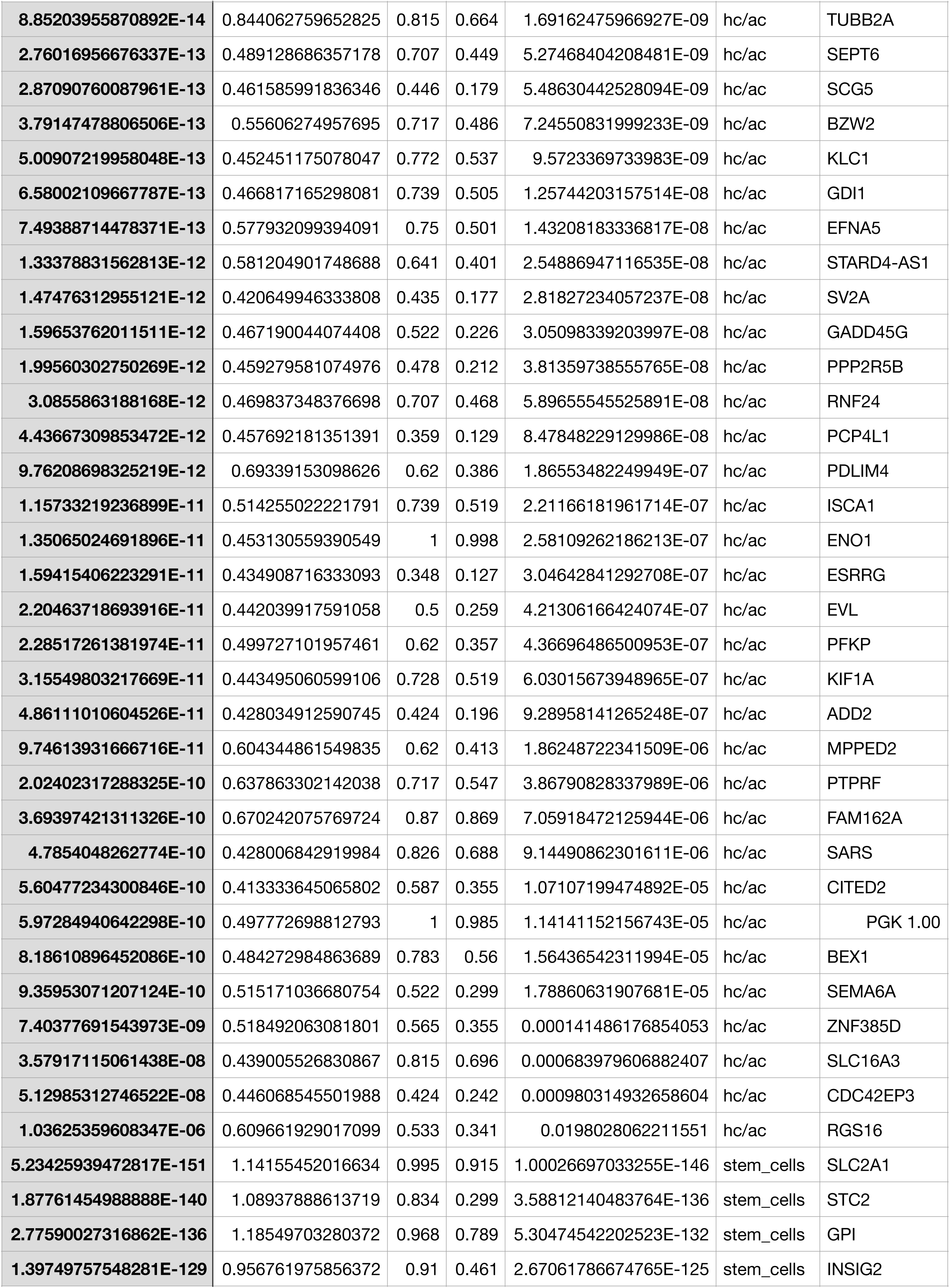
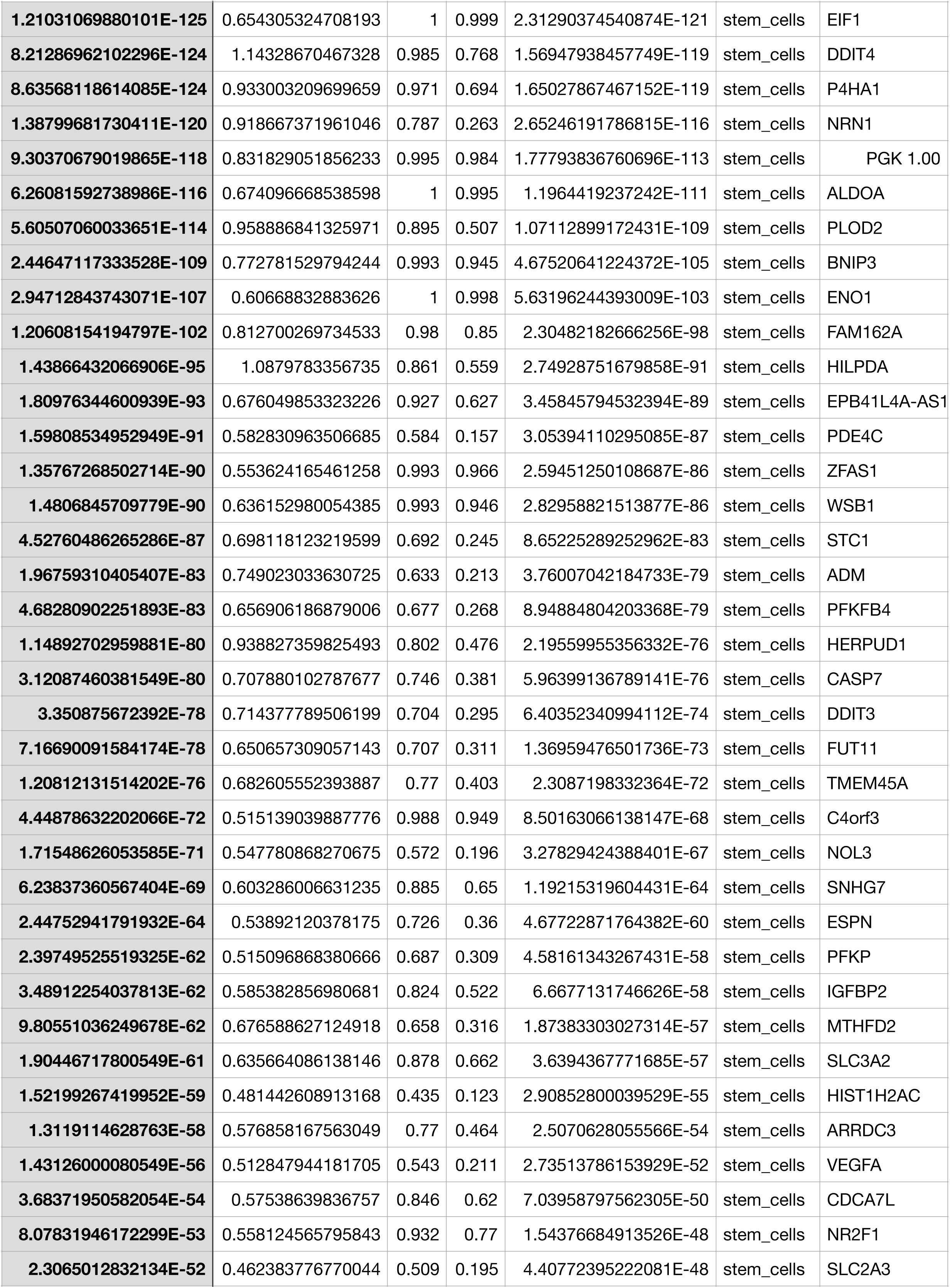
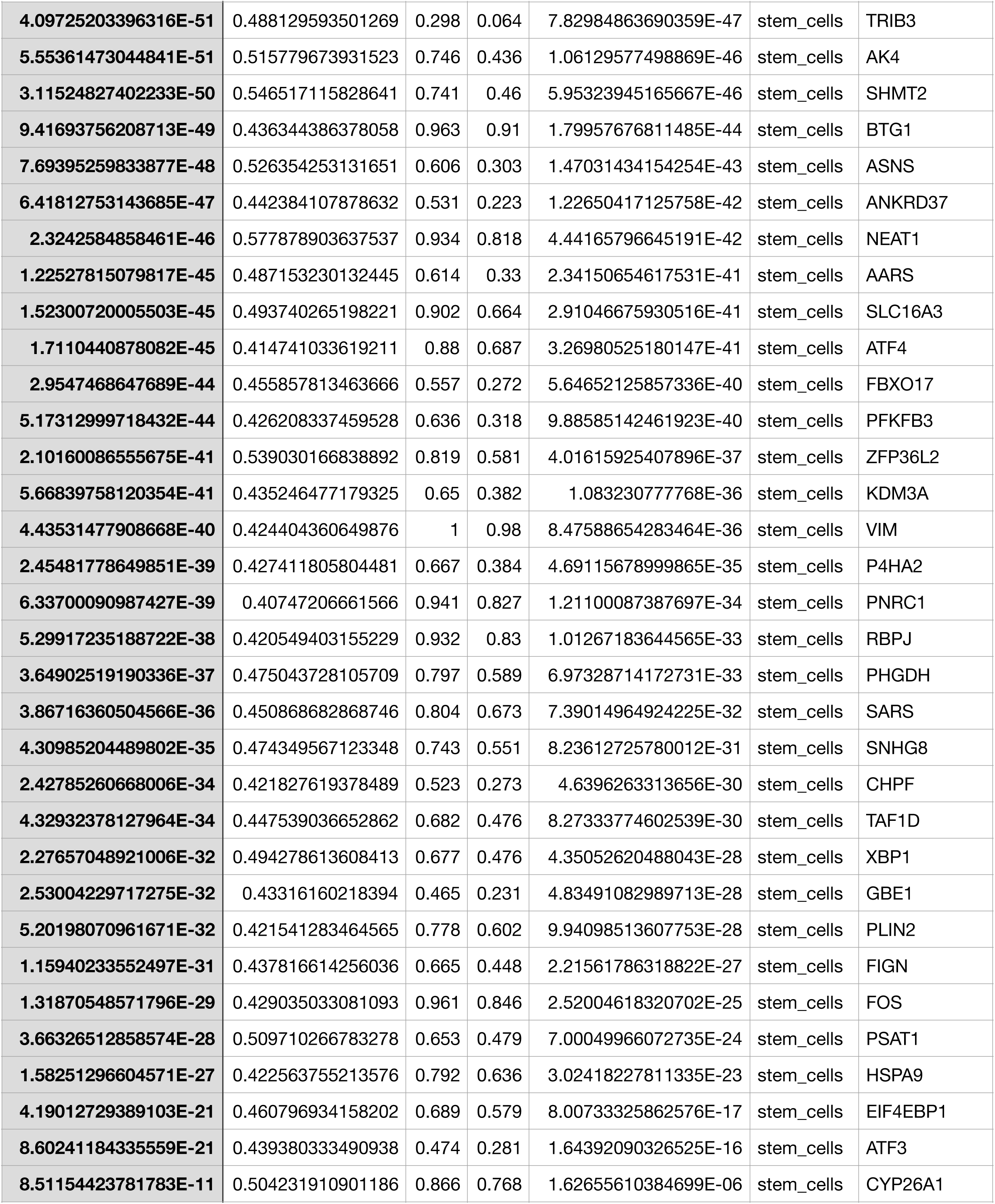
enriched_genes GFP

**SupTable 2.**
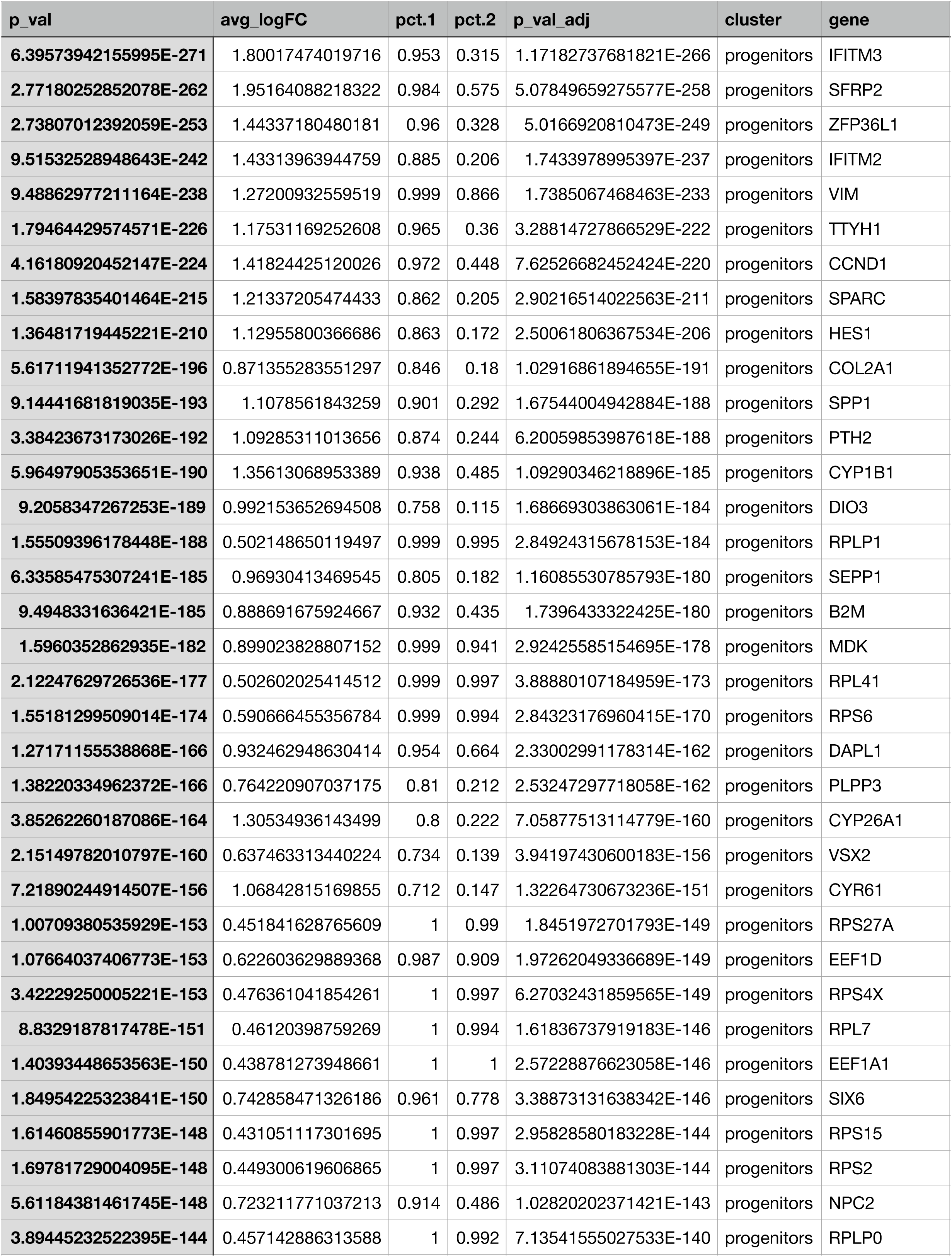
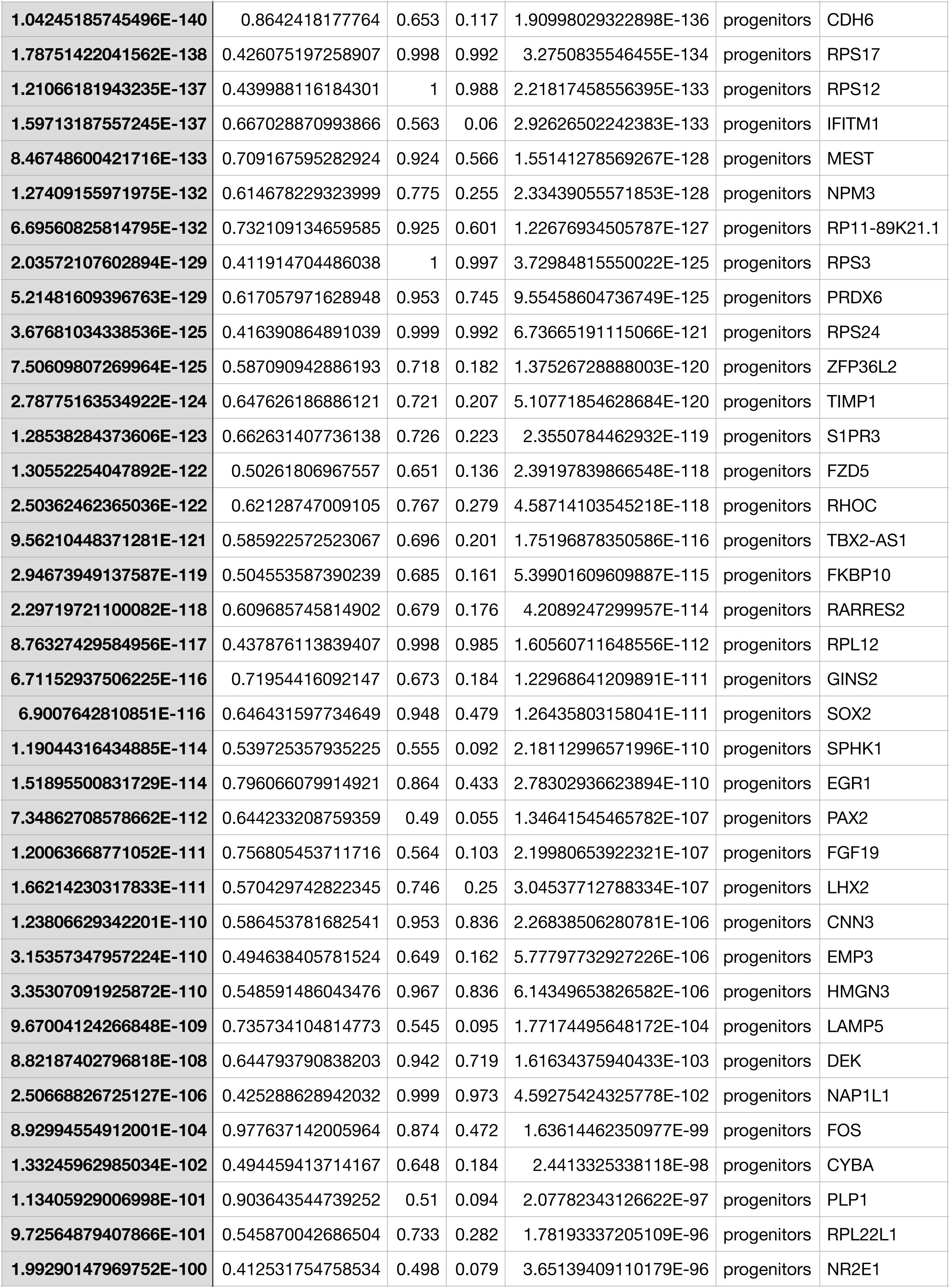
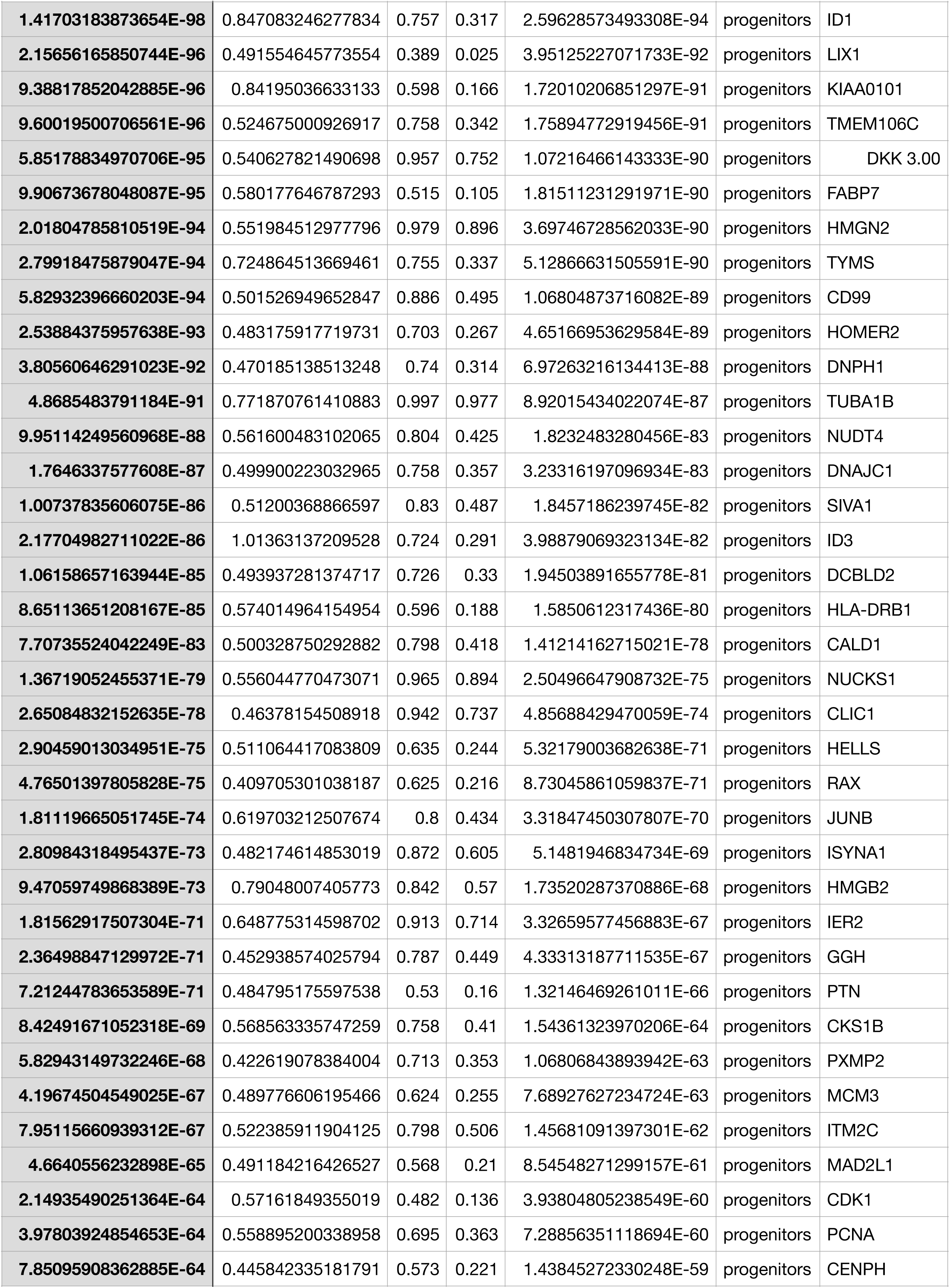
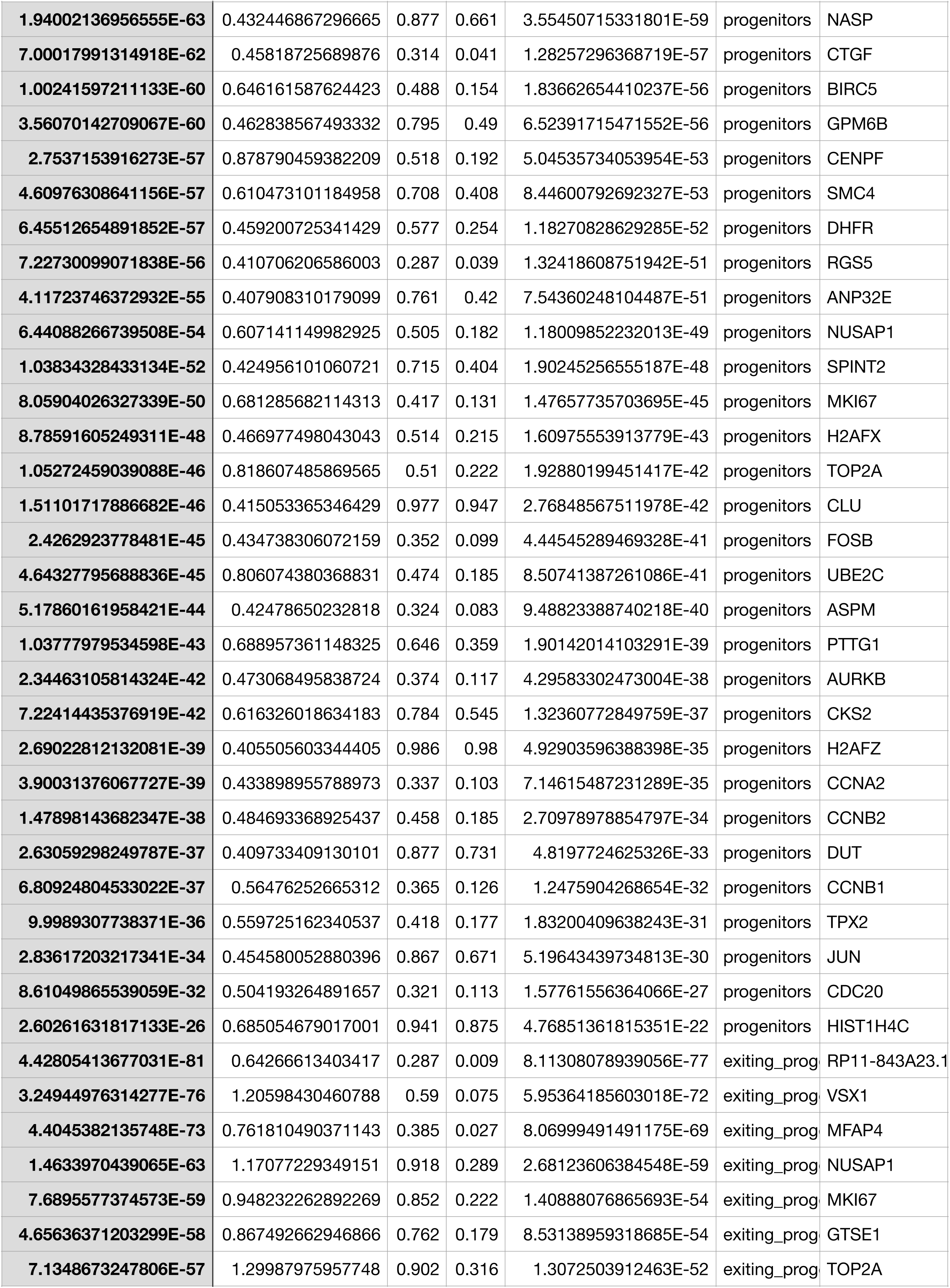
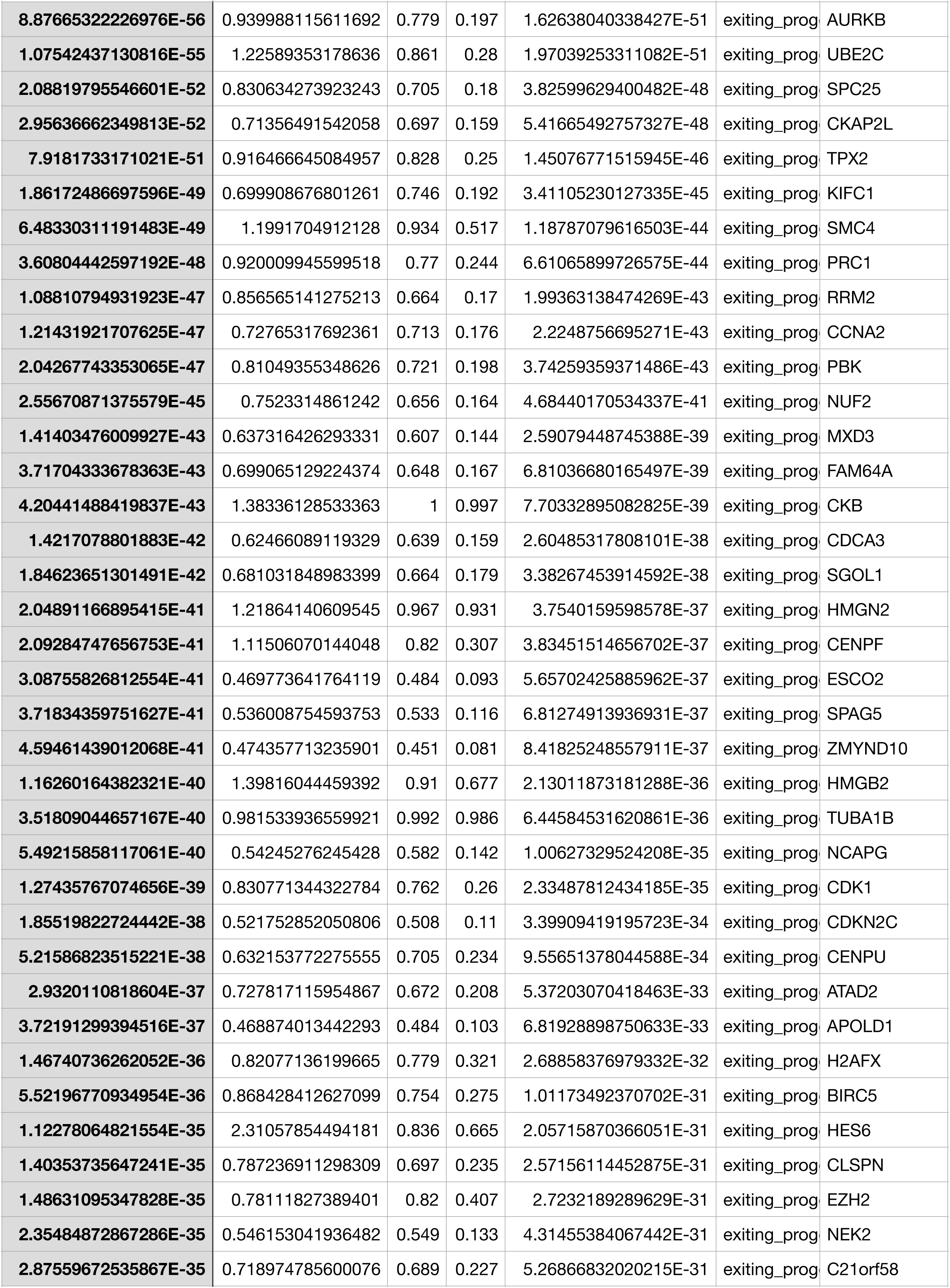
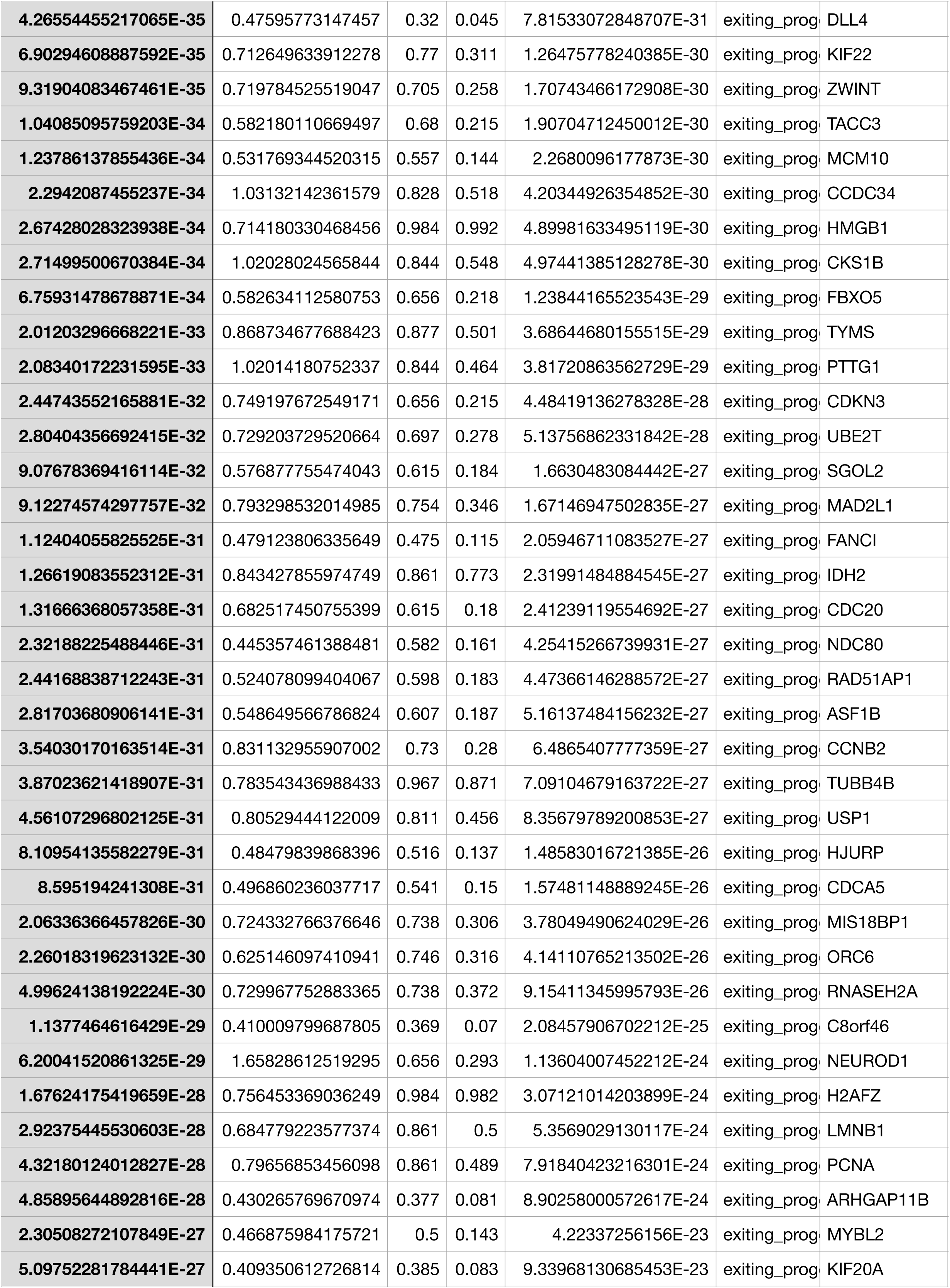
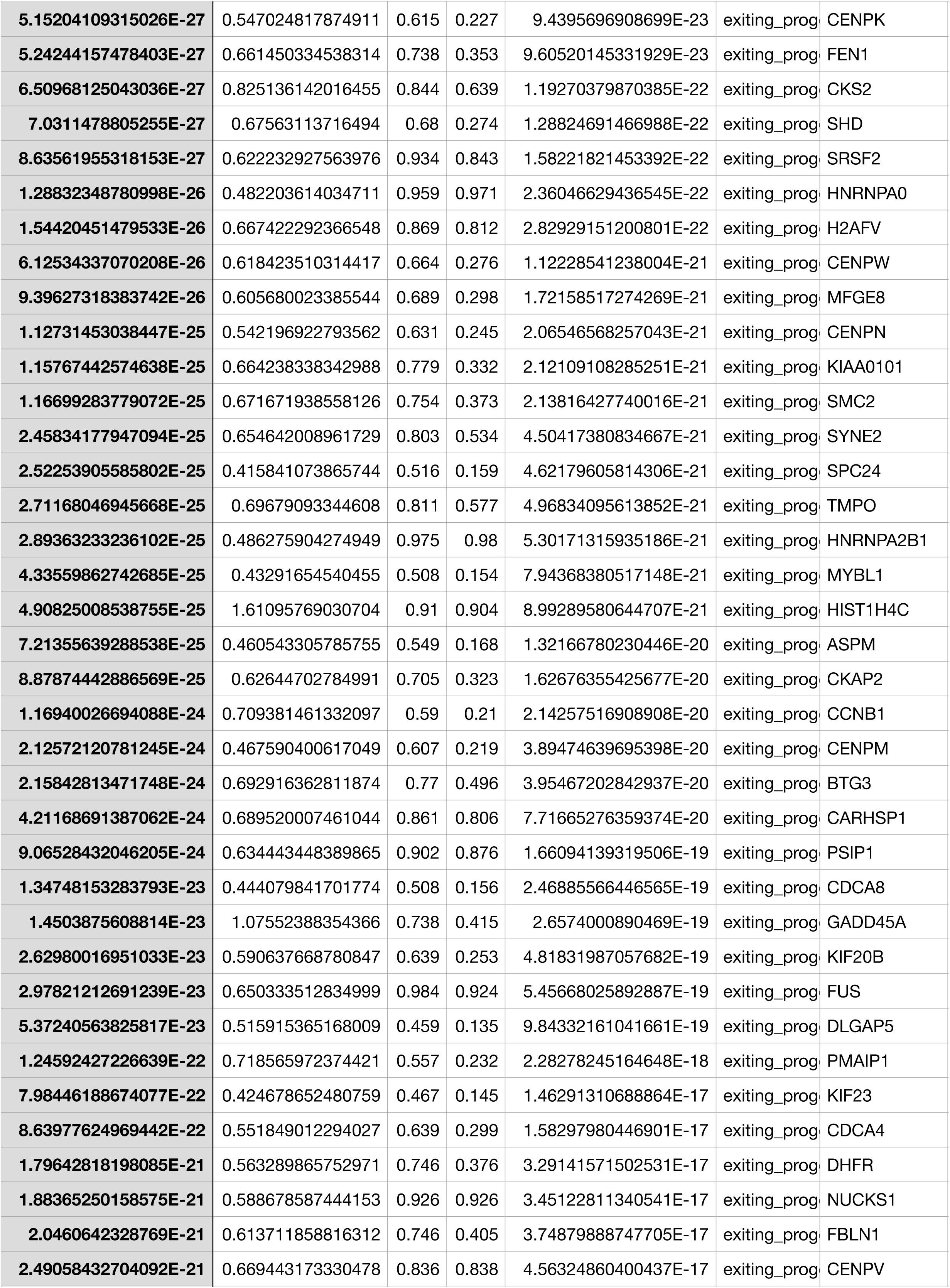
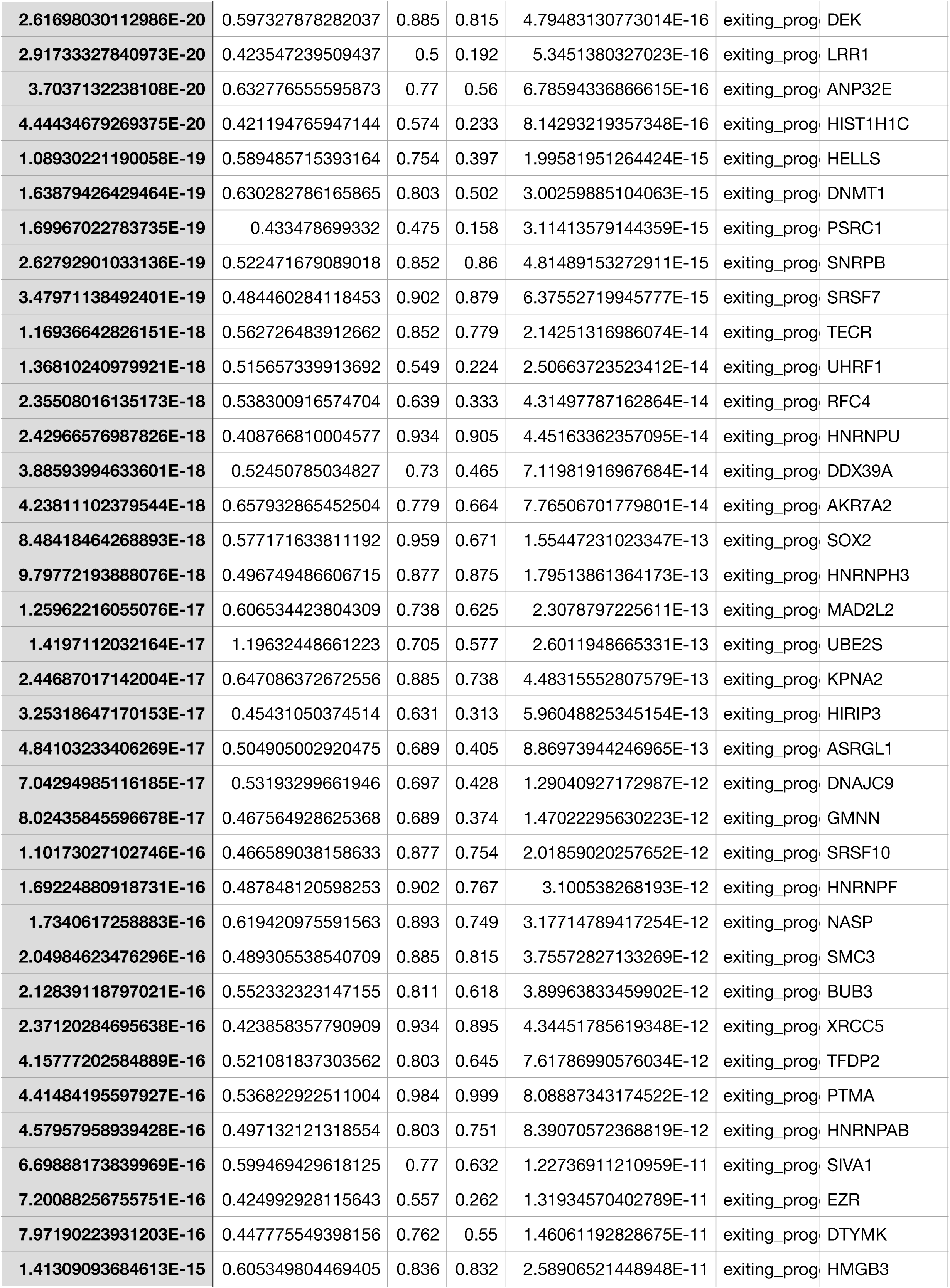
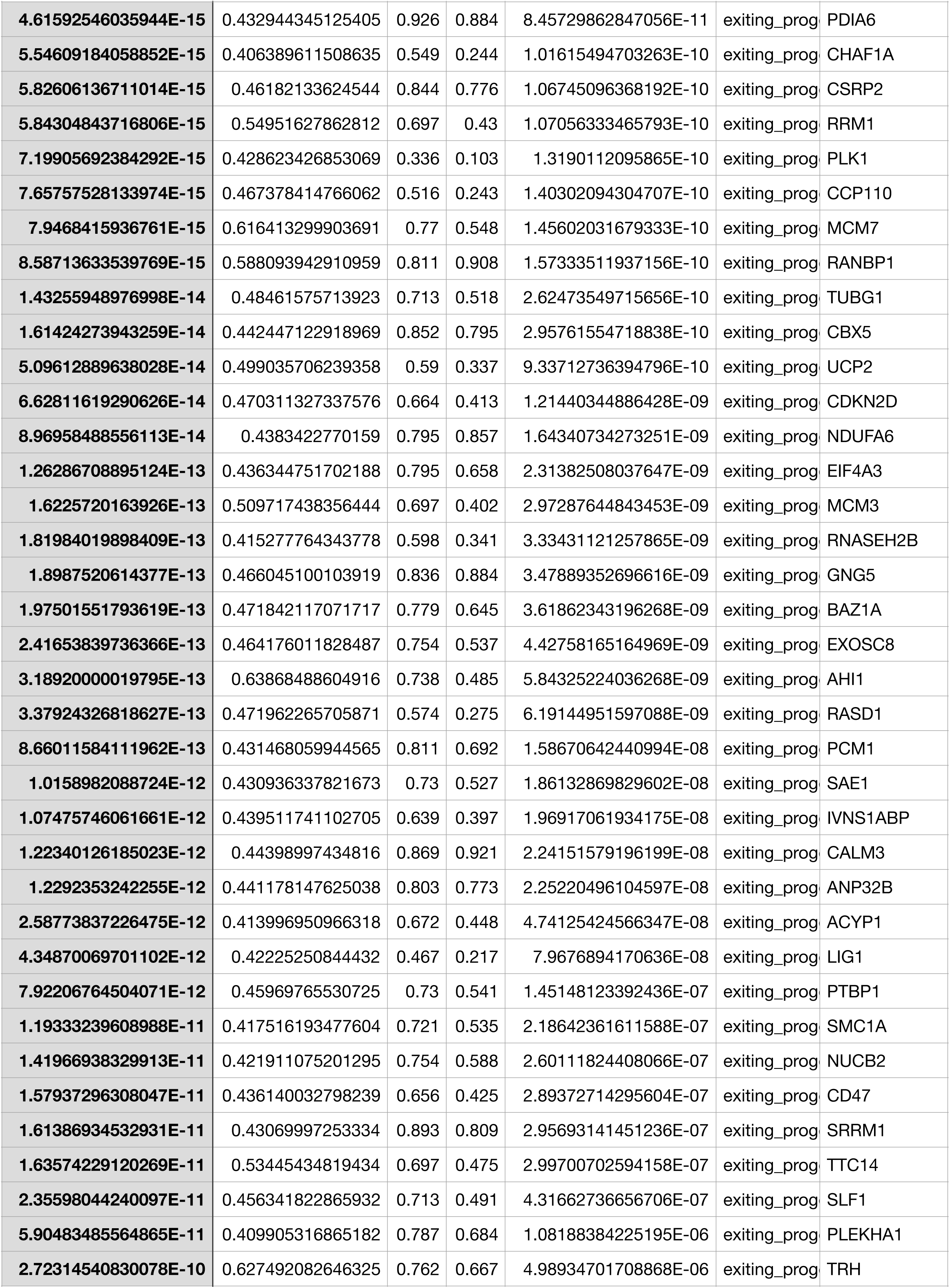
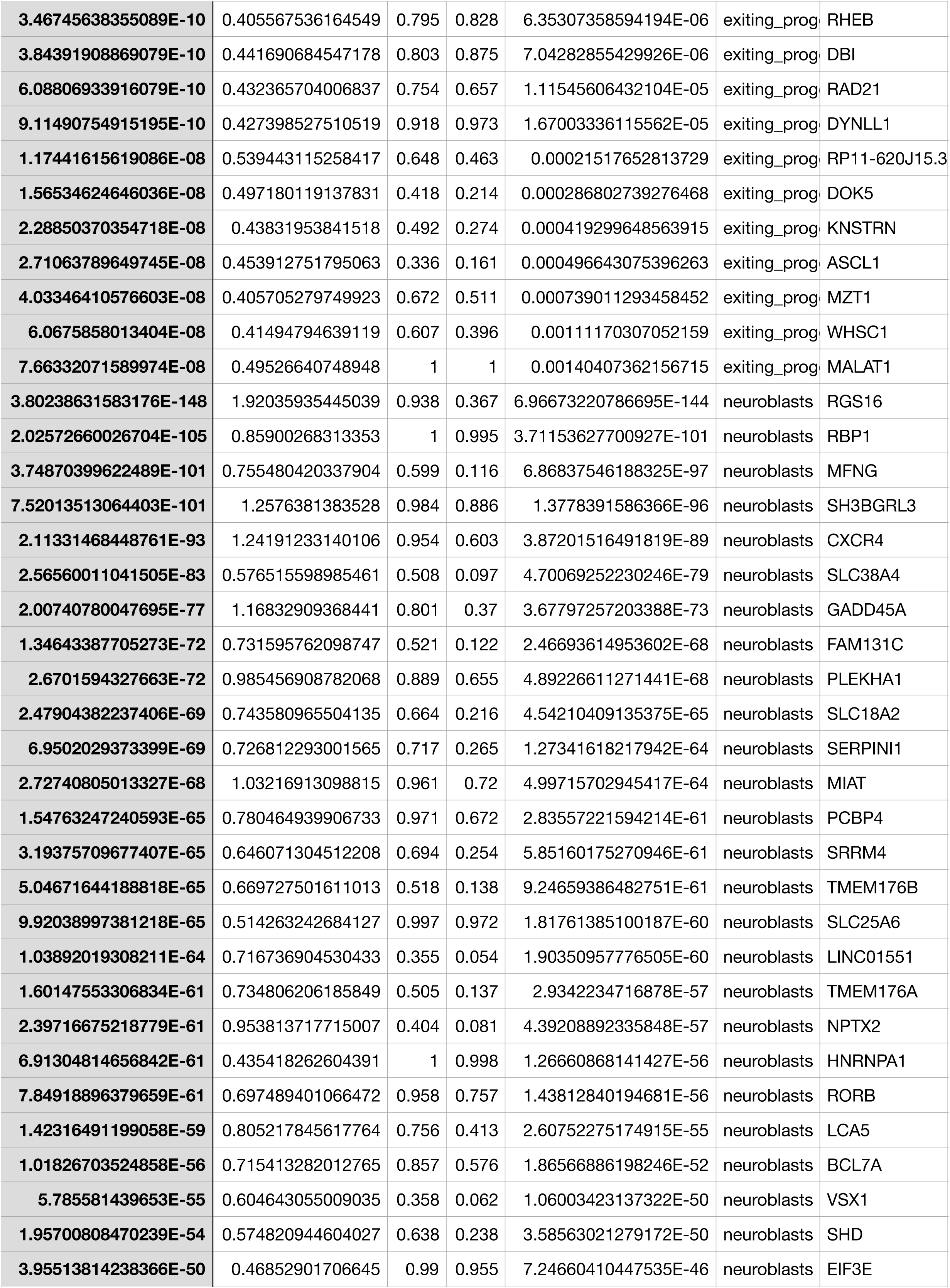
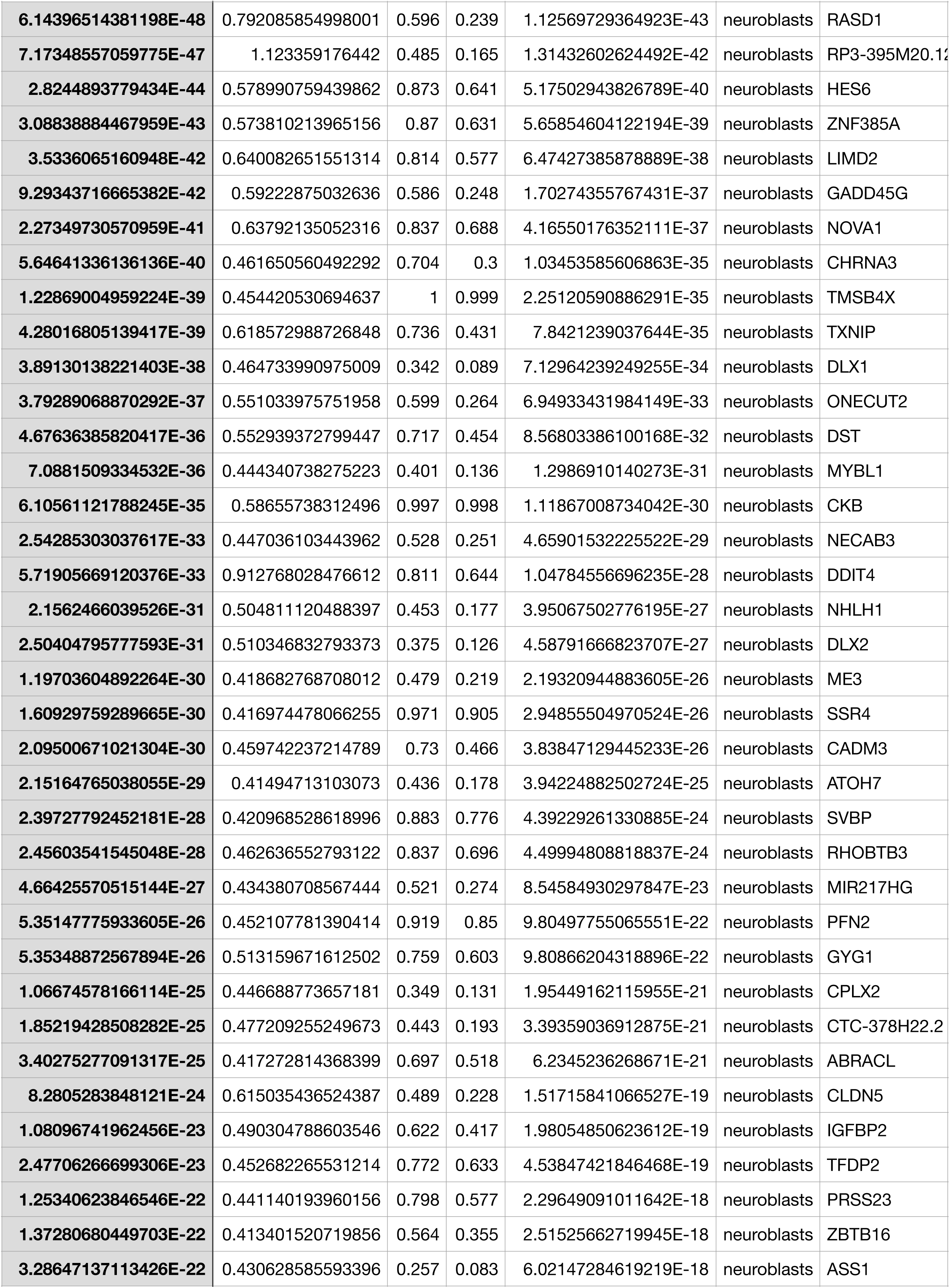
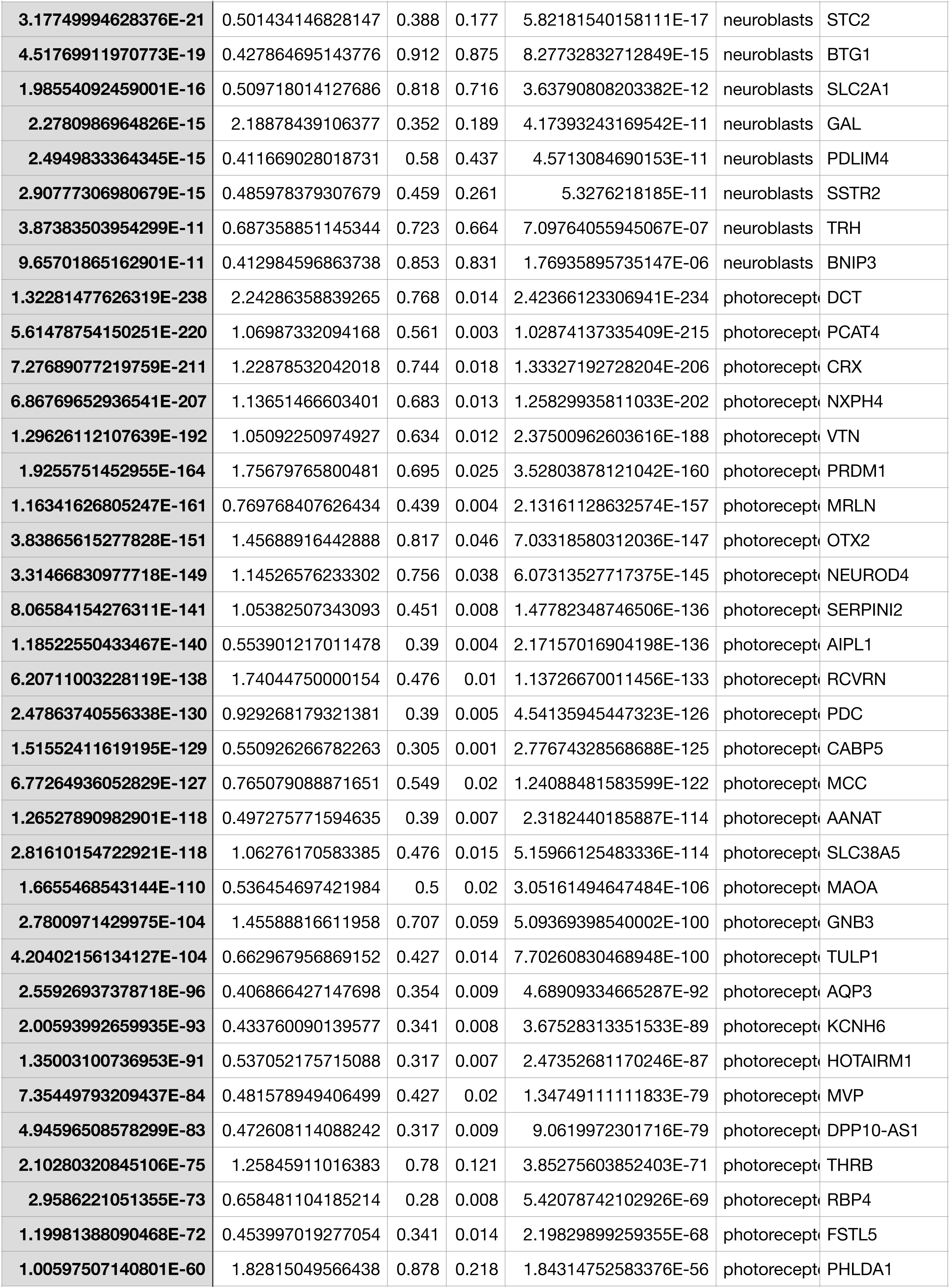
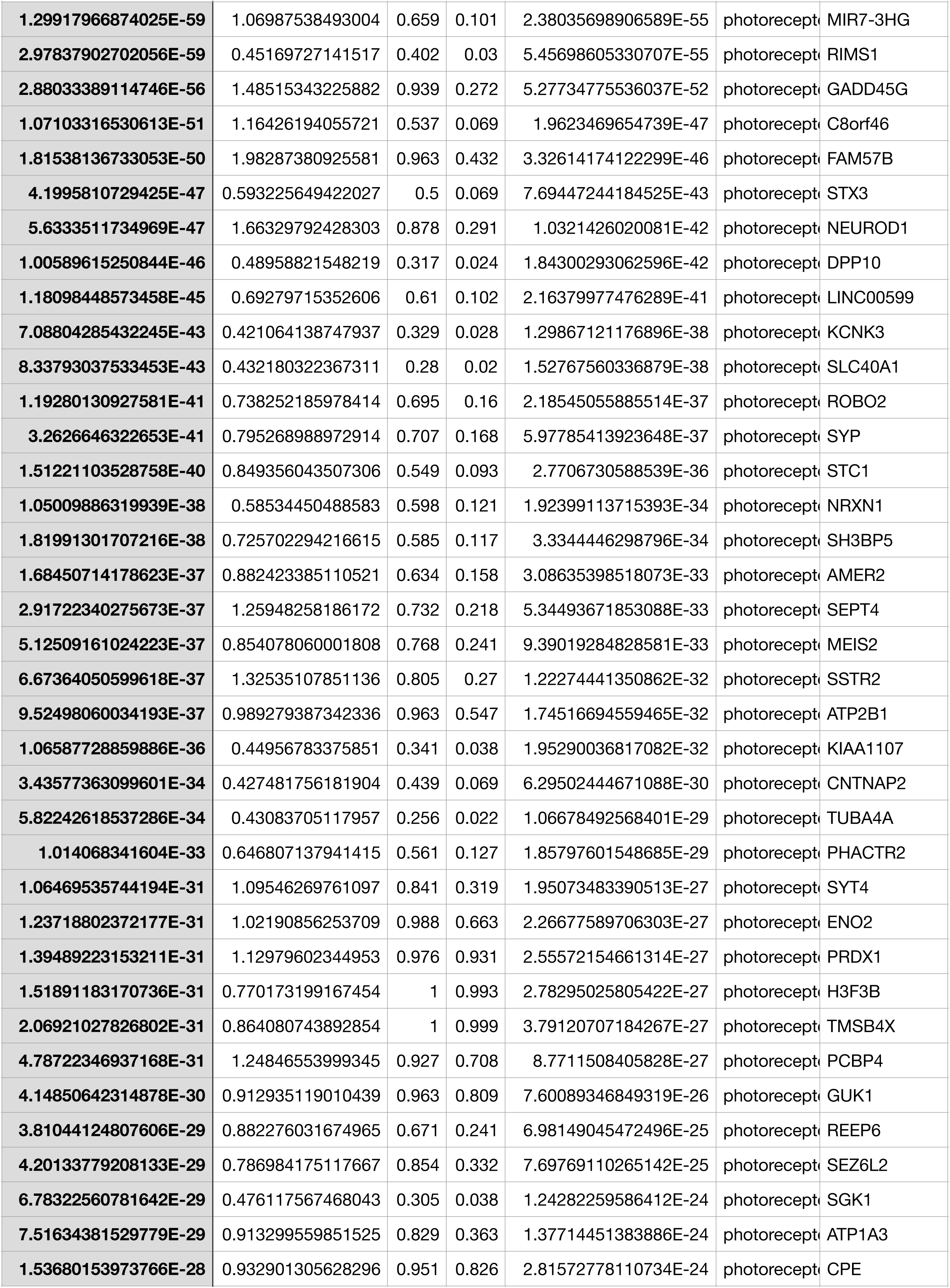
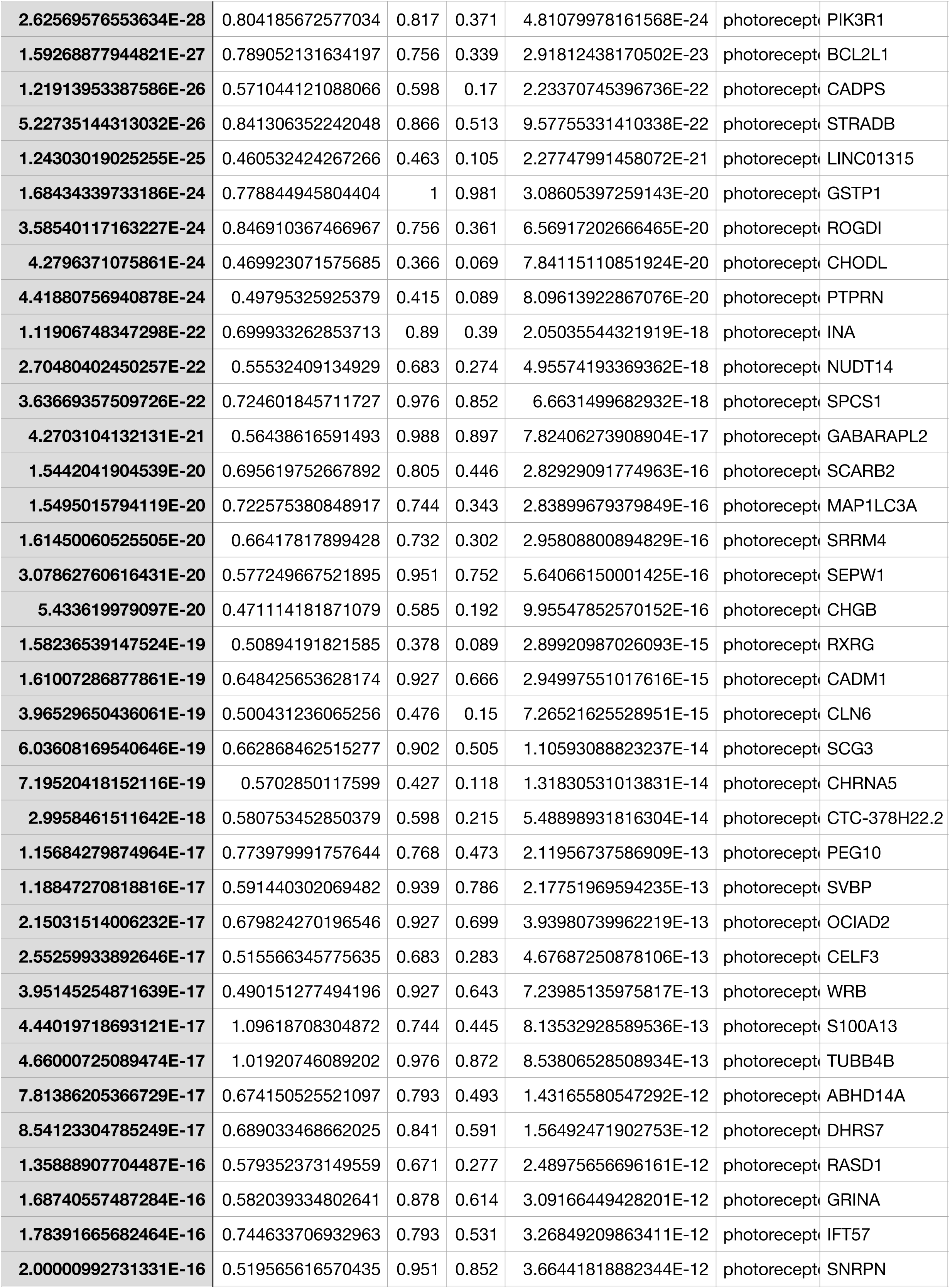
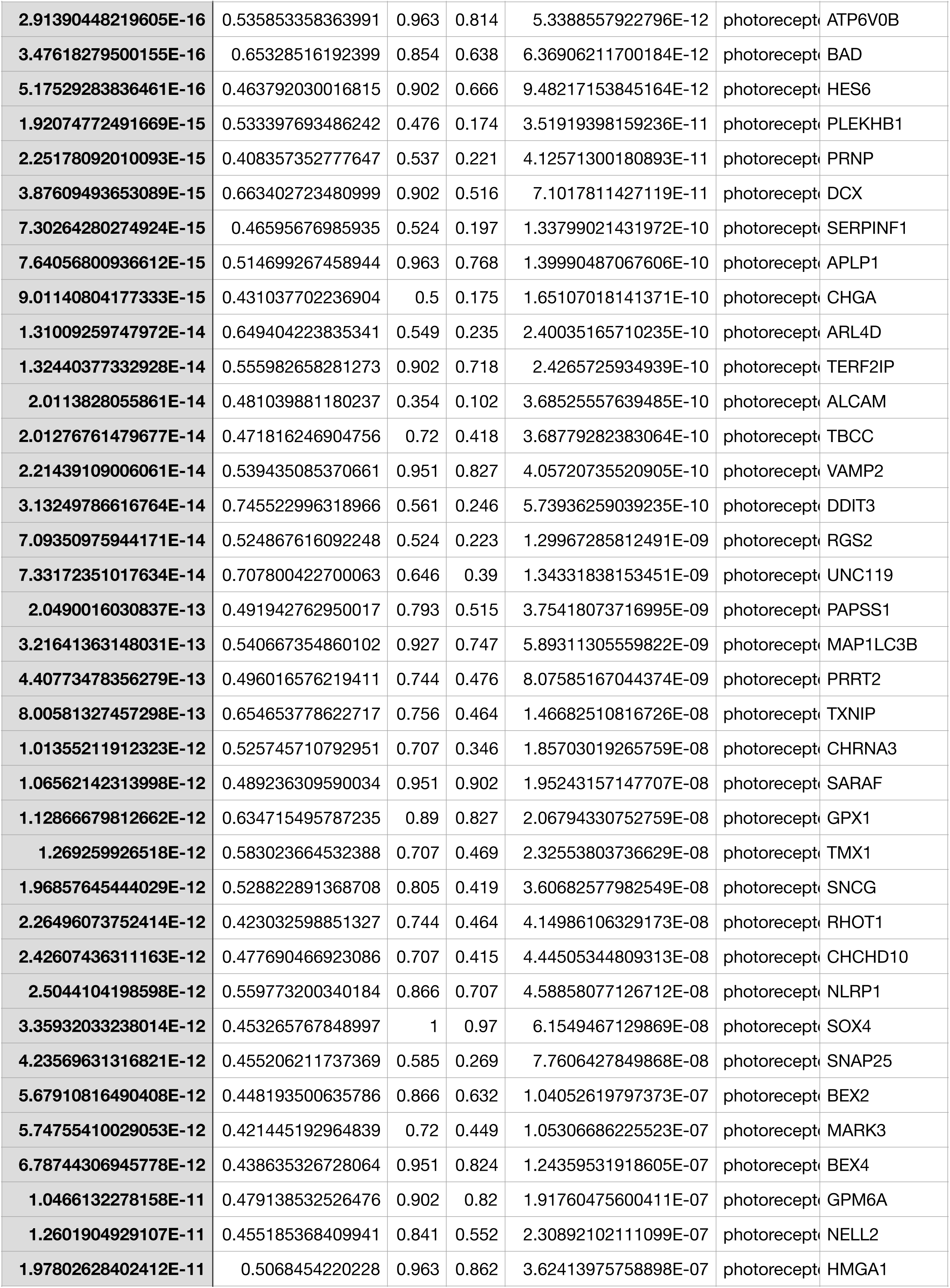
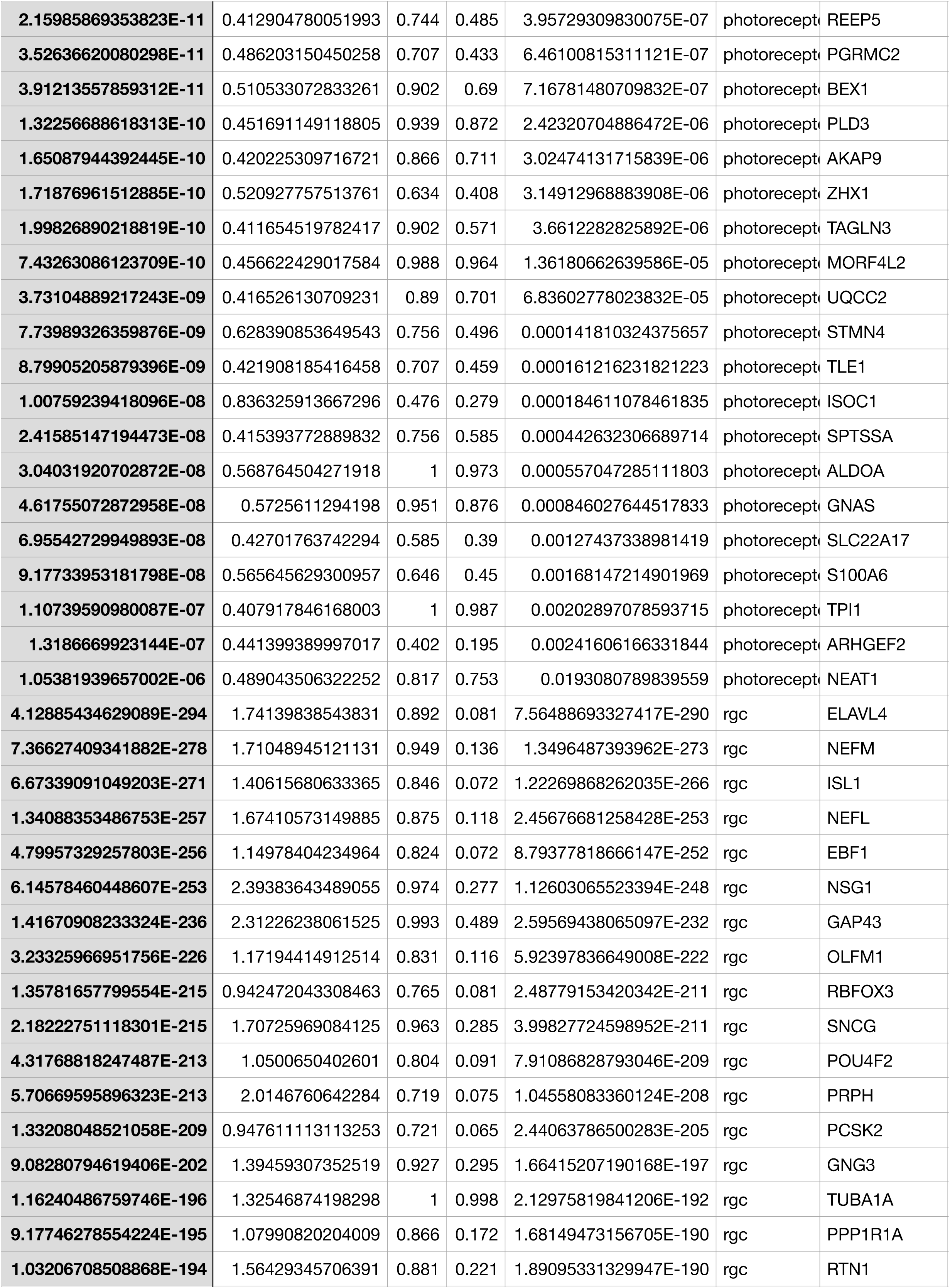
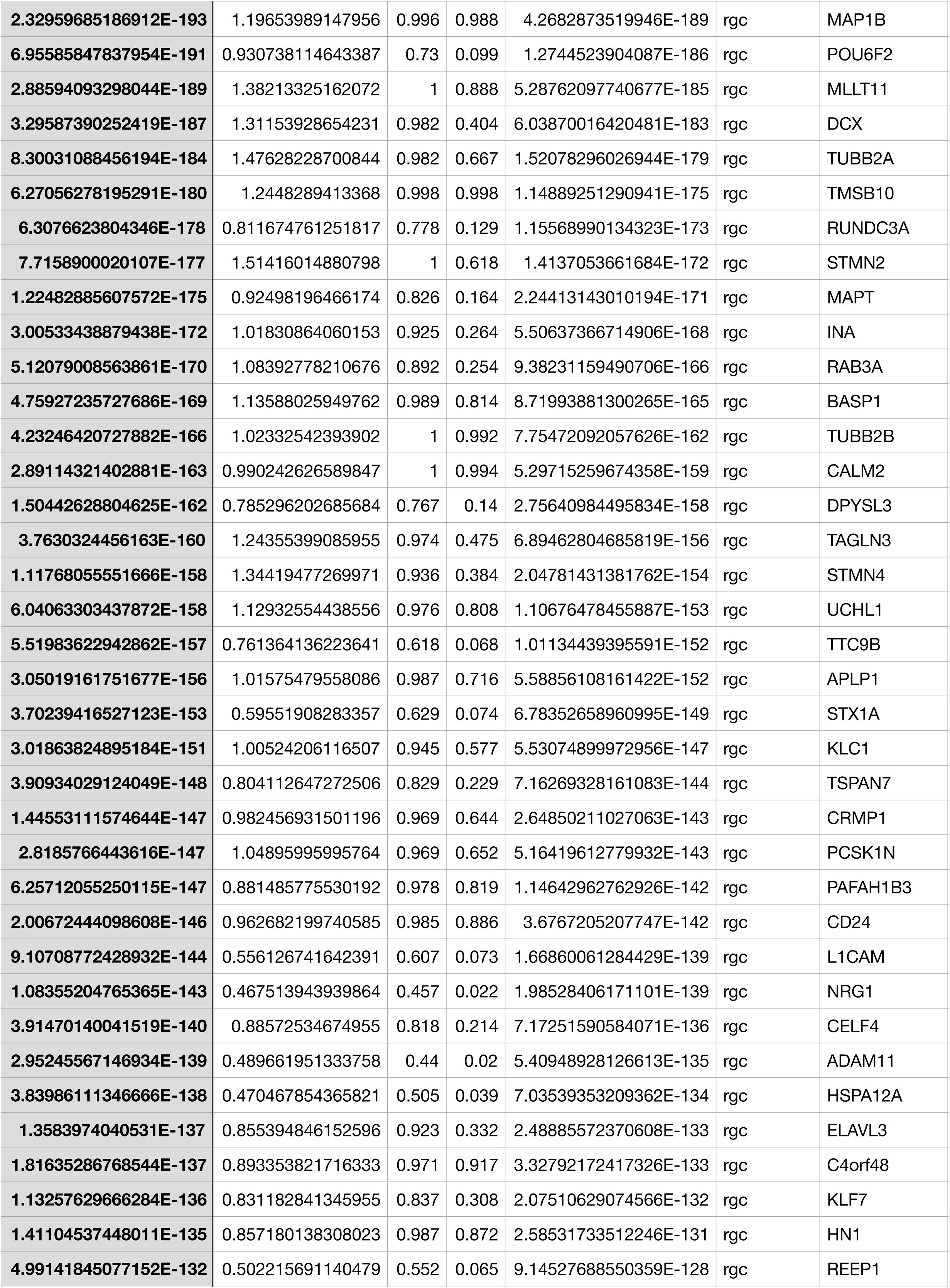
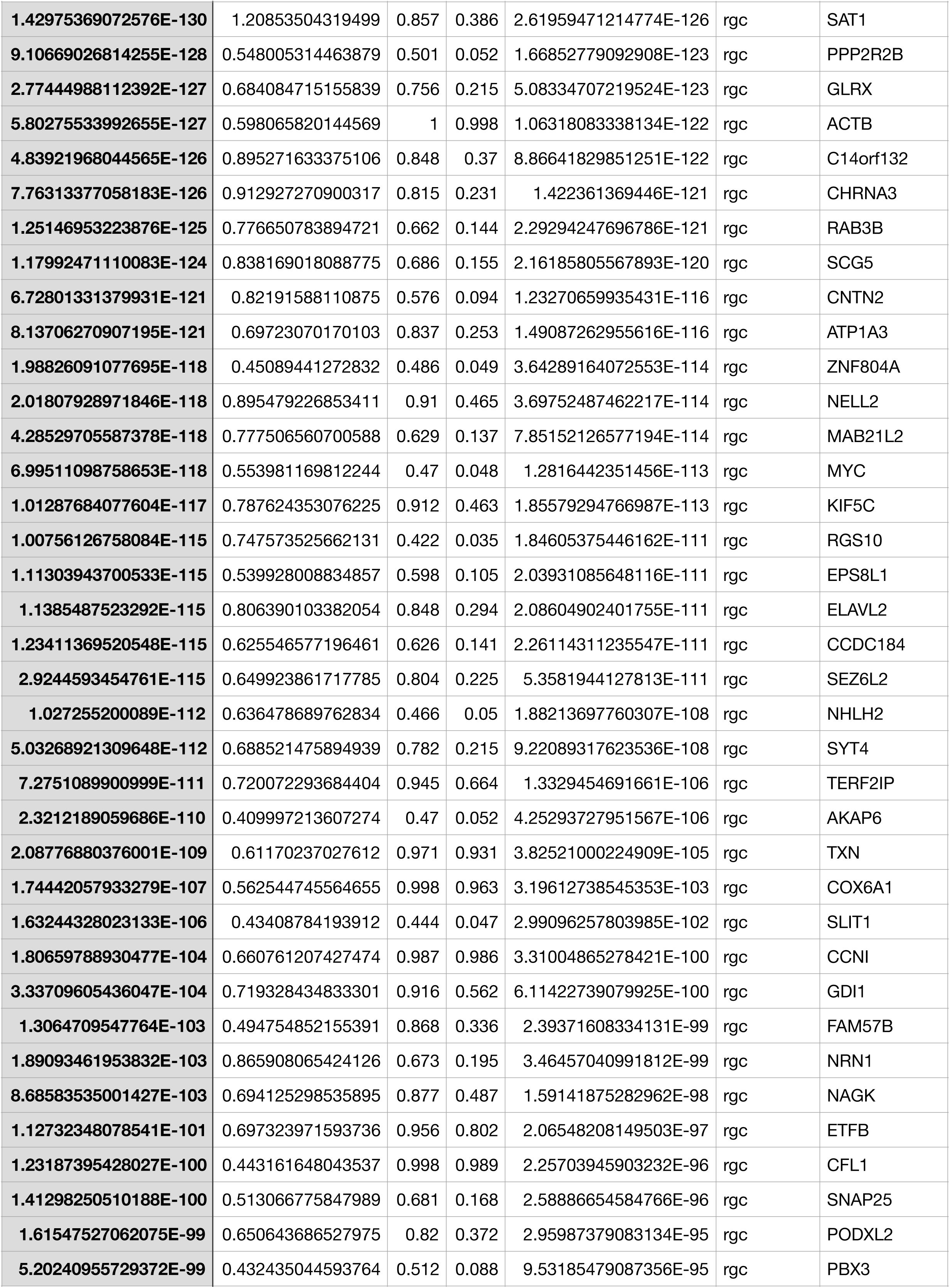
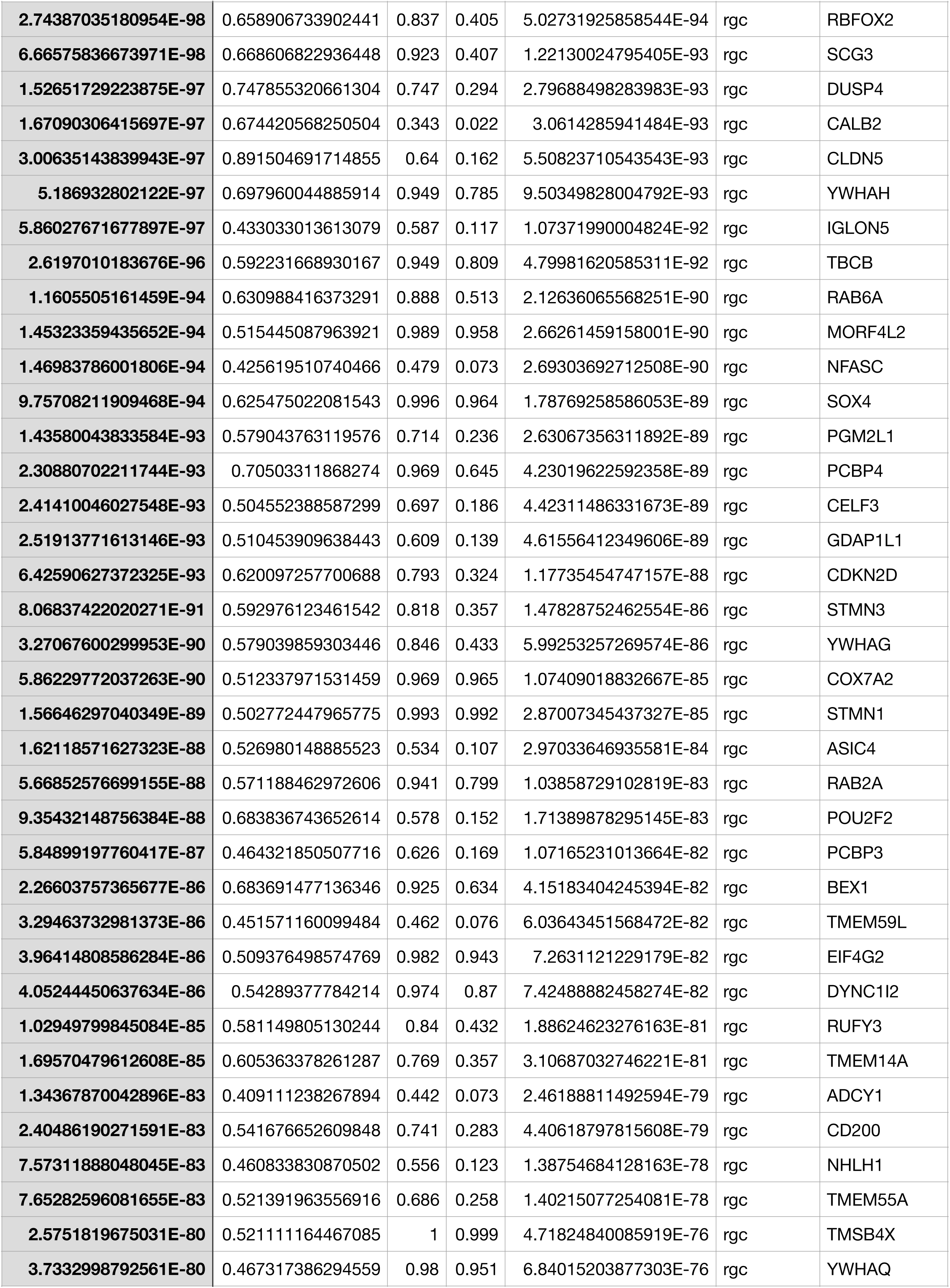
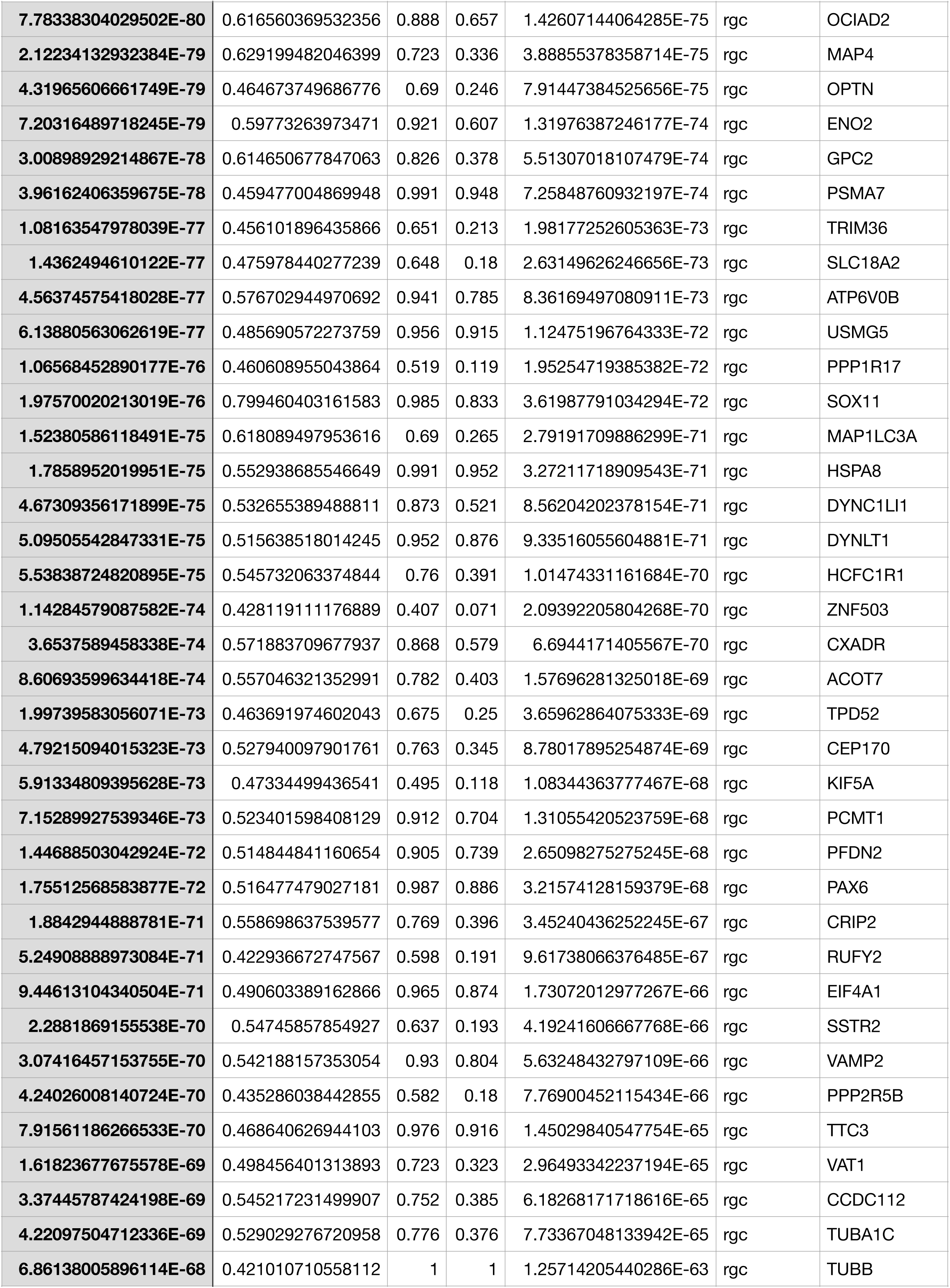
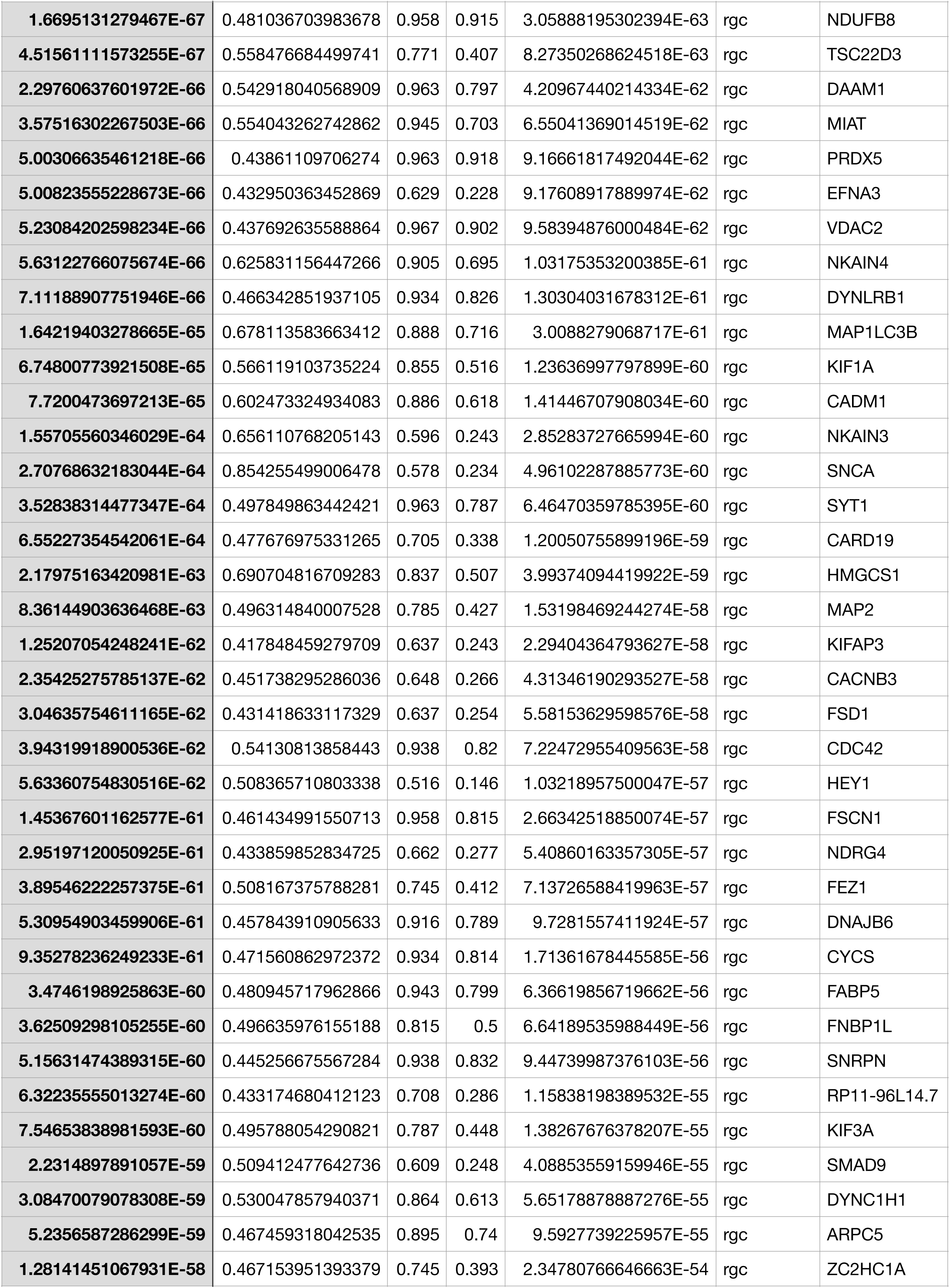
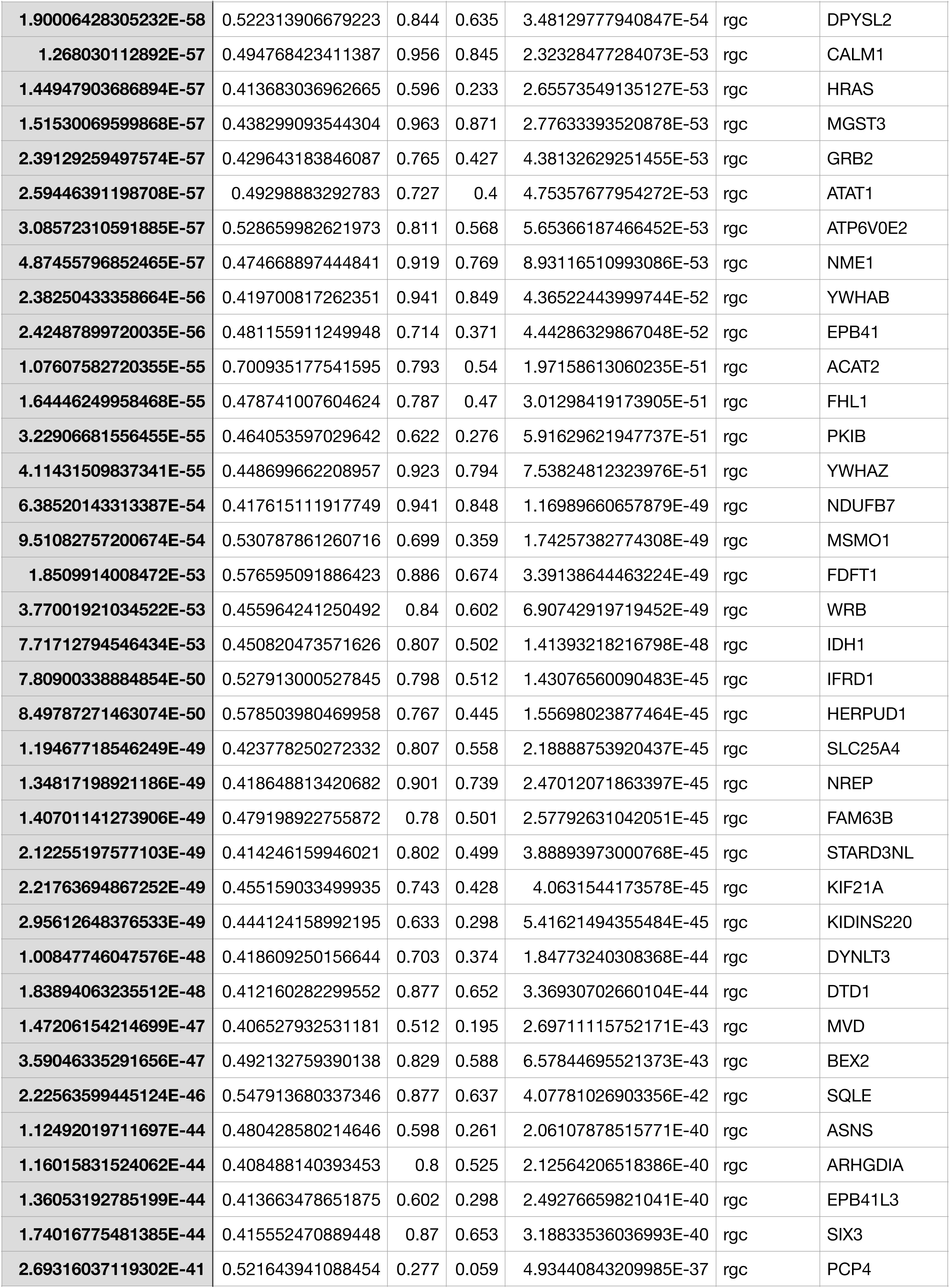
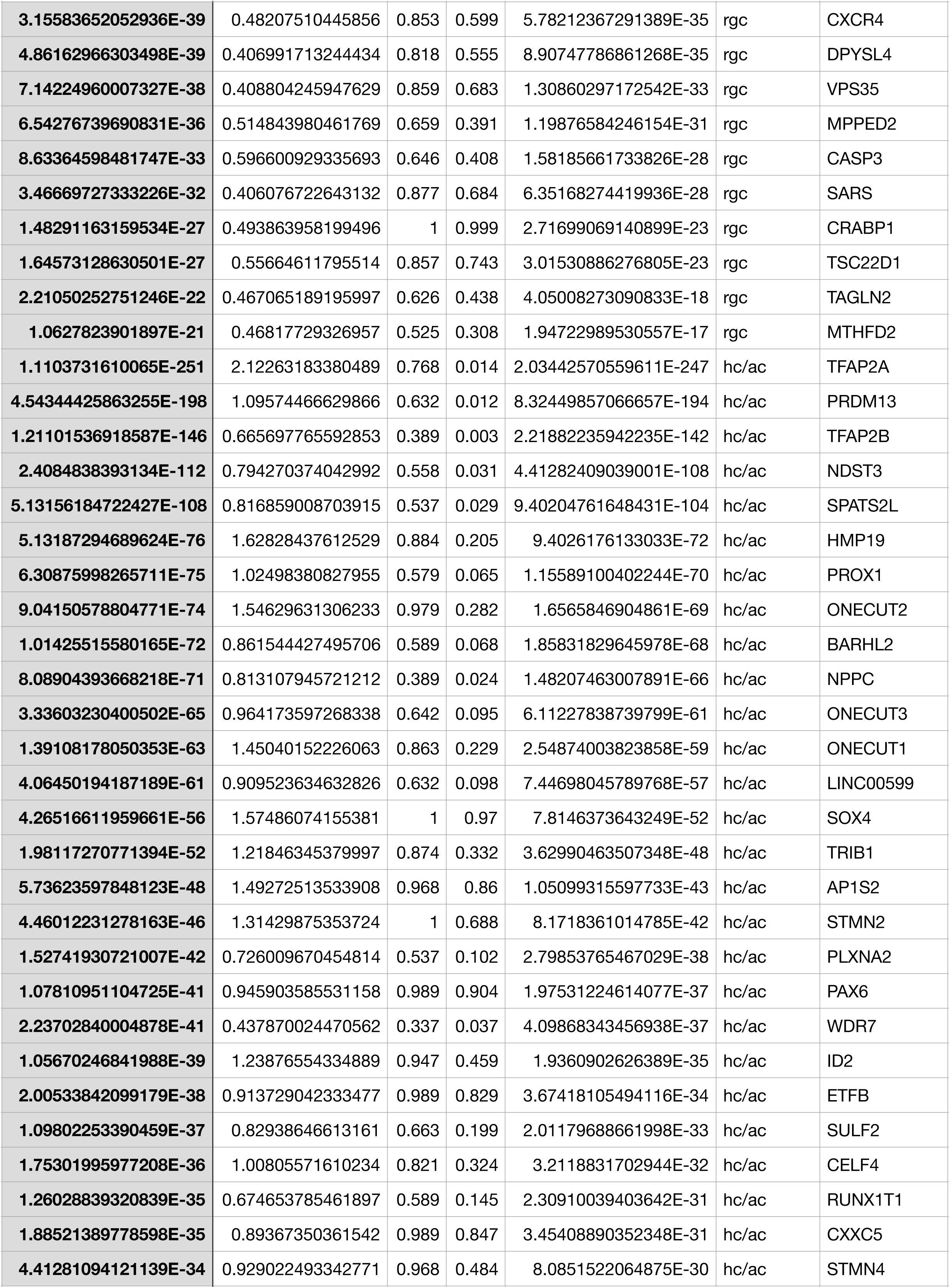
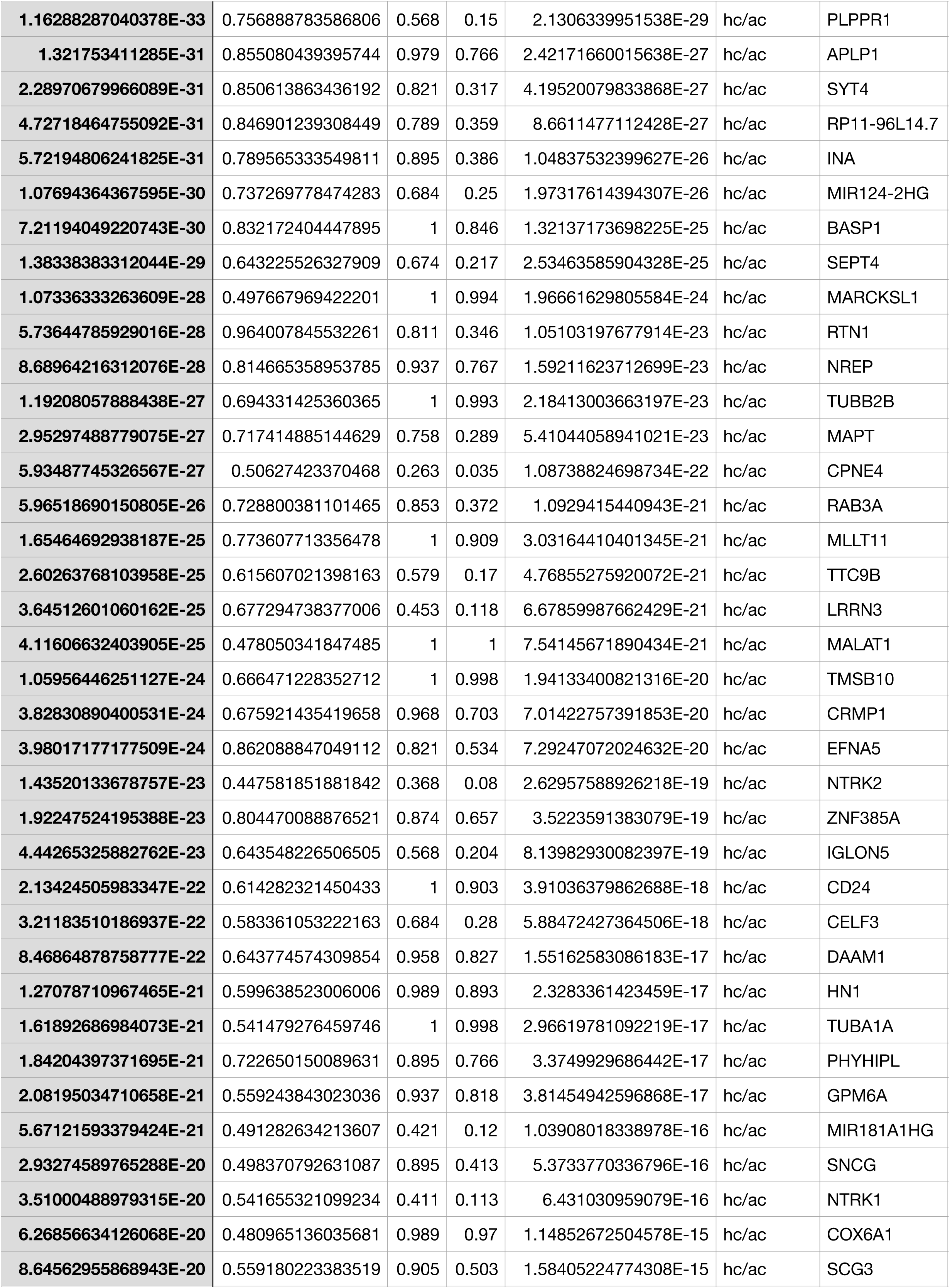
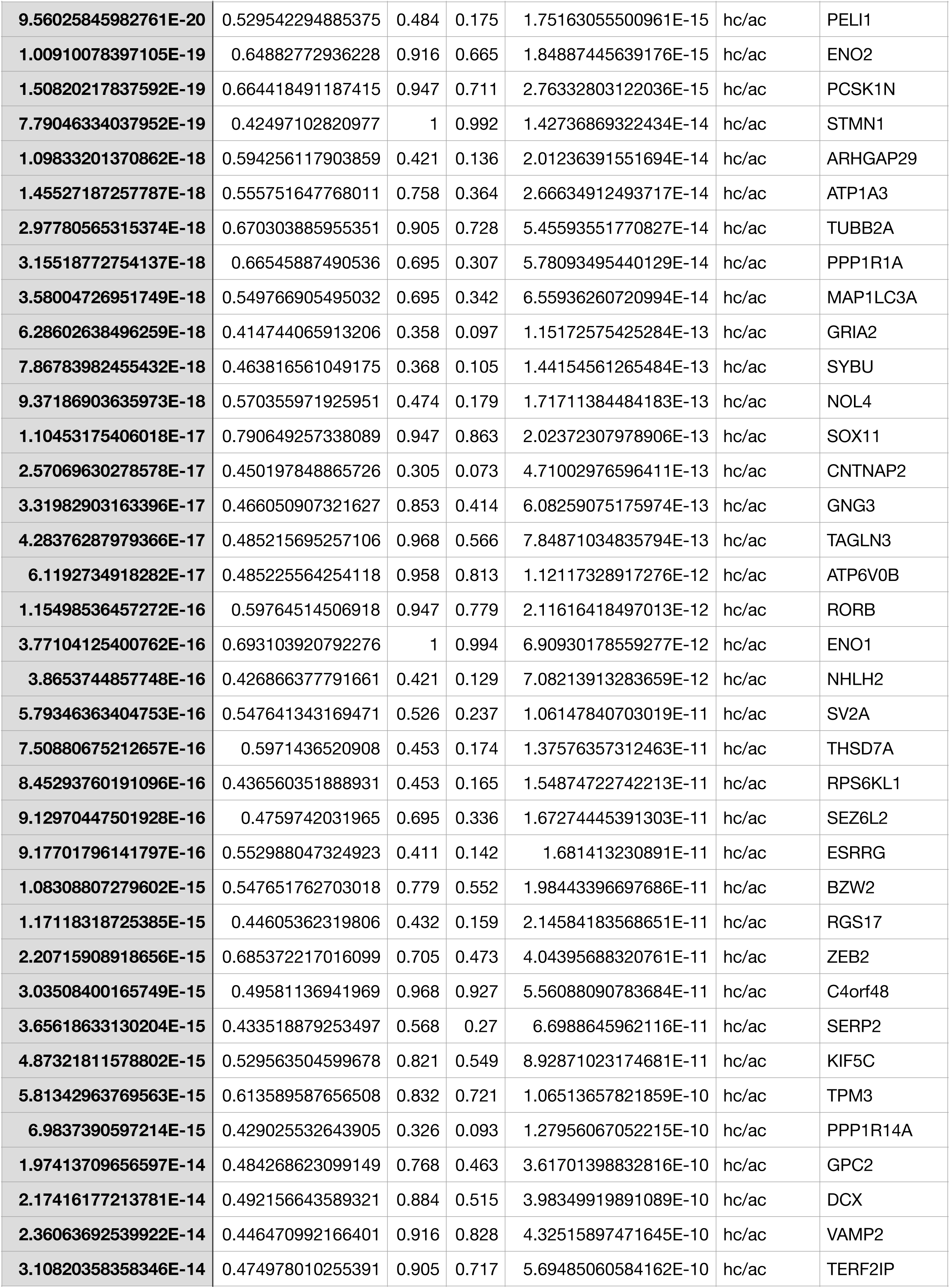
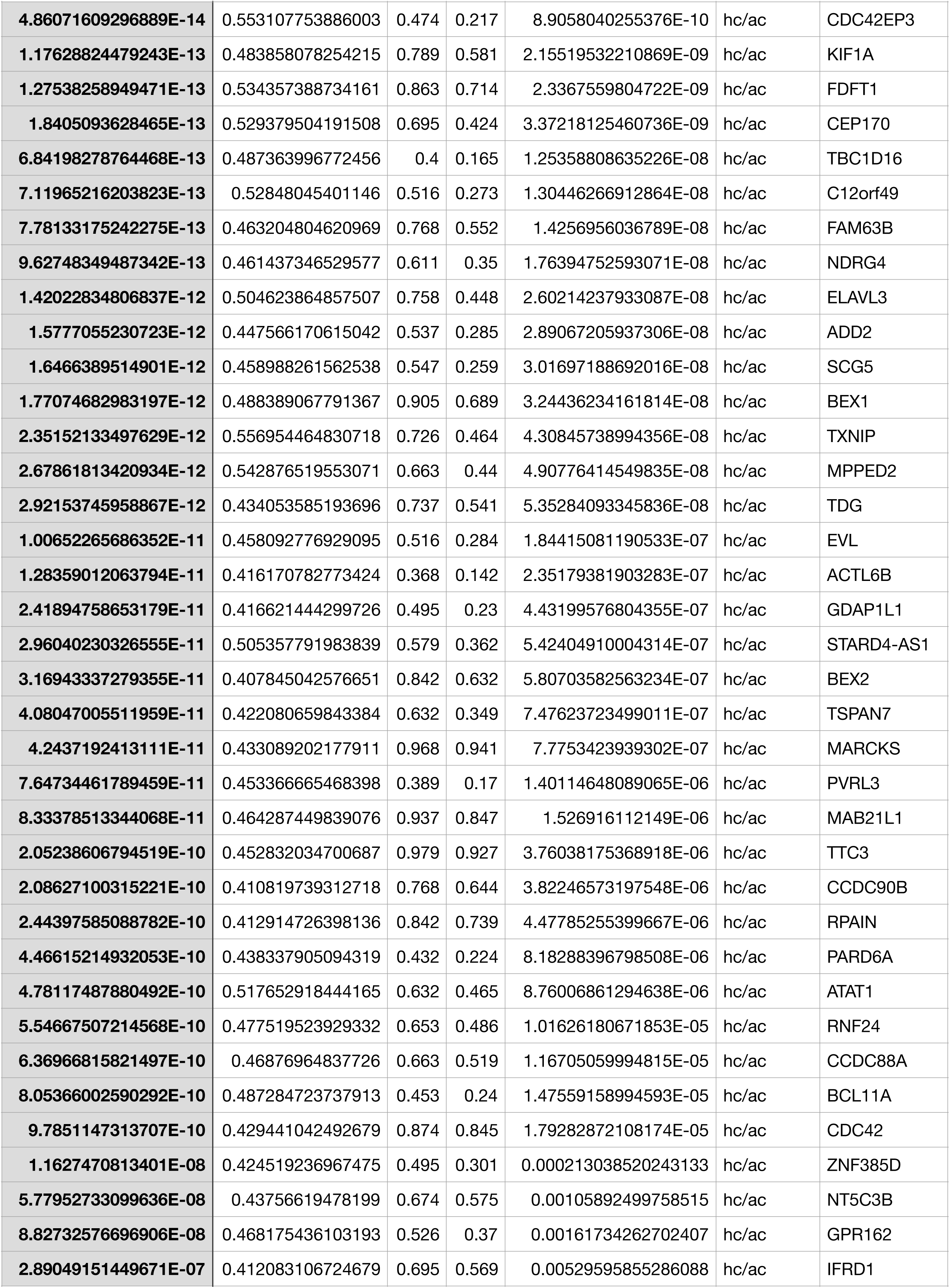
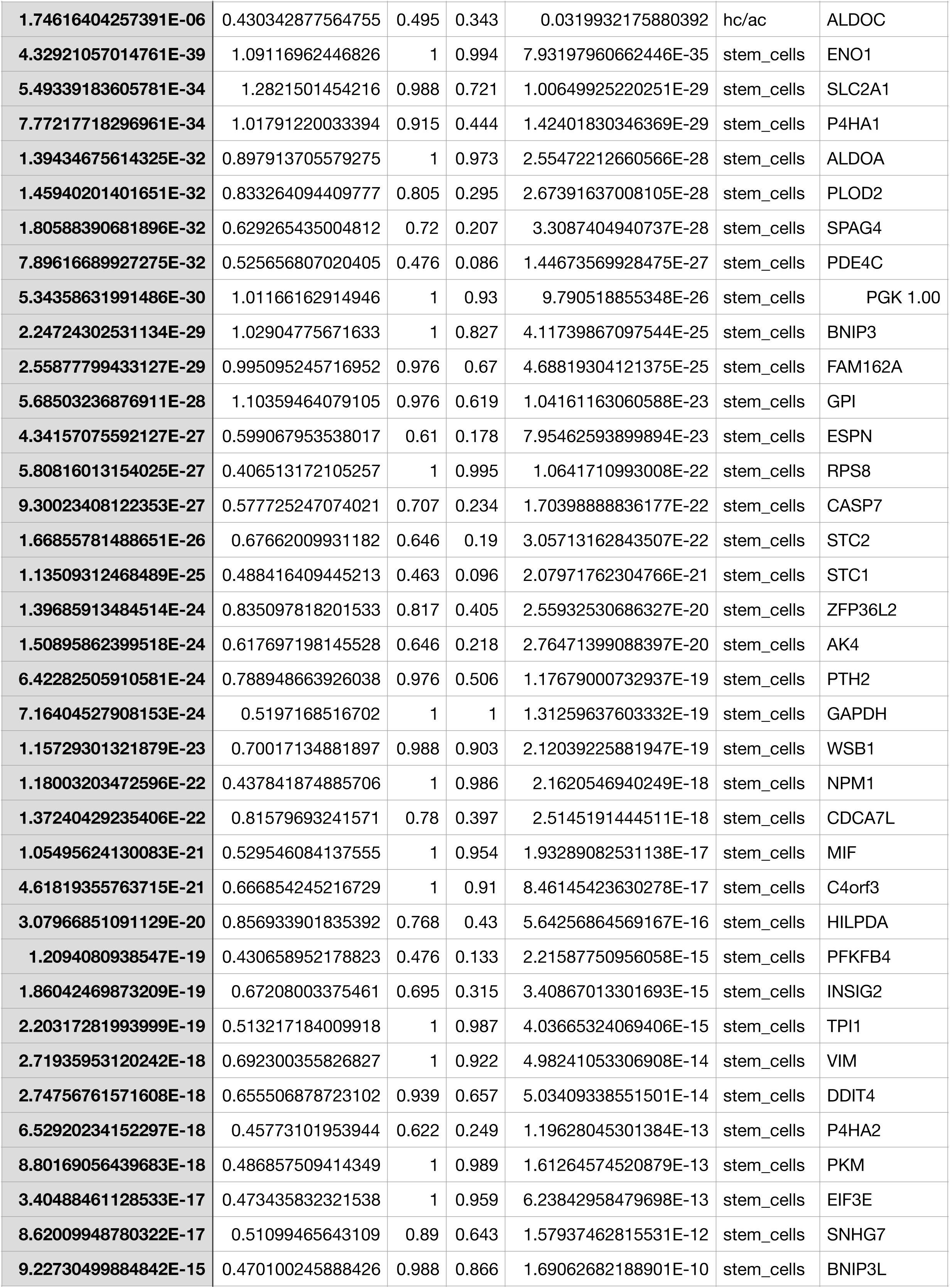
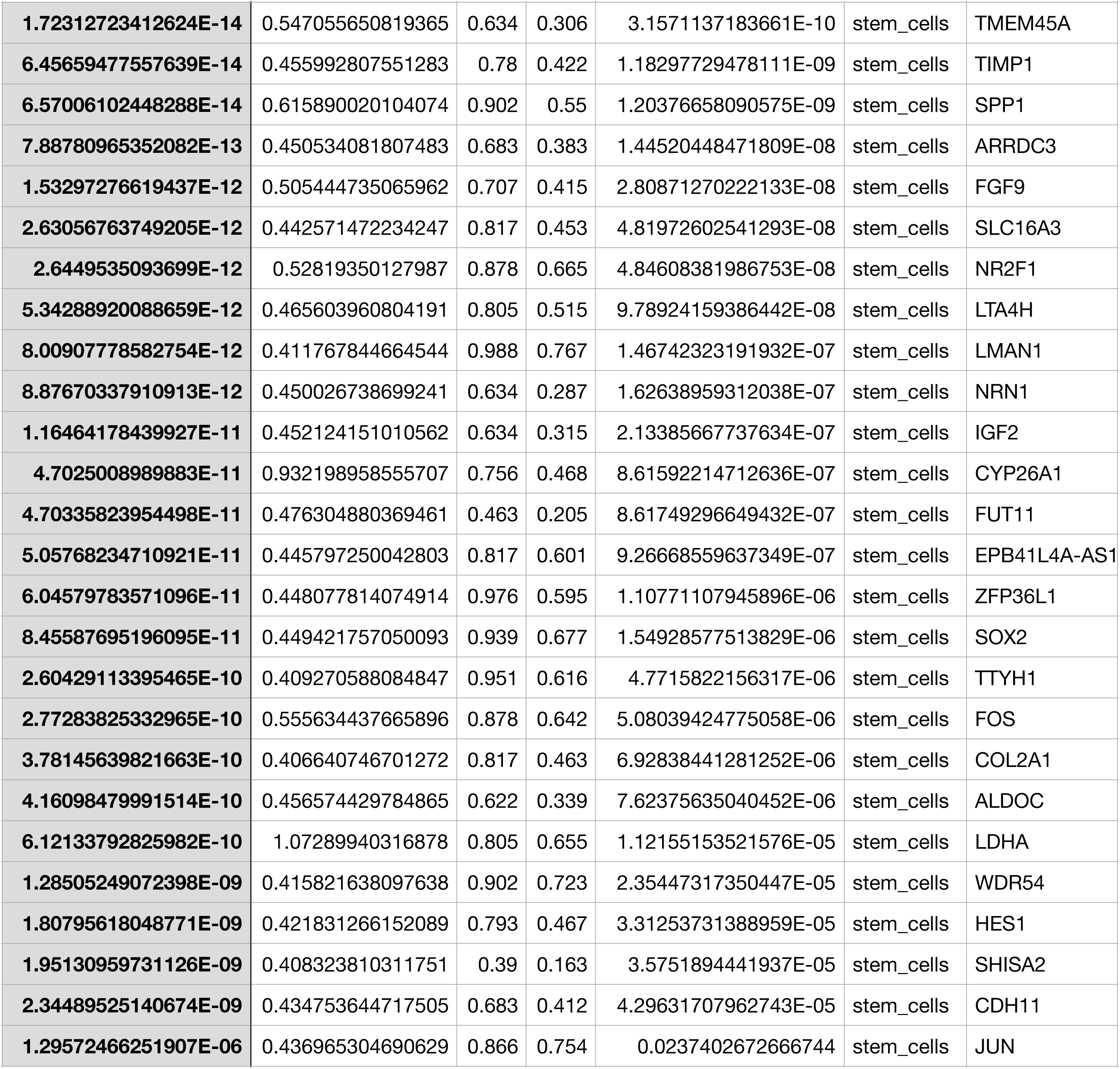
enriched_genes AEP

**SupTable 3.**
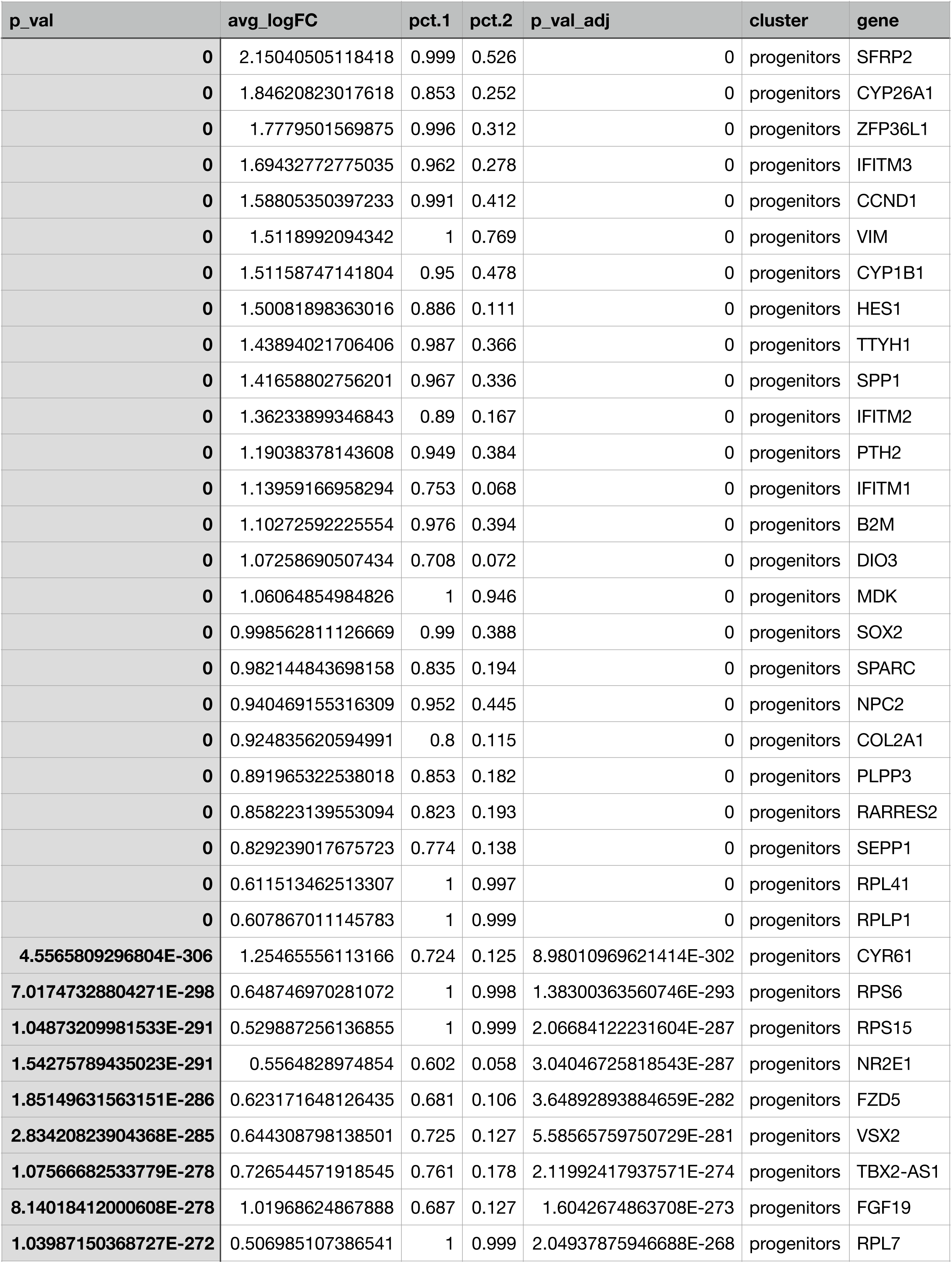
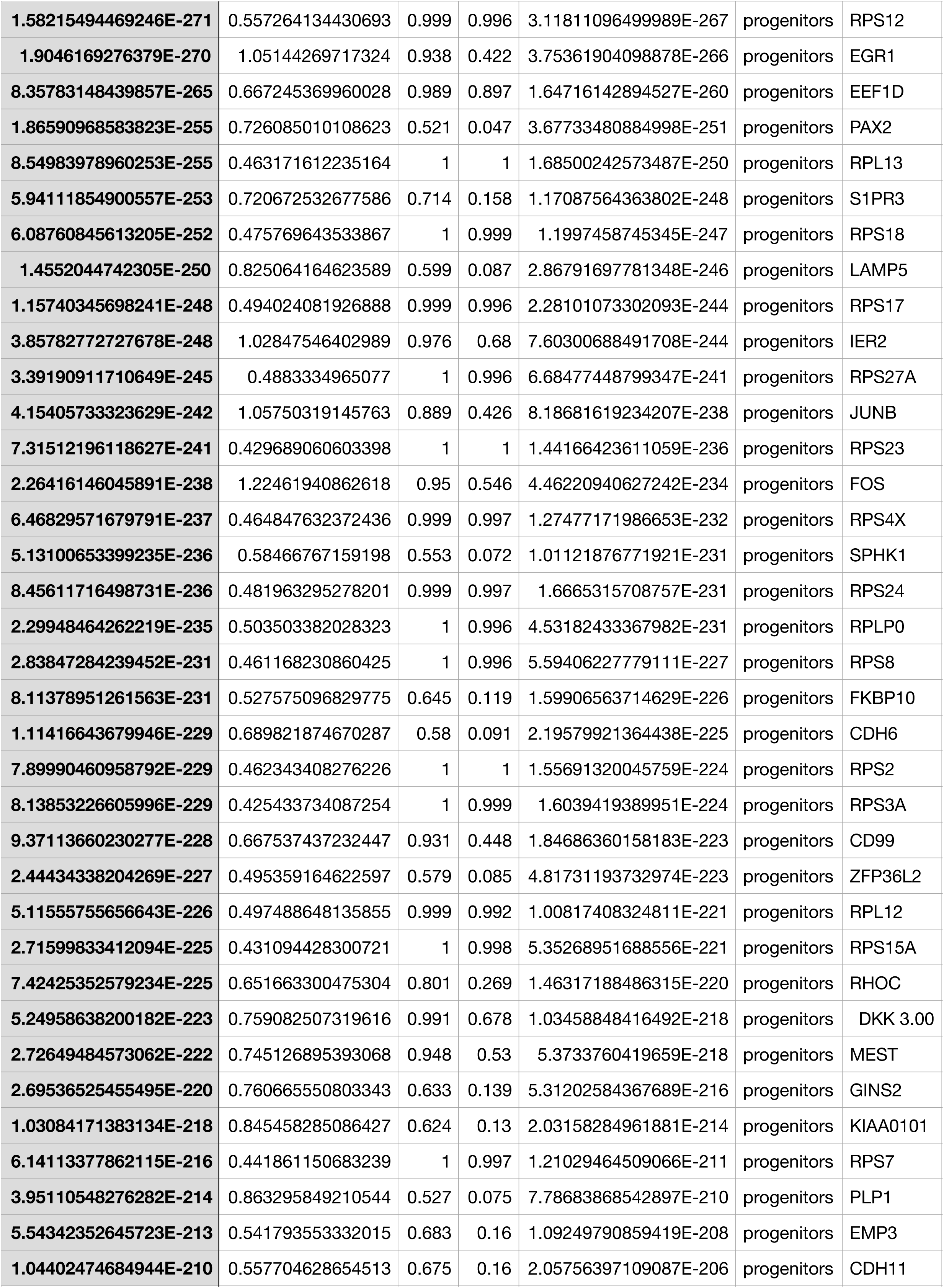
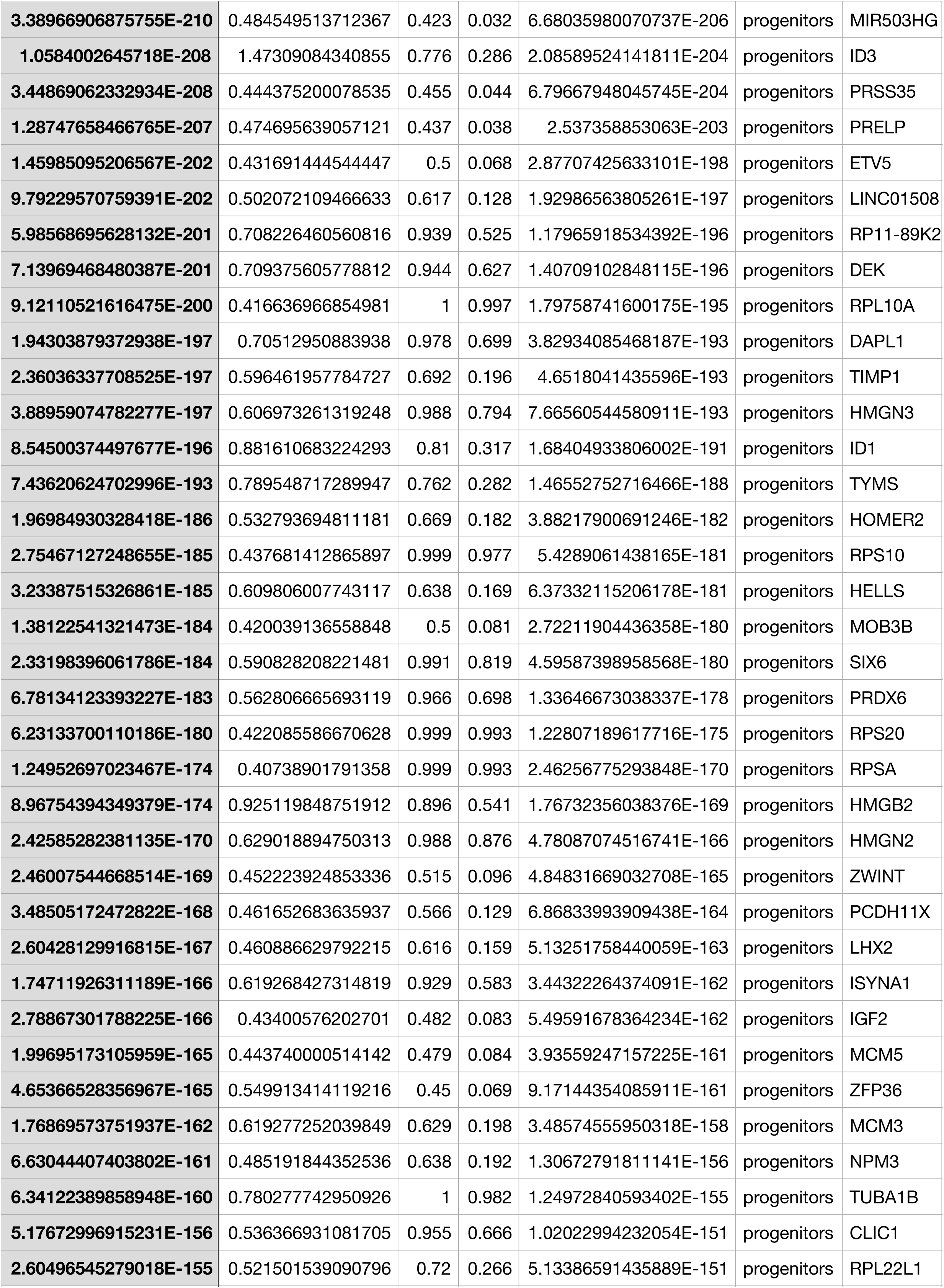
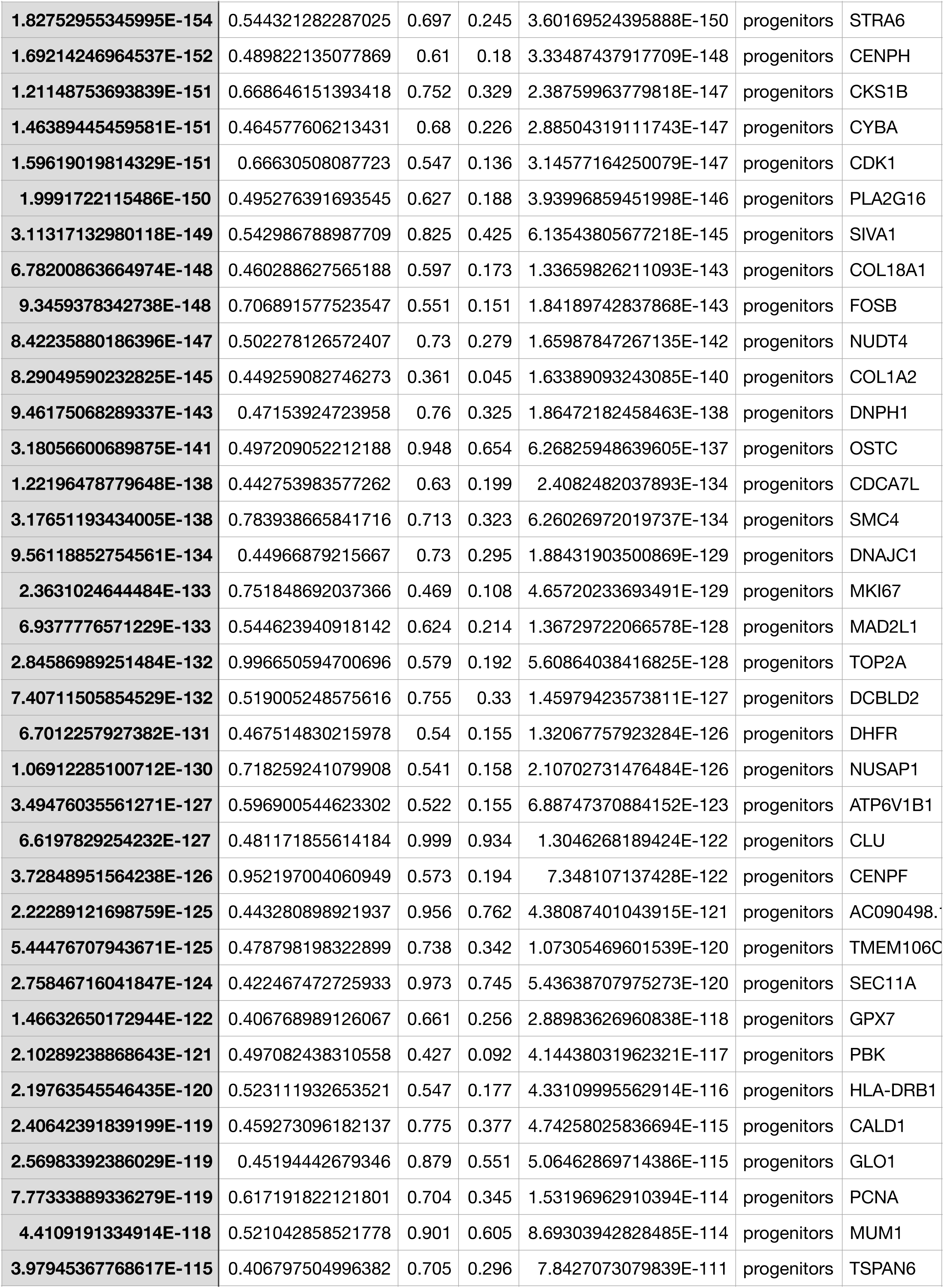
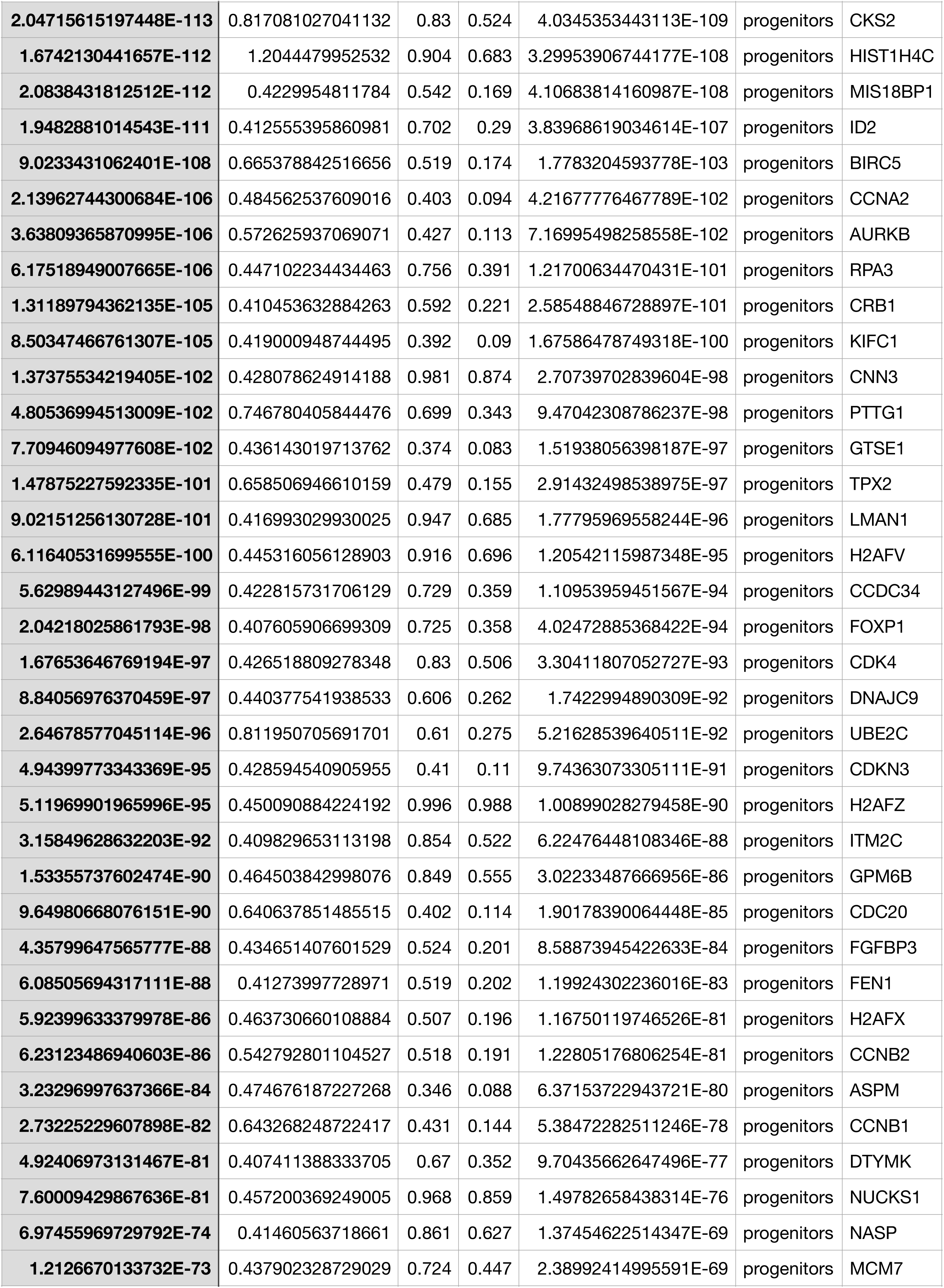
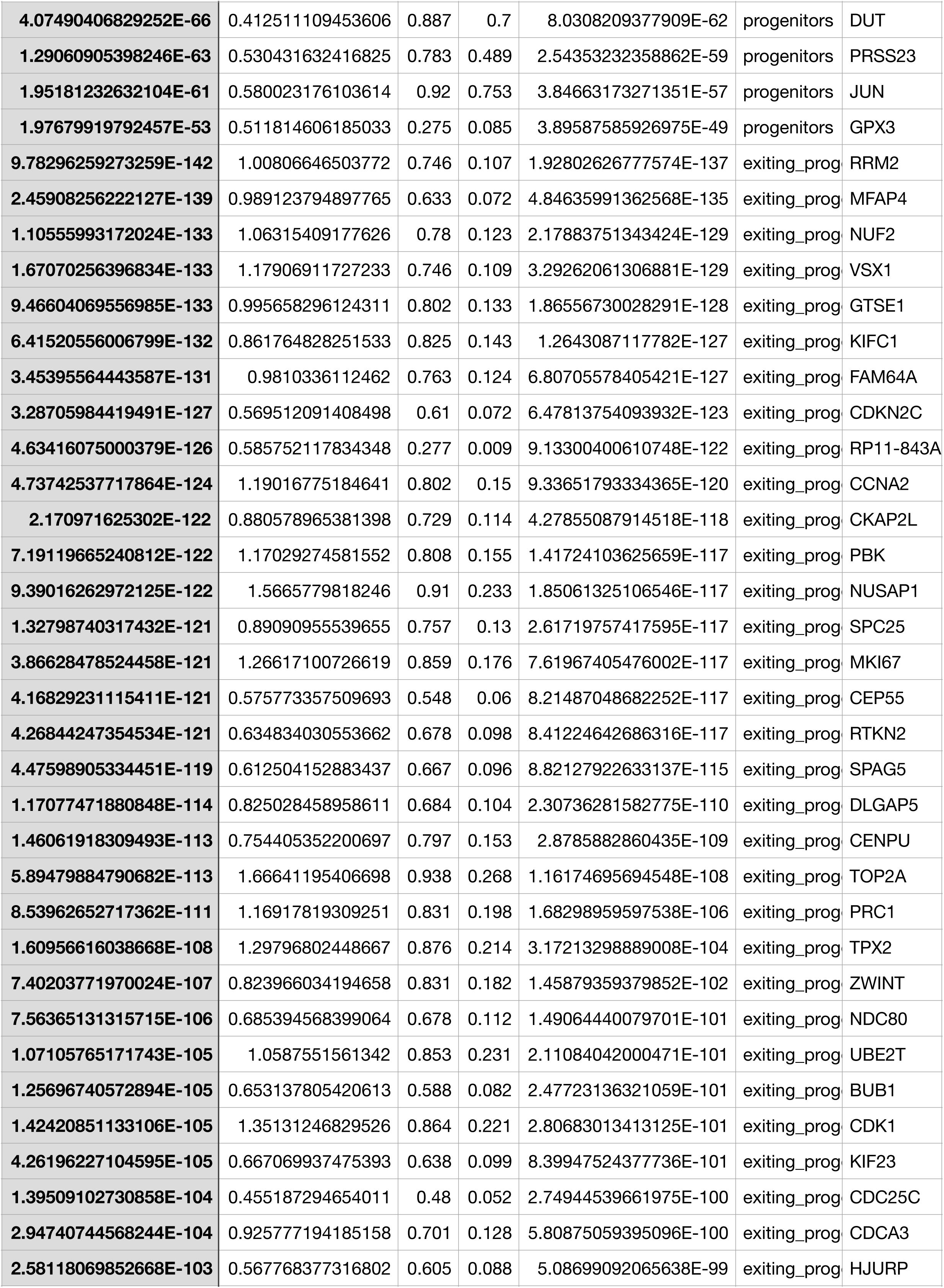
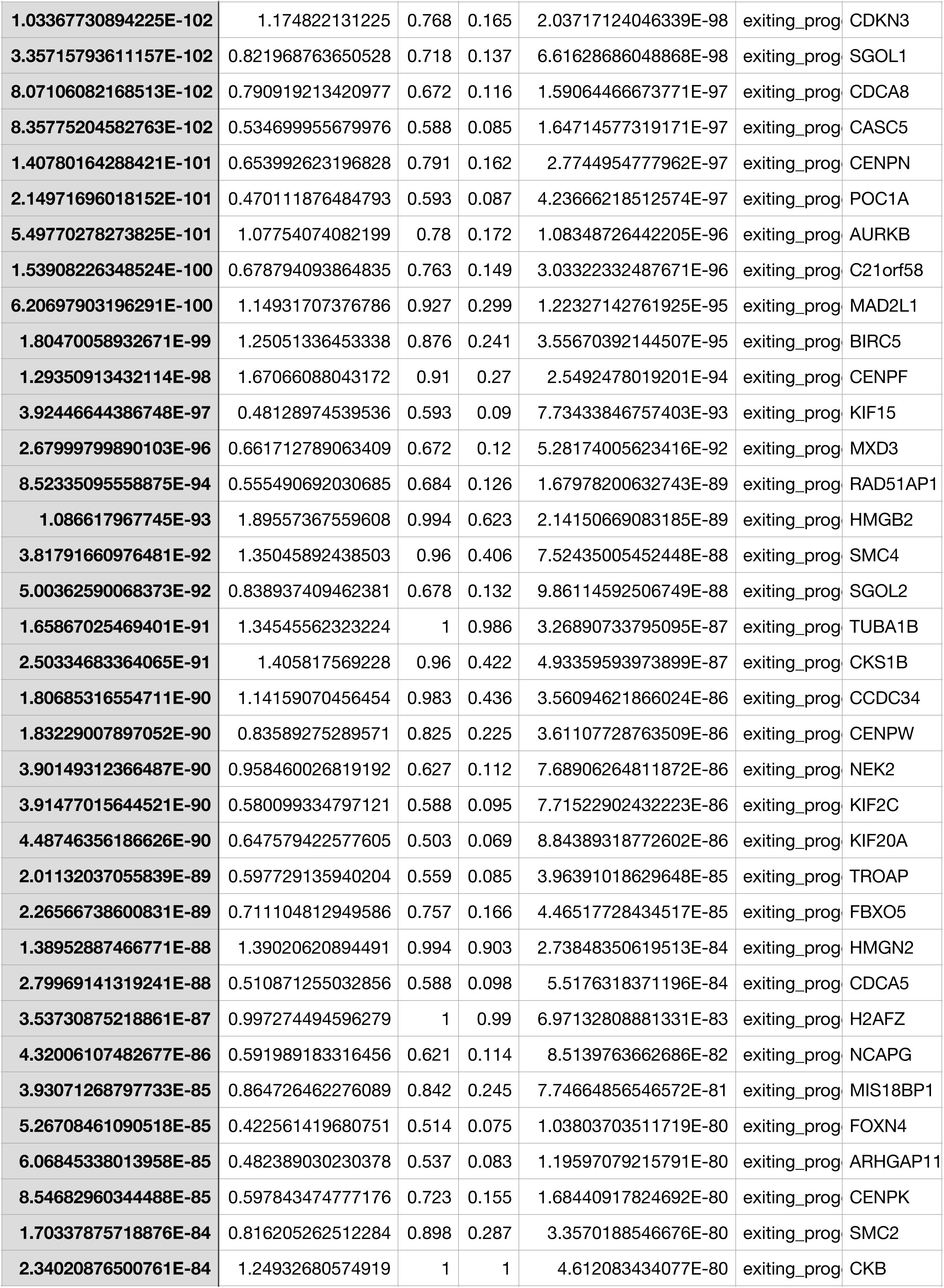
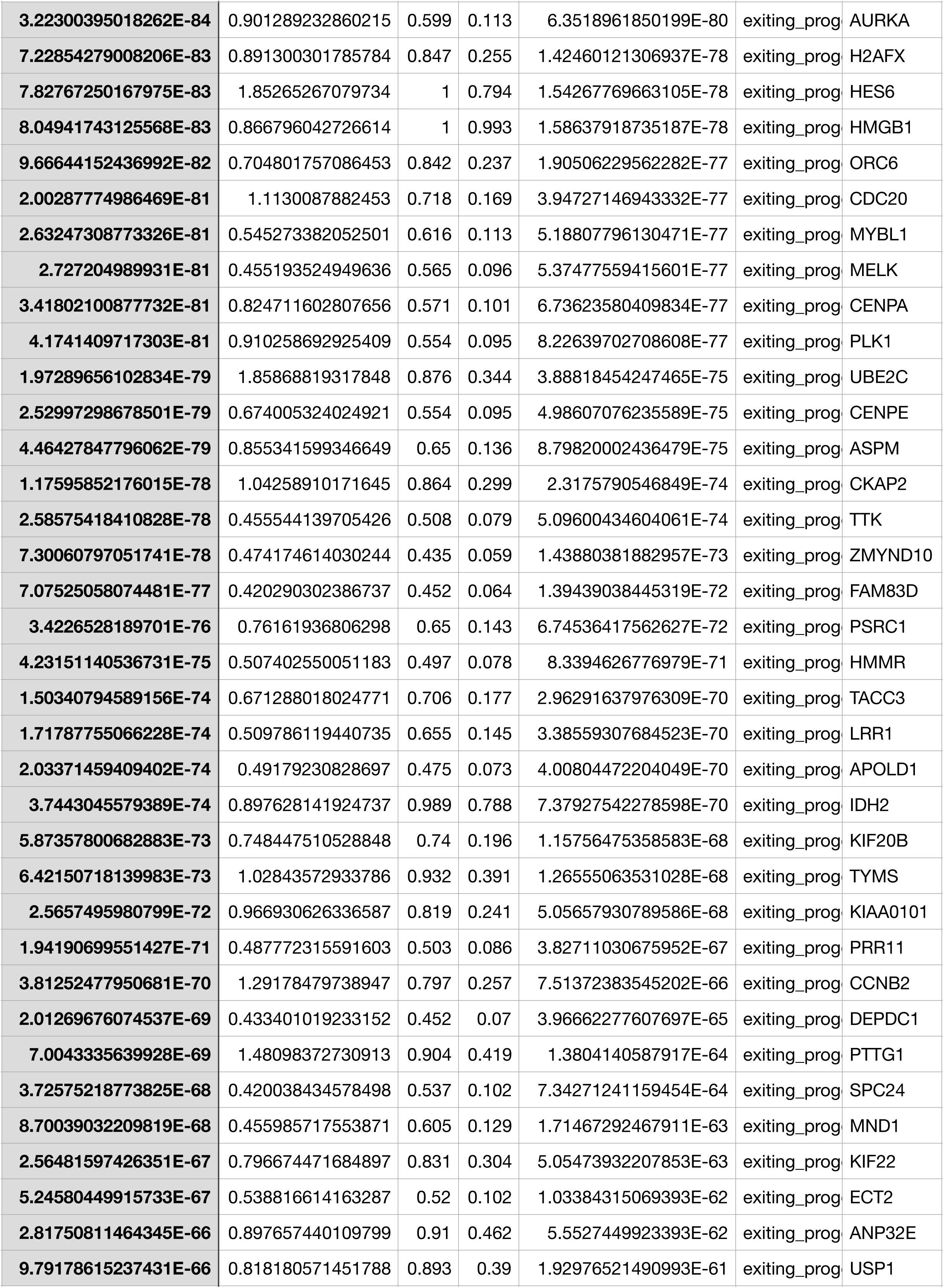
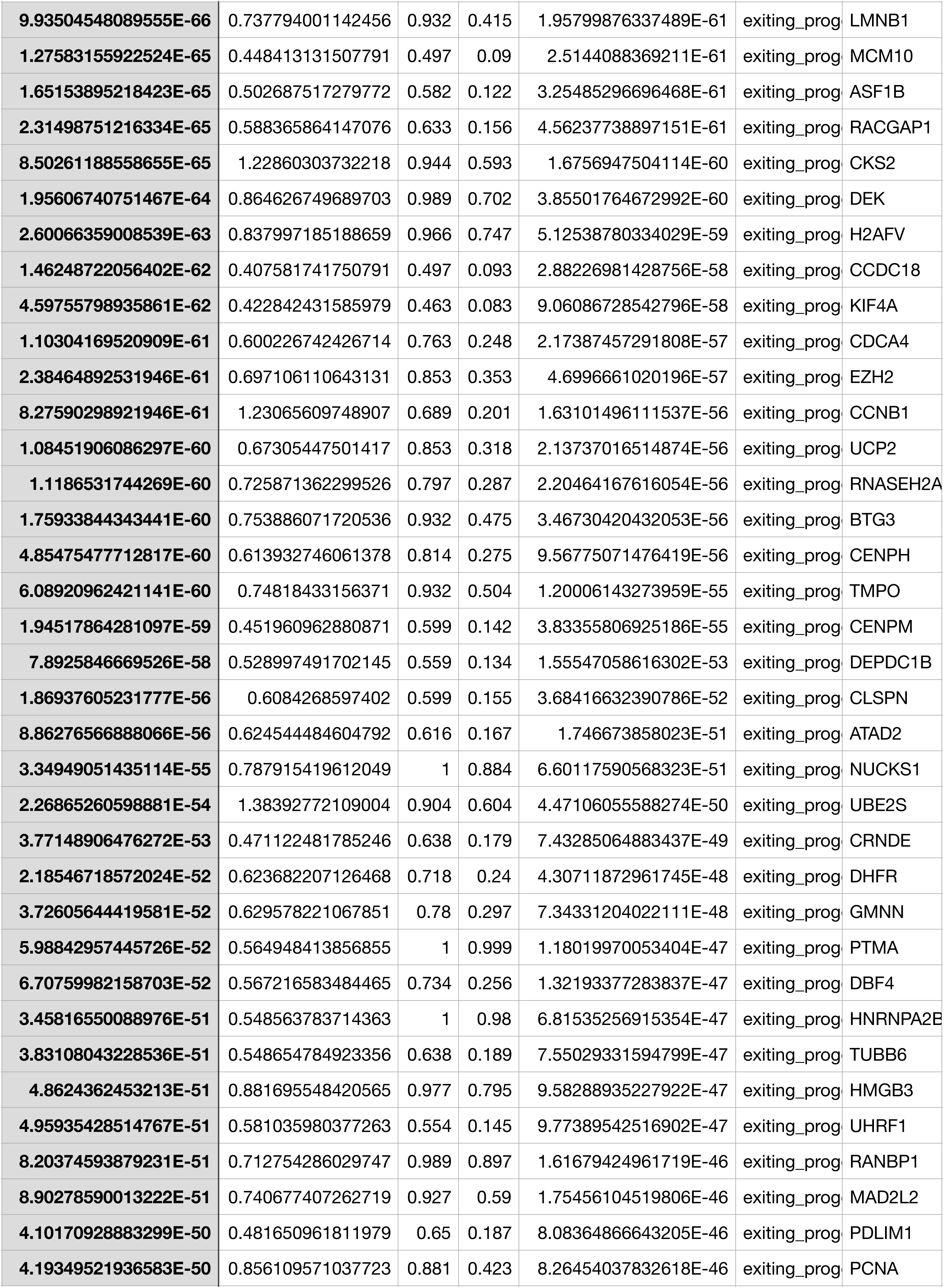
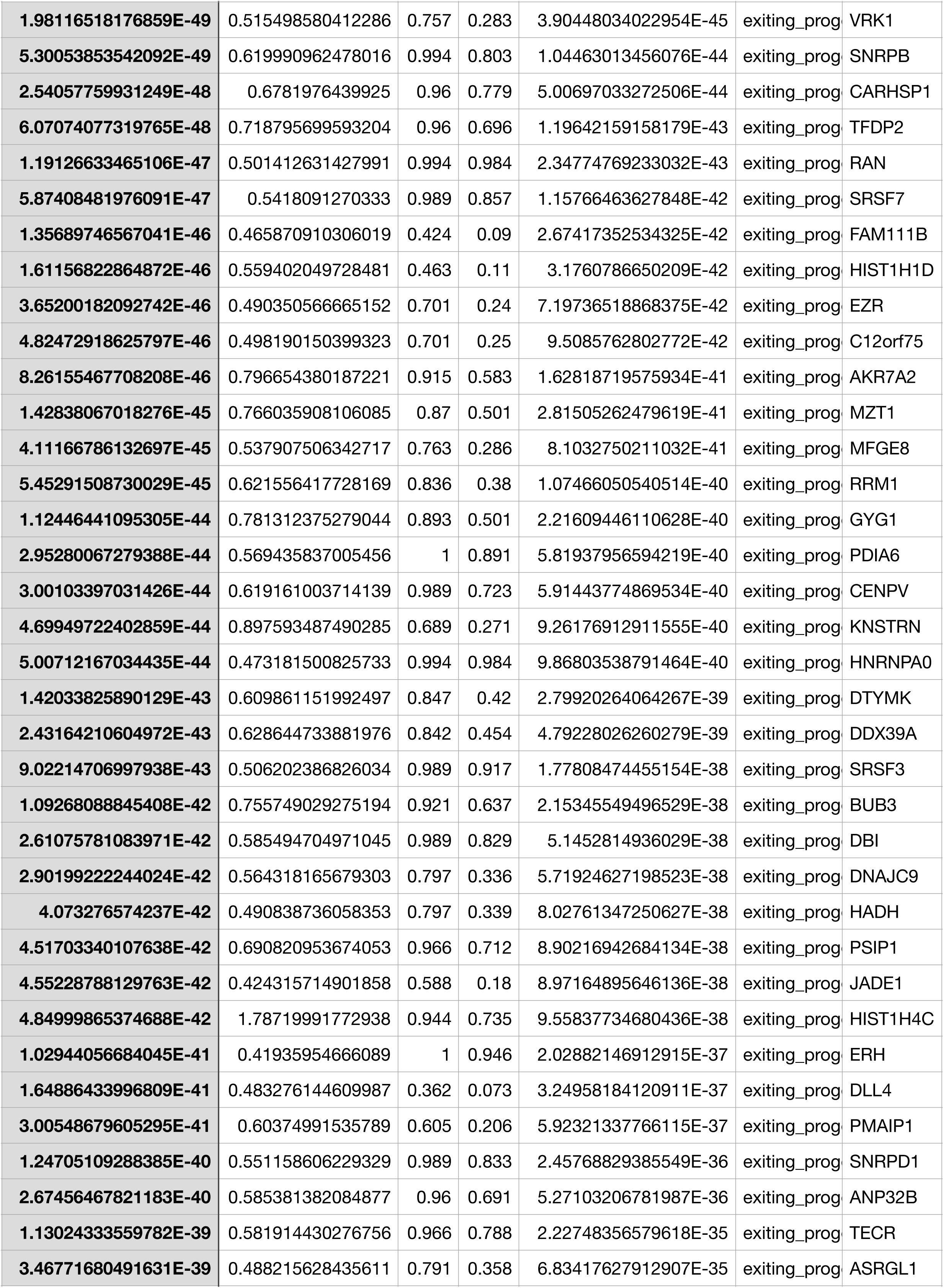
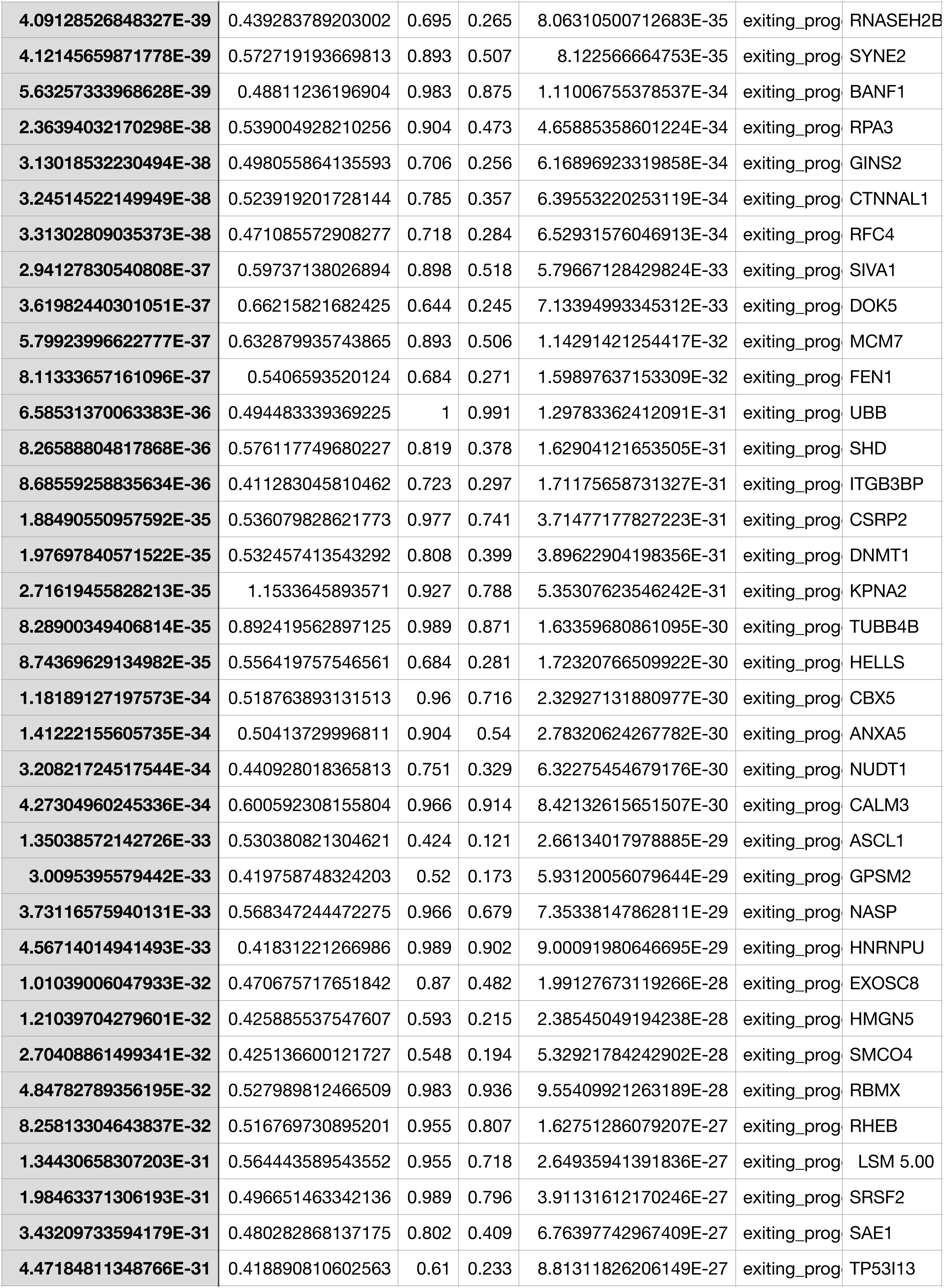
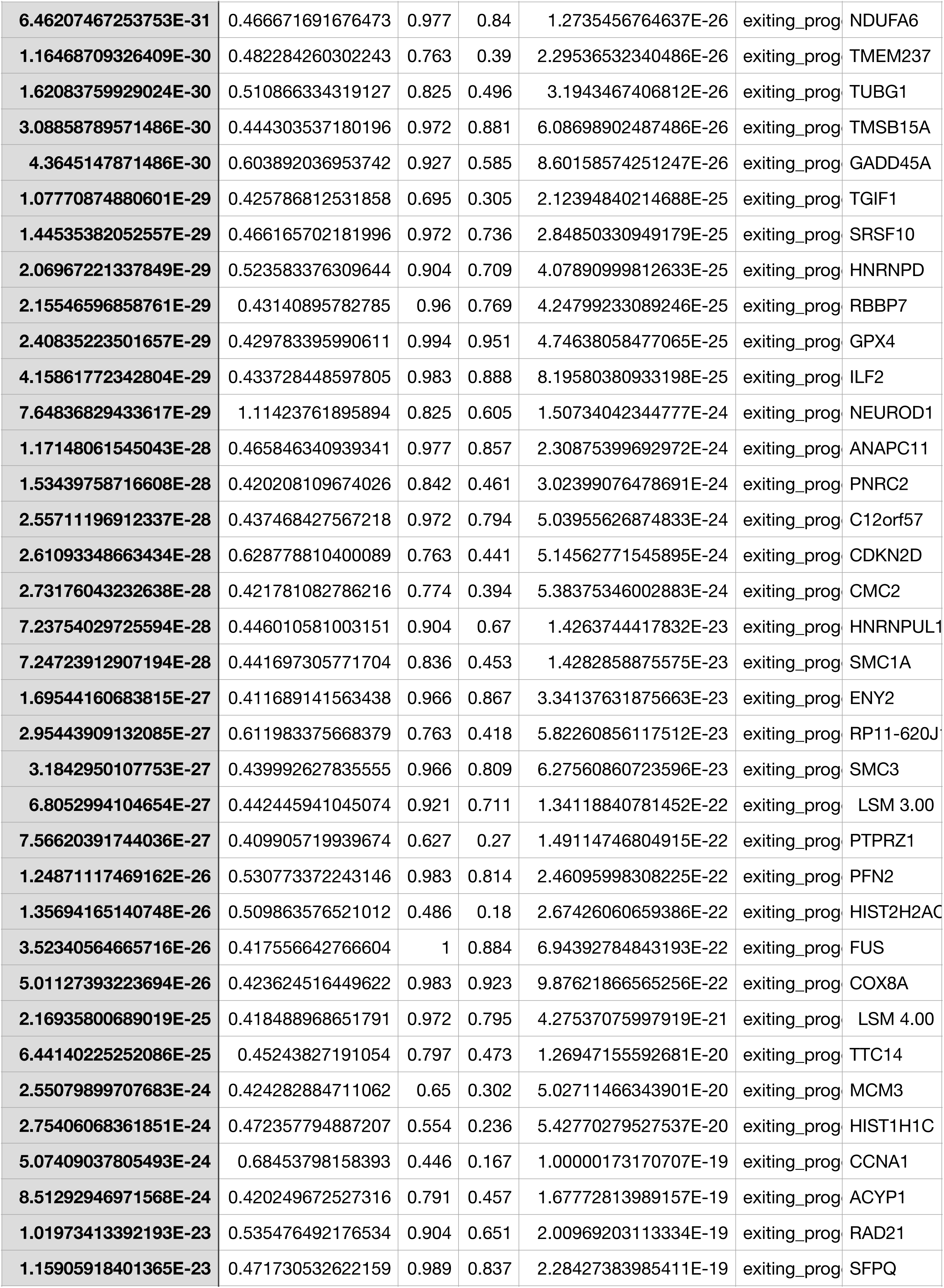
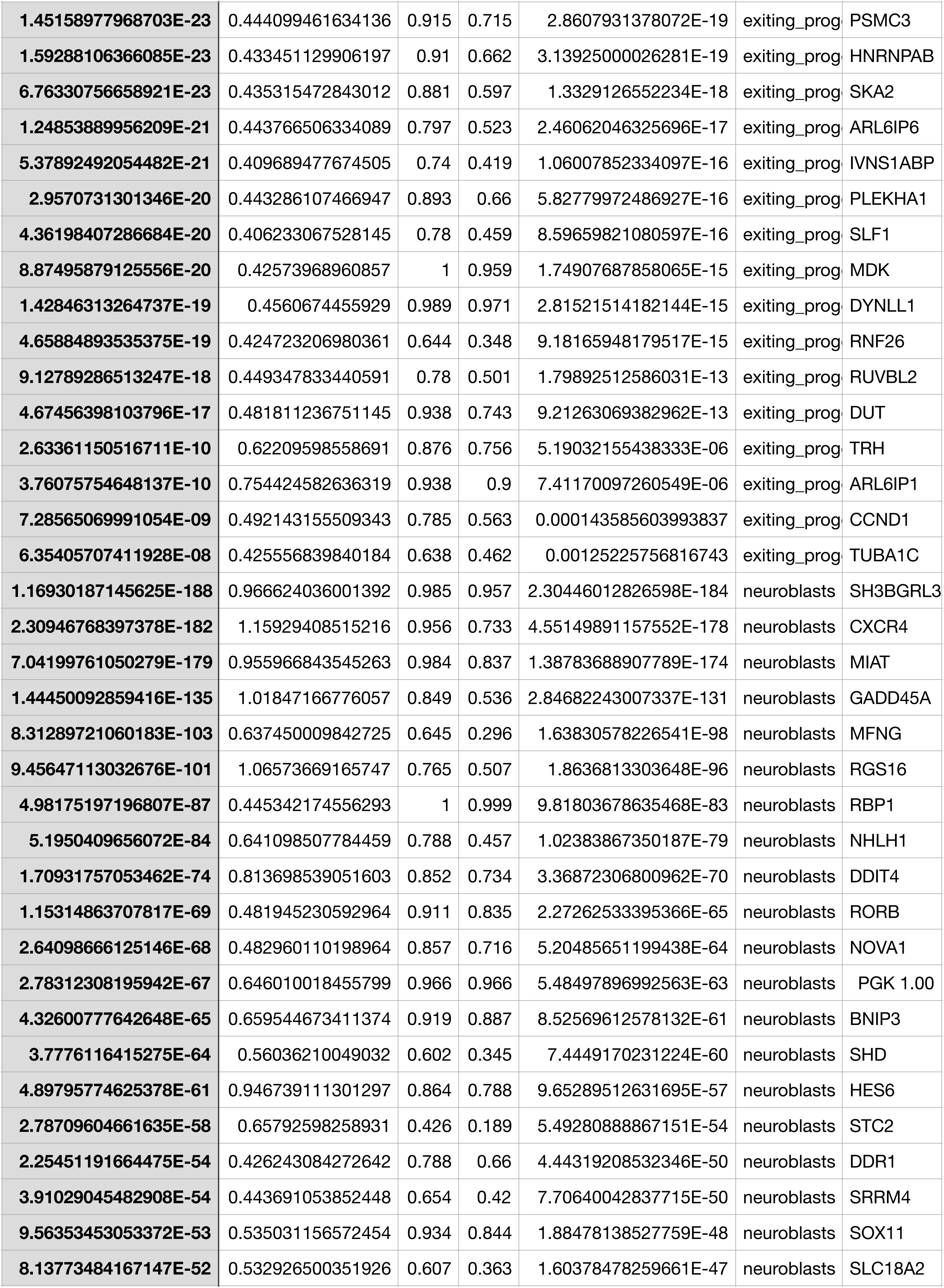
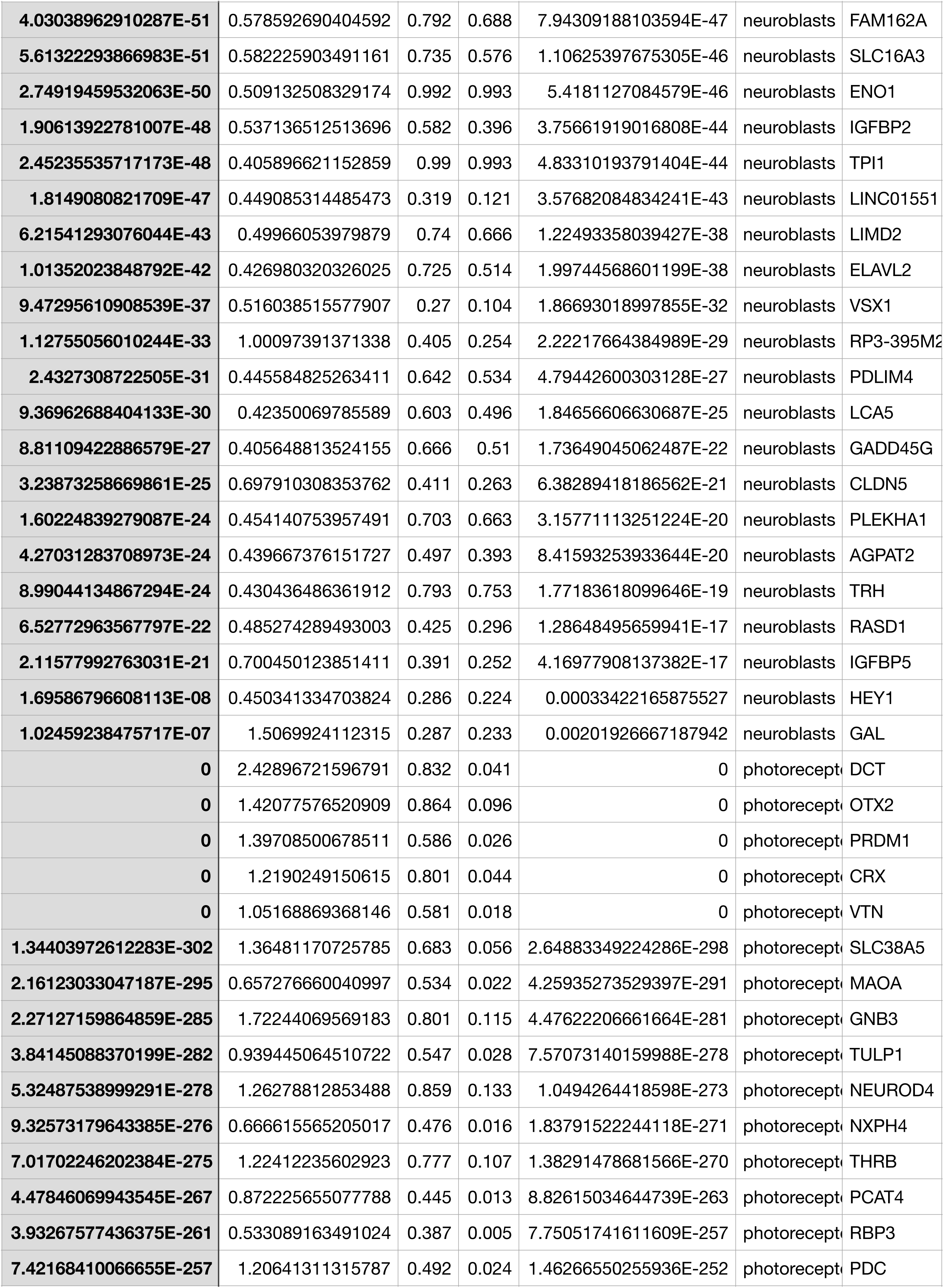
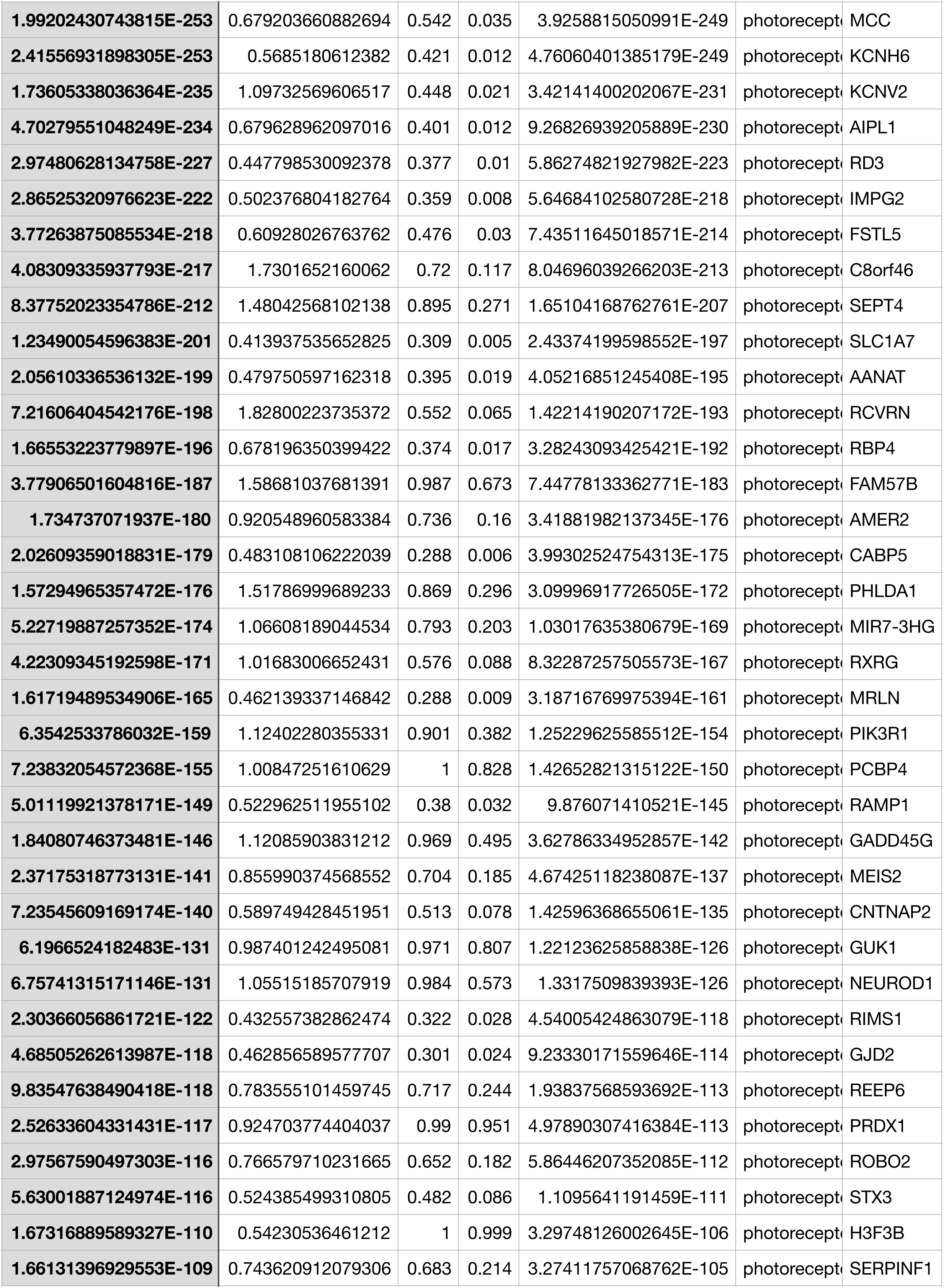
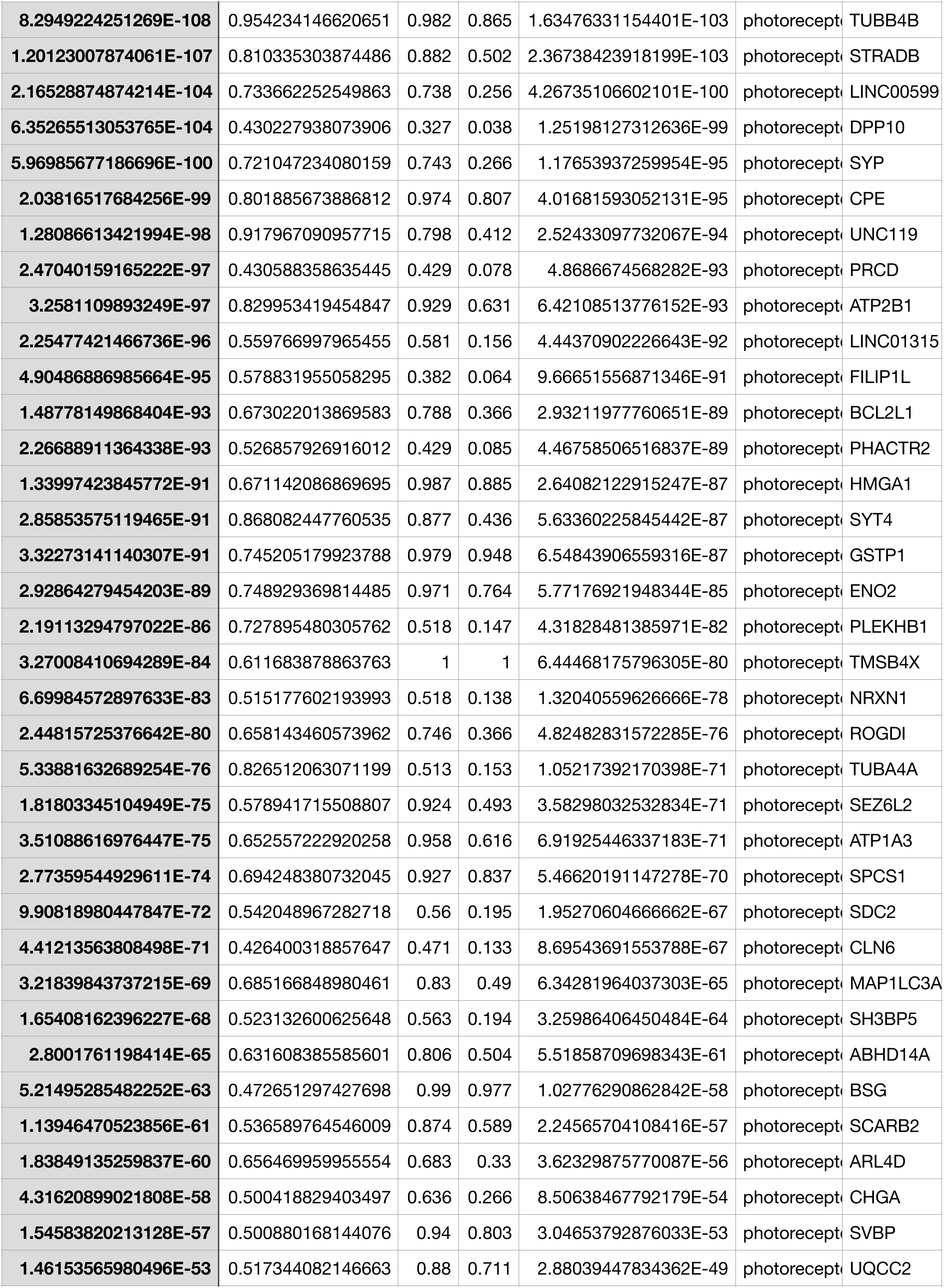
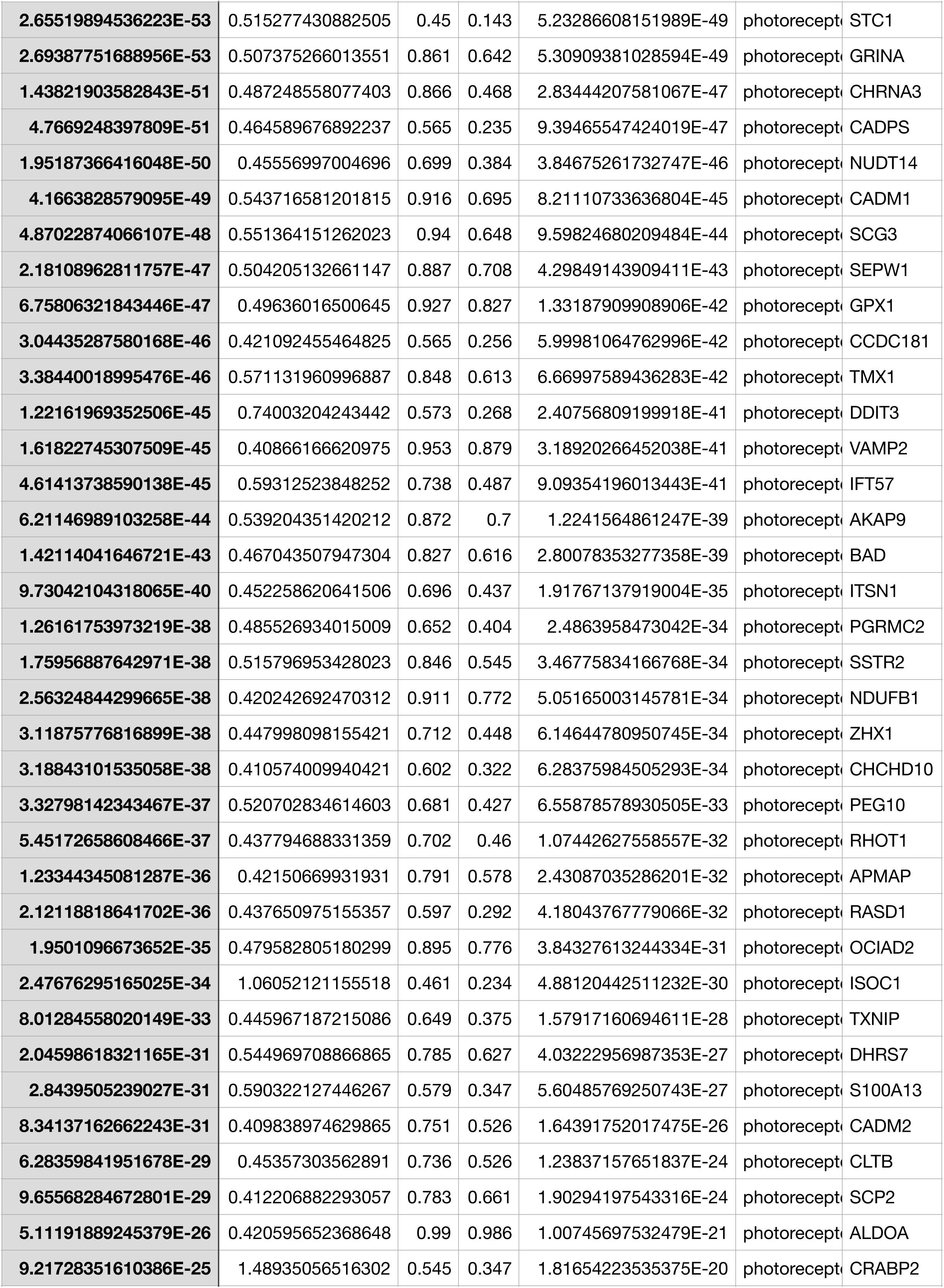
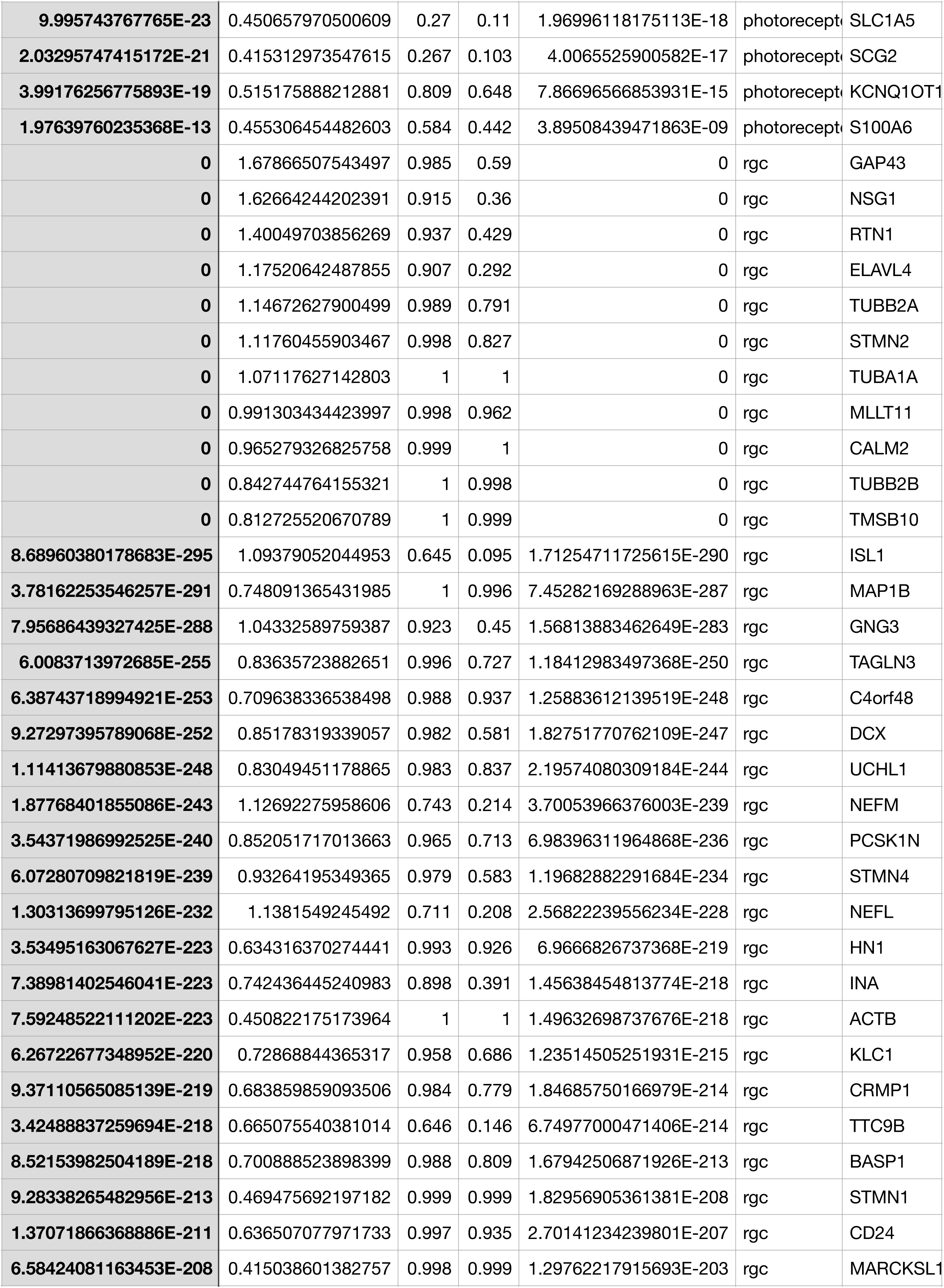
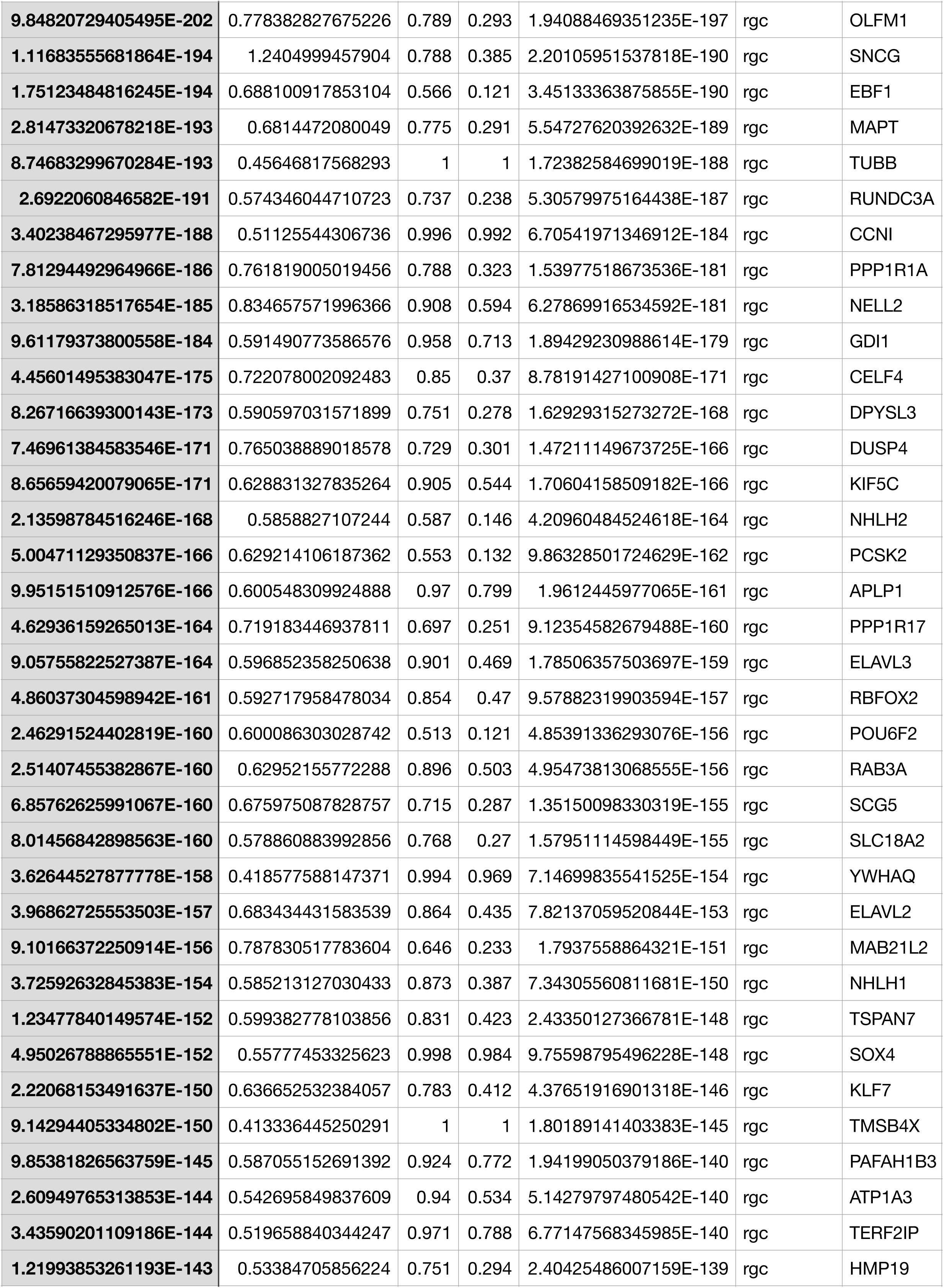
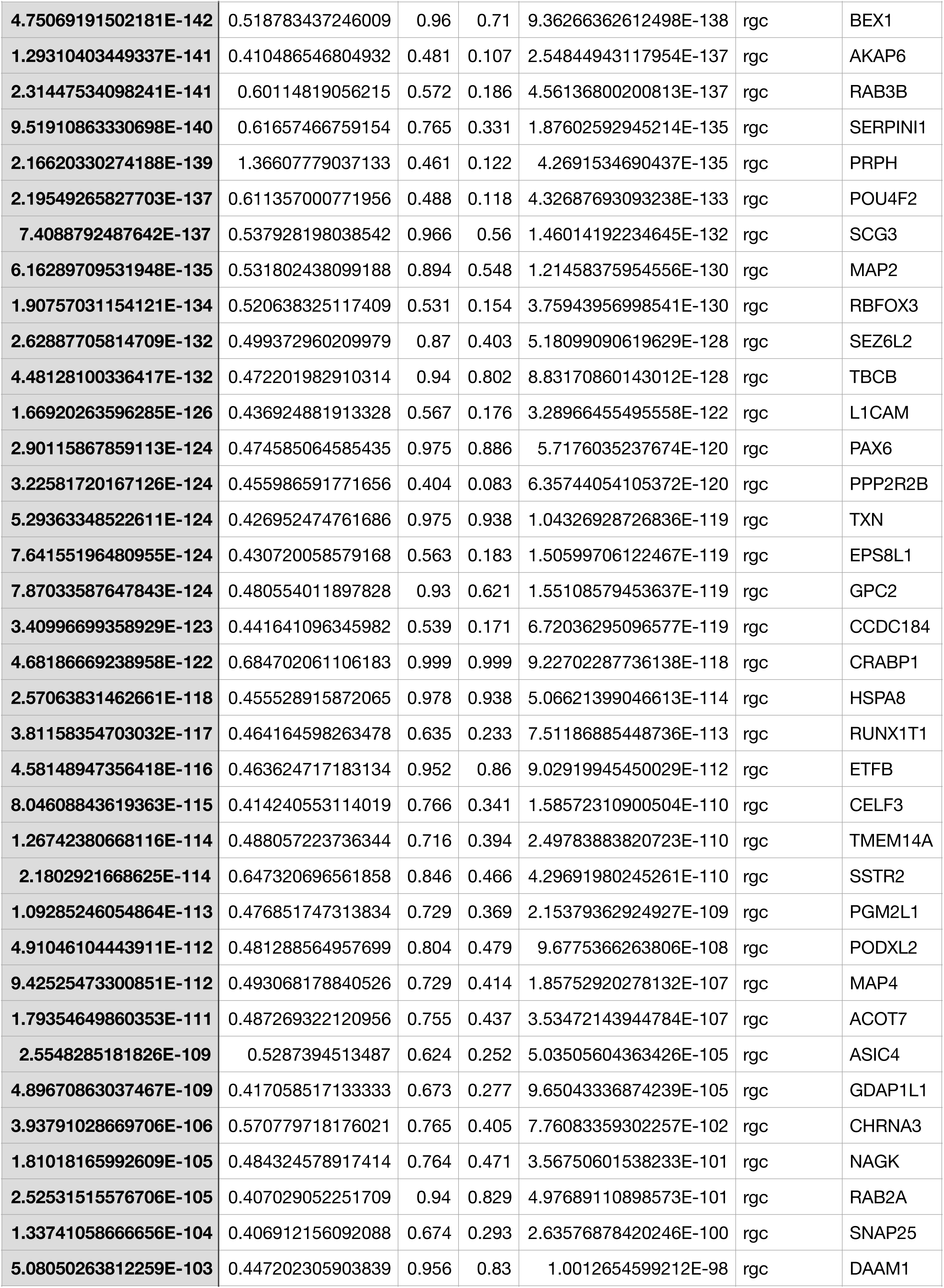
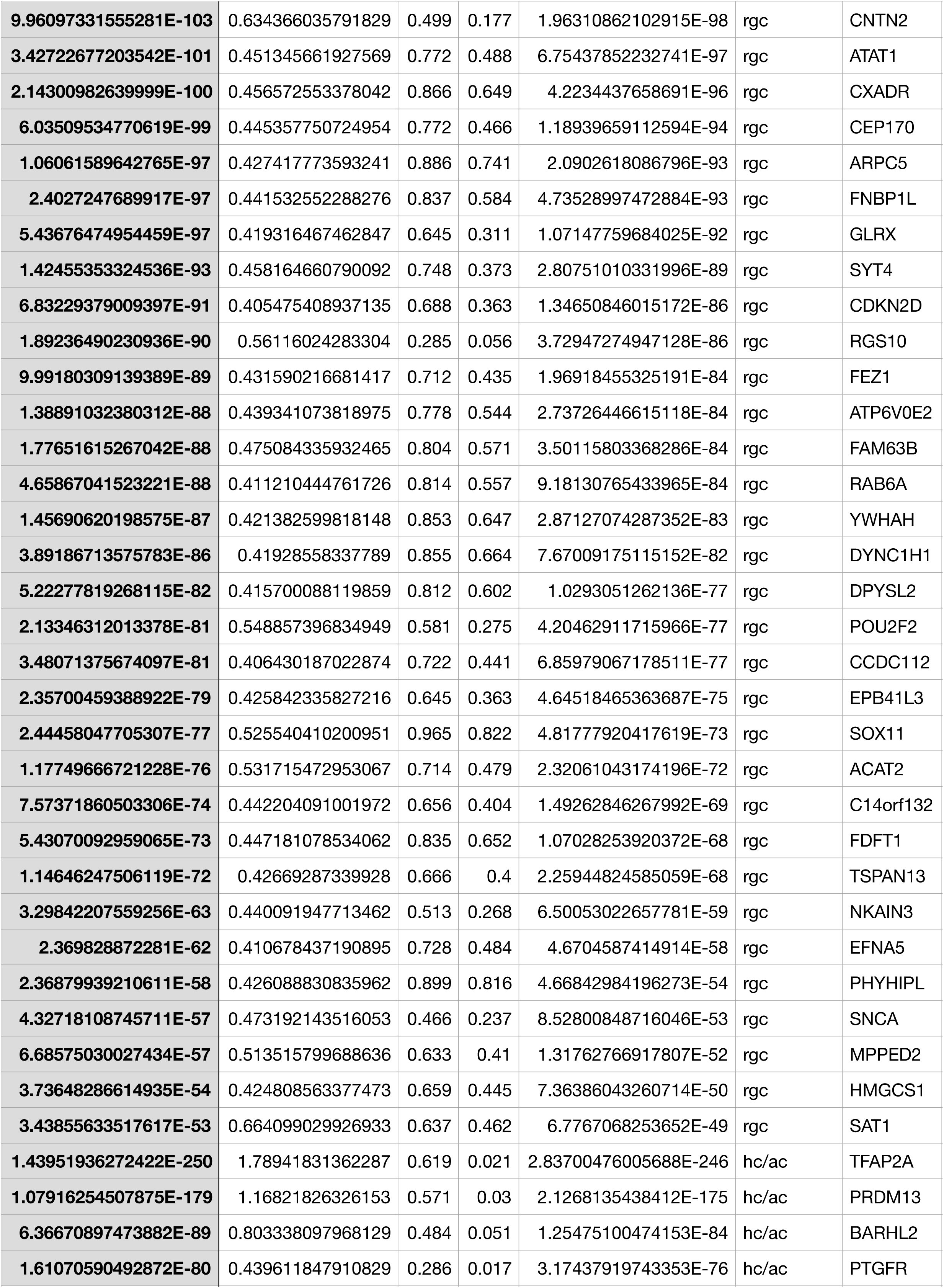
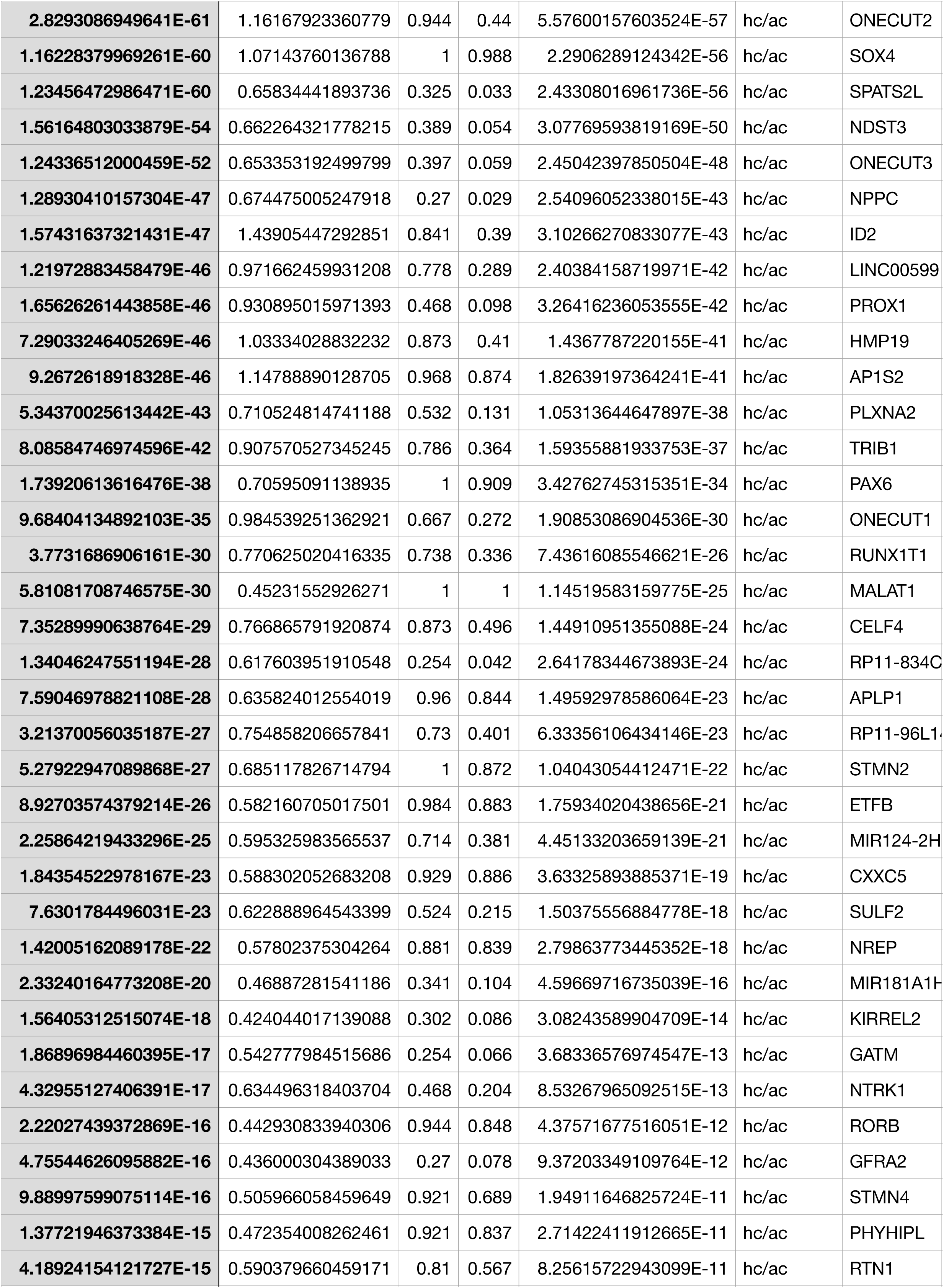
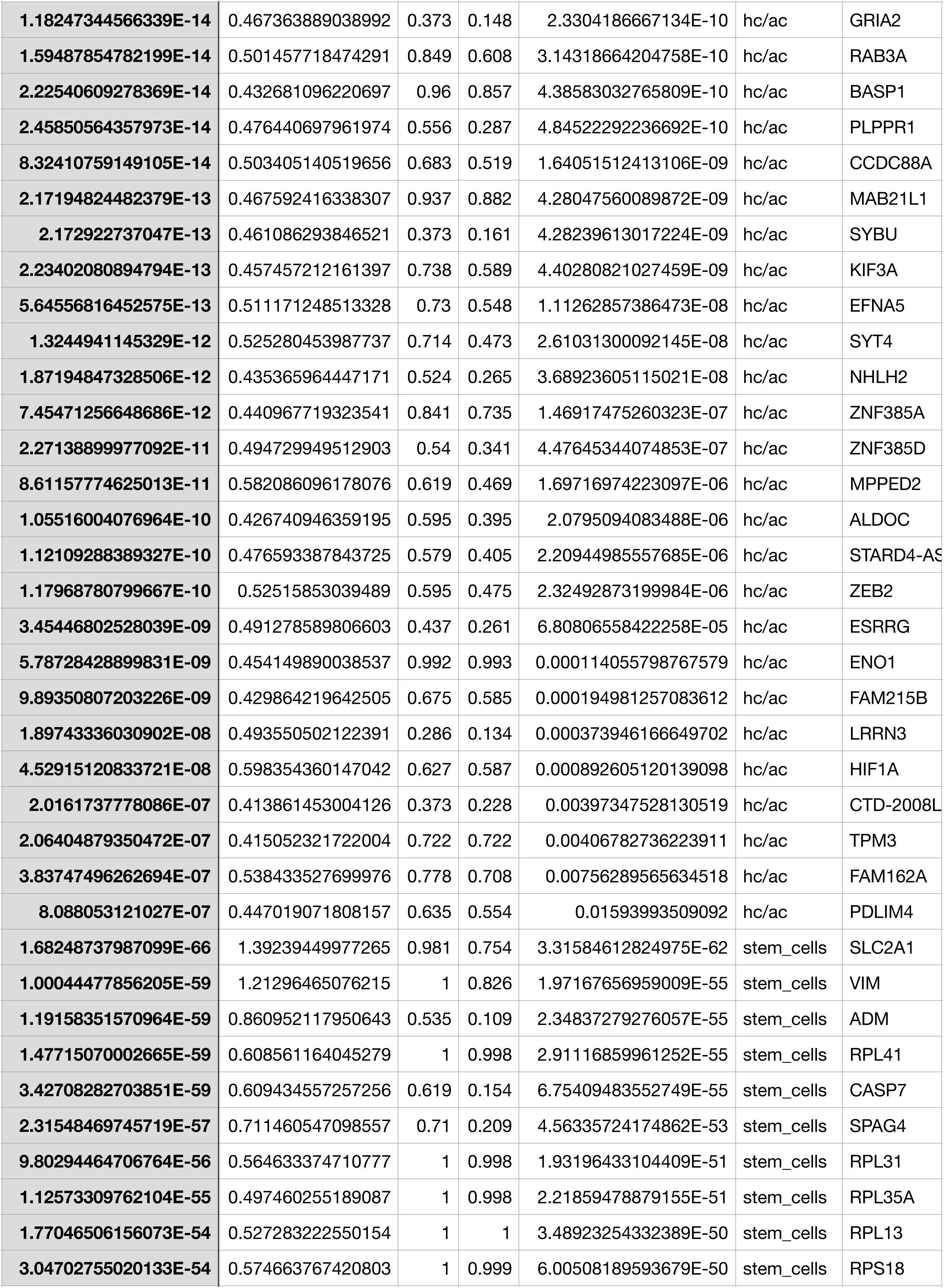
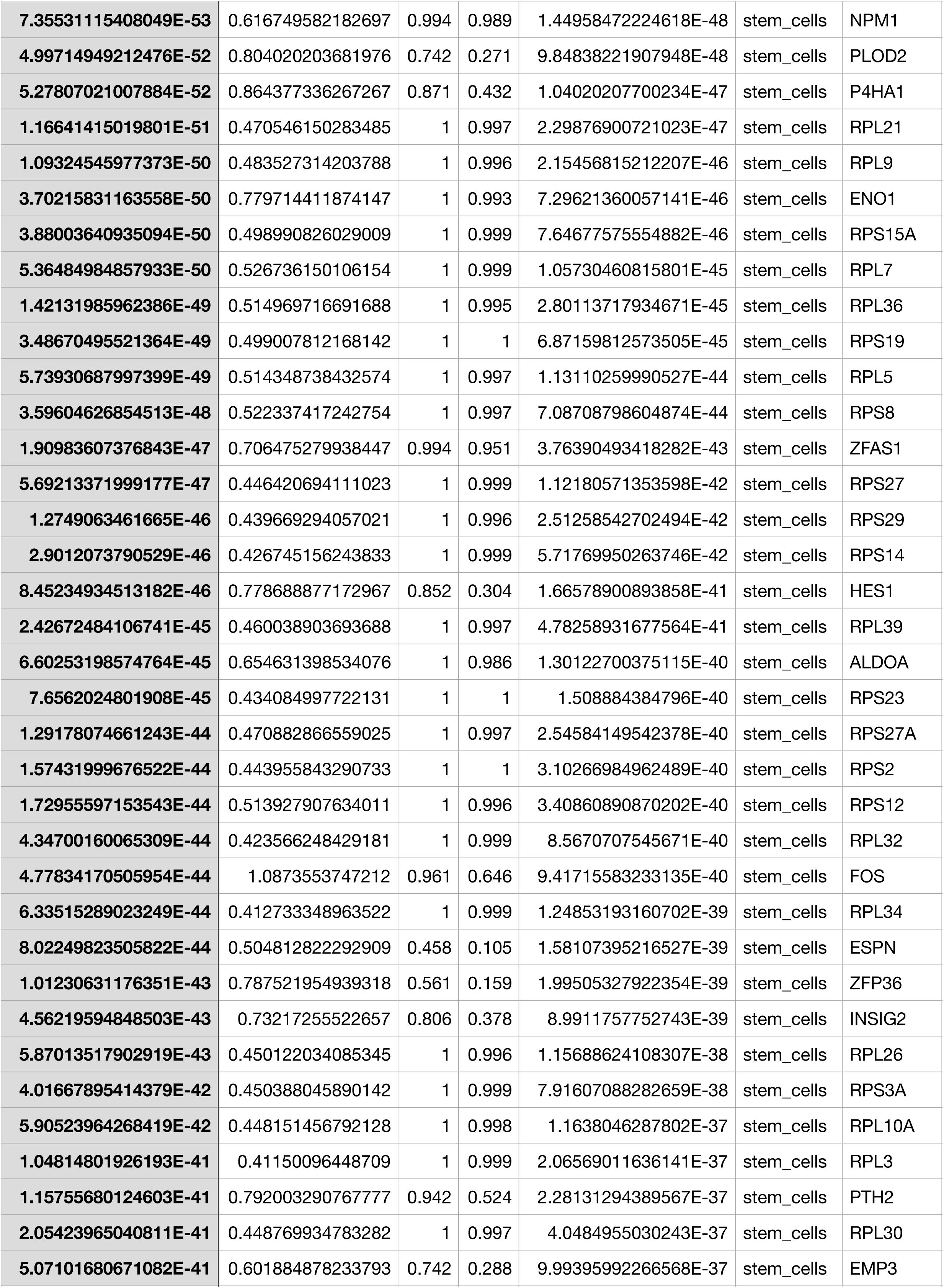
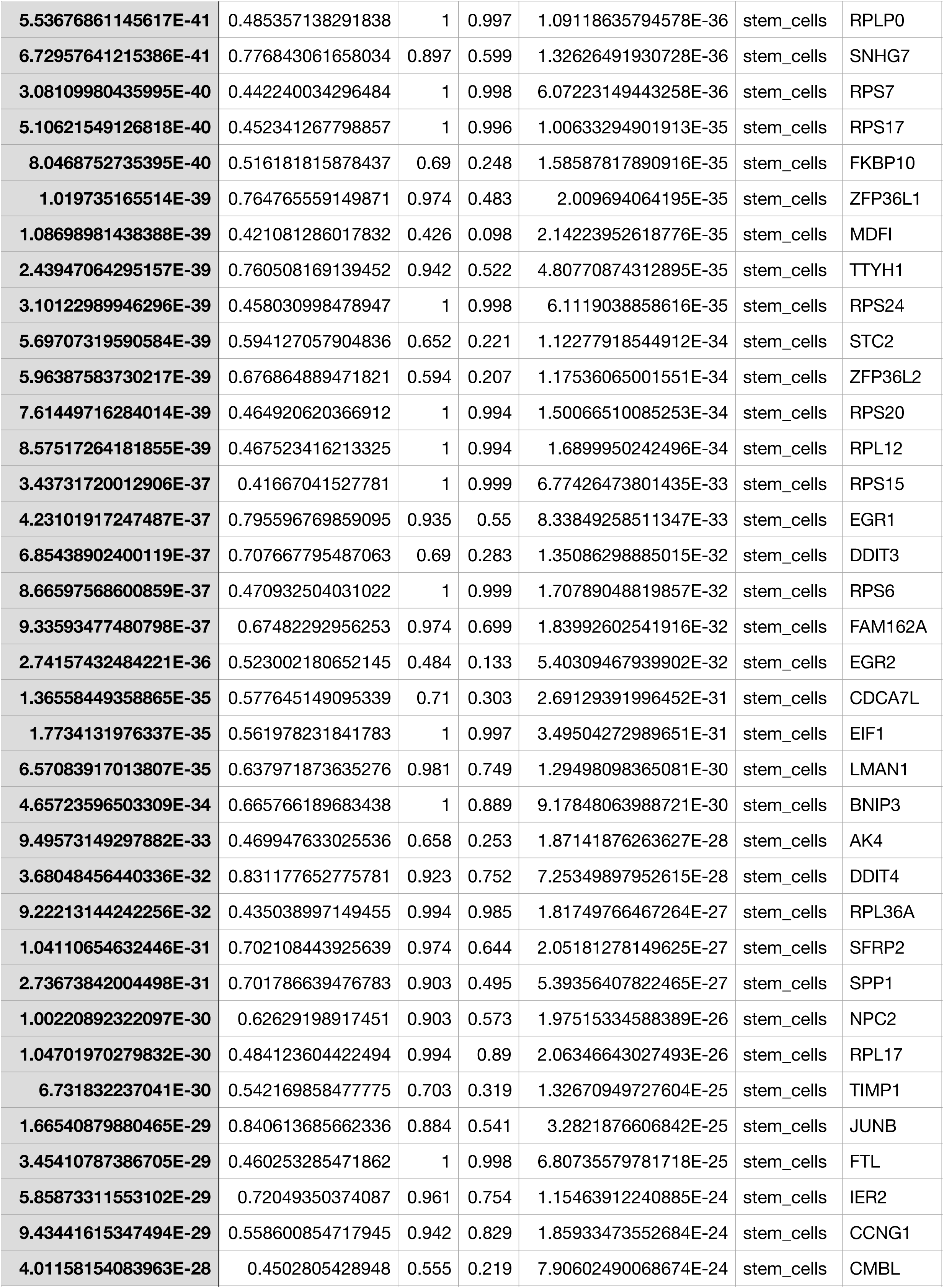
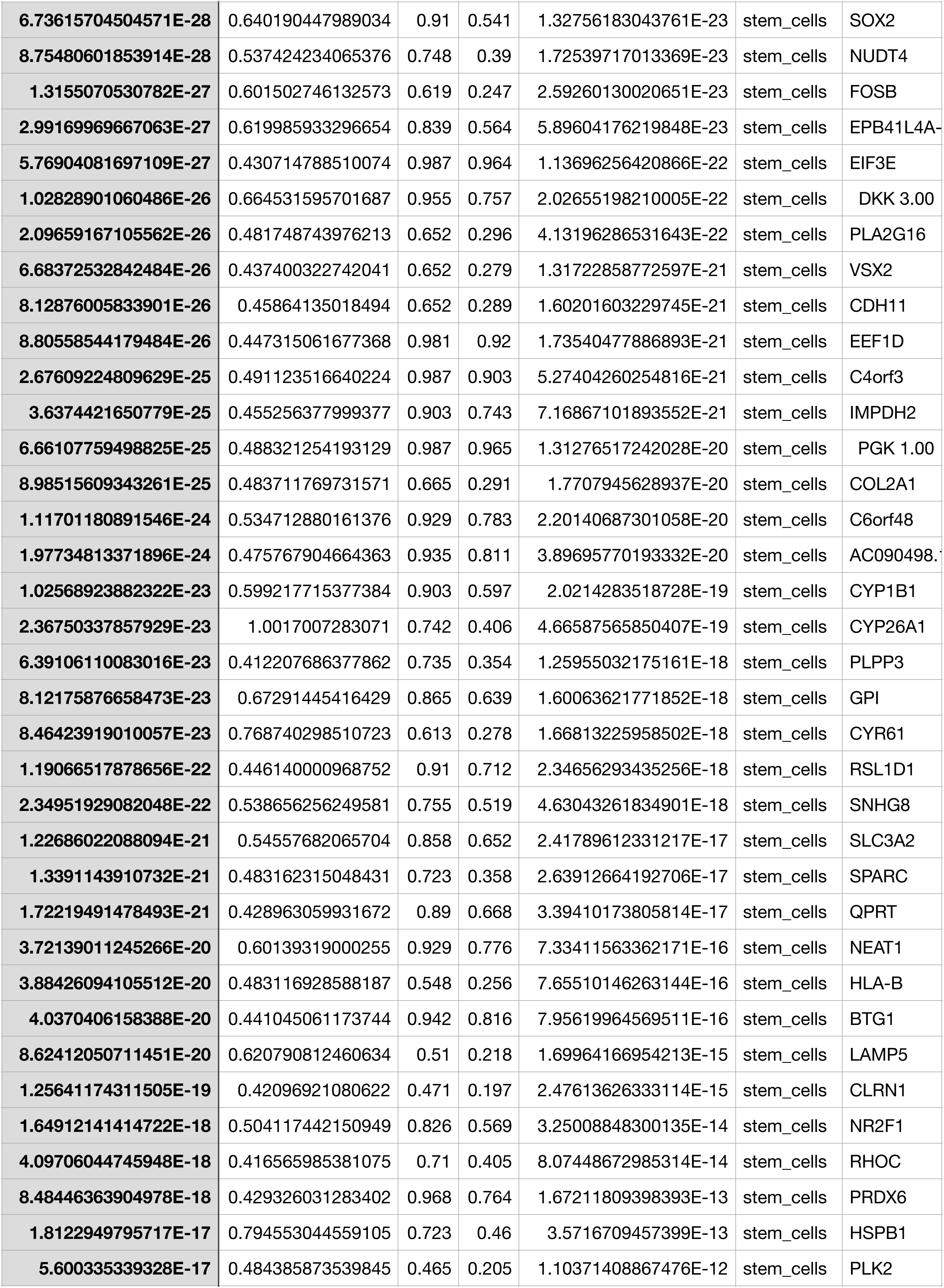
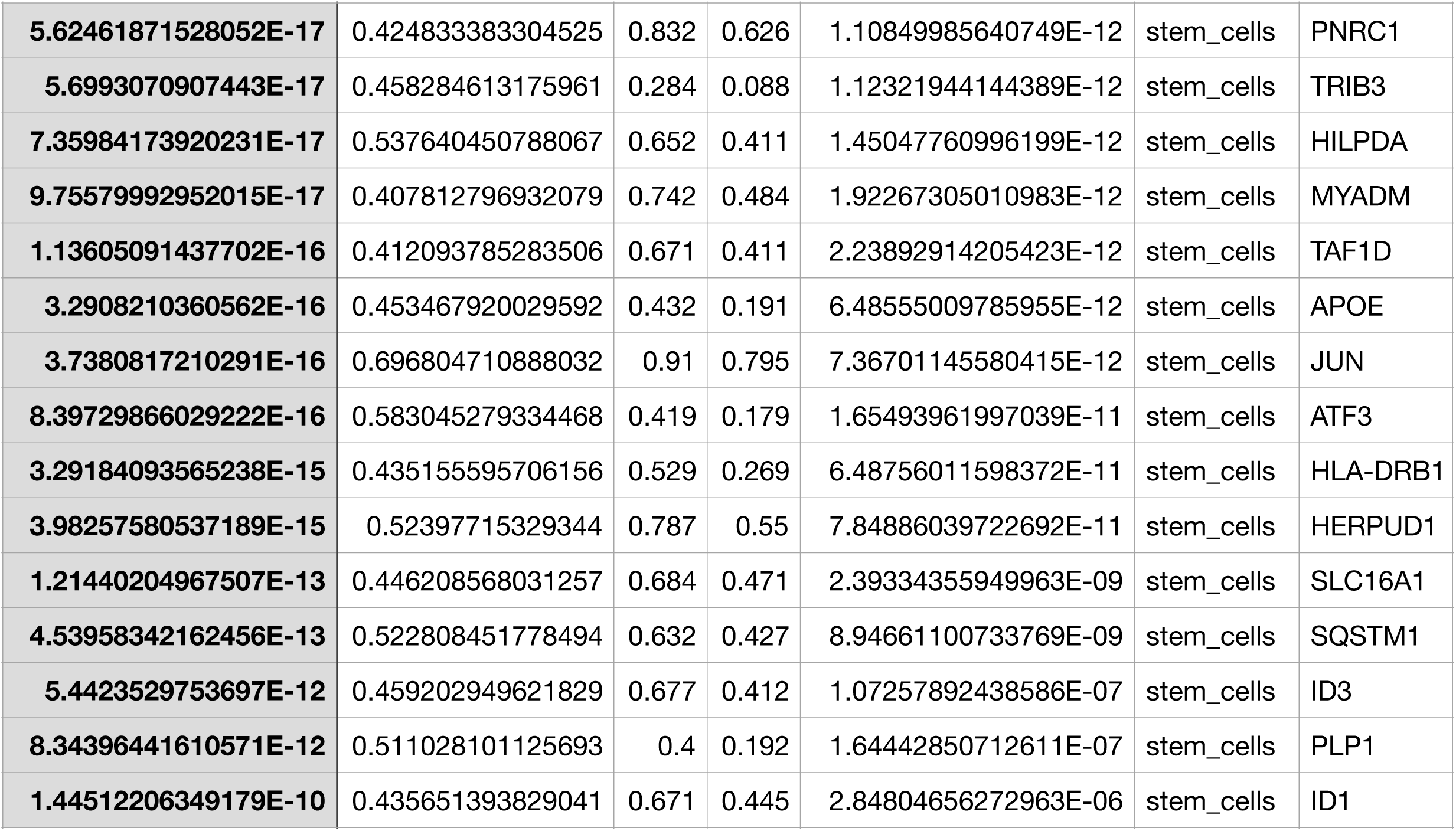
enriched_genes NEP

**SupTable 4.** GFP DEG10

| p_val | avg_logFC | pct.1 | pct.2 | p_val_adj | cluster | gene |
| --- | --- | --- | --- | --- | --- | --- |
| 7.9755606001145E-273 | 1.40687509154545 | 0.994 | 0.774 | 1.52412963068188E-268 | progenitors | SFRP2 |
| 2.49792717412938E-215 | 1.10792614355096 | 0.979 | 0.679 | 4.77353882976125E-211 | progenitors | CCND1 |
| 1.85787725777464E-183 | 1.02937240932241 | 0.917 | 0.527 | 3.55040343960734E-179 | progenitors | IFITM3 |
| 7.4066933197728E-179 | 0.819761985581814 | 0.976 | 0.648 | 1.41541909340858E-174 | progenitors | ZFP36L1 |
| 9.07551000912606E-155 | 0.881976464884904 | 0.794 | 0.328 | 1.73432996274399E-150 | progenitors | IFITM2 |
| 5.76141481742139E-154 | 0.799704968799598 | 0.94 | 0.567 | 1.10100637160923E-149 | progenitors | SPP1 |
| 6.64350211498513E-148 | 0.840515644141872 | 0.996 | 0.969 | 1.26957325417366E-143 | progenitors | TUBA1B |
| 4.98192161231705E-75 | 0.875412079859905 | 0.733 | 0.414 | 9.52045220113788E-71 | progenitors | ID3 |
| 3.7046619019519E-64 | 0.798452198708058 | 0.521 | 0.202 | 7.07960889463007E-60 | progenitors | TOP2A |
| 1.85546245477534E-31 | 0.814642413278677 | 0.935 | 0.873 | 3.54578875107567E-27 | progenitors | HIST1H4C |
| 7.2954676474322E-155 | 1.59801553415597 | 0.866 | 0.05 | 1.39416386742429E-150 | exiting_progenitors | VSX1 |
| 1.03911716785467E-64 | 2.12932606929404 | 0.896 | 0.173 | 1.98575290777027E-60 | exiting_progenitors | NEUROD1 |
| 1.24149239692652E-47 | 1.40548563638122 | 0.985 | 0.295 | 2.37249197052657E-43 | exiting_progenitors | GADD45A |
| 1.46083426287943E-43 | 2.63111846143287 | 1 | 0.588 | 2.79165427636259E-39 | exiting_progenitors | HES6 |
| 4.82477763184945E-39 | 1.56099638527471 | 1 | 0.999 | 9.2201500544643E-35 | exiting_progenitors | CKB |
| 1.08268544410143E-38 | 1.41338645627875 | 0.985 | 0.386 | 2.06901188367782E-34 | exiting_progenitors | TOP2A |
| 1.41196897993069E-34 | 1.47472975445024 | 1 | 0.82 | 2.69827272064755E-30 | exiting_progenitors | HMGB2 |
| 6.04793414891691E-34 | 1.62890152812534 | 0.925 | 0.393 | 1.15576021585802E-29 | exiting_progenitors | UBE2C |
| 1.2588052172725E-31 | 1.47249990993975 | 0.94 | 0.46 | 2.40557677020775E-27 | exiting_progenitors | UBE2S |
| 2.45762647695137E-19 | 1.52401547361851 | 1 | 0.909 | 4.69652419745406E-15 | exiting_progenitors | HIST1H4C |
| 1.58465566595118E-106 | 1.23781838970503 | 0.781 | 0.136 | 3.02827697763271E-102 | neuroblasts | RASD1 |
| 2.07794986521511E-86 | 1.76935434095007 | 0.907 | 0.315 | 3.97096219242608E-82 | neuroblasts | RGS16 |
| 5.83735191721759E-83 | 1.80571880822077 | 0.861 | 0.28 | 1.11551795138028E-78 | neuroblasts | GADD45A |
| 1.36643906209204E-71 | 1.45433707165045 | 0.993 | 0.805 | 2.61126504765789E-67 | neuroblasts | SH3BGRL3 |
| 1.95708713178716E-71 | 1.03370974860396 | 1 | 0.992 | 3.73999350884525E-67 | neuroblasts | RBP1 |
| 9.30910987572058E-66 | 1.86753029882987 | 0.967 | 0.577 | 1.7789708972502E-61 | neuroblasts | HES6 |
| 4.03321739470783E-65 | 1.6520567018224 | 0.695 | 0.195 | 7.70747844128665E-61 | neuroblasts | RP3-395M20.12 |
| 1.00948029377479E-52 | 2.70905334751122 | 0.596 | 0.163 | 1.92911684140362E-48 | neuroblasts | GAL |
| 8.11922275965311E-46 | 1.39163024172413 | 0.894 | 0.589 | 1.55158346936971E-41 | neuroblasts | TRH |
| 1.3582387027706E-34 | 1.21826979255327 | 0.384 | 0.089 | 2.59559416099461E-30 | neuroblasts | CLDN5 |
| 0 | 2.53809497444755 | 0.891 | 0.016 | 0 | photoreceptors | DCT |
| 0 | 1.79281106208741 | 0.832 | 0.018 | 0 | photoreceptors | PRDM1 |
| 1.79493797776546E-196 | 1.98740664572594 | 0.653 | 0.024 | 3.4301264755098E-192 | photoreceptors | RCVRN |
| 6.4160580545524E-183 | 1.75792736202719 | 0.792 | 0.049 | 1.22610869422496E-178 | photoreceptors | GNB3 |
| 1.03472421379948E-112 | 1.62778005261522 | 0.871 | 0.125 | 1.9773579725708E-108 | photoreceptors | SEPT4 |
| 9.85220748449355E-109 | 1.98729829403436 | 0.95 | 0.161 | 1.88275685028672E-104 | photoreceptors | NEUROD1 |
| 5.41587624456727E-93 | 1.97736147740527 | 0.931 | 0.216 | 1.0349739503368E-88 | photoreceptors | PHLDA1 |
| 2.98941308612055E-92 | 1.66753426505955 | 0.812 | 0.126 | 5.71276840757638E-88 | photoreceptors | SSTR2 |
| **2.79256254814977E-90** | 2.32822297749944 | 1 | 0.301 | 5.33658702951421E-86 | photoreceptors | FAM57B |
| **2.79670025374746E-65** | 1.84728491349718 | 1 | 0.537 | 5.3444941849114E-61 | photoreceptors | PCBP4 |
| **6.83826479193271E-283** | 1.74475653920269 | 0.876 | 0.063 | 1.30679240173834E-278 | rgc | NEFM |
| **1.21117925290698E-253** | 1.60314977443565 | 0.742 | 0.043 | 2.31456355230524E-249 | rgc | ELAVL4 |
| **4.21051803814329E-197** | 1.61871436608795 | 0.732 | 0.07 | 8.04629997089184E-193 | rgc | NEFL |
| **4.54528126733512E-169** | 1.86078476725769 | 0.919 | 0.178 | 8.68603250187741E-165 | rgc | SNCG |
| **6.25681677417204E-157** | 2.39364505699298 | 0.852 | 0.171 | 1.19567768554428E-152 | rgc | NSG1 |
| **7.88674169471337E-143** | 2.46425886392925 | 0.952 | 0.331 | 1.50715633785973E-138 | rgc | GAP43 |
| **5.18627711382802E-142** | 1.94591534625095 | 0.603 | 0.068 | 9.91097556452535E-138 | rgc | PRPH |
| **1.27579741404468E-132** | 1.6192275686326 | 0.967 | 0.338 | 2.43804885823938E-128 | rgc | TAGLN3 |
| **1.28574486485278E-130** | 2.21079637469672 | 0.995 | 0.426 | 2.45705843673367E-126 | rgc | STMN2 |
| **4.61190659072856E-99** | 1.58325292053938 | 0.976 | 0.644 | 8.81335349488229E-95 | rgc | TUBB2A |
| **2.34856100902955E-258** | 1.9415508564384 | 0.652 | 0.011 | 4.48810008825547E-254 | hc/ac | TFAP2A |
| **4.01413646679045E-123** | 1.76249219831456 | 0.957 | 0.132 | 7.67101478803654E-119 | hc/ac | ONECUT2 |
| **3.97561951181722E-99** | 1.3497925095472 | 0.707 | 0.076 | 7.59740888708271E-95 | hc/ac | HMP19 |
| **1.20881240855296E-61** | 1.22079022460648 | 0.63 | 0.099 | 2.31004051274471E-57 | hc/ac | ONECUT1 |
| **4.59065478460819E-60** | 2.12190030735959 | 1 | 0.451 | 8.77274129338626E-56 | hc/ac | STMN2 |
| **2.15654880866339E-52** | 1.73862906000784 | 1 | 0.977 | 4.12116477335574E-48 | hc/ac | SOX4 |
| **3.08417096870498E-45** | 1.21129892303644 | 1 | 0.876 | 5.89385072119522E-41 | hc/ac | MLLT11 |
| **1.0126107081594E-36** | 1.22880326073114 | 0.967 | 0.848 | 1.93509906329262E-32 | hc/ac | AP1S2 |
| **2.81432353183936E-32** | 1.20097554358389 | 0.924 | 0.587 | 5.37817226934501E-28 | hc/ac | ID2 |
| **4.6942112596226E-32** | 1.27061487486526 | 0.837 | 0.419 | 8.9706377171388E-28 | hc/ac | STMN4 |
| **5.23425939472817E-151** | 1.14155452016634 | 0.995 | 0.915 | 1.00026697033255E-146 | stem_cells | SLC2A1 |
| **1.87761454988888E-140** | 1.08937888613719 | 0.834 | 0.299 | 3.58812140483764E-136 | stem_cells | STC2 |
| **2.77590027316862E-136** | 1.18549703280372 | 0.968 | 0.789 | 5.30474542202523E-132 | stem_cells | GPI |
| **1.39749757548281E-129** | 0.956761975856372 | 0.91 | 0.461 | 2.67061786674765E-125 | stem_cells | INSIG2 |
| **8.21286962102296E-124** | 1.14328670467328 | 0.985 | 0.768 | 1.56947938457749E-119 | stem_cells | DDIT4 |
| **8.63568118614085E-124** | 0.933003209699659 | 0.971 | 0.694 | 1.65027867467152E-119 | stem_cells | P4HA1 |
| **1.38799681730411E-120** | 0.918667371961046 | 0.787 | 0.263 | 2.65246191786815E-116 | stem_cells | NRN1 |
| **5.60507060033651E-114** | 0.958886841325971 | 0.895 | 0.507 | 1.07112899172431E-109 | stem_cells | PLOD2 |
| **1.43866432066906E-95** | 1.0879783356735 | 0.861 | 0.559 | 2.74928751679858E-91 | stem_cells | HILPDA |
| **1.14892702959881E-80** | 0.938827359825493 | 0.802 | 0.476 | 2.19559955356332E-76 | stem_cells | HERPUD1 |

**SupTable 5.**
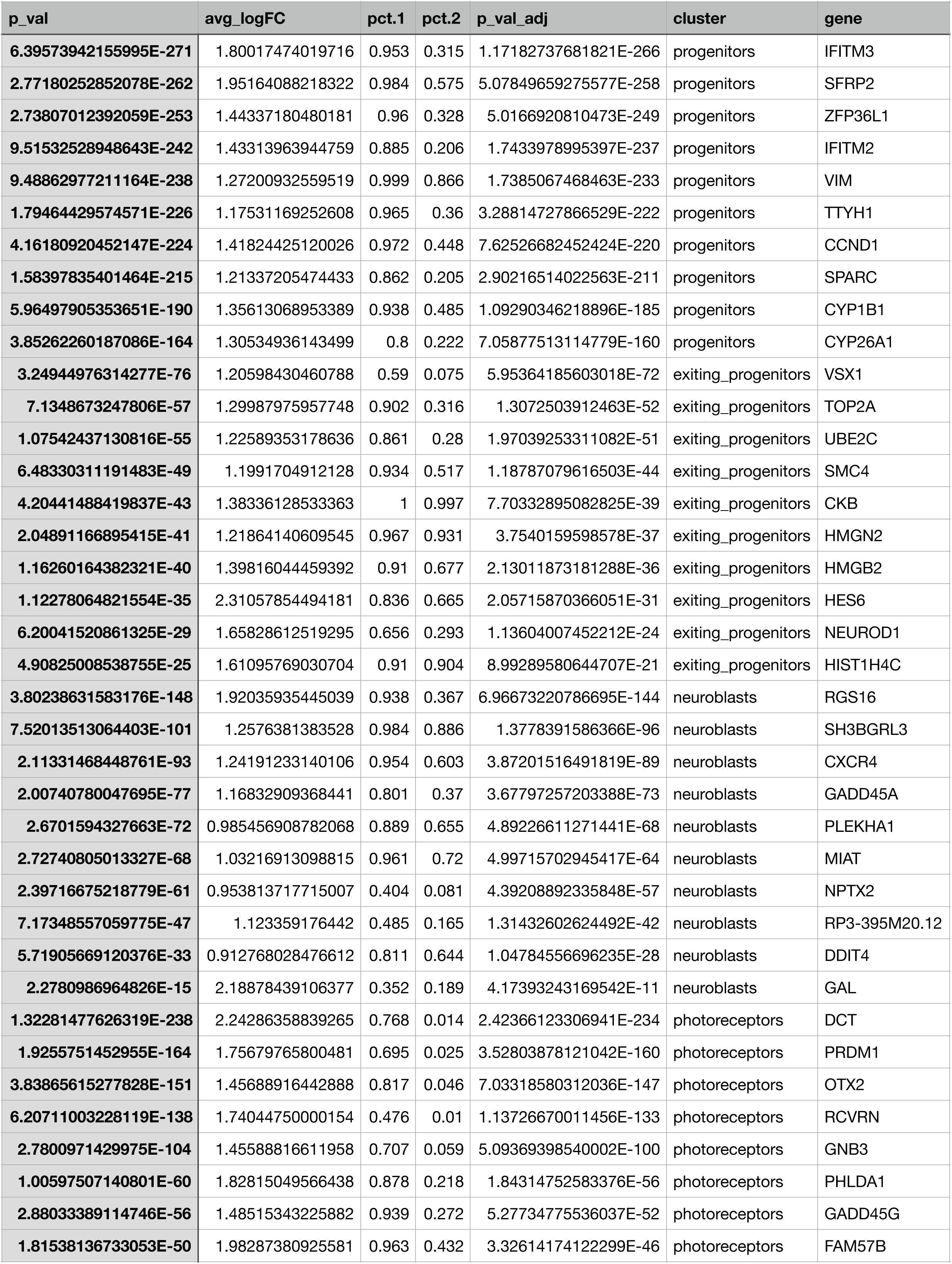
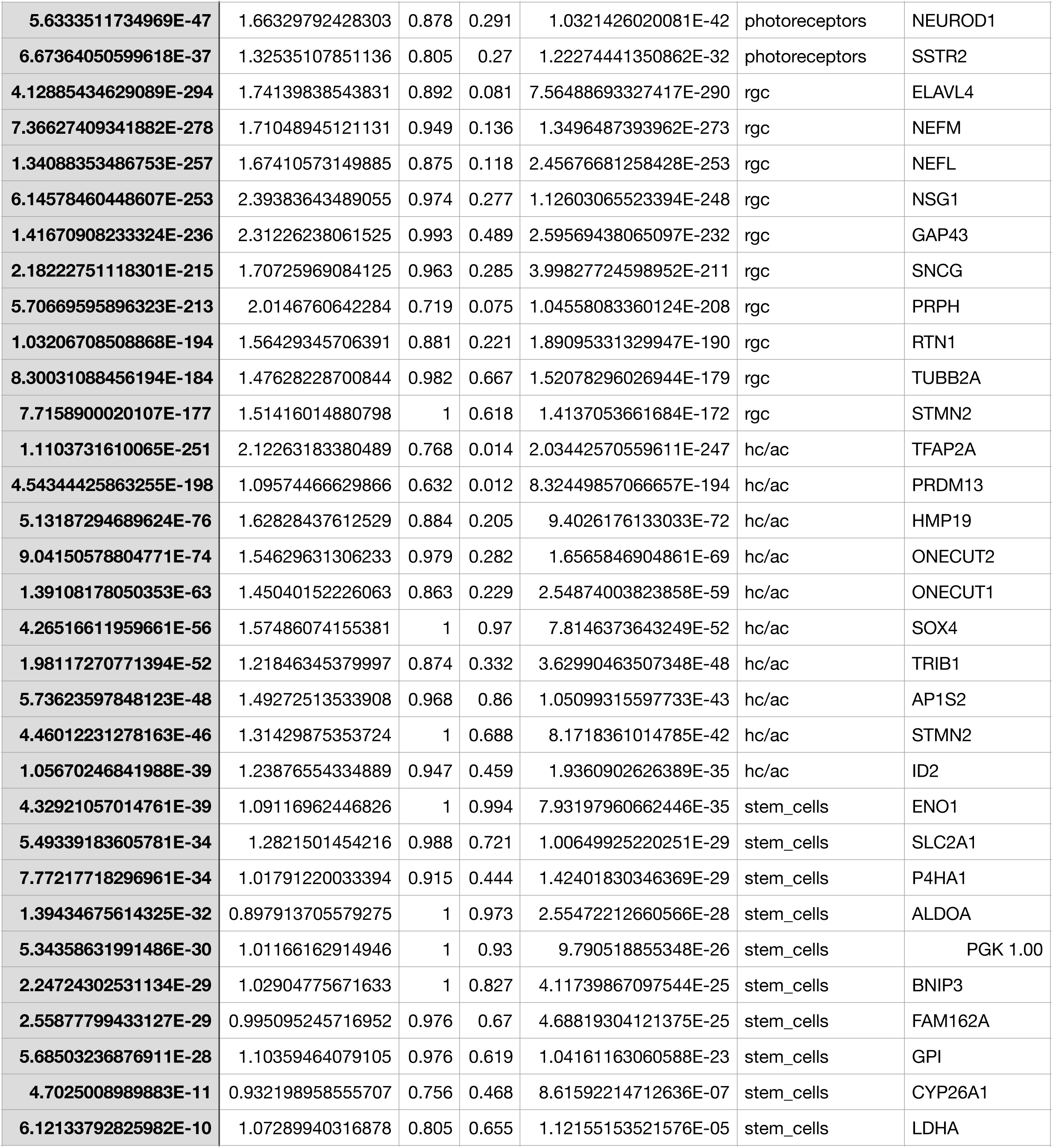
AEP DEG10

| p_val | avg_logFC | pct.1 | pct.2 | p_val_adj | cluster | gene |
| --- | --- | --- | --- | --- | --- | --- |
| 6.39573942155995E-271 | 1.80017474019716 | 0.953 | 0.315 | 1.17182737681821E-266 | progenitors | IFITM3 |
| 2.77180252852078E-262 | 1.95164088218322 | 0.984 | 0.575 | 5.07849659275577E-258 | progenitors | SFRP2 |
| 2.73807012392059E-253 | 1.44337180480181 | 0.96 | 0.328 | 5.0166920810473E-249 | progenitors | ZFP36L1 |
| 9.51532528948643E-242 | 1.43313963944759 | 0.885 | 0.206 | 1.7433978995397E-237 | progenitors | IFITM2 |
| 9.48862977211164E-238 | 1.27200932559519 | 0.999 | 0.866 | 1.7385067468463E-233 | progenitors | VIM |
| 1.79464429574571E-226 | 1.17531169252608 | 0.965 | 0.36 | 3.28814727866529E-222 | progenitors | TTYH1 |
| 4.16180920452147E-224 | 1.41824425120026 | 0.972 | 0.448 | 7.62526682452424E-220 | progenitors | CCND1 |
| 1.58397835401464E-215 | 1.21337205474433 | 0.862 | 0.205 | 2.90216514022563E-211 | progenitors | SPARC |
| 5.96497905353651E-190 | 1.35613068953389 | 0.938 | 0.485 | 1.09290346218896E-185 | progenitors | CYP1B1 |
| 3.85262260187086E-164 | 1.30534936143499 | 0.8 | 0.222 | 7.05877513114779E-160 | progenitors | CYP26A1 |
| 3.24944976314277E-76 | 1.20598430460788 | 0.59 | 0.075 | 5.95364185603018E-72 | exiting_progenitors | VSX1 |
| 7.1348673247806E-57 | 1.29987975957748 | 0.902 | 0.316 | 1.3072503912463E-52 | exiting_progenitors | TOP2A |
| 1.07542437130816E-55 | 1.22589353178636 | 0.861 | 0.28 | 1.97039253311082E-51 | exiting_progenitors | UBE2C |
| 6.48330311191483E-49 | 1.1991704912128 | 0.934 | 0.517 | 1.18787079616503E-44 | exiting_progenitors | SMC4 |
| 4.20441488419837E-43 | 1.38336128533363 | 1 | 0.997 | 7.70332895082825E-39 | exiting_progenitors | CKB |
| 2.04891166895415E-41 | 1.21864140609545 | 0.967 | 0.931 | 3.7540159598578E-37 | exiting_progenitors | HMGN2 |
| 1.16260164382321E-40 | 1.39816044459392 | 0.91 | 0.677 | 2.13011873181288E-36 | exiting_progenitors | HMGB2 |
| 1.12278064821554E-35 | 2.31057854494181 | 0.836 | 0.665 | 2.05715870366051E-31 | exiting_progenitors | HES6 |
| 6.20041520861325E-29 | 1.65828612519295 | 0.656 | 0.293 | 1.13604007452212E-24 | exiting_progenitors | NEUROD1 |
| 4.90825008538755E-25 | 1.61095769030704 | 0.91 | 0.904 | 8.99289580644707E-21 | exiting_progenitors | HIST1H4C |
| 3.80238631583176E-148 | 1.92035935445039 | 0.938 | 0.367 | 6.96673220786695E-144 | neuroblasts | RGS16 |
| 7.52013513064403E-101 | 1.2576381383528 | 0.984 | 0.886 | 1.3778391586366E-96 | neuroblasts | SH3BGRL3 |
| 2.11331468448761E-93 | 1.24191233140106 | 0.954 | 0.603 | 3.87201516491819E-89 | neuroblasts | CXCR4 |
| 2.00740780047695E-77 | 1.16832909368441 | 0.801 | 0.37 | 3.67797257203388E-73 | neuroblasts | GADD45A |
| 2.6701594327663E-72 | 0.985456908782068 | 0.889 | 0.655 | 4.89226611271441E-68 | neuroblasts | PLEKHA1 |
| 2.72740805013327E-68 | 1.03216913098815 | 0.961 | 0.72 | 4.99715702945417E-64 | neuroblasts | MIAT |
| 2.39716675218779E-61 | 0.953813717715007 | 0.404 | 0.081 | 4.39208892335848E-57 | neuroblasts | NPTX2 |
| 7.17348557059775E-47 | 1.123359176442 | 0.485 | 0.165 | 1.31432602624492E-42 | neuroblasts | RP3-395M20.12 |
| 5.71905669120376E-33 | 0.912768028476612 | 0.811 | 0.644 | 1.04784556696235E-28 | neuroblasts | DDIT4 |
| 2.2780986964826E-15 | 2.18878439106377 | 0.352 | 0.189 | 4.17393243169542E-11 | neuroblasts | GAL |
| 1.32281477626319E-238 | 2.24286358839265 | 0.768 | 0.014 | 2.42366123306941E-234 | photoreceptors | DCT |
| 1.9255751452955E-164 | 1.75679765800481 | 0.695 | 0.025 | 3.52803878121042E-160 | photoreceptors | PRDM1 |
| 3.83865615277828E-151 | 1.45688916442888 | 0.817 | 0.046 | 7.03318580312036E-147 | photoreceptors | OTX2 |
| 6.20711003228119E-138 | 1.74044750000154 | 0.476 | 0.01 | 1.13726670011456E-133 | photoreceptors | RCVRN |
| 2.7800971429975E-104 | 1.45588816611958 | 0.707 | 0.059 | 5.09369398540002E-100 | photoreceptors | GNB3 |
| 1.00597507140801E-60 | 1.82815049566438 | 0.878 | 0.218 | 1.84314752583376E-56 | photoreceptors | PHLDA1 |
| 2.88033389114746E-56 | 1.48515343225882 | 0.939 | 0.272 | 5.27734775536037E-52 | photoreceptors | GADD45G |
| 1.81538136733053E-50 | 1.98287380925581 | 0.963 | 0.432 | 3.32614174122299E-46 | photoreceptors | FAM57B |
| 5.6333511734969E-47 | 1.66329792428303 | 0.878 | 0.291 | 1.0321426020081E-42 | photoreceptors | NEUROD1 |
| 6.67364050599618E-37 | 1.32535107851136 | 0.805 | 0.27 | 1.22274441350862E-32 | photoreceptors | SSTR2 |
| 4.12885434629089E-294 | 1.74139838543831 | 0.892 | 0.081 | 7.56488693327417E-290 | rgc | ELAVL4 |
| 7.36627409341882E-278 | 1.71048945121131 | 0.949 | 0.136 | 1.3496487393962E-273 | rgc | NEFM |
| 1.34088353486753E-257 | 1.67410573149885 | 0.875 | 0.118 | 2.45676681258428E-253 | rgc | NEFL |
| 6.14578460448607E-253 | 2.39383643489055 | 0.974 | 0.277 | 1.12603065523394E-248 | rgc | NSG1 |
| 1.41670908233324E-236 | 2.31226238061525 | 0.993 | 0.489 | 2.59569438065097E-232 | rgc | GAP43 |
| 2.18222751118301E-215 | 1.70725969084125 | 0.963 | 0.285 | 3.99827724598952E-211 | rgc | SNCG |
| 5.70669595896323E-213 | 2.0146760642284 | 0.719 | 0.075 | 1.04558083360124E-208 | rgc | PRPH |
| 1.03206708508868E-194 | 1.56429345706391 | 0.881 | 0.221 | 1.89095331329947E-190 | rgc | RTN1 |
| 8.30031088456194E-184 | 1.47628228700844 | 0.982 | 0.667 | 1.52078296026944E-179 | rgc | TUBB2A |
| 7.7158900020107E-177 | 1.51416014880798 | 1 | 0.618 | 1.4137053661684E-172 | rgc | STMN2 |
| 1.1103731610065E-251 | 2.12263183380489 | 0.768 | 0.014 | 2.03442570559611E-247 | hc/ac | TFAP2A |
| 4.54344425863255E-198 | 1.09574466629866 | 0.632 | 0.012 | 8.32449857066657E-194 | hc/ac | PRDM13 |
| 5.13187294689624E-76 | 1.62828437612529 | 0.884 | 0.205 | 9.4026176133033E-72 | hc/ac | HMP19 |
| 9.04150578804771E-74 | 1.54629631306233 | 0.979 | 0.282 | 1.6565846904861E-69 | hc/ac | ONECUT2 |
| 1.39108178050353E-63 | 1.45040152226063 | 0.863 | 0.229 | 2.54874003823858E-59 | hc/ac | ONECUT1 |
| 4.26516611959661E-56 | 1.57486074155381 | 1 | 0.97 | 7.8146373643249E-52 | hc/ac | SOX4 |
| 1.98117270771394E-52 | 1.21846345379997 | 0.874 | 0.332 | 3.62990463507348E-48 | hc/ac | TRIB1 |
| 5.73623597848123E-48 | 1.49272513533908 | 0.968 | 0.86 | 1.05099315597733E-43 | hc/ac | AP1S2 |
| 4.46012231278163E-46 | 1.31429875353724 | 1 | 0.688 | 8.1718361014785E-42 | hc/ac | STMN2 |
| 1.05670246841988E-39 | 1.23876554334889 | 0.947 | 0.459 | 1.9360902626389E-35 | hc/ac | ID2 |
| 4.32921057014761E-39 | 1.09116962446826 | 1 | 0.994 | 7.93197960662446E-35 | stem_cells | ENO1 |
| 5.49339183605781E-34 | 1.2821501454216 | 0.988 | 0.721 | 1.00649925220251E-29 | stem_cells | SLC2A1 |
| 7.77217718296961E-34 | 1.01791220033394 | 0.915 | 0.444 | 1.42401830346369E-29 | stem_cells | P4HA1 |
| 1.39434675614325E-32 | 0.897913705579275 | 1 | 0.973 | 2.55472212660566E-28 | stem_cells | ALDOA |
| 5.34358631991486E-30 | 1.01166162914946 | 1 | 0.93 | 9.790518855348E-26 | stem_cells | PGK 1.00 |
| 2.24724302531134E-29 | 1.02904775671633 | 1 | 0.827 | 4.11739867097544E-25 | stem_cells | BNIP3 |
| 2.55877799433127E-29 | 0.995095245716952 | 0.976 | 0.67 | 4.68819304121375E-25 | stem_cells | FAM162A |
| 5.68503236876911E-28 | 1.10359464079105 | 0.976 | 0.619 | 1.04161163060588E-23 | stem_cells | GPI |
| 4.7025008989883E-11 | 0.932198958555707 | 0.756 | 0.468 | 8.61592214712636E-07 | stem_cells | CYP26A1 |
| 6.12133792825982E-10 | 1.07289940316878 | 0.805 | 0.655 | 1.12155153521576E-05 | stem_cells | LDHA |

**SupTable 6.** NEP DEG10

| p_val | avg_logFC | pct.1 | pct.2 | p_val_adj | cluster | gene |
| --- | --- | --- | --- | --- | --- | --- |
| 0 | 2.15040505118418 | 0.999 | 0.526 | 0 | progenitors | SFRP2 |
| 0 | 1.84620823017618 | 0.853 | 0.252 | 0 | progenitors | CYP26A1 |
| 0 | 1.7779501569875 | 0.996 | 0.312 | 0 | progenitors | ZFP36L1 |
| 0 | 1.69432772775035 | 0.962 | 0.278 | 0 | progenitors | IFITM3 |
| 0 | 1.58805350397233 | 0.991 | 0.412 | 0 | progenitors | CCND1 |
| 0 | 1.5118992094342 | 1 | 0.769 | 0 | progenitors | VIM |
| 0 | 1.51158747141804 | 0.95 | 0.478 | 0 | progenitors | CYP1B1 |
| 0 | 1.50081898363016 | 0.886 | 0.111 | 0 | progenitors | HES1 |
| 0 | 1.43894021706406 | 0.987 | 0.366 | 0 | progenitors | TTYH1 |
| 1.0584002645718E-208 | 1.47309084340855 | 0.776 | 0.286 | 2.08589524141811E-204 | progenitors | ID3 |
| 9.39016262972125E-122 | 1.5665779818246 | 0.91 | 0.233 | 1.85061325106546E-117 | exiting_progenitors | NUSAP1 |
| 5.89479884790682E-113 | 1.66641195406698 | 0.938 | 0.268 | 1.16174695694548E-108 | exiting_progenitors | TOP2A |
| 1.29350913432114E-98 | 1.67066088043172 | 0.91 | 0.27 | 2.5492478019201E-94 | exiting_progenitors | CENPF |
| 1.086617967745E-93 | 1.89557367559608 | 0.994 | 0.623 | 2.14150669083185E-89 | exiting_progenitors | HMGB2 |
| 2.50334683364065E-91 | 1.405817569228 | 0.96 | 0.422 | 4.93359593973899E-87 | exiting_progenitors | CKS1B |
| 1.38952887466771E-88 | 1.39020620894491 | 0.994 | 0.903 | 2.73848350619513E-84 | exiting_progenitors | HMGN2 |
| 7.82767250167975E-83 | 1.85265267079734 | 1 | 0.794 | 1.54267769663105E-78 | exiting_progenitors | HES6 |
| 1.97289656102834E-79 | 1.85868819317848 | 0.876 | 0.344 | 3.88818454247465E-75 | exiting_progenitors | UBE2C |
| 7.0043335639928E-69 | 1.48098372730913 | 0.904 | 0.419 | 1.3804140587917E-64 | exiting_progenitors | PTTG1 |
| 4.84999865374688E-42 | 1.78719991772938 | 0.944 | 0.735 | 9.55837734680436E-38 | exiting_progenitors | HIST1H4C |
| 1.16930187145625E-188 | 0.966624036001392 | 0.985 | 0.957 | 2.30446012826598E-184 | neuroblasts | SH3BGRL3 |
| 2.30946768397378E-182 | 1.15929408515216 | 0.956 | 0.733 | 4.55149891157552E-178 | neuroblasts | CXCR4 |
| 7.04199761050279E-179 | 0.955966843545263 | 0.984 | 0.837 | 1.38783688907789E-174 | neuroblasts | MIAT |
| 1.44450092859416E-135 | 1.01847166776057 | 0.849 | 0.536 | 2.84682243007337E-131 | neuroblasts | GADD45A |
| 9.45647113032676E-101 | 1.06573669165747 | 0.765 | 0.507 | 1.8636813303648E-96 | neuroblasts | RGS16 |
| 1.70931757053462E-74 | 0.813698539051603 | 0.852 | 0.734 | 3.36872306800962E-70 | neuroblasts | DDIT4 |
| 4.89795774625378E-61 | 0.946739111301297 | 0.864 | 0.788 | 9.65289512631695E-57 | neuroblasts | HES6 |
| 1.12755056010244E-33 | 1.00097391371338 | 0.405 | 0.254 | 2.22217664384989E-29 | neuroblasts | RP3-395M20.12 |
| 2.11577992763031E-21 | 0.700450123851411 | 0.391 | 0.252 | 4.16977908137382E-17 | neuroblasts | IGFBP5 |
| 1.02459238475717E-07 | 1.5069924112315 | 0.287 | 0.233 | 0.00201926667187942 | neuroblasts | GAL |
| 0 | 2.42896721596791 | 0.832 | 0.041 | 0 | photoreceptors | DCT |
| 0 | 1.42077576520909 | 0.864 | 0.096 | 0 | photoreceptors | OTX2 |
| 0 | 1.39708500678511 | 0.586 | 0.026 | 0 | photoreceptors | PRDM1 |
| 2.27127159864859E-285 | 1.72244069569183 | 0.801 | 0.115 | 4.47622206661664E-281 | photoreceptors | GNB3 |
| 4.08309335937793E-217 | 1.7301652160062 | 0.72 | 0.117 | 8.04696039266203E-213 | photoreceptors | C8orf46 |
| 8.37752023354786E-212 | 1.48042568102138 | 0.895 | 0.271 | 1.65104168762761E-207 | photoreceptors | SEPT4 |
| 7.21606404542176E-198 | 1.82800223735372 | 0.552 | 0.065 | 1.42214190207172E-193 | photoreceptors | RCVRN |
| 3.77906501604816E-187 | 1.58681037681391 | 0.987 | 0.673 | 7.44778133362771E-183 | photoreceptors | FAM57B |
| **1.57294965357472E-176** | 1.51786999689233 | 0.869 | 0.296 | 3.09996917726505E-172 | photoreceptors | PHLDA1 |
| **9.21728351610386E-25** | 1.48935056516302 | 0.545 | 0.347 | 1.81654223535375E-20 | photoreceptors | CRABP2 |
| **0** | 1.67866507543497 | 0.985 | 0.59 | 0 | rgc | GAP43 |
| **0** | 1.62664244202391 | 0.915 | 0.36 | 0 | rgc | NSG1 |
| **0** | 1.40049703856269 | 0.937 | 0.429 | 0 | rgc | RTN1 |
| **0** | 1.17520642487855 | 0.907 | 0.292 | 0 | rgc | ELAVL4 |
| **0** | 1.14672627900499 | 0.989 | 0.791 | 0 | rgc | TUBB2A |
| **0** | 1.11760455903467 | 0.998 | 0.827 | 0 | rgc | STMN2 |
| **1.87768401855086E-243** | 1.12692275958606 | 0.743 | 0.214 | 3.70053966376003E-239 | rgc | NEFM |
| **1.30313699795126E-232** | 1.1381549245492 | 0.711 | 0.208 | 2.56822239556234E-228 | rgc | NEFL |
| **1.11683555681864E-194** | 1.2404999457904 | 0.788 | 0.385 | 2.20105951537818E-190 | rgc | SNCG |
| **2.16620330274188E-139** | 1.36607779037133 | 0.461 | 0.122 | 4.2691534690437E-135 | rgc | PRPH |
| **1.43951936272422E-250** | 1.78941831362287 | 0.619 | 0.021 | 2.83700476005688E-246 | hc/ac | TFAP2A |
| **1.07916254507875E-179** | 1.16821826326153 | 0.571 | 0.03 | 2.1268135438412E-175 | hc/ac | PRDM13 |
| **2.8293086949641E-61** | 1.16167923360779 | 0.944 | 0.44 | 5.57600157603524E-57 | hc/ac | ONECUT2 |
| **1.16228379969261E-60** | 1.07143760136788 | 1 | 0.988 | 2.2906289124342E-56 | hc/ac | SOX4 |
| **1.57431637321431E-47** | 1.43905447292851 | 0.841 | 0.39 | 3.10266270833077E-43 | hc/ac | ID2 |
| **1.21972883458479E-46** | 0.971662459931208 | 0.778 | 0.289 | 2.40384158719971E-42 | hc/ac | LINC00599 |
| **1.65626261443858E-46** | 0.930895015971393 | 0.468 | 0.098 | 3.26416236053555E-42 | hc/ac | PROX1 |
| **7.29033246405269E-46** | 1.03334028832232 | 0.873 | 0.41 | 1.4367787220155E-41 | hc/ac | HMP19 |
| **9.2672618918328E-46** | 1.14788890128705 | 0.968 | 0.874 | 1.82639197364241E-41 | hc/ac | AP1S2 |
| **9.68404134892103E-35** | 0.984539251362921 | 0.667 | 0.272 | 1.90853086904536E-30 | hc/ac | ONECUT1 |
| **1.68248737987099E-66** | 1.39239449977265 | 0.981 | 0.754 | 3.31584612824975E-62 | stem_cells | SLC2A1 |
| **1.00044477856205E-59** | 1.21296465076215 | 1 | 0.826 | 1.97167656959009E-55 | stem_cells | VIM |
| **1.19158351570964E-59** | 0.860952117950643 | 0.535 | 0.109 | 2.34837279276057E-55 | stem_cells | ADM |
| **4.99714949212476E-52** | 0.804020203681976 | 0.742 | 0.271 | 9.84838221907948E-48 | stem_cells | PLOD2 |
| **5.27807021007884E-52** | 0.864377336267267 | 0.871 | 0.432 | 1.04020207700234E-47 | stem_cells | P4HA1 |
| **4.77834170505954E-44** | 1.0873553747212 | 0.961 | 0.646 | 9.41715583233135E-40 | stem_cells | FOS |
| **4.23101917247487E-37** | 0.795596769859095 | 0.935 | 0.55 | 8.33849258511347E-33 | stem_cells | EGR1 |
| **3.68048456440336E-32** | 0.831177652775781 | 0.923 | 0.752 | 7.25349897952615E-28 | stem_cells | DDIT4 |
| **1.66540879880465E-29** | 0.840613685662336 | 0.884 | 0.541 | 3.2821876606842E-25 | stem_cells | JUNB |
| **2.36750337857929E-23** | 1.0017007283071 | 0.742 | 0.406 | 4.66587565850407E-19 | stem_cells | CYP26A1 |

## Notes

### Competing Interest Statement

The authors have declared no competing interest.

### Summary of Updates

The revision has -changed the title; -reduced the word count for the abstract to <200; -combined supplementary data with the main manuscript.

